# Ancient gene clusters govern the initiation of monoterpenoid indole alkaloid biosynthesis and C3 stereochemistry inversion

**DOI:** 10.1101/2025.01.07.631695

**Authors:** Jaewook Hwang, Jonathan Kirshner, Daniel André Ramey Deschênes, Matthew Bailey Richardson, Steven J. Fleck, Scott Galeung Alexander Mann, Jun Guo, Jacob Owen Perley, Mohammadamin Shahsavarani, Jorge Jonathan Oswaldo Garza-Garcia, Alyssa Dawn Seveck, Savannah Sadie Doiron, Zhan Mai, Stephen Nelson Silliphant, Sarah Anne Englehart, Barry A. Blight, Larry Calhoun, Di Gao, Jiazhang Lian, Ghislain Deslongchamps, Victor A. Albert, Yang Qu

## Abstract

The inversion of C3 stereochemistry in monoterpenoid indole alkaloids (MIAs), derived from the central precursor strictosidine (3*S*), is a critical step for the biosynthesis of numerous 3*R* MIAs and spirooxindoles, including the antihypertensive drug reserpine. While early MIA biosynthesis preserves the 3*S* configuration, the mechanism underlying C3 inversion has remained unresolved. Here, we identify and biochemically characterize a conserved oxidase-reductase pair in the Gentianales order: the heteroyohimbine/yohimbine/corynanthe C3-oxidase (HYC3O) and C3-reductase (HYC3R), which together invert the 3*S* stereochemistry to 3*R* across diverse substrates. Notably, *HYC3O* and *HYC3R* reside in gene clusters in *Rauvolfia tetraphylla* and *Catharanthus roseus*, homologous to an elusive geissoschizine synthase (GS) cluster we also uncovered. In *R. tetraphylla*, these clusters are in tandem on a single chromosome, likely derived from segmental duplication, whereas in *C. roseus* they reside on separate chromosomes due to translocation. Comparative genomics indicate the GS cluster originated at the base of Gentianales (∼135 Mya), coinciding with the evolution of the strictosidine synthase cluster, while the reserpine cluster arose later in rauvolfioid Apocynaceae. Together, these findings uncover the genomic and biochemical basis for key events in MIA evolution and diversification, providing insights beyond the canonical vinblastine and ajmaline biosynthetic pathways.

## Introduction

The elucidation of the nearly complete, >30-step biosynthetic pathway for the anticancer drug vinblastine marked a milestone in monoterpenoid indole alkaloid (MIA) biosynthesis ^1–5^. This pathway, embedded within a family of over 3,000 MIAs, has provided a biochemical framework for studying other medicinally important MIAs in nature. In the model plant *Catharanthus roseus* (Madagascar’s periwinkle), MIA biosynthesis begins with enzymes encoded from a three-gene biosynthetic gene cluster (BGC) known as the strictosidine synthase (STR) BGC. This cluster encodes tryptophan decarboxylase (*TDC*), strictosidine synthase (*STR*), and Multidrug and Toxic Compound Extrusion (MATE) transporter 1 (*MATE1*) ^6,7^. Tryptamine, produced by TDC, is condensed with secologanin by STR. This pivotal reaction merges the secoiridoid and shikimate pathways. MATE1 facilitates this reaction by importing secologanin into the vacuole, where STR resides ^8,9^. Strictosidine then serves as the universal precursor for all downstream MIAs (Fig. 1) ^10–15^.

**Figure 1.**
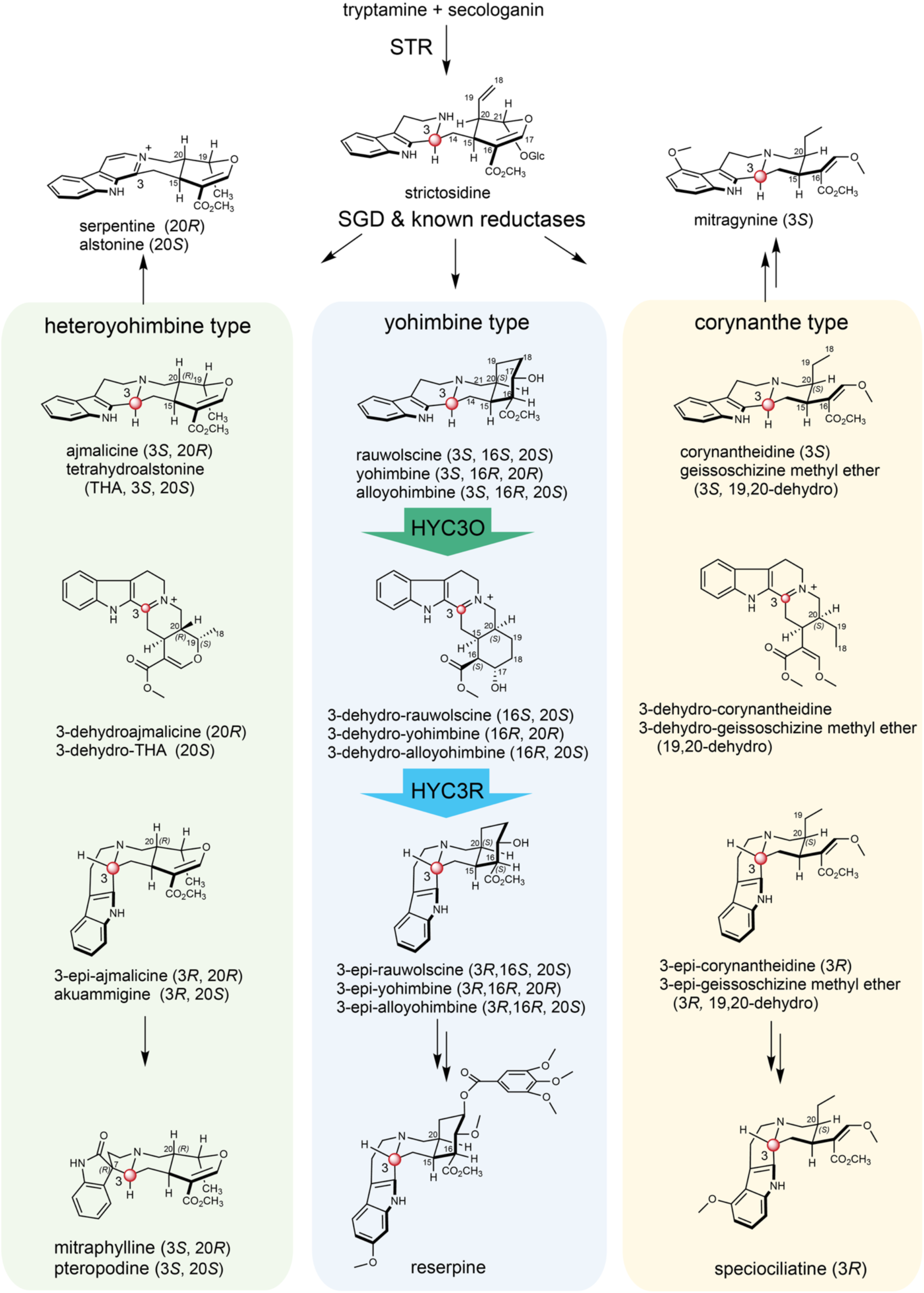
HYC3Os and HYC3Rs mediate C3 stereochemical inversion in the biosynthesis of reserpine and diverse 3*R* MIAs and their oxindole derivatives. Following the biosynthesis of strictosidine by the strictosidine synthase (STR) and its deglycosylation by the strictosidine β-glucosidase (SGD) ^10,11^, the resulting aglycones are reduced by various strictosidine aglycone reductases to form heteroyohimbine (green shade), yohimbine (blue shade), and corynanthe (orange shade) skeletons, all retaining the inherent 3*S* stereochemistry derived from strictosidine ^12–15^ . In this study, we identify conserved enzyme pairs: the heteroyohimbine/yohimbine/corynathe C3-oxidases and C3 reductases (HYC3Os and HYC3Rs) across three plant families that collaboratively catalyze C3 epimerization in diverse MIAs. This tandem reaction involves H3-hydride abstraction by HYC3Os, followed by *si*-face C3-reduction by HYC3Rs, leading to the formation of 3*R* MIAs. The indole ring of these 3*R* MIAs adopts a unique perpendicular orientation relative to the core scaffold, which is critical for their bioactivities and required for the biosynthesis of various spirooxindoles (e.g., mitraphylline) and derivatives (e.g., reserpine and speciociliatine).

The STR BGC is broadly conserved among MIA-producing species within the Gentianales order, including *Gelsemium sempervirens* (Gelsemiaceae), *C. roseus* (Apocynaceae), *Mitragyna speciosa* (kratom, Rubiaceae), and *Ophiorrhiza pumila* (Rubiaceae) ^7,16^. Phylogenomic studies suggest this cluster arose early in Gentianales evolution, likely close to the split between Gentianales and other core eudicots, ∼135 million years ago (Mya) ^17^.

Strictosidine β-glucosidase (SGD) hydrolyzes strictosidine to yield the highly labile strictosidine aglycone. This reactive intermediate is stabilized by reduction via multiple cinnamyl alcohol dehydrogenase (CAD)-like enzymes, such as yohimbine synthase (YOS), geissoschizine synthase (GS), and tetrahydroalstonine synthase (THAS), to form stable isomers that serve as scaffolds for further modification ^1–3,12–14,18–20^. Despite extensive knowledge of downstream modifications into major MIA classes such as iboga, aspidosperma, and sarpagan alkaloids, the biosynthesis of a distinct group of MIAs bearing 3*R* stereochemistry (Fig. 1), exemplified by the antihypertensive drug reserpine, has remained unresolved.

Reserpine is produced by *Rauvolfia* species such as *R. tetraphylla* (devil-pepper) and *R. serpentina* (Indian snakeroot), both members of the Apocynaceae family. Unlike 3*S* MIAs, 3*R* MIAs like reserpine exhibit a perpendicular indole orientation that profoundly alters their bioactivity and reactivity (Fig. 1). The 3*S* stereochemistry originates from the central precursor strictosidine. It has been well established by enzyme assays and feeding experiments that STRs from various species (e.g., *C. roseus*, *R. serpentina*, *M. speciosa*, *O. pumila*, and *G. sempervirens* across three plant families) exclusively produce strictosidine (3*S*) rather than its 3*R* epimer, vincoside ^10,16,21–23^. Subsequently, all characterized strictosidine aglycone reductases (e.g., GS, THAS, YOS) exclusively yield 3*S* products ^12–15^ (Fig. 1).

Beyond reserpine, 3*R* MIAs are widely distributed and pharmacologically relevant. Examples include 3-epi-THA (akuammigine) from *C. roseus* and *Picralima nitida* (akuamma; Apocynaceae) ^24,25^, reserpilline from *Ochrosia elliptica* (bloodhorn; Apocynaceae) and *Rauvolfia spp*. ^26,27^, 3-epi-mitragynine (speciociliatine) from *M. speciosa* (Rubiaceae), and the oxindoles mitraphylline, speciophylline, and corynoxine, which are biosynthesized from 3*R* intermediates in *M. speciosa*, *Mitragyna parvifolia*, *Cephalanthus occidentalis* (button bush; Rubiaceae), and *Hamelia patens* (firebush; Rubiaceae) ^14,28–31^ (Fig. 1). The widespread occurrence of 3*R* MIAs across diverse plant families suggests a significant, yet unresolved, biochemical pathway in nature.

The recent genome assembly of *R.tetraphylla* ^15,32^ provides an opportunity to investigate 3*R*-MIA biosynthesis at a genomic level. Since reserpine carries a 3*R* yohimbine-type scaffold, YOS likely represents the entry point into its biosynthesis. Intriguingly, *YOS* is located within a CAD-rich locus that contains *GS*, *THAS*, homologs of *R. serpentina* ajmaline pathway enzymes vomilenine 1,2-reductase (*VR*) and dihydrovomilenine 19,20-reductase (*DHVR*)^19^, and many uncharacterized genes, suggesting a coordinated genomic architecture for alkaloid diversification.

Here, we take a comparative genomic approach across *R. tetraphylla*, C*. roseus* and additional Gentianales species (*Asclepias syriaca*, *Gelsemium elegans, M. speciosa*, and *O. pumila*), with *Vitis vinifera* (grapevine) serving as the early-diverging core eudicot outgroup. We identify an oxidase and reductase pair heteroyohimbine/yohimbine/corynanthe C3-oxidase (HYC3O) and C3-reductase (HYC3R). Together, these enzymes catalyze stereochemical inversion at C3 by oxidizing 3*S* MIAs to iminium intermediates (HYC3O), which are then reduced by HYC3R to yield 3*R* products. Enzyme assays across five additional Gentianales species reveal that HYC3O/HYC3R represent a conserved mechanism for 3*R* MIA formation.

In *R. tetraphylla* and *C. roseus*, HYC3O and HYC3R co-occur within a BGC, which we designate the reserpine BGC. Phylogenomic analysis further uncovers a related cluster, which we term the geissoschizine synthase (GS) BGC. Our results indicate that these clusters originate from a large segmental duplication of a CAD-rich ancestral block that predates the Gentianales crown group and is syntenic with *Vitis*. The GS BGC represents the ancestral function, supporting corynanthe biosynthesis as early as the diversification of Rubiaceae (∼118 Mya) ^17^, whereas the reserpine BGC represents a later innovation in the rauvolfioid lineage. Together with a comprehensive synteny analysis of the STR BGC, our work not only resolves the long-standing question of the enzymatic origin of 3*R* MIAs but also reveals the genomic foundations underlying the initiation and diversification of MIA biosynthesis.

## Results

### A flavoprotein and a CAD-like reductase catalyze C3-stereochemistry inversion of rauwolscine

To investigate *Rauvolfia* MIA biosynthesis and its genomic context, we analyzed the published *R. tetraphylla* genome ^15^ for homologs of known pathway genes. This search revealed a highly repetitive, ca. 200 kbp genomic locus rich in genes encoding CAD-like reductases, including the first committed enzyme for reserpine biosynthesis, yohimbane synthase (RtYOS) ^15^ (Supplementary Fig 1a). Interestingly, just 8.4 kbp upstream of *RtYOS*, we identified a gene encoding a flavoprotein, belonging to the berberine-bridge enzyme (BBE) type (Supplementary Fig 1a). This uncharacterized enzyme shared 57% amino acid identity to *O*-acetylstemmadenine oxidase (CrASO), a known BBE-like oxidase in *C. roseus* MIA biosynthesis ^1^. The close genomic proximity of the flavoprotein gene to *RtYOS* suggested a functional association, motivating us to investigate its biochemical role.

First, we synthesized the gene encoding the *R. tetraphylla* flavoprotein, expressed it in *Saccharomyces cerevisiae*, and purified the His-tagged protein using affinity chromatography. (Supplementary Fig. 1b). Liquid chromatography tandem mass spectrometry (LC-MS/MS) analysis showed that the enzyme could oxidize rauwolscine (*m/z* 355) to a new MIA (*m/z* 353) (Fig. 2a). The *R. serpentina* homolog (90% amino acid identify) also oxidized rauwolcine to generate the same product. The change in the UV absorption maxima of a typical indole (280 nm) as substrate to that with longer wavelengths (350 nm) as product indicated a double bond formation between C3-*N*4, leading to extended conjugation (Fig. 2b, Supplementary Fig. 2a). The formation of a 3-dehydro iminium intermediate by hydride abstraction was also consistent with the flavin chemistry in other BBE-like oxidases, such as CrASO and the California poppy BBE ^1,33^. Reduction of this intermediate with NaBH₄ regenerated 3*S*-rauwolscine, since the geometry of the tertiary nitrogen strongly favours hydride addition from the *re*-face (Supplementary Fig.2b).

**Figure 2.**
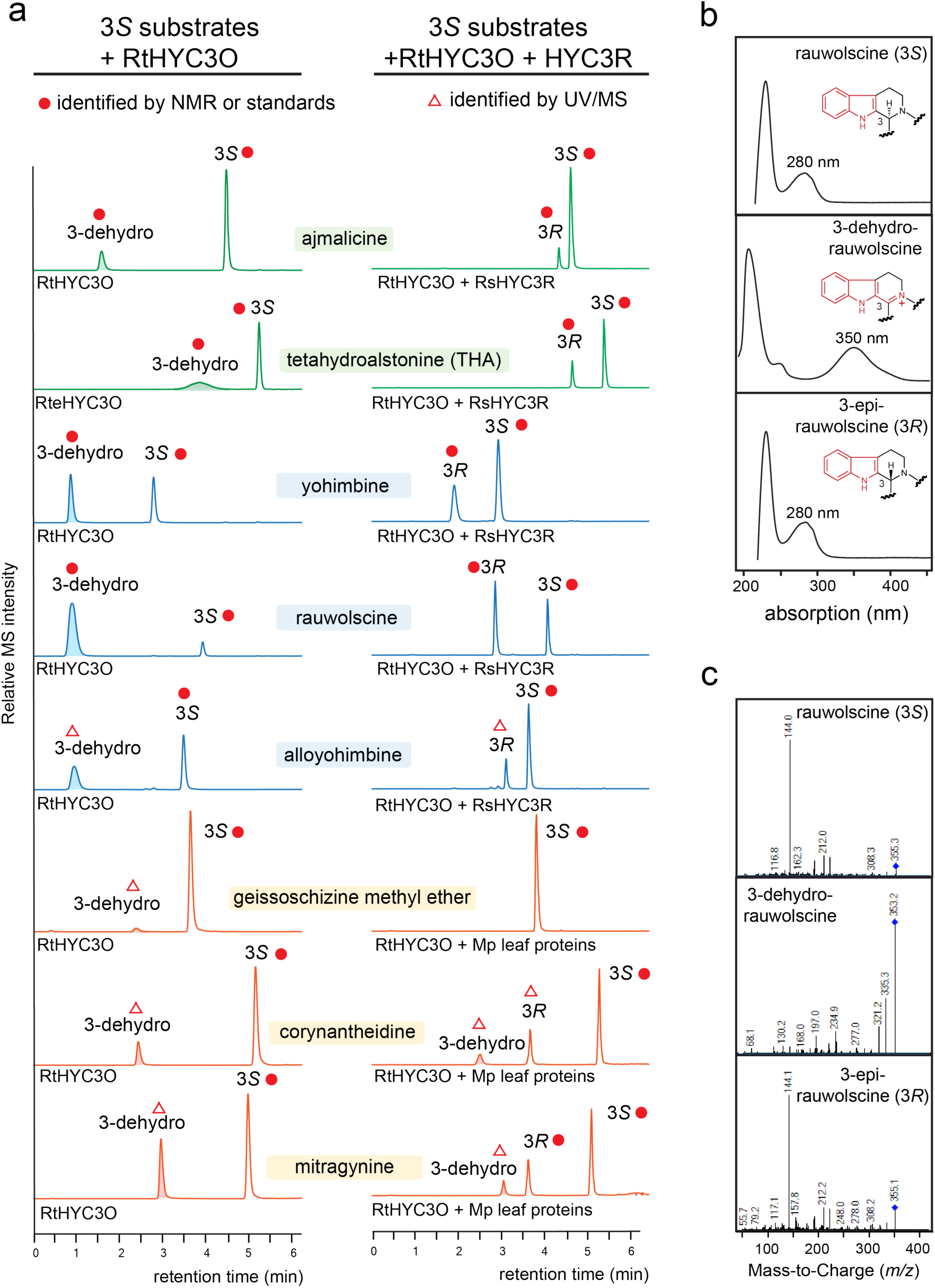
*Rauvolfia* HYC3O and HYC3R catalyze C3 stereochemistry inversion across structurally diverse heteroyohimbine (green), yohimbine (blue), and corynanthe (orange) types of MIA substrates. (**a**) LC-MS/MS [M+H]^+^ Multiple Reaction Monitoring (MRM) chromatograms show the in vitro reaction products generated by RtHYC3O alone, and RtHYC3O in combination with either RsHYC3R or *M. parvifolia* total leaf proteins, using various substrates. A red dot indicates that the MIA was verified by NMR or authentic standards. A red triangle indicates that the MIA was inferred based on its MS and UV profiles. Ajmalicine and THA: MRM 353>144; Yohimbine, rauwolscine, and alloyohimbine: MRM 355>144; geissoschizine methyl ether: MRM 367>144; corynantheidine: MRM 369>144; mitragynine: MRM 399>174; 3-dehydroajmalicine and 3-dehydro-THA: MRM 351>265; 3-dehydro-yohimbine/rauwolscine/alloyohimbine: MRM 353>335; 3-dehydrogeissoschizine methyl ether: MRM 365>249; 3-dehydrocorynantheidine: MRM 367>251; 3-dehydromitragynine: MRM 399>227. (**b**) The UV absorption maximum for 3-dehydro-rauwolscine shifts from 280 nm to 350 nm due to extended conjugation resulting from C3 dehydrogenation. (**c**) C3 dehydrogenation in rauwolscine leads to significant change in its MS/MS fragmentation pattern, compared to those of 3*S*/3*R*-rauwolscine epimers. Product ion scans used for MRM and additional UV absorption profiles are provided in Supplementary Fig. 2.

Because reserpine biosynthesis requires a C3 stereochemical inversion, we hypothesized that a reductase could reduce the C3-*N*4 iminium and generate the 3*R*-epimer. To test this, eight highly expressed, CAD-like reductases (RsCAD1-8) previously identified from *R. serpentina* roots ^19^ were expressed and assayed with 3-dehydro-rauwolscine. Only RsCAD7 reduced the intermediate to 3-epi-rauwolscine (*m/z* 355, Fig. 2a), which displayed identical MS/MS fragmentation and UV spectra to rauwolscine (Figs. 2b and 2c).

Structure elucidation by Nuclear Magnetic Resonance (NMR) experiments confirmed the identities of rauwolscine, 3-dehydro-rauwolscine and 3-epi-rauwolscine (Supplementary Figs. 3-20, Supplementary Tables 1 and 2) ^34^. In 3-dehydro-rauwolscine, disappearance of the H3 resonance and a downfield shift of C3 from 54.1 to 166.1 ppm supported the presence of a C3-*N*4 double bond. For 3-epi-rauwolscine, inversion of the C3 stereocenter reoriented the indole ring perpendicular to the rest of the molecule (Fig. 1). This was evident from the Nuclear Overhauser Effect Spectroscopy (NOESY) correlations observed between H3 and H19 (Supplementary Fig. 7), whereas in rauwolscine (3*S*), H3 correlated with H15 instead (Supplementary Fig. 13). These data provide direct evidence of stereochemical inversion at C3, a key step in the biosynthesis of reserpine.

### The heteroyohimbine/yohimbine/corynanthe C3-oxidase (HYC3O) and C3-reductase (HYC3R) have broad substrate spectra

To assess substrate specificity, we tested the oxidase-reductase pair against MIAs from three major structural classes. *R. tetraphylla* oxidase oxidized rauwolscine, yohimbine, and alloyohimbine (yohimbine type), ajmalicine and tetrahydroalstonine (THA) (heteroyohimbine type), and corynantheidine and mitragynine (corynanthe type) to the corresponding 3-dehydro MIAs (Fig. 2a). In each case, the C3-dehydrogenation was evident from the 2 amu *m/z* loss and product UV absorption maxima shift from 280 nm to 350 nm (Supplementary Fig. 2a). Alloyohimbine, one of the authentic substrates, was isolated from *Corynanthe johimbe* (Yohimbe; Rubiaceae) bark for this study (Supplementary Figs. 21-25, Supplementary Table 1 and 2).

The partner reductase, RsCAD7, reduced 3-dehydro intermediates of the yohimbine and heteroyohimbine classes to their 3*R* epimers, but showed no detectable activity toward corynanthe-type intermediates (Fig. 2a). Based on these results, we named the oxidase and reductase pair the heteroyohimbine/yohimbine/corynanthe C3-oxidase (HYC3O) and C3-reductase (HYC3R), respectively.

Despite the broad substrate scope, RtHYC3O displayed strict stereochemical requirements. This enzyme showed negligible activities with geissoschizine methyl ether (Fig. 2a), rauwolscine’s 20*R*-epimer corynanthine, corynantheidine’s 20*S*-epimer dihydrocorynantheine and mitragynine’s 20*S*-epimer speciogynine. It also did not act on 3-epi-ajmalicine, 3-epi-THA, or 3-epi-mitragynine (speciociliatine), indicating that 3*S*-stereochemistry is needed for these substrates.

We confirmed the structures of 3-dehydro and 3-epi forms of yohimbine, ajmalicine and THA by 1D/2D-NMR from compounds generated in vitro, isolated from plant extracts, or obtained by chemical oxidation (Supplementary Figs. 26-67, Supplementary Table 1 and 2) ^24,34–38^. Additionally, 3-epi-mitragynine (speciociliatine) was identified with a commercial standard. With these results, 3-epi-corynantheidine and 3-epi-alloyohimbine were readily identified by their LC-MS/MS and UV absorption profiles, which mirrored their 3*S*-epimers (Supplementary Fig. 2a).

Interestingly, NMR analysis revealed tautomerization between 3-dehydro and 3,14-dehydro forms. In 3-dehydro-ajmalicine, -yohimbine, and -rauwolscine, the absence of H14 methylene signals indicated equilibrium with the 3,14-dehydro isomers (Supplementary Figs. 15-20, 26-37, Supplementary table 1 and 2). For 3-dehydro-THA, the H14 alkene resonances (δ 4.98, δ 96.4, Supplementary Figs. 38-43) confirmed complete rearrangement to 3,14-dehydro-THA. This dynamic behavior likely explains the broad LC-MS/MS peak observed for 3-dehydro-THA (Fig. 2a). Notably, heating 3-epi-THA greatly reduced its conformer exchange in NMR ^36^.

### Diverse HYC3O and HYC3R enzymes are responsible for MIA C3 stereochemistry inversion in three plant families

Reserpine and other 3*R*-MIAs occur broadly across MIA-producing lineages, suggesting that HYC3O/HYC3R oxidoreductases contribute to their biosynthesis beyond *Rauvolfia*. Phylogenetic analysis confirmed that HYC3O homologs from Rubiaceae, Gelsemiaceae, and Apocynaceae form a distinct clade of BBE-like oxidases, closely related to but separate from *O*-acetylstemmadenine oxidases (ASOs) involved in late-stage Apocynaceae MIA metabolism (Fig. 3a, See Supplementary Fig. 68 and Supplementary data for the complete tree). Similarly, HYC3Rs formed a CAD-like reductase clade distinct from strictosidine aglycone reductases such as GS or dihydrocorynantheine synthase (DCS), but grouping with reductases known to reduce analogous double bonds (e.g., *Strychnos nux-vomica* Wieland-Gumlich aldehyde synthase: SnvWS, CrTHAS2, *R. serpentina* vomilenine reductase: RsVR) (Fig. 3b, See Supplementary Fig. 69 and Supplementary data for the complete trees).

**Figure 3.**
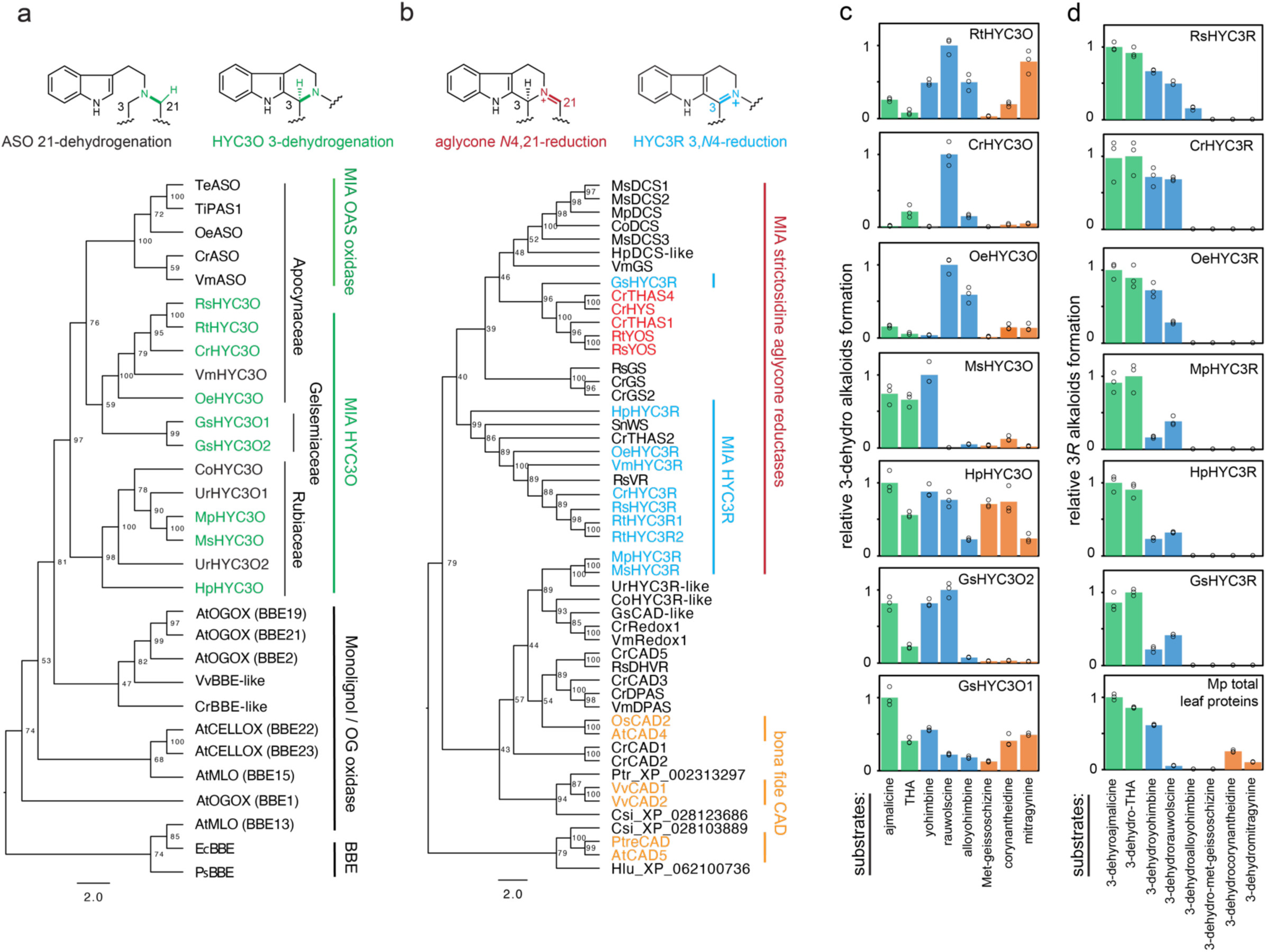
Phylogenetic and biochemical analysis of HYC3O and HYC3R enzymes across Gentianales. (**a**) Phylogenetic analysis reveals a monophyletic origin for the MIA-oxidizing enzymes HYC3O (green) and *O*-acetylstemmadenine oxidase (ASO) within the Gentianales order. These enzymes evolved from broader families of BBE-like flavin-containing oxidases, which include monolignol oxidases (AtMLO) and oligosaccharide oxidases (AtOGOX and AtCELLOX) from *Arabidopsis thaliana*. (**b**) Phylogenetic analysis demonstrates that MIA-reducing CAD-like reductases share a common ancestry with bona fide monolignol dehydrogenases (orange) and other CAD-like enzymes across diverse plant species. All characterized HYC3Rs (blue) from Apocynaceae, along with the HYC3R from *Hamelia patens* (Rubiaceae, an early diverging family in the Gentianales order) form a monophyletic clade, indicating a shared evolutionary origin. In contrast, HYC3Rs from *Mitragyna speciosa*, *M. parvifolia* (Rubiaceae), and *Gelsemium sempervirens* (Gelsemiaceae) fall outside this clade, suggesting they have evolved independently. This pattern points to multiple evolutionary origins of HYC3R function in distinct lineages of MIA-producing plants. The strictosidine aglycone reductases are labeled in red. Co: *Cephalanthus occidentalis*; Cr: *Catharanthus roseus*; Csi: *Camellia sinensis*; Gs: *Gelsemium sempervirens*; Hp: *Hamelia patens*; Hlu: *Humulus lupulus*; Ms: *Mitragyna speciosa*; Mt: *Mitragyna parvifolia*; Oe: *Ochrosia elliptica*; Rs: *Rauvolfia serpentina*; Rt: *Rauvolfia tetraphylla*; Rst: *Rhazya stricta*; Snv: *Strychnos nux-vomica*; Te: *Tabernaemontana elegans*; Ti: *Tabernanthe iboga*; Ur: *Uncaria rhynchophylla*; Vm: *Vinca minor*; At: *Arabidopsis thaliana*; Csi: *Camellia sinensis*; Ec: *Eschscholzia californica*; Os: *Oryza sativa*; Ps: *Papaver somniferum*; Pt: *Populus tremuloides*; Ptr: *Populus trichocarpa*; and Vv: *Vitis vinifera*. Phylogenetic analyses were performed using protein sequences (Source Data) with IQ-TREE 1.6.12, employing the LG substitution model. The full phylogenetic trees including more sequences from diverse plant species are found in Supplementary Figs. 68 and 69 and Supplementary data. **c**) Relative enzyme activities of various HYC3Os with different substrates. (**d**) Relative enzyme activities of various HYC3Rs with different substrates. For (**d**) and (**e**), the activity for the substrate with the highest conversion rate was set as 1, with other rates normalized to this value. Substrates include heteroyohimbine type (green): ajmalicine, tetrahydroalstonine (THA); yohimbine type (blue): yohimbine, rauwolscine, alloyohimbine; and corynanthe type: geissoschizine methyl ether, corynantheidine, mitragynine, and their respective 3-dehydro derivatives. Data were generated from three technical replicates, and the bar graphs display mean values, with individual data points shown as circles for each mean (Source Data).

To probe their biochemical functions, we expressed homologs from *C. roseus* and *O. elliptica* (Apocynaceae), *H. patens*, *M. speciosa*, and *M. parvifolia* (Rubiaceae), and *G. sempervirens* (Gelsemiaceae) in yeast. Despite sequence diversity (57-76% identity), all HYC3Os exhibited dehydrogenase activity, though with distinct substrate preferences (Fig. 3c). RtHYC3O, consistent with its native role in reserpine biosynthesis, was most active on rauwolscine but also unexpectedly accepted mitragynine, a unique MIA from *M. speciosa*. MsHYC3O preferred yohimbine, ajmalicine, and THA but was inactive on rauwolscine. HpHYC3O and GsHYC3O2 showed balanced activity across all tested substrates, while GsHYC3O1 favored yohimbine/heteroyohimbine types but not corynanthe substrates. Strictosidine was universally excluded, highlighting a strict requirement for 3*S* geometry and aglycone-like scaffolds.

The partner reductases also showed functional divergence. HYC3Rs from *C. roseus*, *O. elliptica*, and *H. patens* efficiently reduced 3-dehydro intermediates of yohimbine and heteroyohimbine types (Fig. 3d) but were inactive on fully aromatized β-carbolines such as serpentine and alstonine (See Fig. 1 for their structures). SnvWS, CrTHAS2, and RsVR lacked HYC3R activity, despite their close phylogenetic relationship to the HYC3Rs (Fig. 3b). Surprisingly, canonical HYC3Rs were absent from *M. speciosa*, *M. parvifolia*, and *G. sempervirens* transcriptomes. Yet, leaf protein extracts from *Mitragyna* species reduced most 3-dehydro MIAs (Fig. 3d), implying the existence of alternative reductases. Screening candidate CAD-like enzymes revealed two atypical HYC3Rs: MpHYC3R, previously annotated as a THAS ^39^, clustered near CrRedox1 for stemmadenine biosynthesis, while GsHYC3R grouped with strictosidine aglycone reductases (Fig. 3b). Both could reduce yohimbine/heteroyohimbine intermediates but not corynanthe types. The observed reduction of corynanthe intermediates in *Mitragyna* leaves suggests additional, non-CAD-like reductases are involved for speciociliatine and related 3*R*-corynanthe MIAs.

### Active site architectures reveal the substrate spectra of HYC3Os and the dual catalytic activities of HYC3Rs

Homology modeling (AlphaFold 3) and substrate docking [Molecular Operating Environment (MOE)] revealed how active site architectures shape the substrate preferences of HYC3Os and HYC3Rs (Figs. 4a-e, Supplementary Figs. 70-92). The high rauwolscine binding affinity for RtHYC3O (*K*_M_ 1.15 µM, Supplementary Fig. 93) was supported by the observed hydrogen bonds between Q432/E434 and rauwolscine’s indole/carbomethoxy groups in the model, orienting the H3 position toward FAD’s *N*5 for hydride abstraction (Figs. 4a and c). Mitragynine docked similarly, its 9-methoxy accommodated in a spacious pocket (Fig. 4a), supporting the observed dehydrogenase activity. The dehydrogenase activity is consistent with the presence of a conserved “gatekeeper” residue V176 (equivalent to V169 in California poppy EcBBE) located within the GLCPTV oxygen-binding pocket (GWCPTV in EcBBE). This domain is positioned at the *re*-face of the FAD isoalloxazine ring and plays a critical role in enabling FAD re-oxidation by molecular oxygen, a key step in the dehydrogenation cycle ^40^ .

**Figure 4.**
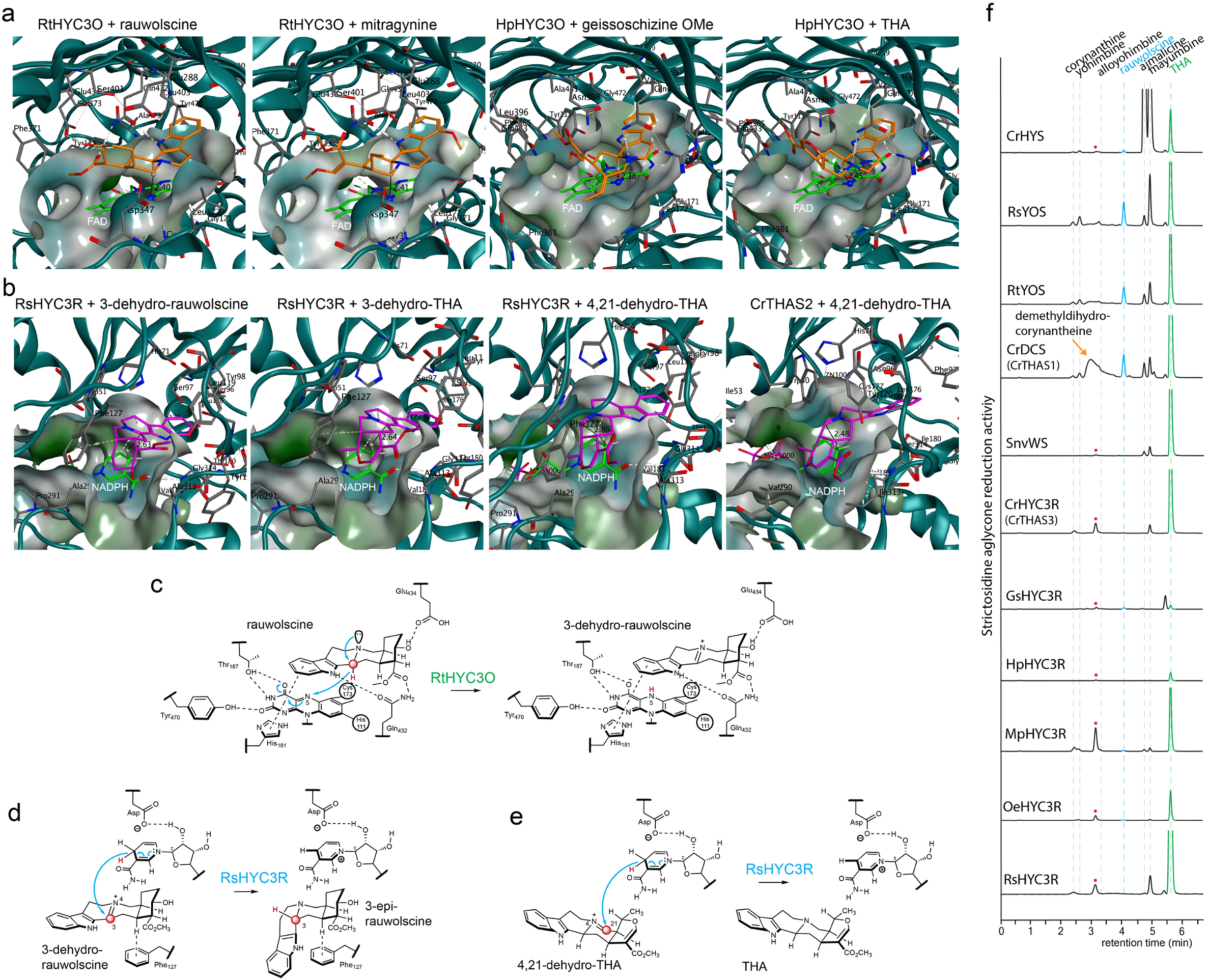
Homology modeling and substrate docking experiments reveal the basis for substrate promiscuity of HYC3Os and dual catalytic activity of HYC3Rs for both 3-dehydro MIA and strictosidine aglycone substrates. (**a**) Docking studies of rauwolscine, mitragynine, geissoschizine methyl ether, and tetrahydroalstonine (THA) at the active sites of RtHYC3O and HpHYC3O homology models. See Supplementary Figs. 73, 77, 79, and 83 for larger representation of these docking models. The surface surrounding the active sites are shown. Hydrogen bond (white dashed lines) networks and π-stacking interactions orient the substrates (orange) for efficient H3 hydride abstraction by FAD (green). The distance between the substrate’s H3 and the *N*5 of FAD is indicated by a black line, with corresponding distances labeled in angstroms (Å). Variations in active site architecture correlate with the observed substrate preferences (**b**) Docking studies of 3-dehydrorauwolscine, 3-dehydro-THA, and 4,21-dehydro-THA at the active sites of the RsHYC3R homology model and CrTHAS2 crystal structure demonstrate key differences in substrate accommodation. See Supplementary Figs. 87, 89, 91, and 92 for larger representation of these docking models. The spacious active site in RsHYC3R supports the binding and reduction of both 3-dehydro-THA and 4,21-dehydro-THA (a form of strictosidine aglycone mixture), accounting for its dual catalytic activity. In contrast, the narrower active site of CrTHAS2 selectively facilitates the reduction of 4,21-dehydro-THA but not 3-dehydro-THA. NADPH is shown in green, and alkaloid substrates are in magenta. White dotted lines show the hydrogen bonds. The distance between the substrate’s H3 or H21 and the hydride donor of NADPH is indicated by a black line, with corresponding distances labeled in angstroms (Å). (**c**) Illustration of rauwolscine binding and oxidation by FAD at the RtHYC3O active site. (**d**) Illustration of 3-dehydrorauwolscine binding and reduction by NADPH at the RsHYC3R active site. (**e**) Illustration of 3-dehydro-THA binding and reduction by NADPH at the RsHYC3R active site. (**f**) LC-MS/MS MRM [M+H]^+^ (355>144 and 353>144) chromatograms show the *in vivo* reduction of strictosidine aglycone by strictosidine aglycone reductases and HYC3Rs. These reductases were expressed in yeast *Saccharomyces cerevisiae* strain AJM-dHYS engineered for *de novo* production of strictosidine aglycone. The reductases exhibited diverse product spectra, reflecting the structural diversity of strictosidine aglycone in equilibrium. An unknown *m/z* 353 peak is labeled with a red dot.

Most HYC3Os contained the hallmark bicovalent Cys/His-FAD bonds of the BBE-like oxidases ^41^. Uniquely, HpHYC3O carried a C173A substitution, forming a single His-FAD bond, a feature also observed in ASOs from *C. roseus*, *V. minor* and *T. elegans* ^1^ . This modification lowers the FAD’s redox potential while enlarging the active site, consistent with HpHYC3O’s broader substrate spectra (Fig. 4a). Key hydrogen bonds (Q400, R286, N398, substrate N4) and π-stacking of indole with FAD in the model supported substrate docking orientations, while an E431A substitution likely contributed to accommodating bulky carbomethoxy groups (e.g., geissoschizine methyl ether). Distances between substrate H3 and FAD *N*5 (<3 Å) correlated with activity profiles across HYC3Os (Supplementary Figs. 70-85).

HYC3Rs, belonging to the CAD-like reductase family, displayed an unexpected dual function. When expressed in a strictosidine aglycone-producing yeast strain (AJM-dHYS) ^19,42,43^, all HYC3Rs, as well as SnvWS and CrTHAS2, reduced strictosidine aglycone to THA, with some also yielding ajmalicine, mayumbine, and an *m/z* 353 isomer (Fig. 4f). By contrast, canonical strictosidine aglycone reductases outside the HYC3R clade (YOS, GS, and DCS) lacked HYC3R activity. These enzymes produced a much broader mixture of yohimban and heteroyohimban epimers (Fig. 4f). These activities aligned with the structural diversity of strictosidine aglycones ^12–15^, reflecting their homologous nature.

Docking and modeling revealed why HYC3Rs tolerate multiple substrates. Strictosidine aglycone naturally exists as a mixture in equilibrium, including two primary forms: cathenamine (20,21-dehydro-THA) and 19-epi-cathenamine. Cathenamine can spontaneously protonate at C20 in solution, forming 4,21-dehydro-THA, which is then reduced by NADPH to produce THA ^13^ (Fig. 4e). Reducing 3,4-dehydro-THA or 4,21-dehydro-THA requires slight adjustments in substrates positioning for double bond reduction.

Compared to CrTHAS2 that lacks HYC3R activity, RsHYC3R possesses a more spacious active site, accommodating both substrates via van der Waals forces, including F127’s arene-C20 interaction suggested by the model (Figs. 4b, 4e, Supplementary Figs. 86-92). This flexibility enabled it to accommodate both 3-dehydro MIAs and strictosidine aglycone intermediates, consistent with its high affinity for 3-dehydro-rauwolscine (*K*_M_ 1.38 µM, Supplementary Fig. 93). In contrast, CrTHAS2’s active site appears sterically restricted, as bulky residues V290 and I313 replace two alanine found in RsHYC3R (Fig. 4b). Additionally, the inward conformation of W60 in CrTHAS2 further narrows the active site, preventing the binding of 3-dehydro substrates.

### HYC3O/HYC3R are encoded in reserpine BGC, which likely arose from a segmental duplication of the geissoschizine synthase (GS) BGC predating Gentianales diversification

Investigating in both *Rauvolfia* and *Catharanthus* revealed that HYC3O and HYC3R are co-localized within a syntenic gene cluster (Fig. 5a). In *R. tetraphylla*, a ∼200 kbp locus begins with *HYC3O* and *YOS*, followed by multiple tandem homologs of *R. serpentina* reductases, vomilenine 1,2-reductase (*VR*) and dihydrovomilenine 19,20-reductase (*DHVR)*, both required for ajmaline biosynthesis ^19^. The cluster further contains an E3 ubiquitin ligase (*E3UL*) and tandem *HYC3R* duplicates. In *C. roseus*, a homologous cluster contains single copies of *HYC3O*, *HYC3R*, *DCS* (*THAS1*), and *E3UL*, but lacks additional CAD-like reductases. Given its role, we refer to this region as the reserpine BGC.

**Figure 5.**
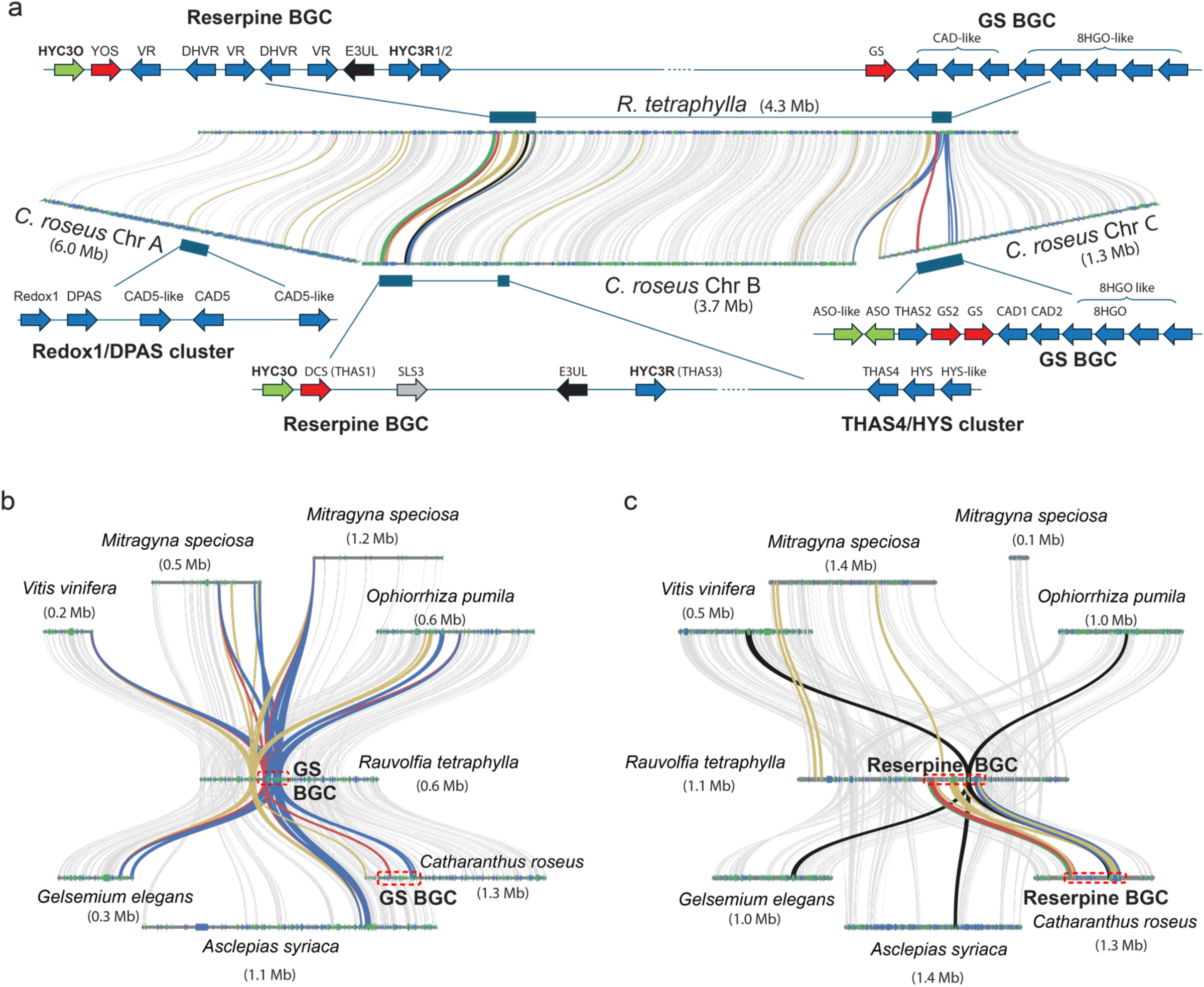
Syntenic reserpine (*HYC3O*/*HYC3R*) and geissoschizine synthase (*GS*) clusters in *R. tetraphylla* and *C. roseus* encode key MIA biosynthetic enzymes, tracing their origin to a genomic block in the common ancestor of Gentianales and grapevine *(Vitis vinifera)*. (**a**) Synteny analysis on a genomic scale between *R. tetraphylla* and *C. roseus* reveals a ∼200 kpb reserpine biosynthetic gene cluster (BGC), comprising genes encoding strictosidine aglycone reductases: yohimbine synthase (YOS) and dimethyldihydrocorynantheine synthase (DCS or THAS1), the redox pair HYC3O/HYC3R that further epimerize YOS/DCS products at the C3 position, and an uncharacterized E3 ubiquitin ligase (E3UL). *R. tetraphylla* reserpine BGC additionally encodes CAD-like reducdtases VR and DHVR for ajmaline biosynthesis. *C. roseus* reserpine BGC additionally encodes a secologanin synthase homolog (SLS3). The GS cluster is conserved between the two species and encodes GS, 8-hydroxygeraniol oxidoreductase (8HGO), and uncharacterized CAD-like reductases and their homologs. In *C. roseus*, the GS BGC also incorporates genes encoding *O*-acetylstemmadenoine oxidase (ASO) and an ASO-like oxidase. *C. roseus* additionally contains two CAD-rich gene clusters encoding Redox1/dihydroprecondylocarpine synthase (DPAS) and tetrahydroalstonine synthase 4 (THAS4)/heteroyohibine synthase (HYS). While both the reserpine and GS BGCs are colocalized on a single chromosome in *R. tetraphylla*, the syntenic blocks in *C. roseus* are dispersed across three chromosomes. Colorations of arrows and syntenic lines correspond with *HYC3O* (green), *HYC3R* (blue), *THAS/GS*/*YOS* (red) encoding strictosidine aglycone reductases, additional CAD-like genes (dark yellow), and other genes (grey). (**b**) Synteny analysis shows that the GS BGC is conserved between *Rauvolfia tetraphylla* and other Gentianales members, including *Mitragyna speciosa*, *Ophiorrhiza pumila*, *Gelsemium elegans*, *Asclepias syriaca*, *Catharanthus roseus*, tracing its origin to *Vitis vinifera*. (**b**) Synteny analysis shows that the reserpine BGC is only conserved between *Rauvolfia tetraphylla* and *Catharanthus roseus*. Genes from *R. tetraphylla* identified in our large-scale phylogenetic trees (primarily encoding CAD-like enzymes; see Supplementary Fig. 69 and Supplementary data) are linked to their corresponding high-scoring pair (HSP) orthologs in other species via dark yellow syntenic lines. Genes homologous to *HYC3R* are highlighted with blue syntenic lines, while genes encoding strictosidine aglycone reductases (*THAS*, *GS*, and *YOS*) are indicated in red. The E3 ubiquitin ligase (*E3UL*) is indicated in black. The remaining HSPs are connected in grey syntenic lines. *M. speciosa*, which underwent a lineage-specific whole-genome duplication, exhibits two subgenomic regions mapping to the *R. tetraphylla* GS and reserpine BGCs. Larger version of synteny graphs are provided in Supplementary Data.

To clarify its evolutionary origin, we compared BBE-like oxidases (*HYC3O, ASO*) and CAD-like reductases (*HYC3R, YOS, GS, HYS, THAS, Redox1, DPAS*) across *R. tetraphylla*, *C. roseus*, and related species. Using the *R. tetraphylla* reserpine BGC scaffold as query, syntenic analyses with SynFind ^44^ application in CoGe ^45^ and MCScan ^46^ uncovered an elusive cluster in *C. roseus* that we designate the geissoschizine synthase (GS) BGC (Fig. 5a). This region encodes ASO and an ASO-like homolog, THAS2, two GS genes, two CAD-like reductases (CrCAD1/2) ^1^, 8-hydroxygeraniol oxidoreductase (8HGO) ^47^, and additional 8HGO homologs. In *R. tetraphylla*, a syntenic GS BGC was also found but lacked *ASO*, consistent with the absence of iboga/aspidosperma MIAs in this lineage.

Additionally, *C. roseus* exhibits a specific genomic organization where *Redox1* and *DPAS* (iboga/aspidosperma MIA biosynthesis) are tightly clustered with *CrCAD5* homologs, and *HYS* and *THAS4* are associated with the reserpine BGC (Fig. 5a). This organization is not conserved in *R. tetraphylla*, where homologs of these genes are absent from the syntenic loci. Notably, these *C. roseus* loci, which map to a single *R. tetraphylla* scaffold, are distributed across three chromosomes (Fig. 5a) These results suggest a complex history of translocations, tandem duplications, and gene losses following divergence of *C. roseus* and *R. tetraphylla*.

Remarkably, the genomic context of the GS BGC is conserved across both MIA producing (*O. pumila, M. speciosa, G. elegans, C. roseus*) and non-producing (*A. syriaca*) Gentianales species, tracing its origin to the Gentianales common ancestor with *V. vinifera* (Fig. 5b). Due to a lineage-specific whole-genome duplication (WGD), *M. speciosa* possesses two subgenomes, both of which map to the *R. tetraphylla* GS BGC (Fig. 5b)

In contrast, the reserpine BGC is restricted to *R. tetraphylla* and *C. roseus*, as the genomic segments containing this cluster are absent in the other species examined (Fig. 5c). The proximity and gene content similarity (e.g., BBE-like oxidases and CAD-like reductases) between the reserpine and GS BGCs support the hypothesis that the reserpine BGC originated from a segmental duplication of the ancient GS BGC in a rauvolfioid ancestor. Their present forms were likely shaped by subsequent duplications, gene losses (e.g., *ASO* loss in *Rauvolfia*), and neofunctionalization (e.g., recruitment of *VR* and *DHVR* in *Rauvofia*).

Functional testing supported this evolutionary trajectory. Two syntenic CAD-like reductases from *V. vinifera* (*VsCAD1* and *VsCAD2*, Fig. 3b), located in the GS BGC region, were confirmed as canonical CADs reducing cinnamyl and coniferyl aldehydes to their corresponding alcohols (Supplementary Fig. 94), suggesting the ancestral lignin-related function of this genomic block. In *G. sempervirens*, we located *GsHYC3R* within its GS BGC, further highlighting this cluster as a source of neofunctionalization. The recruitment of *VR* and *DHVR* into the *R. tetraphylla* reserpine BGC further illustrates how this genomic region has served as fertile ground for the emergence of multiple biosynthetic pathways and products, many of which remain undiscovered.

### **S**trictosidine biosynthesis evolved once in the Gentianales stem lineage

Strictosidine biosynthesis provides the universal precursor immediately upstream of the CAD-like reductases that act on strictosidne aglycone. The STR BGC, containing genes encoding STR, TDC, and a MATE transporter, have been reported in *C. roseus* ^6^, *Rhazya stricta* (Apocynaceae) ^48^, *O. pumila* ^7^, *M. speciosa* ^49,50^, and *G. sempervirens* ^16^. We also identified a STR BGC in *R. tetraphylla* (Fig. 6). However, in *A. syriaca* (also Apocynaceae), the corresponding syntenic block contains only a downstream *TDC* homolog, while a block in *Gardenia jasminoides* (Rubiaceae) contains homologs for *TDC* and *MATE* but lacks *STR* (Fig. 6). This indicates diversification of STR BGC gene content within Gentianales.

**Figure 6.**
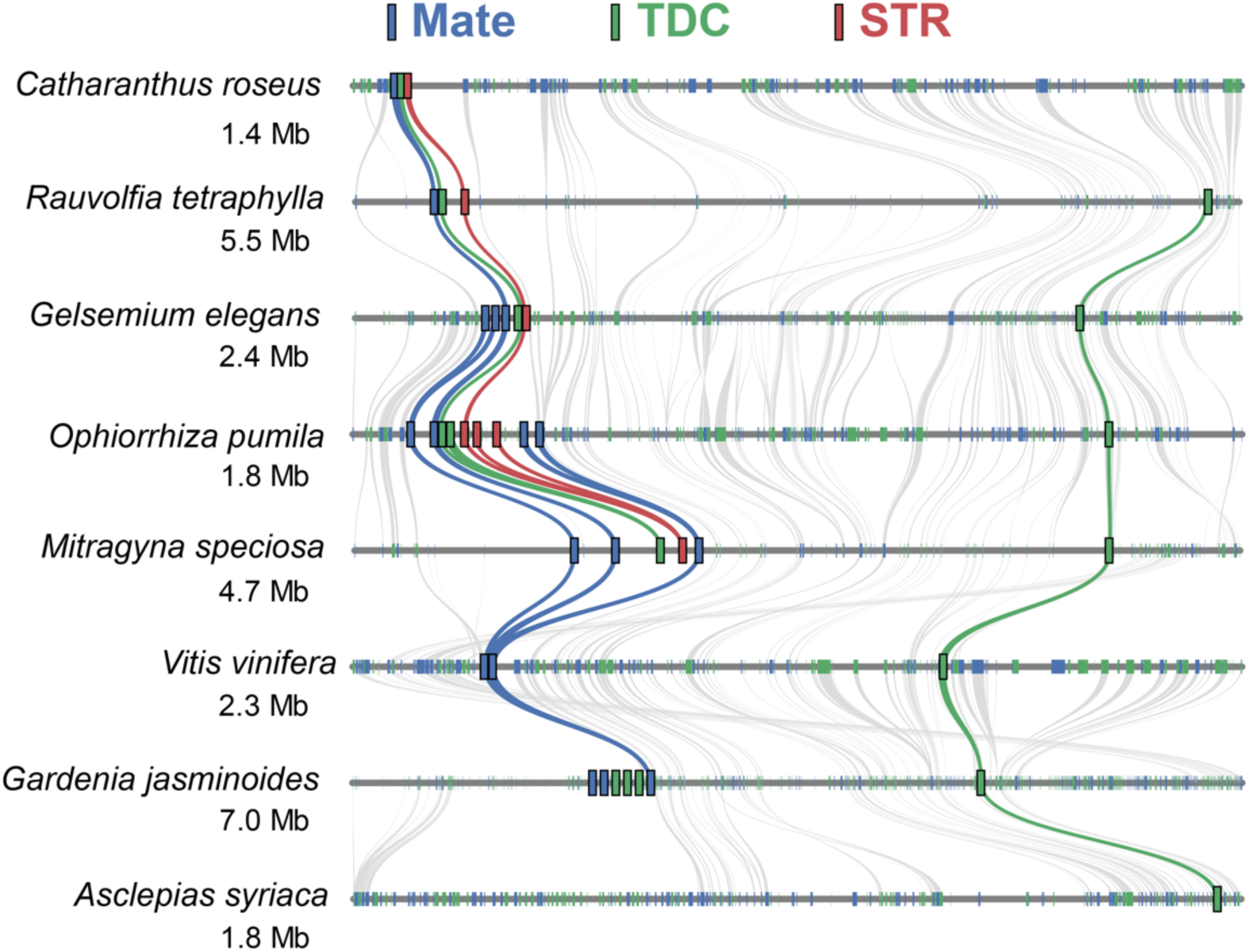
Genomic-scale synteny around the STR BGC for members of the order Gentianales, with grapevine (*Vitis vinifera*) as an outgroup. A syntenic view reveals conservation of the genomic block surrounding the *TDC*/*STR*/*MATE* gene cluster responsible for biosynthesis of the MIA precursor strictosidine. All MIA-producing species in Rubiaceae (*O. pumila* and *M. speciosa*), Gelsemiaceae (*G. elegans*), and Apocynaceae (*C. roseus* and *Rauvolfia tetraphylla*) have complete STR BGCs, with parsimony suggesting the BGC assembled in the Gentianales stem lineage. Evolutionary inferences include: (1) a *MATE* was already in position in the Gentianales/*V. vinifera* common ancestor, (2) a *TDC* also existed downstream of the *MATE*, (3) this (or another) *TDC* duplicated in the Gentianales stem lineage and translocated proximal to the preexisting *MATE*, completing the BGC, (4) a *STR* translocated to the region containing the *MATE* and *TDC* duplicate, completing the BGC, the genes of which subsequently (5) underwent alternative tandem duplications, most notably within Rubiaceae species, (6) the *STR* was deleted in the non-MIA-producing Rubiaceae species *G. jasminoides*, (7) the entire STR BGC was deleted in the non-MIA Apocynaceae species *A. syriaca*, and (8) the downstream *TDC* was deleted in MIA-producing *C. roseus*. Syntenic lines are colored green for *TDC* genes, red for *STR*s, and blue for *MATE*s; grey represents other homologous genes. Where colored lines are multiple within the STR BGC, tandem duplications of *TDC*/*STR*/*MATE* genes can be inferred. Only one of the two subgenomes of tetraploid *M. speciosa* is shown. Numbers below species names indicate lengths of chromosomal regions in Mb.

With complete STR BGCs present in some Apocynaceae, *Gelsemium*, *M. speciosa* and *O. pumila* —all MIA producing species —it is most parsimonious that STR BGC assembled once in the Gentianales stem lineage, with subsequent lineage-specific gene losses. This is consistent with previous genome studies of *O. pumila* and *Pachypodium lamerei* ^51^.

Supporting this hypothesis, synthetic analysis with *V. vinifera* (grapevine), which diverged from other core eudicots ∼148 Mya ^17^, uncovered a syntenic block containing only *TDC* and *MATE* homologs, analogous to the *Gardenia* arrangement (Fig. 7). We therefore propose that STR became physically linked to these pre-existing *TDC* and *MATE* genes in the Gentianales stem lineage (∼135 My) prior to diversification of the Gentianales crown group (∼118 My) ^17^. This timing suggests that the first committed step of complex MIA biosynthesis originated in the early-mid Cretaceous, coinciding with the emergence of the ancestral GS BGC.

**Figure 7.**
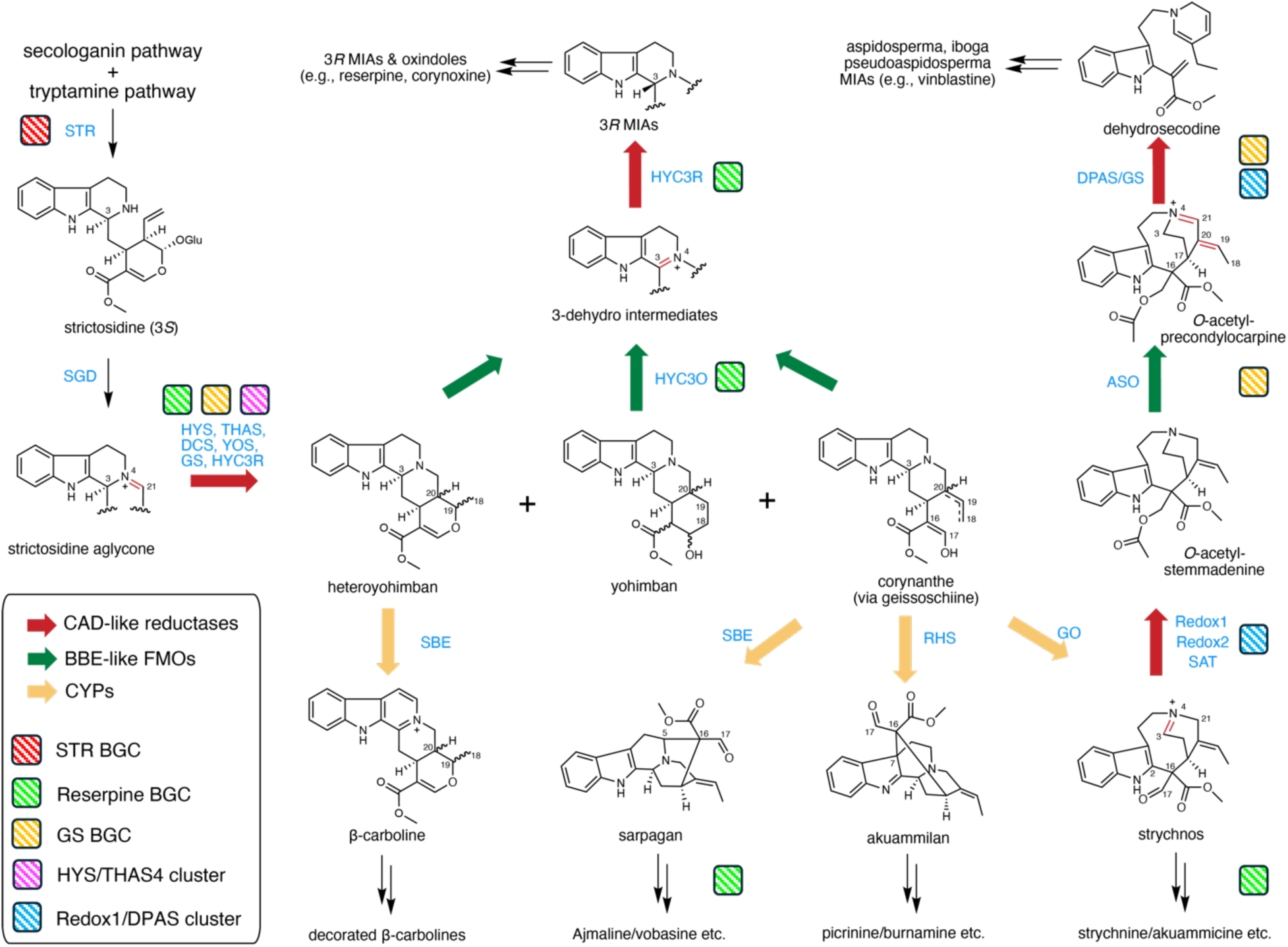
MIA structure diversification is primarily driven by the activities of physically clustered CAD-like reductases, BBE-like oxidases, and non-clustered CYPs. The biosynthetic framework of major MIA subclasses is depicted with representative intermediate structures. Arrows indicate the biosynthetic direction, with corresponding enzyme names and illustrations of the gene clusters to which they belong, positioned adjacent to each transformation step. This schematic highlights the central role of gene clusters (STR, GS, reserpine, Redox1/DPAS, and HYS/THAS4 clusters) in directing the biosynthesis of strictosidine and stepwise conversion of strictosidine aglycone into diverse strychnos, akuammiline, aspidosperma, iboga, spirooxindole, and other scaffolds. SGD: strictosidine β-glucosidase; GS: geissoschizine synthase; THAS: tetrahydroalstonine synthase; HYS: heteroyohimbine synthase; YOS: yohimban synthase; DCS: demethyldihydrocorynantheine/demethylcorynantheidine synthase; SBE: sarpagan bridge enzyme; RHS: rhazimal synthase; GO: geissoschizine oxidase; SAT: stemmadinine *O*-acetyltransferase; ASO: *O*-acetylstemmadenine oxidase; DPAS: dihydroprecondylocarpine synthase; HYC3O: heteroyohimbine/yohimbine/corynanthe 3-oxidase; HYC3R: heteroyohimbine/yohimbine/corynanthe 3-reductase.

### Gene clusters organize the initiation and diversification of MIA biosynthesis

Our synteny analyses of the conserved STR, GS and reserpine BGCs reveal a genomic framework for the initiation and diversification of MIA biosynthesis (Fig. 7). Initiation begins with the STR BGC, the gene products of which facilitate strictosidine biosynthesis, the universal MIA precursor. Following strictosidine deglycosylation, CAD-like reductases, such as HYS, DCS, YOS, and GS encoded by several gene clusters, reduce the reactive aglycone into heteroyohimbine, yohimbine, and corynanthe scaffolds, all of which retain the initial 3*S* stereochemistry. Geissoschizine cyclization via cytochrome P450 monooxygenases represents a critical diversification point, producing the sarpagan, akuammiline, and strychnos scaffolds ^3,52–54^.

In parallel, HYC3O and HYC3R catalyze C3 epimerization of heteroyohimbine, yohimbine, and corynanthe substrates to create a distinct 3*R* MIA branch. The perpendicular indole orientation of these 3*R* structures serves as the substrate for subsequent spirooxindole biosynthesis, formed via C7-oxidation and pinacol rearrangement ^31,55,56^.

In *Catharanthus*, additional enzymes encoded within its GS BGC, *Redox1/DPAS* cluster, and *HYS/THAS4* cluster participate in both upstream and downstream steps of geissoschizine biosynthesis (Fig, 7), leading to diverse aspidosperma and iboga alkaloids like vinblastine. Our results illustrate a previously unrecognized level of gene clustering that organizes and facilitates MIA structural diversification.

## Discussion

### HYC3O and HYC3R phylogeny and function in Gentianales

Our biochemical characterization of HYC3O and HYC3R enzymes resolves a long-standing enigma of the biosynthesis of diverse 3*R* MIAs. The stereochemical modification profoundly alters the structural and biological properties of these metabolites While homologs of HYC3O and HYC3R are broadly distributed across Gentianales, our genomic analyses reveal that their physical clustering is uniquely restricted to the rauvolfioid Apocynaceae lineage.

While HYC3Os form a monophyletic clade closely related with ASOs, HYC3R function has a polyphyletic distribution. In *G. sempervirens*, a CAD-like reductase encoded within the GS cluster independently evolved HYC3R activity, despite being more closely related to strictosidine aglycone reductases like HYS and YOS. Similarly, in *M. speciosa* and *M. parvifolia*, HYC3Rs are more closely allied with Redox1 than with canonical HYC3Rs. A recent study further reported that an isoflavone reductase homolog, unrelated to CAD-like reductases, functions as a HYC3R in *M. speciosa* for corynanthe type substrates ^56^. Conversely *H. patens*, a member of the Rubiaceae family like *M. speciosa*, instead encodes a HYC3R that falls within the canonical HYC3R clade, suggesting descent from a common ancestral gene. However, the absence of a genome assembly precluded confirmation of a corresponding *HYC3O/HYC3R* cluster. Collectively, these findings indicate that HYC3R activity has independently re-emerged in multiple Gentianales lineages, reflecting recurrent evolutionary innovation to enable 3*R*-MIA biosynthesis.

Our biochemical analyses also highlighted broad substrate spectra for HYC3Os, with notable lineage-specific adaptations. Notably, *H. patens* HYC3O naturally carries a C173A substitution, resulting a single FAD-histidine bond, unlike canonical BBE-like oxidases that bind FAD via both cysteine and histidine residues ^41^. Structural modeling suggests this substitution may enlarge the active site and contribute to HpHYC3O’s broad substrate range. Interestingly, the homologous oxidase ASO in *C. roseus*, *V. minor*, and *T. elegans*, central to iboga and aspidosperma biosynthesis, also lacks the cysteine residue and retains only the histidine-FAD linkage ^1,5^. Loss of this bond is expected to lower the redox potential of the FAD cofactor ^41^. The functional implications of this reduced redox potential, and the evolutionary pressures that drove loss of the cysteine-FAD linkage in both HpHYC3O and diverse ASOs across Apocynaceae, remain open questions for future study.

### Evolution of the GS and reserpine BGC

By combining phylogenomic and synteny analysis, we reconstructed the origin of the GS and reserpine BGCs. *Vitis vinifera*, well known for its structurally conserved genome and genome sequence variation compared to many other eudicots ^57–59^, served as an informative outgroup Using the *Vitis* genome, we traced both BGCs to a single ancestral genomic block present ∼148 Mya at the divergence of grapevine from other core eudicots. Synteny between *C. roseus* and *R. tetraphylla* suggests that this ancestral block was subsequently fragmented and distributed across three chromosomes via species-specific translocation events.

Our findings indicate that the reserpine BGC is a more recent innovation, detected only in *R. tetraphylla* and *C. roseus*. Its genomic context suggests that it arose as a rauvolfioid-specific segmental duplication of the GS BGC, followed by functional divergence to specialize in distinct MIA pathways. In contrast, the GS BGC is more ancient and broadly conserved across Gentianales, predating the divergence of Rubiaceae from other families. Our synteny analysis using *Vitis* indicated that the emergence of ancestral GS gene cluster coincides with that of the STR BGC, collectively establishing the genomic and biochemical foundation for MIA diversification.

Consistent with this model, the *THAS4/HYS* and *Redox1/DPAS* clusters appear to have evolved later within Apocynaceae, since the homologous genes are not clustered outside of *C. roseus*. Since all genes in these clusters encode CAD-like reductases, they may have originated either through local duplication within the CAD-rich syntenic block conserved from *Vitis* to Gentianales, or from partial duplications of the older GS BGC. Distinguishing between these scenarios will require further comparative genomic studies, particularly involving strategically positioned Gentianales taxa.

Taken together, our findings highlight the CAD-rich genomic block as a dynamic evolutionary hub for MIA biosynthetic innovation. Beginning with canonical CADs in *Vitis*, this block was progressively reshaped into the GS BGC in Gentianales, facilitating the biosynthesis of corynanthe-type scaffolds such as geissoschizine and dihydrocorynantheine. The reserpine BGC and *HYS*-*THAS4* clusters subsequently enabled the diversification of yohimbine- and heteroyohimbine-type MIAs in Rauvolfioid Apocynaceae, while the emergence of *HYC3Rs* facilitated C3 epimerization and further metabolic branching. The parallel recruitment of BBE-like oxidases like *ASO* and *HYC3O* into these blocks contributed to the diversification of MIA biosynthesis.

Together, the segmental, tandem duplicative ancestry of the reserpine and GS BGCs reflects a broader paradigm in which the duplication of homologous gene clusters and small genomic blocks provides opportunities for chemical innovation in Gentianales ^57,58^. Alongside the large BGCs identified for triterpenoids like QS-21 (9 genes) and benzylisoquinoline alkaloids noscapine and morphine (17 genes) ^60–62^, our data support the prevalence of coordinated specialized metabolism through gene clustering in plants. This work not only uncovers the previously unknown biosynthetic route for 3*R* MIAs but also expands the understanding of MIA genomic organization beyond the canonical vinblastine and ajmaline pathways.

## Materials and Methods

### Cloning

The sequences of HYC3O/HYC3R have been deposited to NCBI GenBank (PP911565-911588, and OQ591889 for RsHYC3R). The open reading frames (ORFs) of all *HYC3O*s/*HYC3R*s except *Rs/CrHYC3R*, *RtYOS*, *SnvWS*, were synthesized and subcloned within the BamHI/Sal sites of pESC-Leu vector (for HYC3O genes) or pESC-Ura vector (for CAD-like reductase genes) (Twist Bioscience, South San Francisco, CA, USA). *RsHYC3R*, *RsYOS*, *CrHYS*, *CrDCS* (*THAS1*), *CrTHAS2*, and *CrHYC3R* (*THAS3*) were amplified from plant cDNA using primer sets 1-12 (Supplementary table 3) and cloned within the BamHI/SalI sites of pESC-Ura and pET30b+ vectors. To generate a *C*-terminally His-tagged RtHYC3O for purification, the ORF was amplified using primer set 13/14 and cloned into the BamHI/SalI restriction sites of the pESC-Leu vector. For yeast expression of *HYC3O*s and *HYC3O/HYC3R* combinations, the pESC vectors were mobilized to *Saccharomyces cerevisiae* strain BY4741 (MATα his3Δ1 leu2Δ0 met15Δ0 ura3Δ0 YPL154c::kanMX4) using a standard lithium acetate/ polyethylene glycol transformation procedure. For de novo production of various reduced strictosidine aglycone products, the vectors containing CAD-like reductases were mobilized to *S. cerevisiae* strain AJM-dHYS ^43^. For RsHYC3R expression in *E. coli*, the pET30b+ construct was introduced into *E. coli* strain BL21(DE3).

### Chemical standards

Authentic chemical standards and substrates were purchased from commercial sources. These included ajmalicine (Sigma Aldrich, St. Louis, MO, USA), yohimbine and corynantheidine (Cayman Chemical, Ann Arbor, MI, USA), corynanthine and rauwolscine (Extrasynthese, Genay, France), and geissoschizine methyl ether (AvaChem Scientific, San Antonio, TX, USA).

### Plant alkaloid purification

For alkaloid purification from plants, fresh leaves (100 g) of *C. occidentalis* or *Luculia pinceana* (Rubiaceae; greenhouse grown) were soaked in ethyl acetate for 1 hr to dissolve alkaloids. The *C. johimbe* bark powder (50 g) was soaked in ethanol for 10 min to dissolve alkaloids. The evaporated extracts were suspended in 1 M HCl and extracted with ethyl acetate. The aqueous phase was basified with NaOH to pH 8 and extracted with ethyl acetate to afford total alkaloids. After evaporation, the crude alkaloids were reconstituted in 0.5 mL methanol and further purified with preparative thin layer chromatography (TLC, Silica gel60 F254, Millipore Sigma, Rockville, MD, USA). THA (Rf 0.77, 3.7 mg) was isolated from *L. pinceana* alkaloids using a mobile phase of acetonitrile:toluene 1:1 (v/v). 3-epi-ajmalicine (Rf 0.38, 3.3 mg) was purified from *C. occidentalis* alkaloids using a mobile phase of toluene: ethyl acetate: methanol 15:4:1 (v/v). For alloyohimbine, the *C. johimbe* alkaloids were separated on TLC with three successive mobile phases. First, acetonitrile: toluene:methanol 15:4:1 (v/v) gave a band with Rf 0.2, which was further separated with acetonitrile:chloroform 1:1 (v/v) to obtain a band with Rf 0.16. Lastly, this was further separated with hexane:ethyl acetate: methanol 5:4:1 (v/v) to obtain pure alloyohimbine (Rf 0.37, 3.2 mg). The structures of purified alkaloids were confirmed with mass spectrometry and 1D/2D NMR analyses.

### De novo alkaloid biosynthesis and alkaloid biotransformation in yeast

Single colonies of the yeasts carrying various vectors were inoculated in 1 mL synthetic complete (SC) media with 2% (w/v) glucose, and incubated at 30 °C, 200 rpm overnight. The yeasts were pelleted by centrifugation, washed once with water, resuspended in 1 mL SC media with 2% (w/v) galactose, and incubated at 30 °C, 200 rpm overnight. For yeast producing alkaloids de novo, the media were directly mixed with equal volumes of methanol for LC-MS/MS analysis. For HYC3O/HYC3R yeasts, the cells were pelleted by centrifugation and resuspended in 0.1 mL 20 mM Tris-HCl pH 7.5 supplemented with 0.5-2 μg alkaloid substrates. The biotransformation took place at 30 °C, 200 rpm overnight, and was mixed with equal volume of methanol for LC-MS/MS analysis.

### Large scale 3-dehydro and 3-epi MIA synthesis and purification

For 3,14-dehydro-THA production, yeast cells expressing RtHYC3O from a 200 ml culture were resuspended in 50 ml of 20 mM Tris-HCl pH 7.5 and fed with 5 mg THA at 30 °C, 200 rpm overnight. After incubation, the reaction was extracted with ethyl acetate, and the crude product was purified by preparative TLC using pure methanol as the mobile phase. The procedure yielded 2 mg of 3,14-dehydro-THA (Rf 0.10).

For 3-dehydro-rauwolscine production, an identical protocol was followed except that 20 mM sodium phosphate buffer (pH 7.5) was used instead of Tris buffer. This substitution was essential, as vacuum drying of 3-dehydro-rauwolscine in Tris buffer led to product degradation, likely due to nucleophilic attack on the electrophilic 3,4-double bond by the amine groups in Tris. No such degradation was observed with phosphate buffer. The dried crude extract was reconstituted in methanol and purified via preparative TLC with pure methanol, yielding 2 mg of 3-dehydro-rauwolscine (Rf 0.13).

For 3-dehydro-yohimbine and 3-dehydro-ajmalicine, a chemical oxidation protocol was followed ^38^ . In brief, to a round bottomed flask, yohimbine (7.7 mg, 0.0217 mmol) or ajmalicine (2.3 mg, 0.0065 mmol) was dissolved in ultra-dry DCM (0.5 mL) and dry Et3N (5.0 uL, 0.0358 mmol for yohimbine and 1.5 uL, 0.0108 mmol for ajmalicine) under N_2_. *t*-BuOCl (5.4 uL, 0.0478 mmol for yohimbine and 1.6 uL, 0.0144 mmol for ajmalicine) was prepared using a previously reported preparation ^63^ and added to the reaction. The solution was stirred for 1 hour at room temperature. The reaction was quenched with water (5 mL) and extracted with DCM (3x3 mL). The organic layer was washed with water (5 mL), then dried over MgSO_4_. The solution was filtered, and solvent was removed under reduced pressure to afford the chloroindolenine. The crude was treated with 3M methanolic HCl (0.77 mL for yohimbine and 0.23 mL for ajmalicine) and allowed to stir for 1 hour under N_2_ at room temperature. The solvent was removed under reduced pressure to obtain the crude iminium products, which were separated by preparative TLC using a solvent system of H₂O:methanol:acetonitrile (1:2:18, v/v/v). This yielded 3-4 mg each of 3-dehydro-rauwolscine (Rf = 0.05) and 3-dehydro-ajmalicine (Rf = 0.06).

For 3-epi-rauwolscine, 3-epi-yohimbine, and 3-epi-THA production, in vitro reactions (50 ml) contained 20 mM Tris HCl pH 7.5, 1 mM NADPH yeast lysates from 200 ml cultures expressing RtHYC3O and RsHYC3R and 3-5 mg 3*S* substrates. Reactions were incubated at 30 °C for 1 hour and subsequently extracted with ethyl acetate The products were purified by preparative TLC using pure methanol, affording 1–2 mg 3-epi-rauwolscine, 3-epi-yohimbine, and 3-epi-THA with Rf values of 0.43, 0.30, and 0.67, respectively. All the purified products were subsequently analyzed by NMR.

### Plant and yeast total protein extraction

Greenhouse-grown *M. speciosa* and *M. parvifolia* leaf tissues (3 g) and 0.2 g polyvinylpolypyrrolidone were ground to a fine powder in liquid nitrogen using a mortar and pestle, which were extracted with ice-cold sample buffer (20 mM Tris-HCl pH 7.5, 100 mM NaCl, 10% (v/v) glycerol). The extracts were centrifuged at 15,000 g for 30 min and desalted into the same sample buffer using a PD10 desalting column (Cytiva, Wilmington, DE, USA) according to the manufacturer’s protocol. The total proteins were desalted once more, and the final samples were stored at -80°C. For HYC3O/HYC3R proteins, yeast cultures (50 ml) expressing various enzymes were pelleted, mixed with 1 mL ice cold sample buffer (20 mM Tris-HCl pH 7.5, 100 mM NaCl, 10% (v/v) glycerol), and mechanically lysed by glass beads (1 mm diameter) using a Qiagen tissuelyser II (Qiagen, Germantown, MD, USA) at 4 °C. The lysate was centrifuged at 20,000g at 4 °C for 10 min. The aqueous phases containing total yeast soluble proteins were stored at -80°C.

### Recombinant protein expression and purification

For RtHYC3O purification, a 500 mL yeast culture expressing *C*-terminal His-tagged RtHYC3O was lysed using glass beads, and the soluble protein fraction was collected by centrifugation as described above. For RsHYC3R purification, *E. coli* BL21(DE3) cells expressing *N*-terminal His-tagged RsHYC3R were grown to an OD₆₀₀ of 0.7, then induced with 0.1 mM isopropyl β-D-1-thiogalactopyranoside (IPTG) at 15 °C and 200 rpm overnight. The cells were lysed by sonication in ice-cold sample buffer (20 mM Tris-HCl, pH 7.5, 100 mM NaCl, 10% (v/v) glycerol) and centrifuged at 10,000 g for 10 minutes at 4 °C to obtain the soluble protein fraction. Both RtHYC3O and RsHYC3R proteins were purified using Ni-NTA affinity chromatography (Cytiva, Marlborough, MA, USA) and subsequently desalted into the sample buffer using a PD-10 desalting column (Cytiva). Purified proteins were stored at -80 °C until further use.

### In vitro enzyme assays

For HYC3O activity, the in vitro assay (100 µL) included 20 mM Tris HCl pH 7.5, 100 ng alkaloid substrate, and 25-45 µL yeast crude lysate containing various HYC3Os (adjusted for the variance of HYC3O activity in each lysate). For each HYC3O, their lysate amounts were kept constant. The assays in triplicates took place at 30 °C for four hours, which was terminated by mixing with equal volumes of methanol for LC-MS analysis. For substrate preference analysis, the 3-dehydro product MS peak areas were compared.

For HYC3O/HYC3R coupled reactions, the in vitro assay (100 µL) included 20 mM Tris HCl pH 7.5, 1 mM NADPH, 100 ng alkaloid substrate, 25-45 µL yeast crude lysate containing various HYC3Os, 1-10 µL yeast crude lysate containing various HYC3Rs, and adjusted for the variance of HYC3R activity in each lysate. For reactions with *Mitragyna* leaf total proteins, 20 µg proteins were used instead of yeast HYC3R lysate. The assays took place at 30 °C for four hours, which was terminated by mixing with equal volume of methanol for LC-MS analysis.

To study HYC3R substrate preference, the seven 3-dehydro substrates were first produced using scaled up (25 ml) reactions with RtHYC3O under otherwise identical HYC3O reaction conditions. The reactions were evaporated to ∼ 1 ml under vacuum as substrates. The in vitro assay (100 µL) included 20 mM Tris HCl pH 7.5, 1 mM NADPH, and 1-5 µL 3-dehydro MIA substrates in excess and adjusted for the variance in substrate amounts, and 1-10 µL yeast crude lysate containing various HYC3Rs. For each HYC3R, their lysate amounts were kept constant, and the substrates should not be completely consumed by end of the assays. The triplicated assays took place at 30 °C for four hours, which was terminated by mixing with equal volumes of methanol for LC-MS analysis. The 3*R* product MS peak areas were compared for HYC3R substrate preference.

For reduction of monolignol aldehydes, the in vitro assay (100 µL) included 20 mM Tris HCl pH 7.5, 1 mM NADPH (for reduction reactions), 2 µg substrates, 2 µg His-tagged and purified VvCAD1/2. The assays took place at 30 °C for 4 hr before LC-MS analysis.

### Enzyme Kinetics

For HYC3O kinetics, each reaction (100 µL) contained 20 mM Tris-HCl (pH 7.5), 0.33 µg of semi-purified RtHYC3O, and rauwolscine substrate at concentrations ranging from 0.078 to 20 µM. For HYC3R kinetics, each reaction (100 µL) contained 20 mM Tris-HCl (pH 7.5), 0.5 µg of purified RsHYC3R, and 3-dehydro-rauwolscine substrate at concentrations ranging from 0.039 to 5 µM. All reactions were carried out at 30 °C for 10 minutes and quenched by the addition of an equal volume of methanol. Samples were centrifuged, filtered, and analyzed by LC-MS/MS. Kinetic parameters were determined by fitting the data to a non-linear regression model using GraphPad Prism version 10.4.2. The data used to generate the enzyme kinetics curves are provided in Source Data.

### LC-MS/MS and NMR

The samples were analyzed using the Ultivo Triple Quadrupole LC-MS/MS system from Agilent (Santa Clara, CA, USA), equipped with an Avantor® ACE® UltraCore C18 2.5 Super C18 column (50×3 mm, particle size 2.5 *µ*m) as well as a photodiode array detector and a mass spectrometer. For alkaloid analysis, the following solvent systems were used: Solvent A, methanol:acetonitrile:ammonium acetate (1M):water at 29:71:2:398; solvent B, methanol: acetonitrile:ammonium acetate (1M):water at 130:320:0.25:49.7. The following linear elution gradient was used: 0-5 min 80% A, 20% B; 5-5.8 min 1% A, 99% B; 5.8-8 min 80% A, 20% B; the flow during the analysis was constant and 0.6 ml/min. The photodiode array detector range was 200 to 500 nm. The mass spectrometer was operated with the gas temperature at 300°C and gas flow of 10 L/min. Capillary voltage was 4 kV from *m/z* 100 to *m/z* 1000 with scan time 500 ms, and the fragmentor performed at 135 V with positive polarity. The MS/MS was operated with gas temperature at 300°C, gas flow of 10 L min^-1^, capillary voltage 4 kV, fragmentor 135 V, and collision energy 30 V with positive polarity. NMR spectra were recorded on an Agilent 400 MR and a Bruker Avance III HD 400 MHz NMR spectrometer in acetone-*d6*, MeOD, or CDCl_3_.

### Homology modeling and substrate docking studies

All computational experiments and visualizations were carried out with MOE version 2022.02 and AlphaFold 3 ^64^ on local computers. All molecular mechanics calculations and simulations employed the AMBER14:EHT forcefield with Reaction Field solvation. Sequence homology searches for HYC3Rs against the Protein Data Bank identified CrTHAS2 complexed with NADP^+^ as the template for homology modeling (PDB ID: 5H81, 68% amino acid identity with RsHYC3R), based on Hidden Markov Model energy scores. Following a QuickPrep of the 5H81 template, homology models were derived in MOE using default settings and scored using the GBVI/WSA dG method. NADPH was modeled and docked into each homology model based on the NADP^+^ binding site in the 5H81. Following NADPH docking, 3,4-dehydro ligands were docked to the HYC3R-NADPH complexes. Rt/HpHYC3O protein models were predicted in AlphaFold3. The FAD-bound models were transferred to MOE, and ligands were docked to the HYC3O-FAD complexes as described above.

For each ligand docking experiment, 17,000 docking poses were initially generated via the Triangle Matcher method and scored by the GBVI/WSA dG function. A subset top-docking poses was refined by the induced fit method, where the bound ligands and active site residues underwent local geometry optimization and rescoring. The top-scoring docking poses with plausible reaction geometry were retained for subsequent analyses. Cartesian coordinates for all homology models and their ligand complexes can be found at DOI: 10.5061/dryad.vdncjsz59.

### Bioinformatics and phylogenetic analyses

The published genome of *R. tetraphylla* (GCA_030512225.1, available at NCBI) was reannotated (with default settings) using Gene Model Mapper version 1.9 by using available reference assembly annotations of several related species *C. roseus* (GCA_024505715.1), *A. syriaca* (CoGe gid61699), *O. pumila* (GCA_016586305.1), and *Calotropis gigantea* (NCBI PRJNA400797). For sequence discovery to synthesize enzymes for experimental analyses, the annotated genomes of *R. tetraphylla* ^15^, *C. roseus* ^65^, *G. sempervirens* ^16^, *G. elegans* ^66^ *, M. speciosa* ^49,50^ were analysed with CoGeBLAST (https://genomevolution.org/coge/) ^67^. Gene family representatives from the *R. tetraphylla* reserpine cluster (our GeMoMa gene model IDs: Catharanthus_roseus_rna_EVM0023822.1_R0, Catharanthus_roseus_rna_EVM0000382.1_R0, and Catharanthus_roseus_rna_EVM0027783.1_R2) were used as query sequences to perform CoGeBLAST searches using TBLASTX with default parameters. The searches were conducted against a selection of species, including *A. scholaris* (gid: 67811)*, E. grandiflorum* (gid: 63766)*, V. vinifera* (v12x; gid: 19990, *O. pumila* (vv1; gid: 63710), *G. elegans* (v1.0; gid: 64491), *A. syriaca* (v0.3; gid: 61699), *Mitragyna speciosa* (vv1; gid 63699), *R. tetraphylla* (v4; gid: 69143), and *C. roseus* (vASM2450571v1; gid: 65259). Protein sequences were aligned using Muscle. The alignments were trimmed to remove poorly aligned regions using GBlocks (all three sensitivity options boxes checked) in Seaview, a sequence processing tool. Phylogenetic trees were constructed using IQ-TREE 1.6.12 with the LG substitution model. A total of 1000 bootstrap replicates were performed to estimate support values for each node in the tree. Phylogenetic trees of HYC3O and HYC3R enzyme families (Figs. 3a and 3b) were constructed in the same manner as the large gene family trees. Inferred amino acid sequences for the latter are given in Source Data.

### Plant genome structural analysis

To begin, we reannotated the available genome of *R. tetraphylla* using GeMoMa as described above. We checked the annotated assembly using SynMap in CoGe to evaluate its whole-genome duplication (WGD; polyploidy) status and any additional WGDs following the *gamma* hexaploidy event ^68^ at the base of all core eudicots, as reported in the *R. tetraphylla* genome study ^15^. While the *R. tetraphylla* genome publication reported that haplotypic contigs were purged and that no WGDs were observed, our self:self syntenic analysis using SynMap in CoGe^69^ showed the presence of internal syntenic blocks that suggest a recent WGD based on the overall low Ks (synonymous substitution rate) values for homologous gene pairs. However, for a putative WGD, the numbers of such low-Ks pairs in *R. tetraphylla* closely matched the number of high-Ks pairs retained since the ancient *gamma* hexaploidy event; *G. elegans* and *V. vinifera* self:self SynMaps and a *R. tetraphylla*:*C. roseus* plot also revealed old *gamma* peaks comprising similarly low numbers of gene pairs (Supplementary Figs. 95, 96). Although heavy fractionation (alternative homolog deletion on different subgenomes) of *gamma* over time was anticipated, it was unusual to observe similar fractionation for such a recent WGD, as would otherwise be suggested by the small number of low-Ks syntenic homolog pairs. Further syntenic analyses executed in MCScan in the JCVI application ^46^ supported these observations. A self:self syntenic dotplot for *R. tetraphylla* showed the same internally duplicated blocks identified by SynMap (Supplementary Fig. 97), but syntenic depth histograms did not yield a sufficient number of doubled blocks to suggest that a WGD accounted for their presence (Supplementary Fig. 98). We further checked these results using the software Ksrates ^70^ (version 1.1.3<u>59</u>), which corrects for unequal Ks rates in different taxa using a phylogenetic tree and provides relative timings via Ks values for species splits and any WGD events inferred. Coding sequence fasta files were extracted using AGAT version 1.0.0<u>65</u> (See Supplementary Fig. 96). Based on this analysis, and the previous SynMap results, we conclude that the extremely recent “syntenic” gene pairs represent alternative haplotypes that remained unpurged in the assembly assembly, resulting in partially diploid regions, despite the reported use of Purge Haplotigs by the original investigators. The conclusion was supported by our examination of two scaffolds, both containing a reserpine cluster, which showed nearly identical sequences in their overlapping regions. Consequently, we focused all subsequent structural analyses on the longer of the two *R. tetraphylla* scaffolds, which contains both the reserpine and GS BGCs.

Color highlighting of genes on the McScan visualization was based on high-scoring pairs (HSPs) identified through synteny-based comparisons, focusing on conserved genomic regions between *R. tetraphylla* and related species. The identity and evolutionary placement of these HSPs were confirmed by assessing their proximity to reference genes within large-scale phylogenetic gene trees. Syntenic analyses across Gentianales and *V. vinifera* provided a comparative framework for conserved genomic blocks. In the resulting diagrams, *R. tetraphylla* genes encoding homologs of YOS and CADs were shaded in dark yellow to indicate their HSP relationships, while experimentally supported biosynthetic genes were linked by color-coded connectors.

For the MCScan analyses noted above (syntenic dot plots, syntenic depth) as well as syntentic karyotype plots between assemblies, input annotation files were converted to gff3 format using AGAT version 1.0.0<u>65</u>. AGAT was also used to extract CDS fasta files using each species’ assembly and reformatted annotation. When detecting synteny between two species with the same ploidy level, a C-score cutoff of 0.99 (–cscore = 0.99) was used to filter out high-Ks pairs (i.e., greater than the *gamma* hexaploidy’s expected Ks) for clearer connections between syntenic blocks. Otherwise, default options were used to generate figures.

## Acknowledgements

This research was supported by a Natural Sciences and Engineering Research Council of Canada (NSERC) Discovery Grant, a New Brunswick Innovation Foundation (NBIF) Research Assistantship Initiative Grant, and a NBIF Research Professional Initiative grant to Y.Q. V.A.A. acknowledges support from U.S. National Science Foundation grant 2030871. The authors thank the Chemical Computing Group ULC (www.chemcomp.com) for MOE licenses. This research was enabled in part by support provided by ACENET (www.ace-net.ca) and the Digital Research Alliance of Canada (<u>alliancecan.ca</u>).

## Conflict of interest

The authors declare that there are no conflicts of interest.

## Data availability

The data supporting the conclusion of this study are included in the article, supplementary information, and Source Data file. RNA-seq data have been deposited in the NCBI Sequence Read Archive under accession numbers SRR32911581, SRR32911592, and SRR32911600. Newly identified gene sequences have been submitted to GenBank under accession numbers PP911565–PP911588 and OQ591889. Additional datasets related to genome annotation and synteny analyses have been deposited at DOI: 10.5061/dryad.vdncjsz59.

## Notes

### Competing Interest Statement

The authors have declared no competing interest.

### Summary of Updates

1. A thorough language/writing editing throughout the manuscript, without altering the experiments or conclusions. 2. Change of figure order to reflect the change in the manuscript writing. 3. Small revision to the title from "Ancient gene clusters initiates monoterpenoid indole alkaloid biosynthesis and C3 stereochemistry inversion" to "Ancient gene clusters govern the initiation of monoterpenoid indole alkaloid biosynthesis and C3 stereochemistry inversion" 4. Inclusion of three new authors who assisted in enzyme kinetics and product analysis during revision.

