## Supplementary information for "Ancient gene clusters govern the initiation of monoterpenoid indole alkaloid biosynthesis and C3 stereochemistry inversion"

**Supplementary Figure 98. MCScan syntenic depth histograms of *Rauvolfia tetraphylla* against *Catharanthus roseus* and the reverse (top), and *Rauvolfia tetraphylla* against *Vitis vinifera* and the reverse (bottom), reveal some duplicated blocks in the published *Rauvolfia tetraphylla* genome ...105**

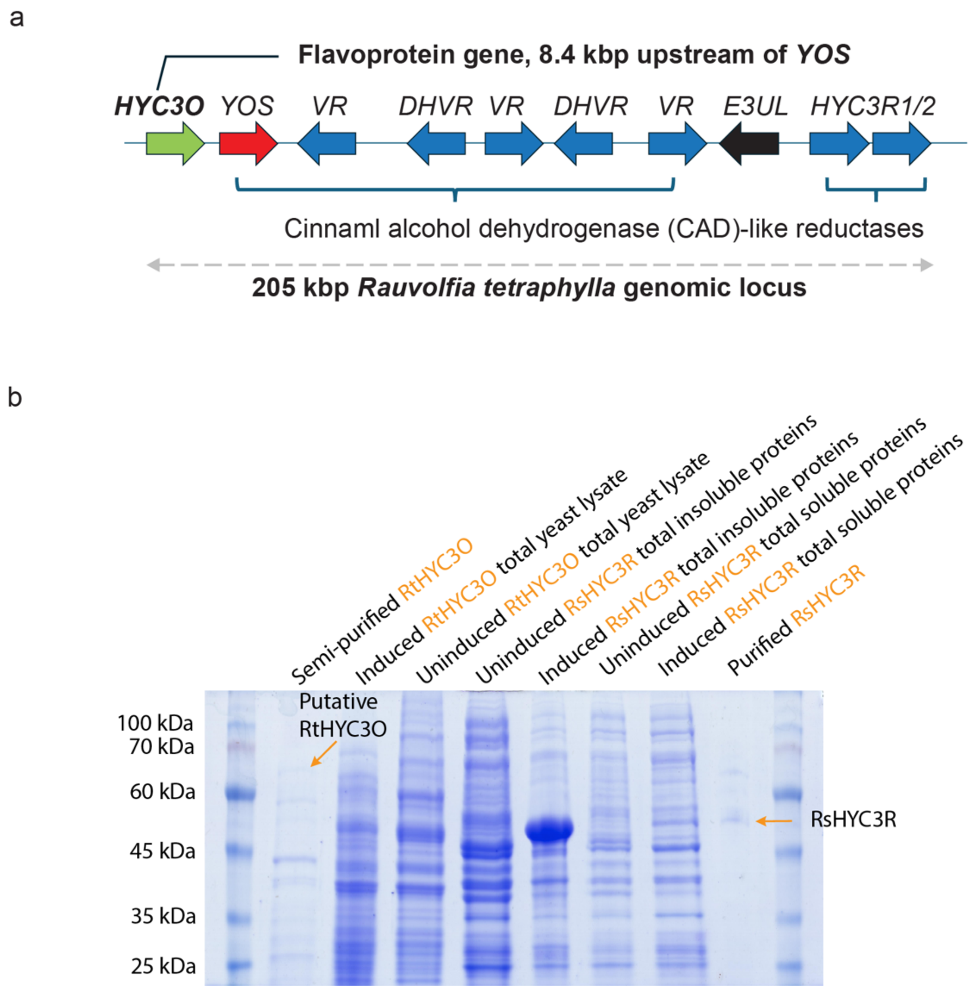

**Supplementary Figure 1. The genomic locus encoding yohimban synthase (YOS) and a newly identified flavoprotein (a) and SDS-PAGE showing purifications of RtHYC3O from yeast and RsHYC3R from *E. coli* (b).**

(**a**) Analysis of the published *Rauvolfia tetraphylla* genome [1] revealed a 205-kbp locus enriched in genes encoding cinnamyl alcohol dehydrogenase (CAD)-like reductases, including the gene for the recently characterized yohimban synthase (YOS). Located 8.4 kbp upstream of *YOS* is an uncharacterized flavoprotein gene, designated in this study as heteroyohimbine/yohimbine/ corynanthe C3-oxidase (HYC3O). Downstream of HYC3O, the locus contains five tandem homologs of *R. serpentina* 1,2-vomilenine reductase (*VR*) and 1,2-dihydrovomilenine 19,20-reductase (*DHVR*) [2] . Following an uncharacterized E3-ubiquitin ligase-like gene (*E3UL*), there are two additional CAD-like genes, which has been designated as heteroyohimbine/yohimbine/ corynanthe C3-reductase (HYC3R) in this study. (**b**) For RtHYC3O, expression of the recombinant protein with a *C*-terminal 6×His-tag was induced by galactose in a 200 mL *Saccharomyces cerevisiae* culture. For RsHYC3R, expression of the *N*-terminal 6×His-tagged protein was induced by isopropyl β-D-1-thiogalactopyranoside (IPTG) in a 150 mL *E. coli* culture. Cell lysates were purified using standard Ni-NTA affinity chromatography and analyzed by SDS-PAGE. While His-tagged RsHYC3R was clearly detected in the SDS-PAGE gel, His-tagged glycoprotein RtHYC3O (theoretical size 60.8 kDa without glycosylation) could only be partially purified, likely due to its low expression level in yeast.

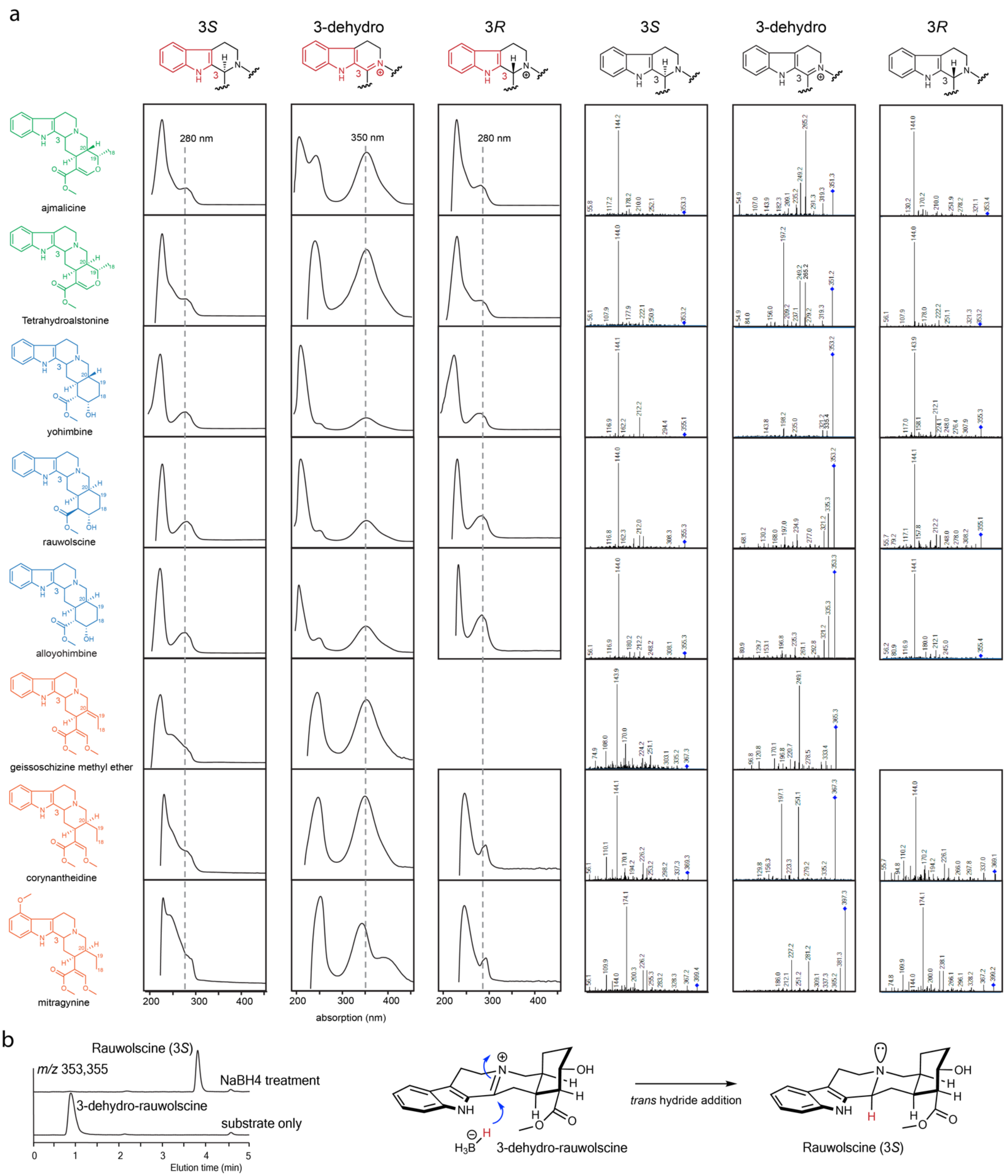

**Supplementary Figure 2. Structure, UV absorption, and MS/MS fragmentation patterns of**

**alkaloids in this study (a) and $NaBH_4$ mediated 3-dehydro-rauwolscine reduction to 3*S*-rauwolscine (b).**

The geometry of the pyridine ring strongly favors *trans* hydride addition, minimizing steric hindrance that would arise from *cis* hydride addition. The LC-MS chromatogram displays the extracted ion chromatogram (EIC) with $[M+H]^+$ *m/z* 353 and 355, corresponding to 3-dehydro-rauwolscine and rauwolscine, respectively.

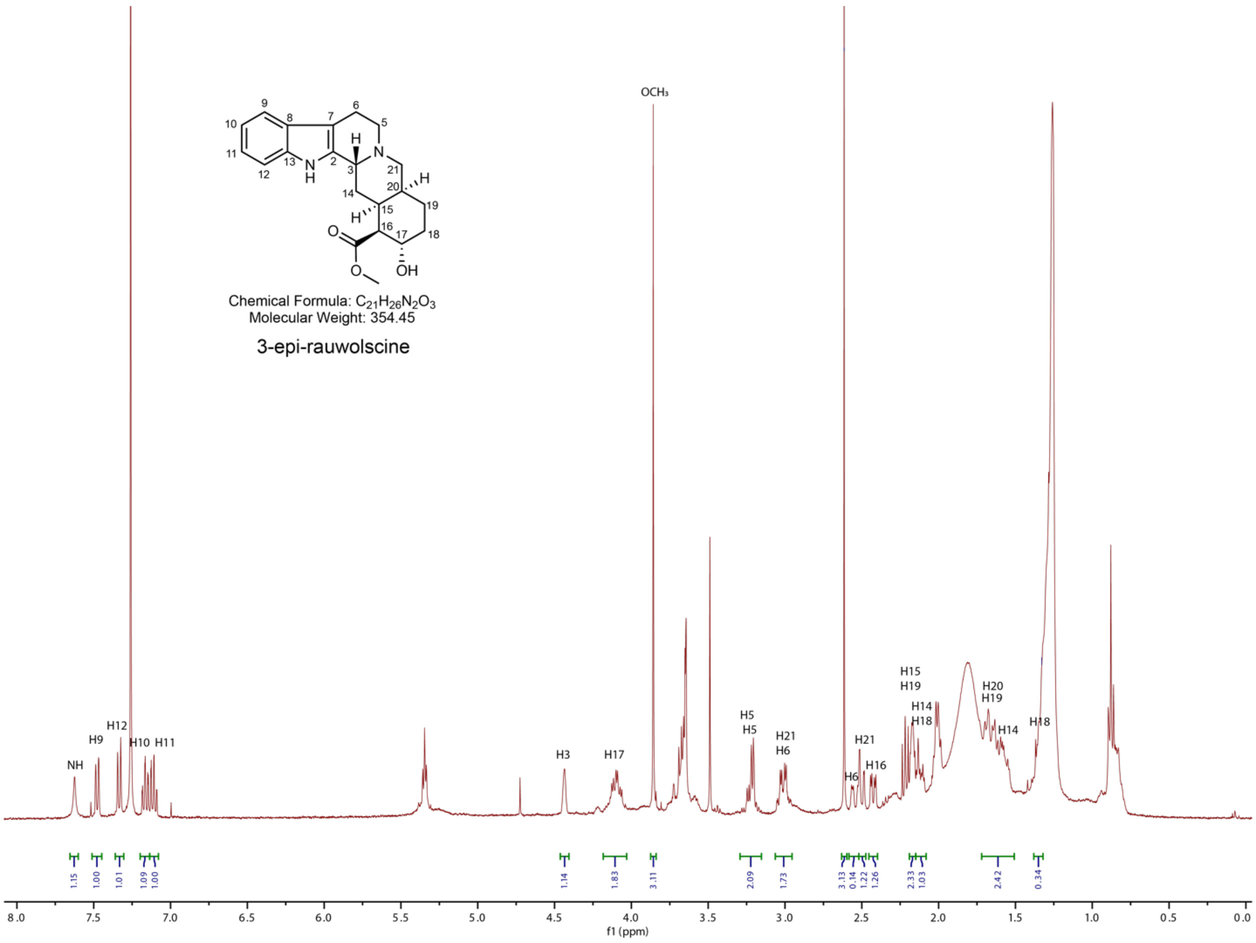

**Supplementary Figure 3. [1]H NMR spectra of 3-epi-rauwolscine in $CDCl_3$.**

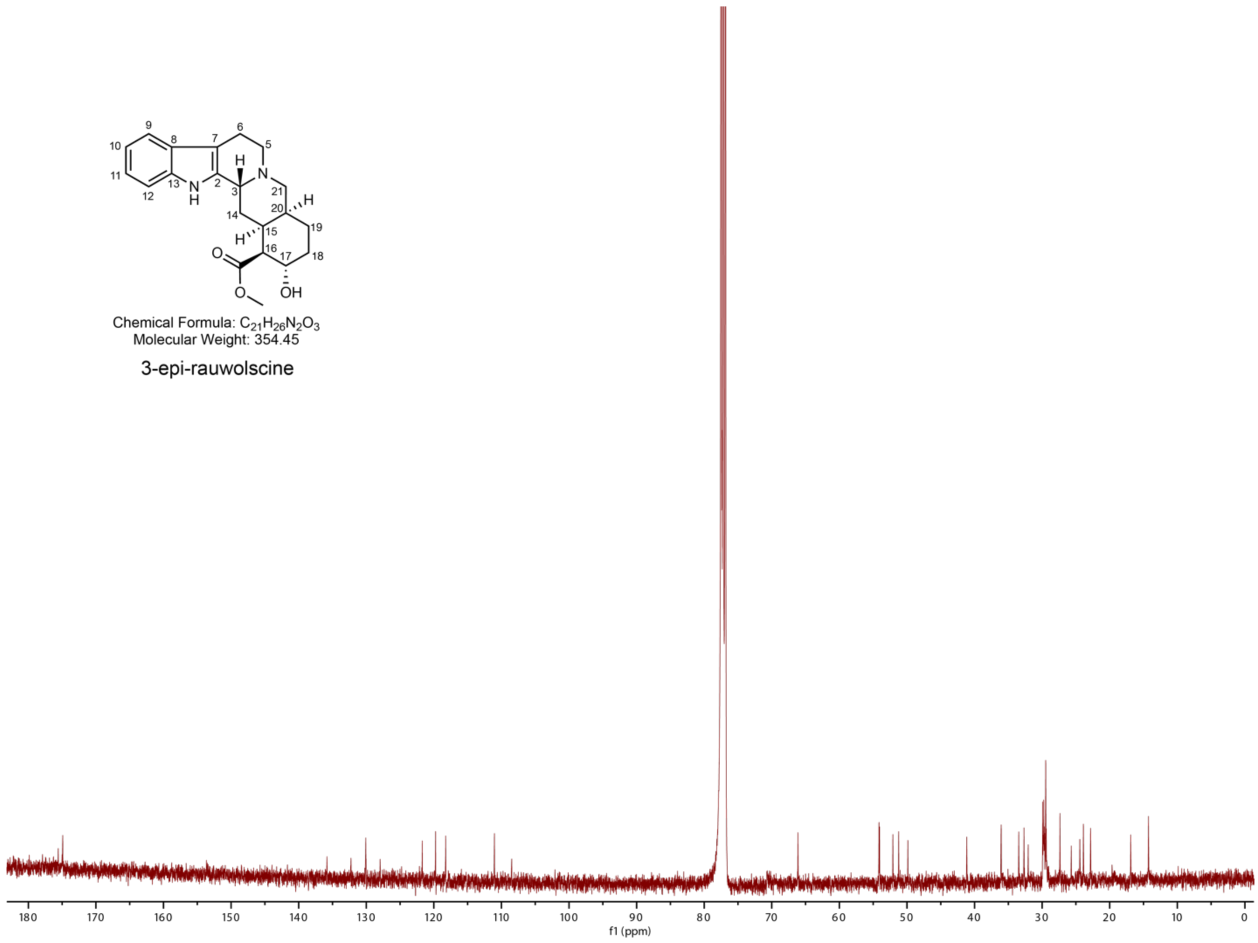

**Supplementary Figure 4. $^{13}$C NMR spectra of 3-epi-rauwolscine in $CDCl_3$.**

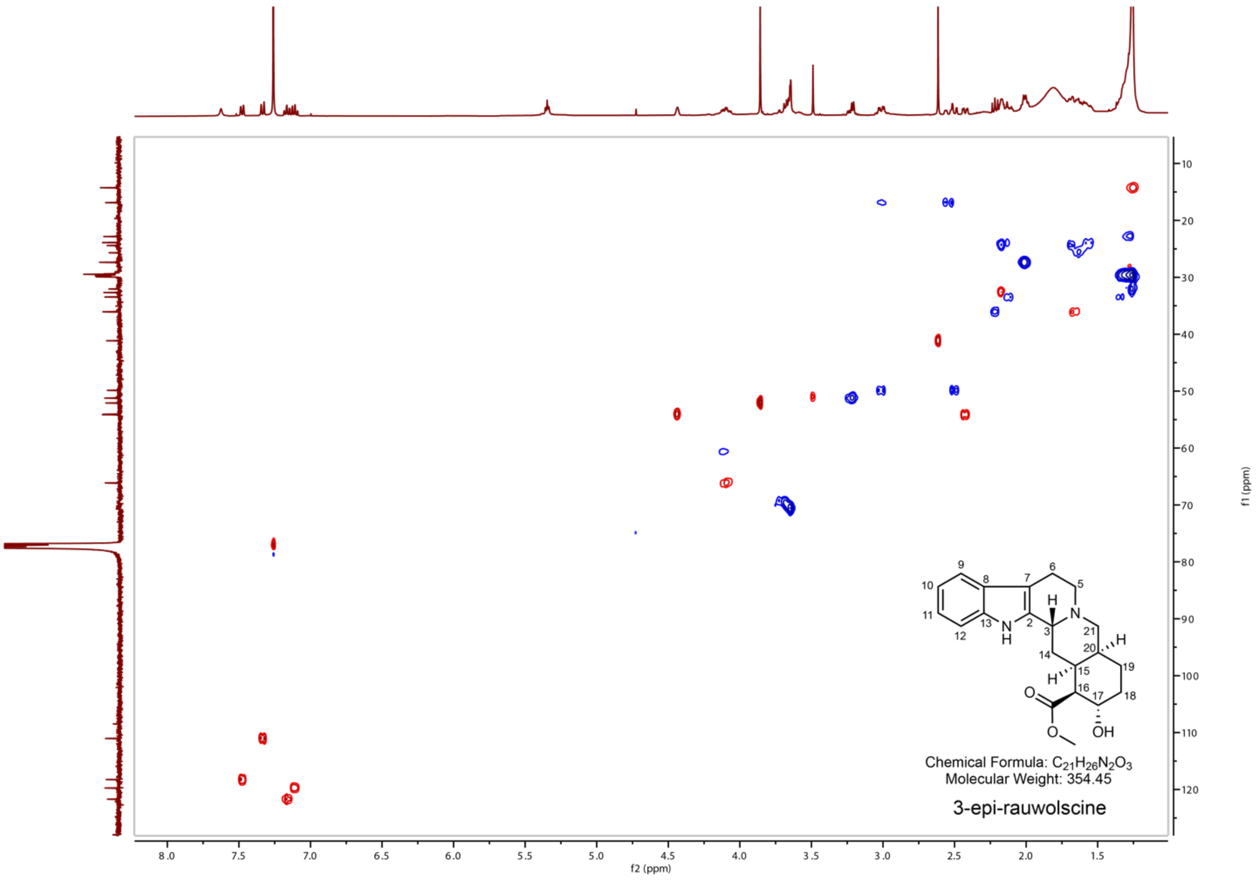

**Supplementary Figure 5. HSQC NMR spectra of 3-epi-rauwolscine in CDCl₃.**

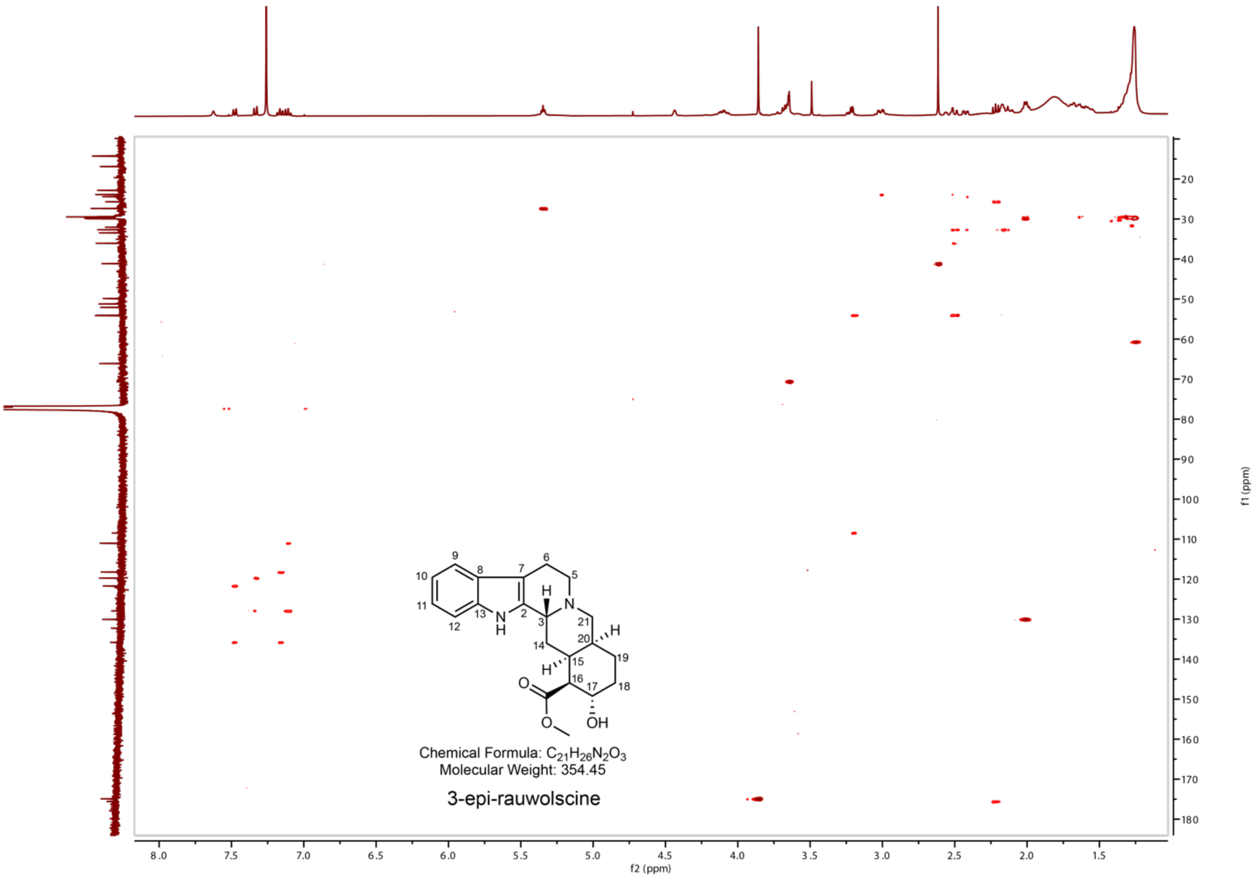

**Supplementary Figure 6. HMBC NMR spectra of 3-epi-rauwolscine in $CDCl_3$.**

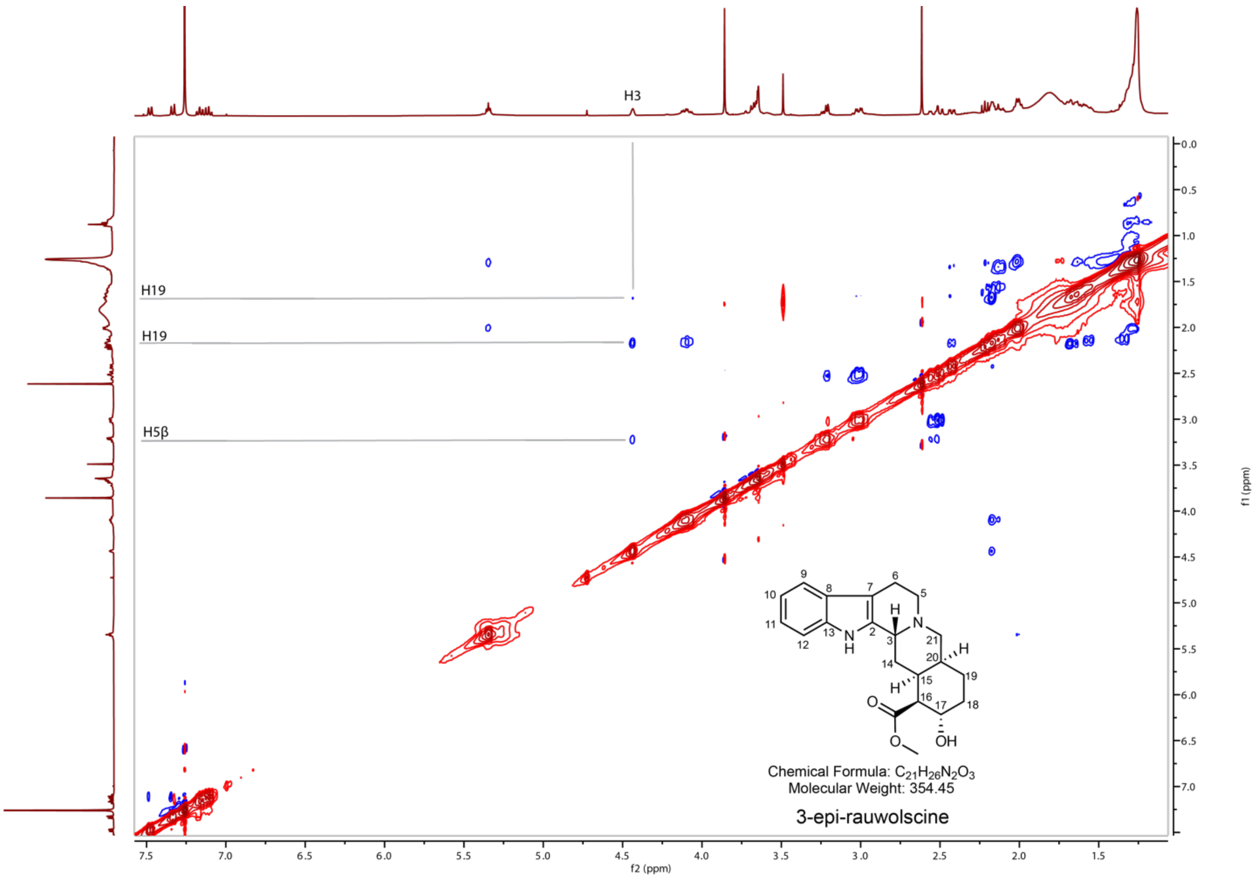

**Supplementary Figure 7. NOESY NMR spectra of 3-epi-rauwolscine in $CDCl_3$.**

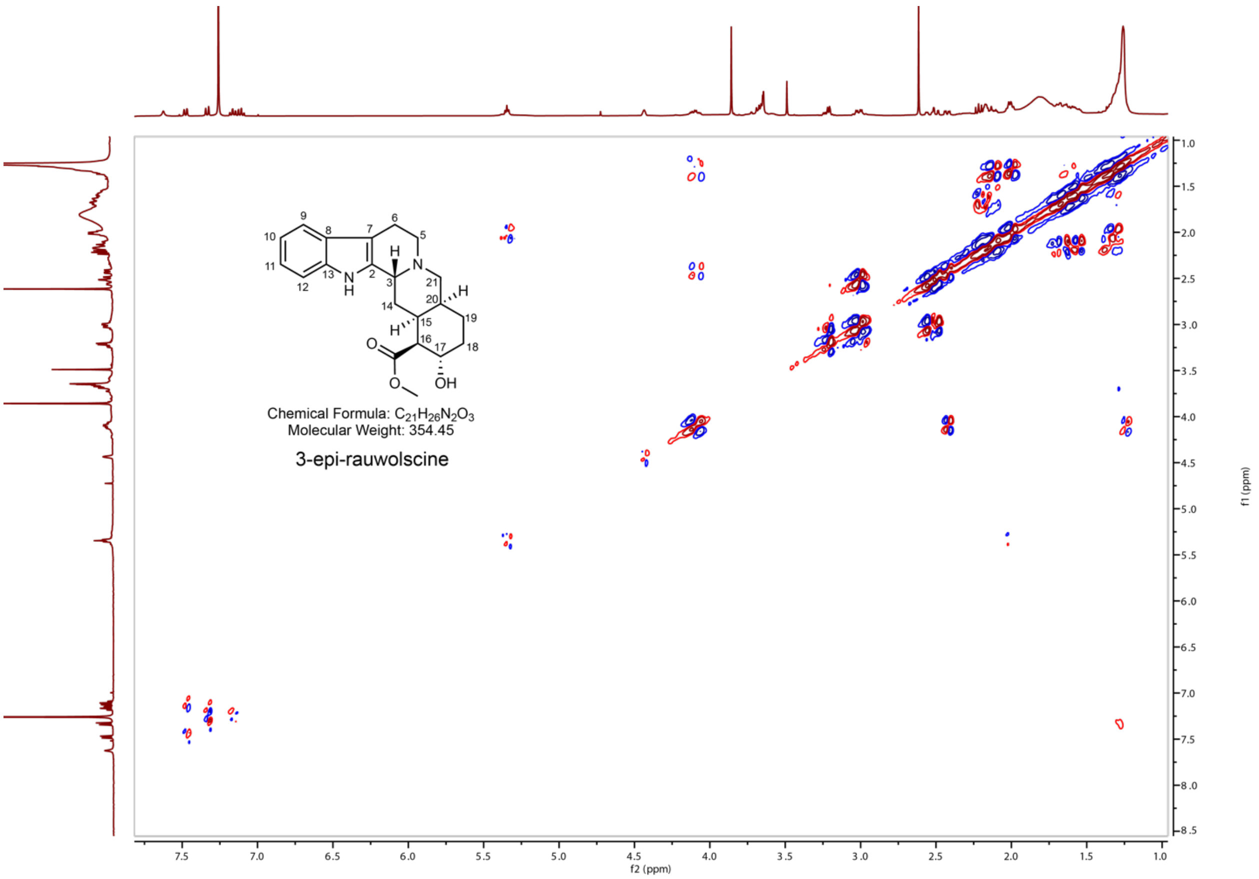

**Supplementary Figure 8. COSY NMR spectra of 3-epi-rauwolscine in $CDCl_3$.**

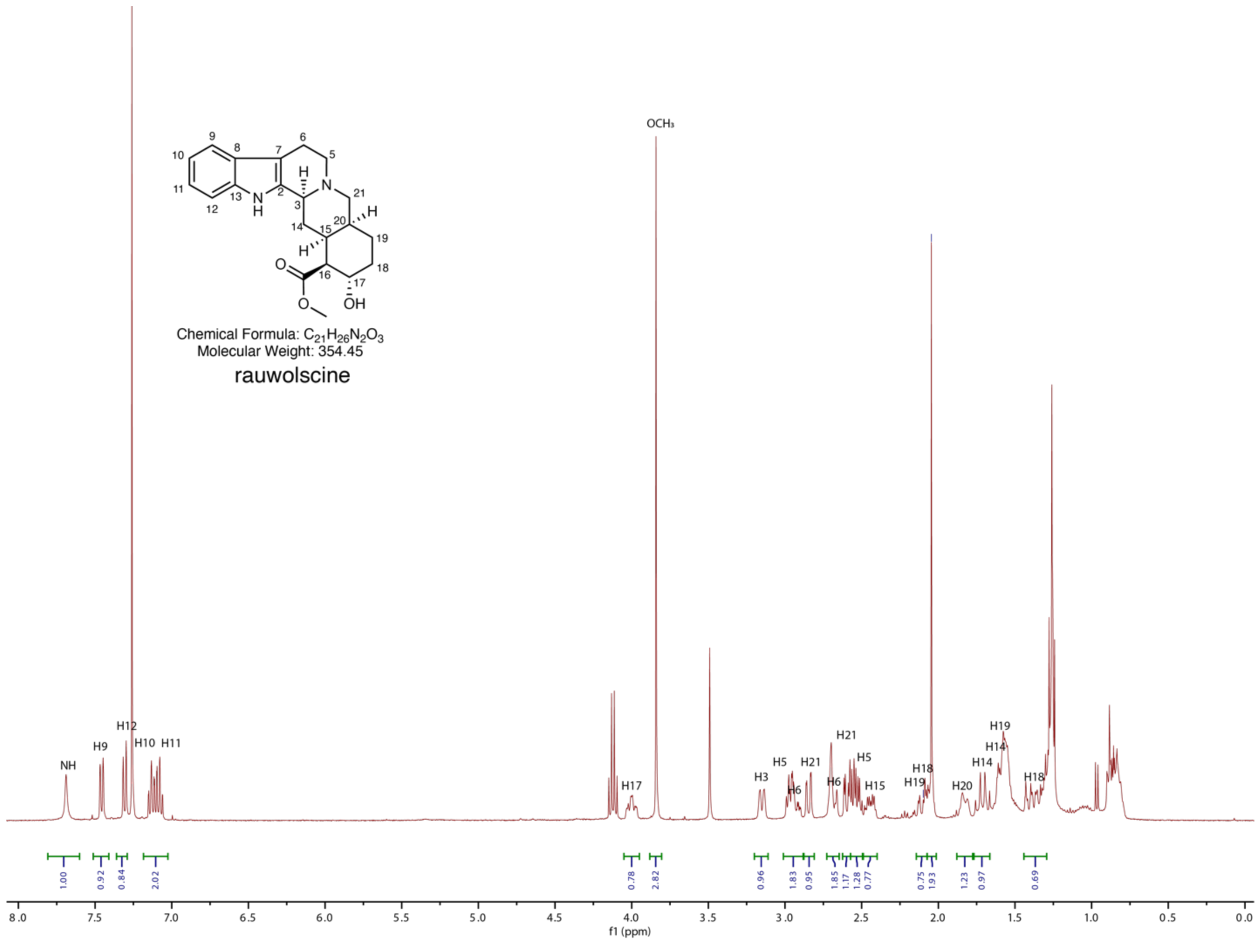

**Supplementary Figure 9. $^1$H NMR spectra of rauwolscine in $CDCl_3$.**

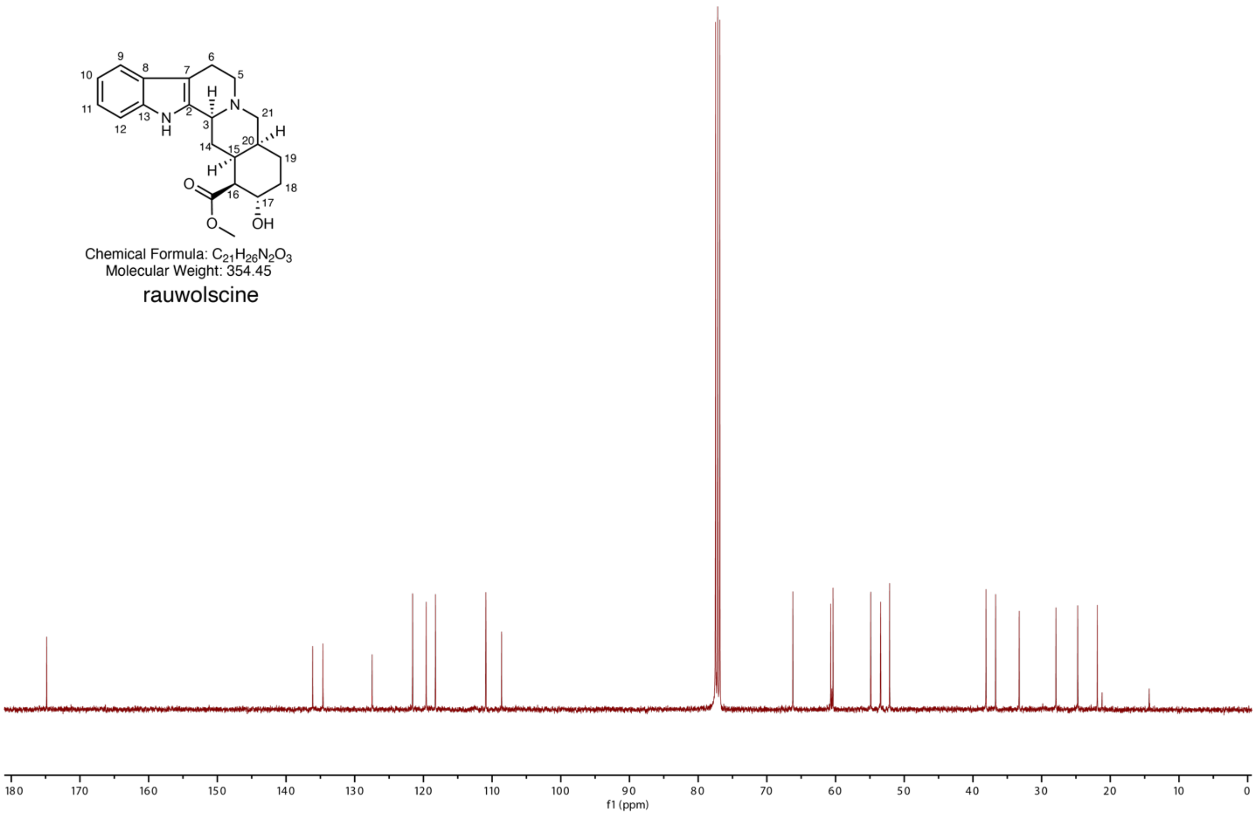

**Supplementary Figure 10. $^{13}$C NMR spectra of rauwolscine in $CDCl_3$.**

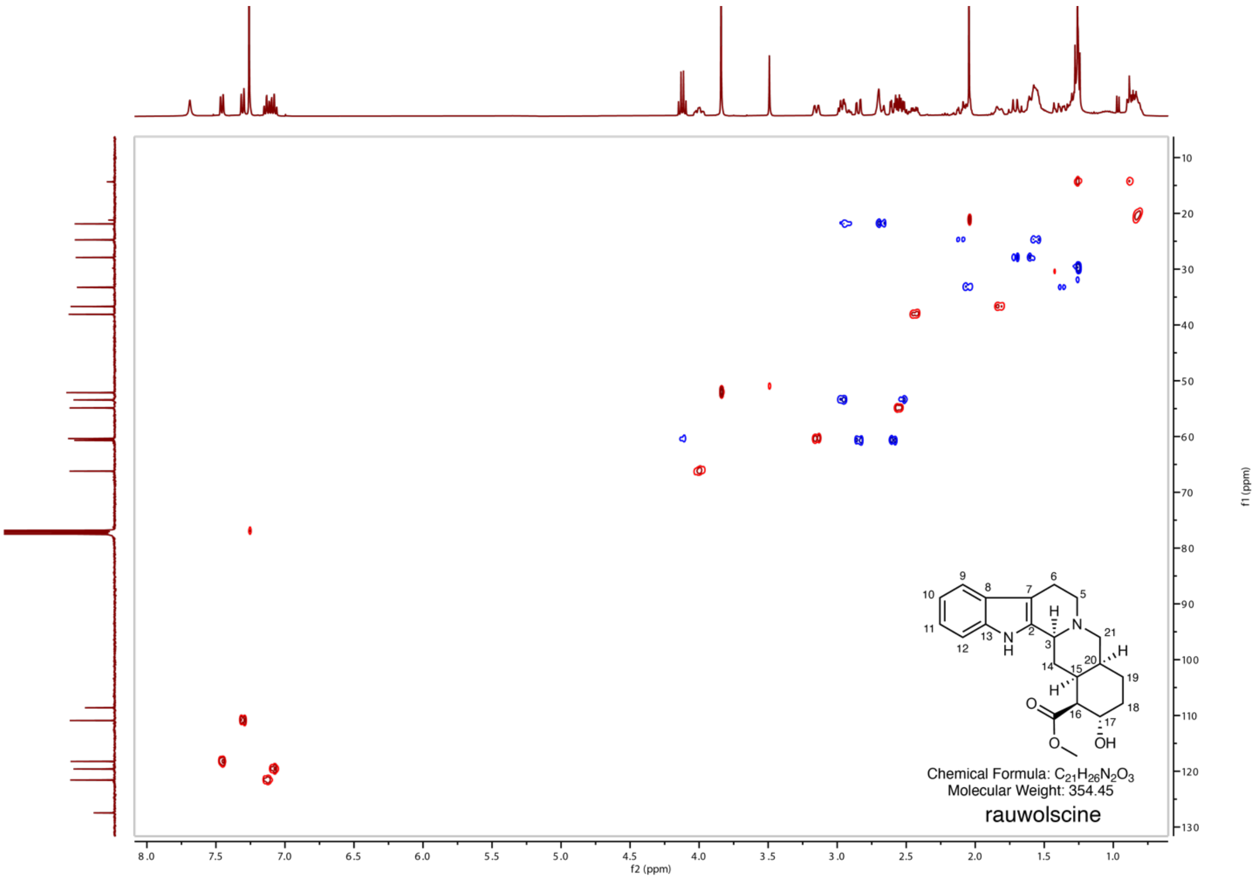

**Supplementary Figure 11. HSQC NMR spectra of rauwolscine in CDCl3.**

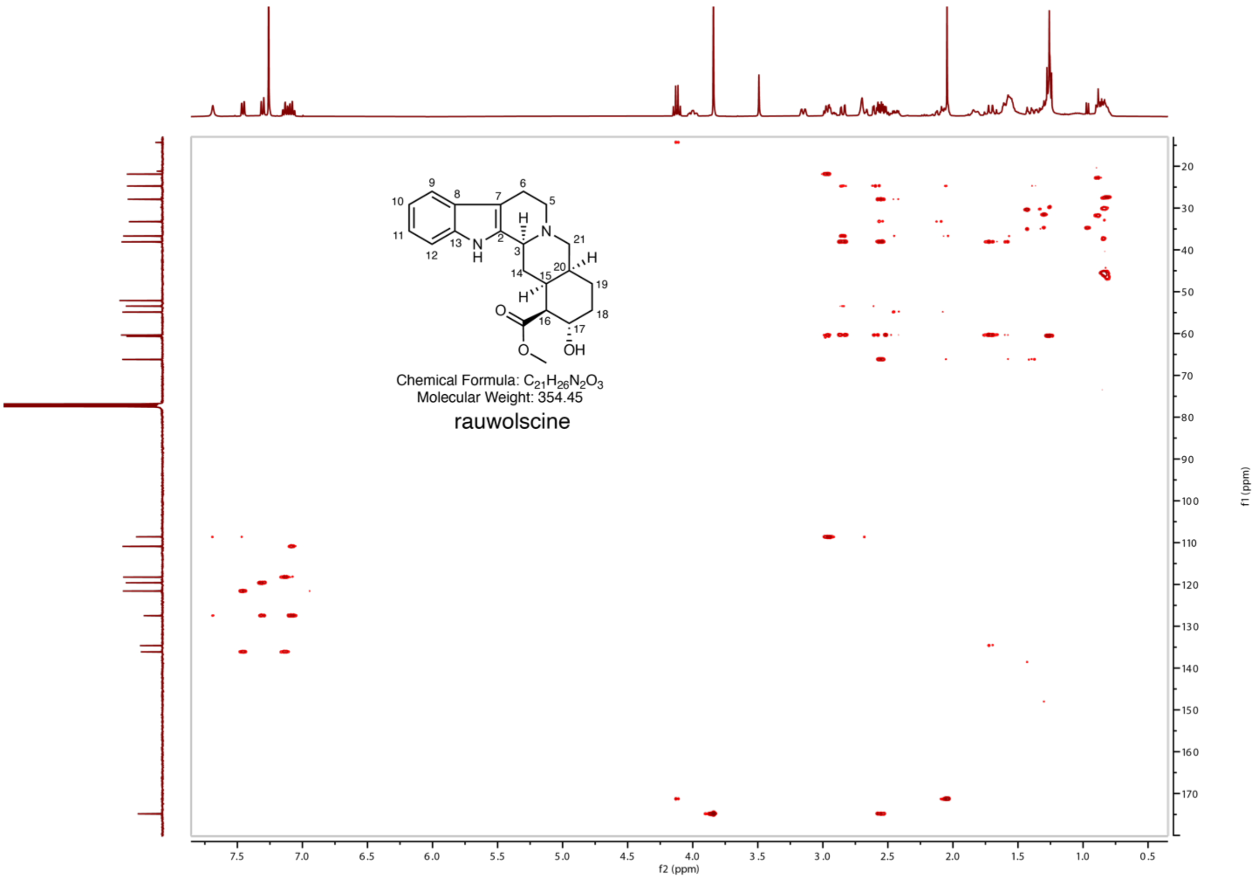

**Supplementary Figure 12. HMBC NMR spectra of rauwolscine in $CDCl_3$.**

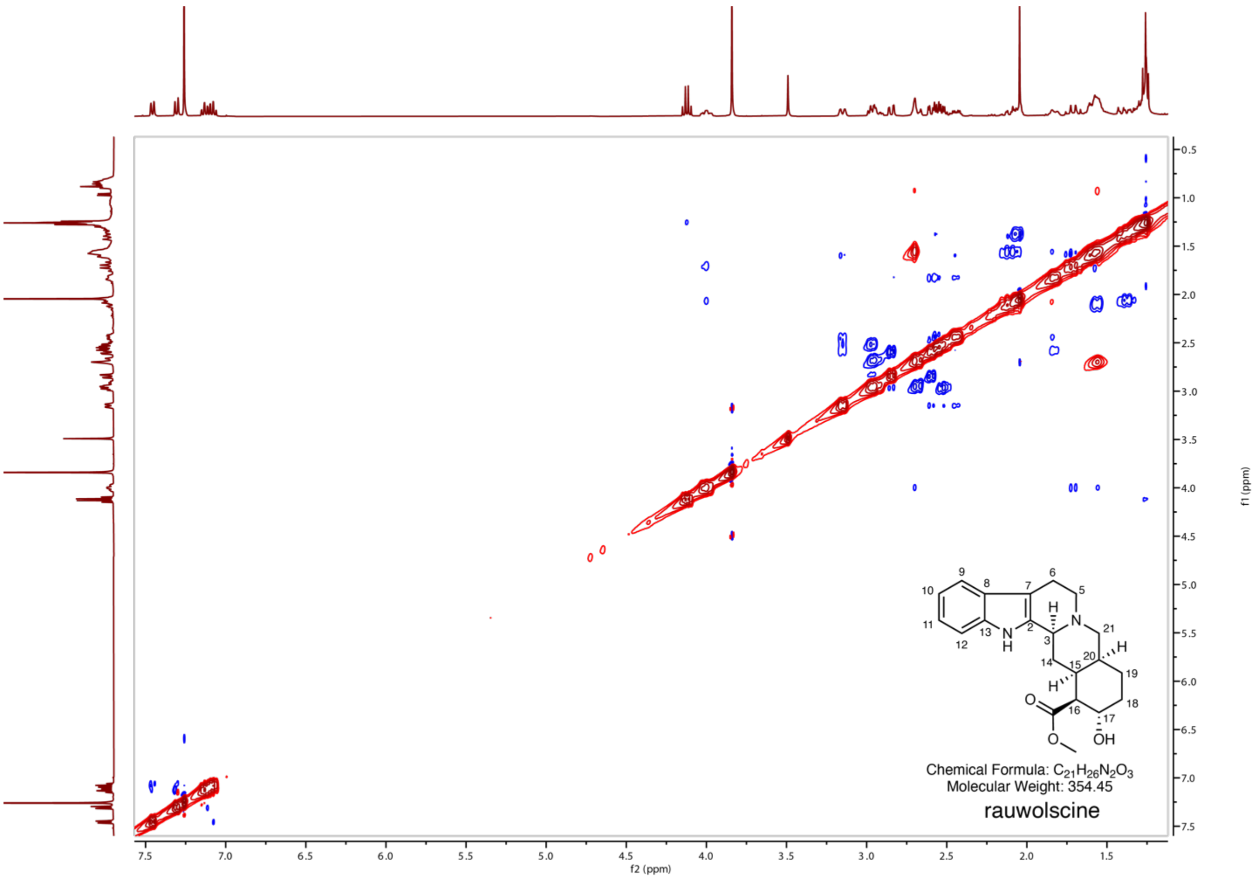

**Supplementary Figure 13. NOESY NMR spectra of rauwolscine in $CDCl_3$.**

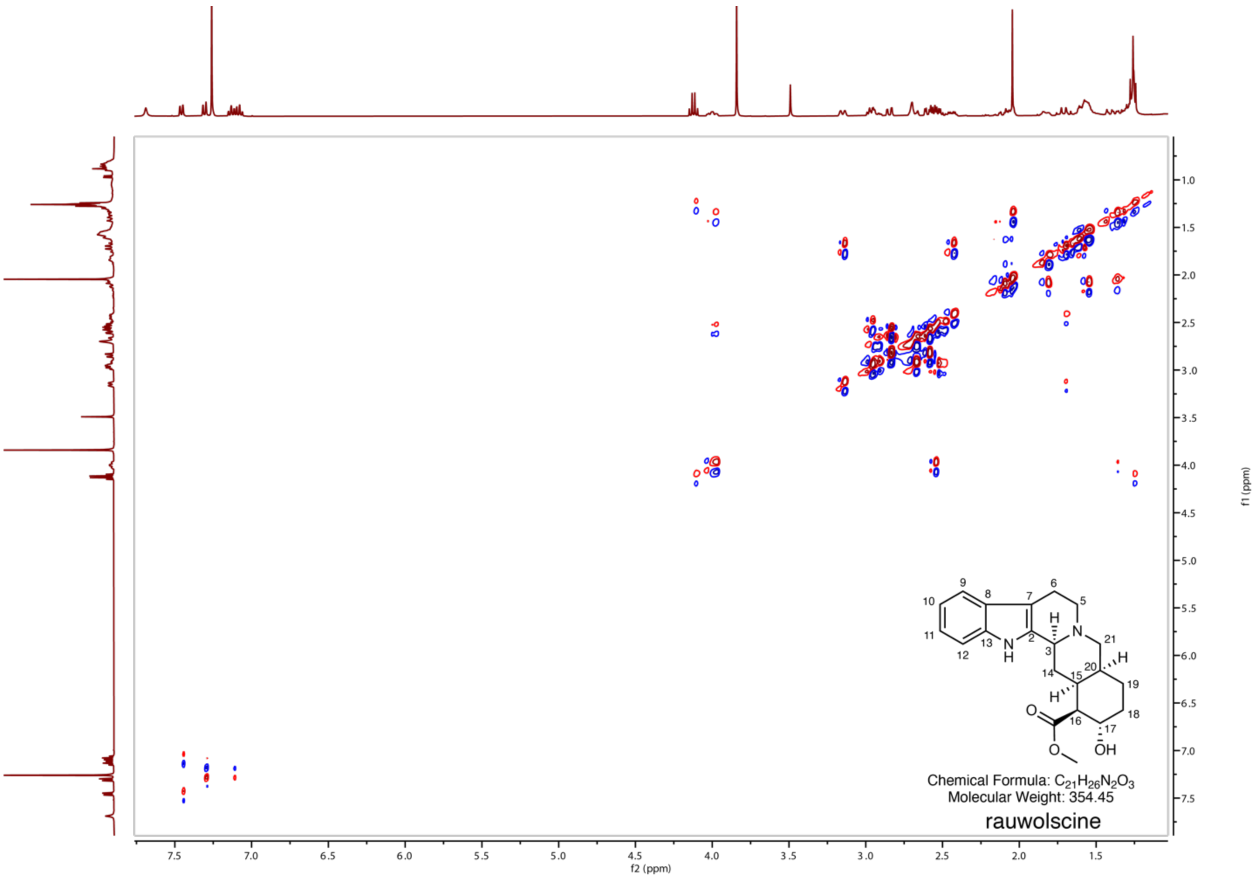

**Supplementary Figure 14. COSY NMR spectra of rauwolscine in $CDCl_3$.**

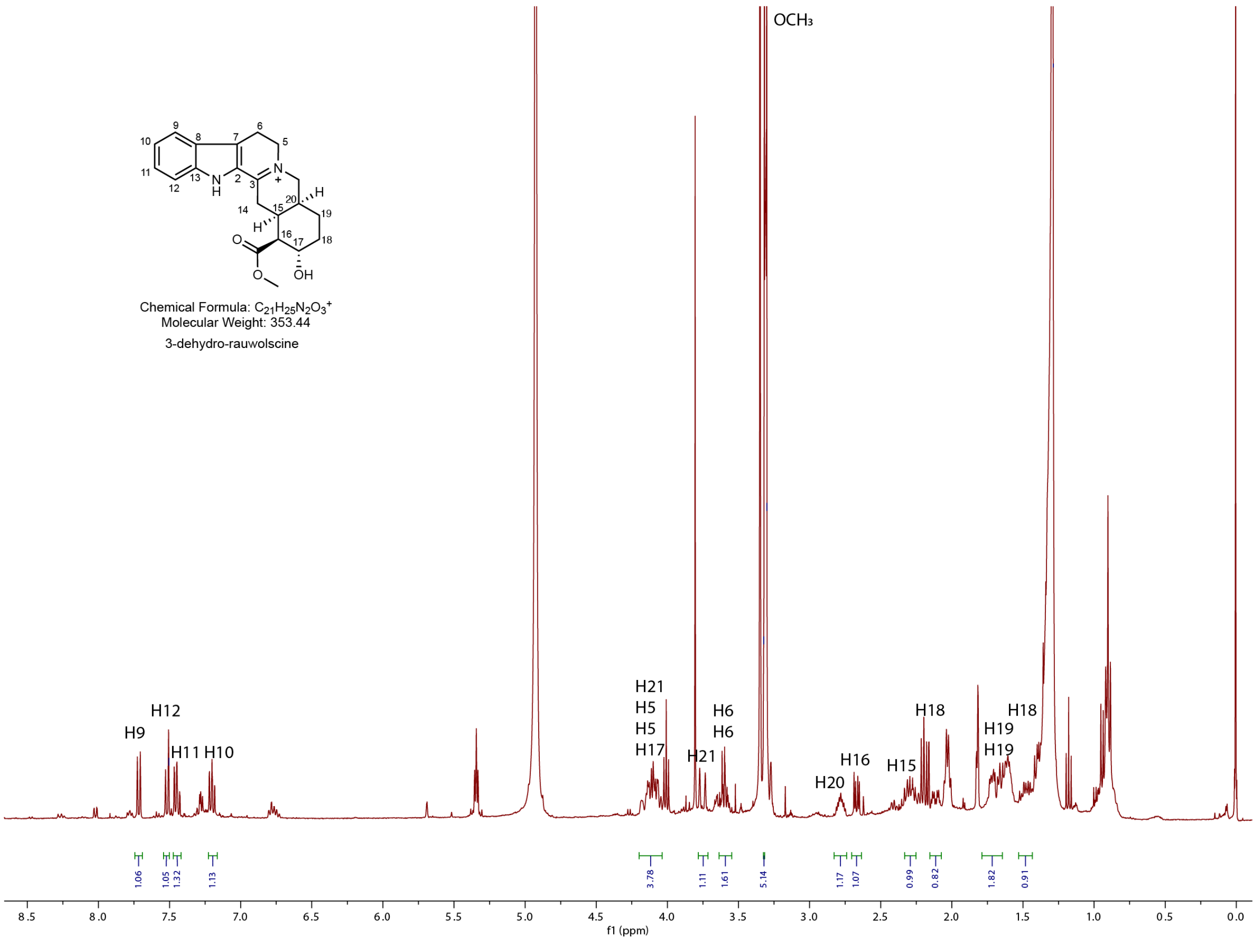

**Supplementary Figure 15. $^{1}$H NMR spectra of 3-dehydro-rauwolscine in MeOD.**
The H14 methylene signals are absent, likely due to tautomeric exchange between the 3,4(*N*)- and 3,14-dehydro forms in MeOD.

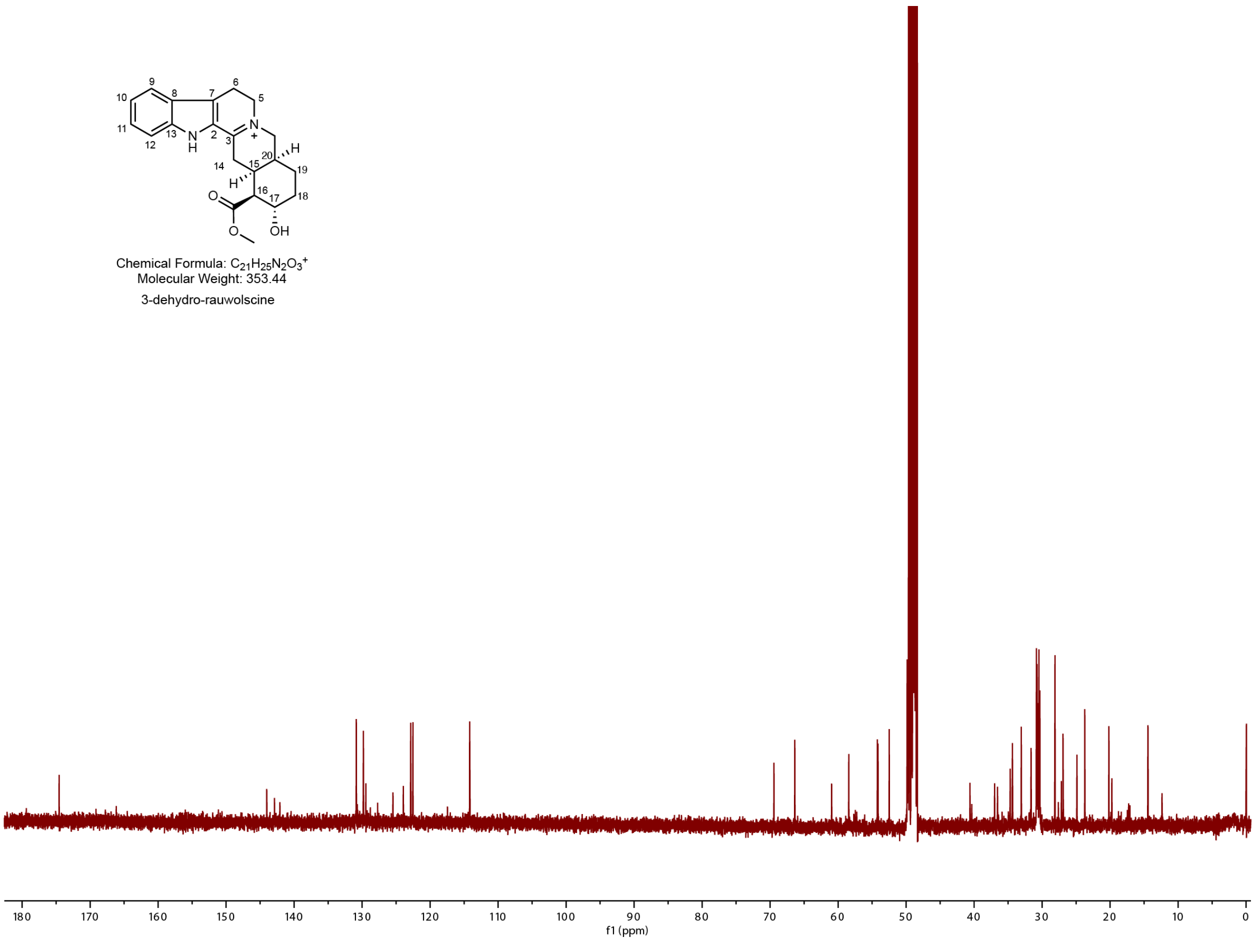

**Supplementary Figure 16. $^{13}C$ NMR spectra of 3-dehydro-rauwolscine in MeOD.**
The C14 methylene signal is absent, likely due to tautomeric exchange between the 3,4(*N*)- and 3,14-dehydro forms in MeOD.

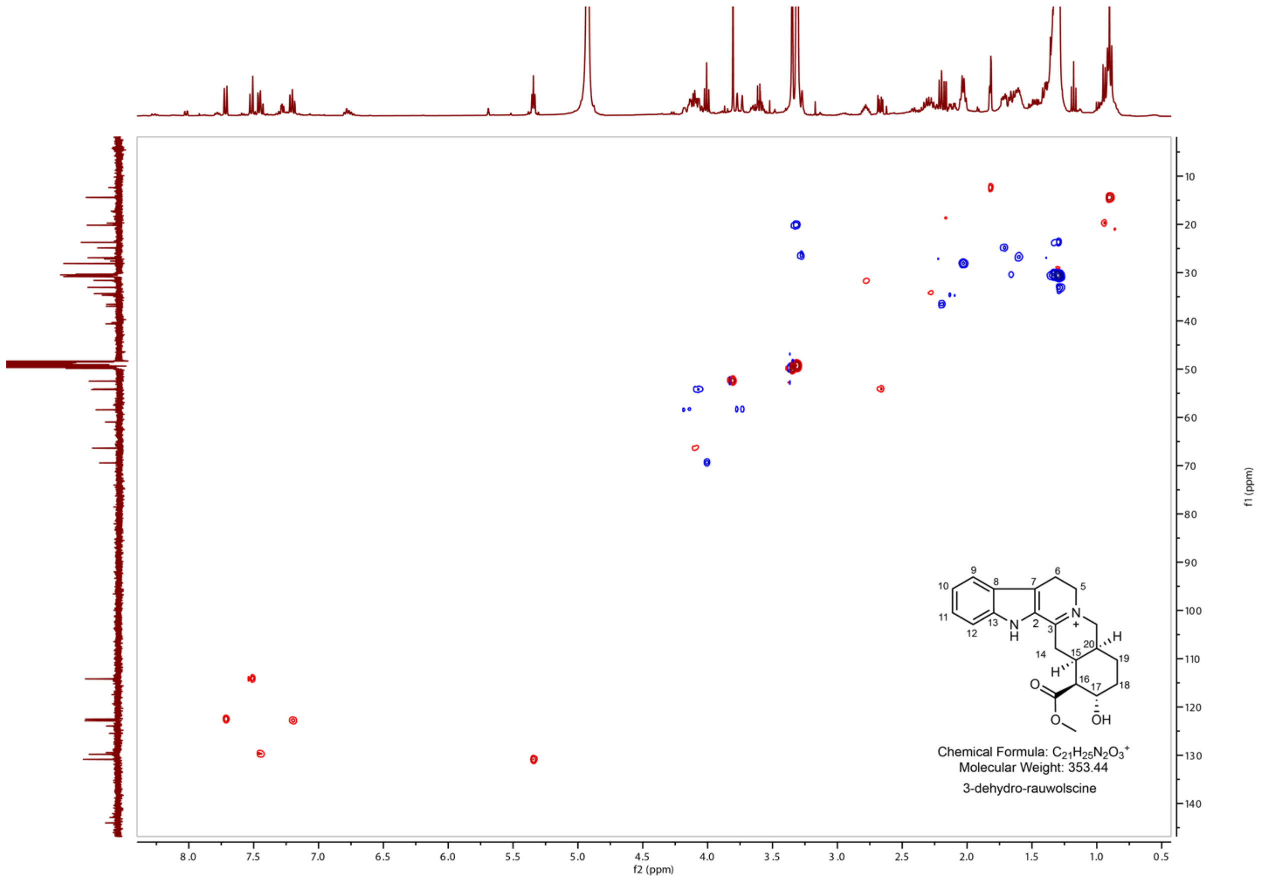

**Supplementary Figure 17. HSQC NMR spectra of 3-dehydro-rauwolscine in MeOD.**

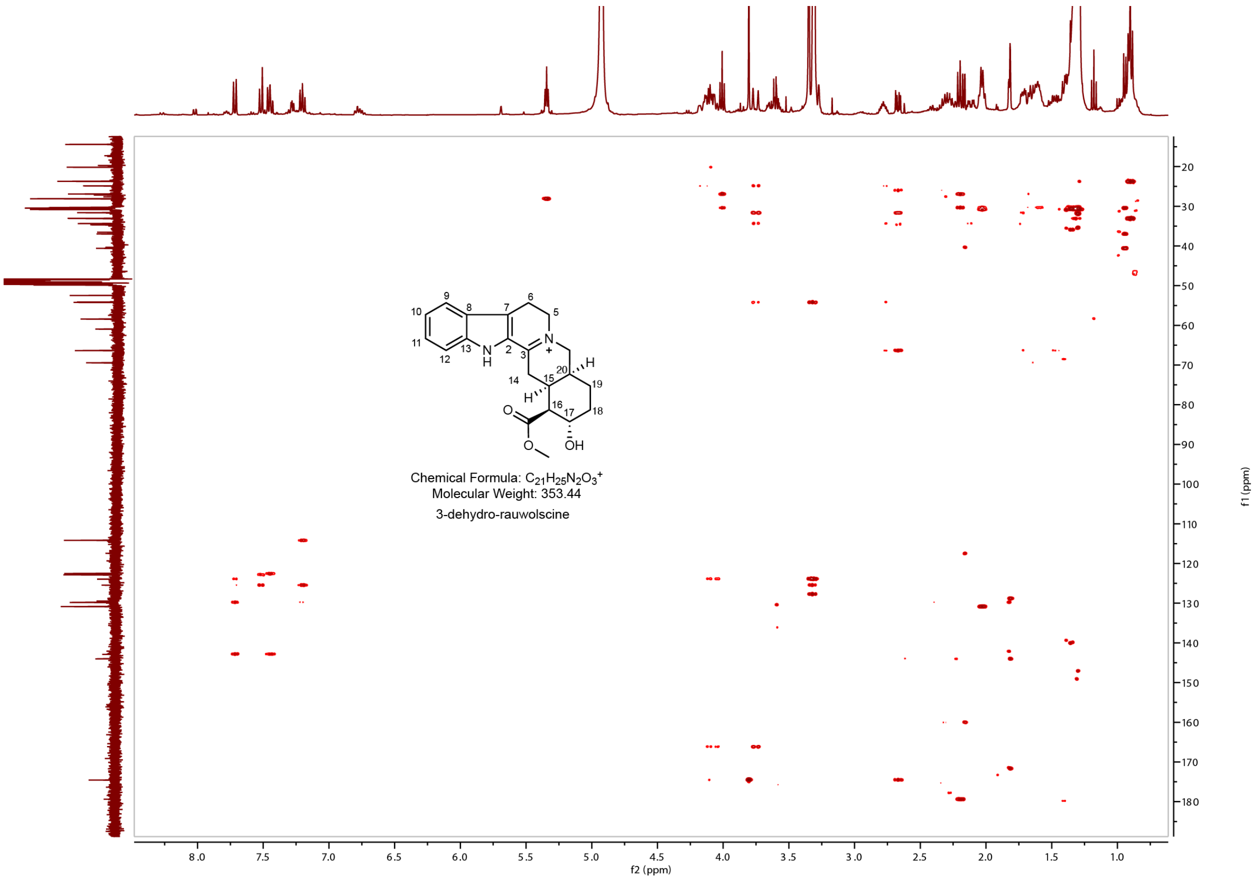

**Supplementary Figure 18. HMBC NMR spectra of 3-dehydro-rauwolscine in MeOD.**

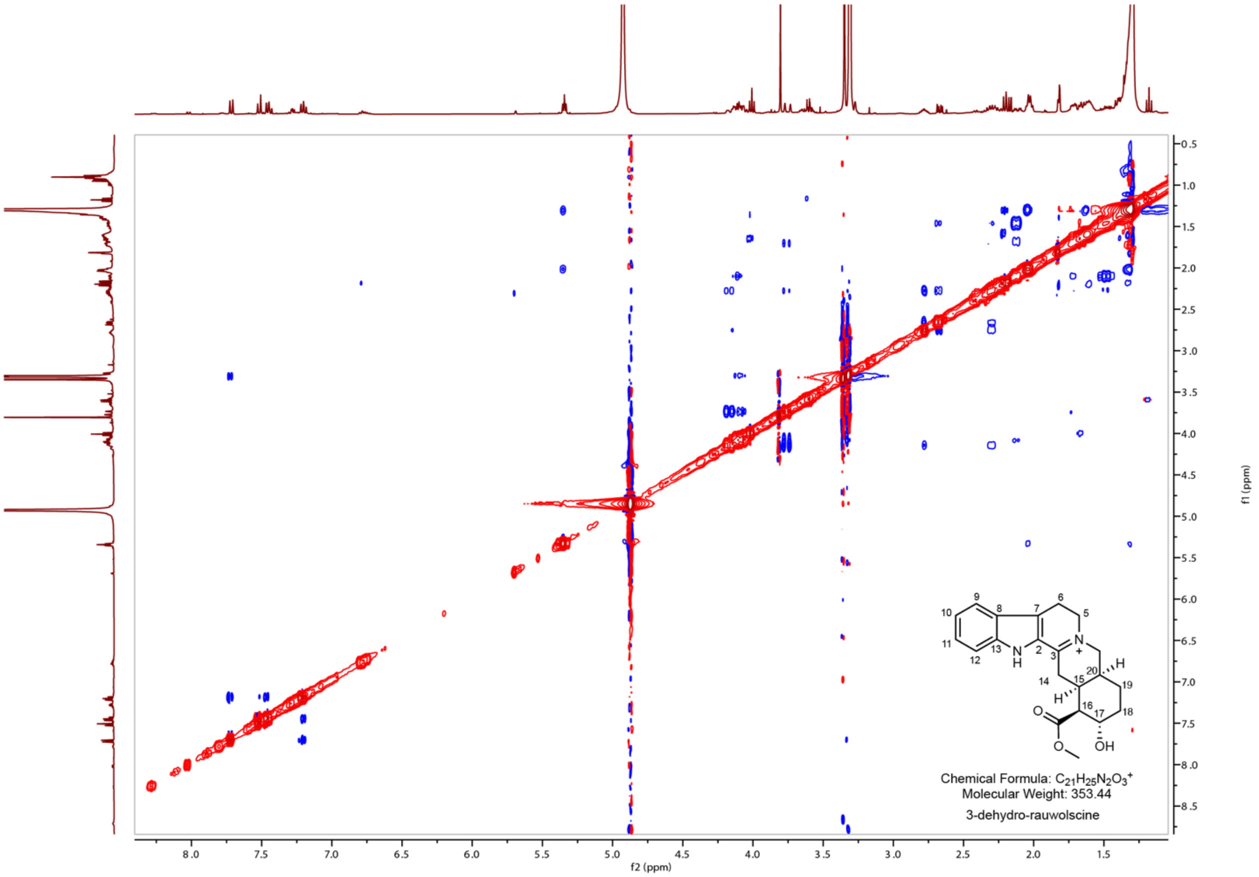

**Supplementary Figure 19. NOESY NMR spectra of 3-dehydro-rauwolscine in MeOD.**

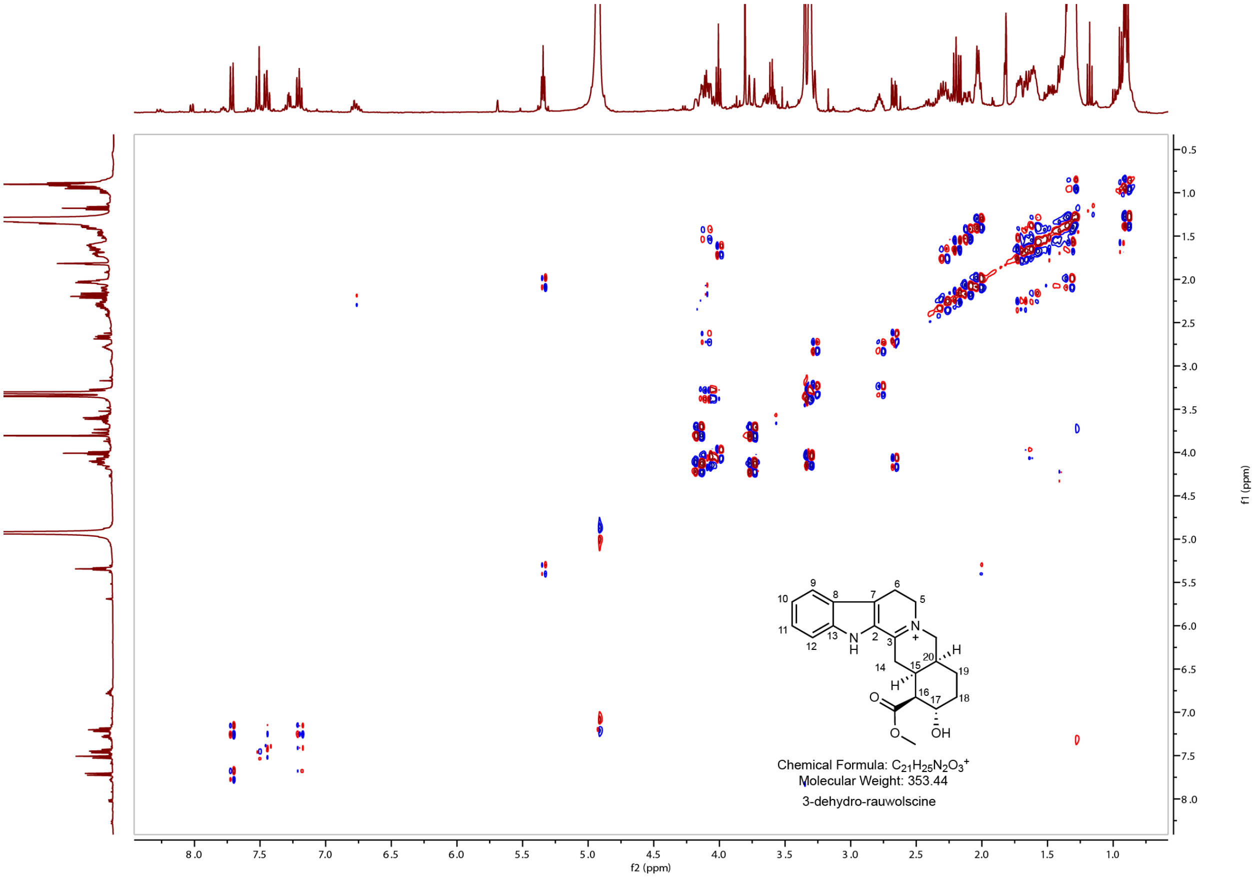

**Supplementary Figure 20. COSY NMR spectra of 3-dehydro-rauwolscine in MeOD.**

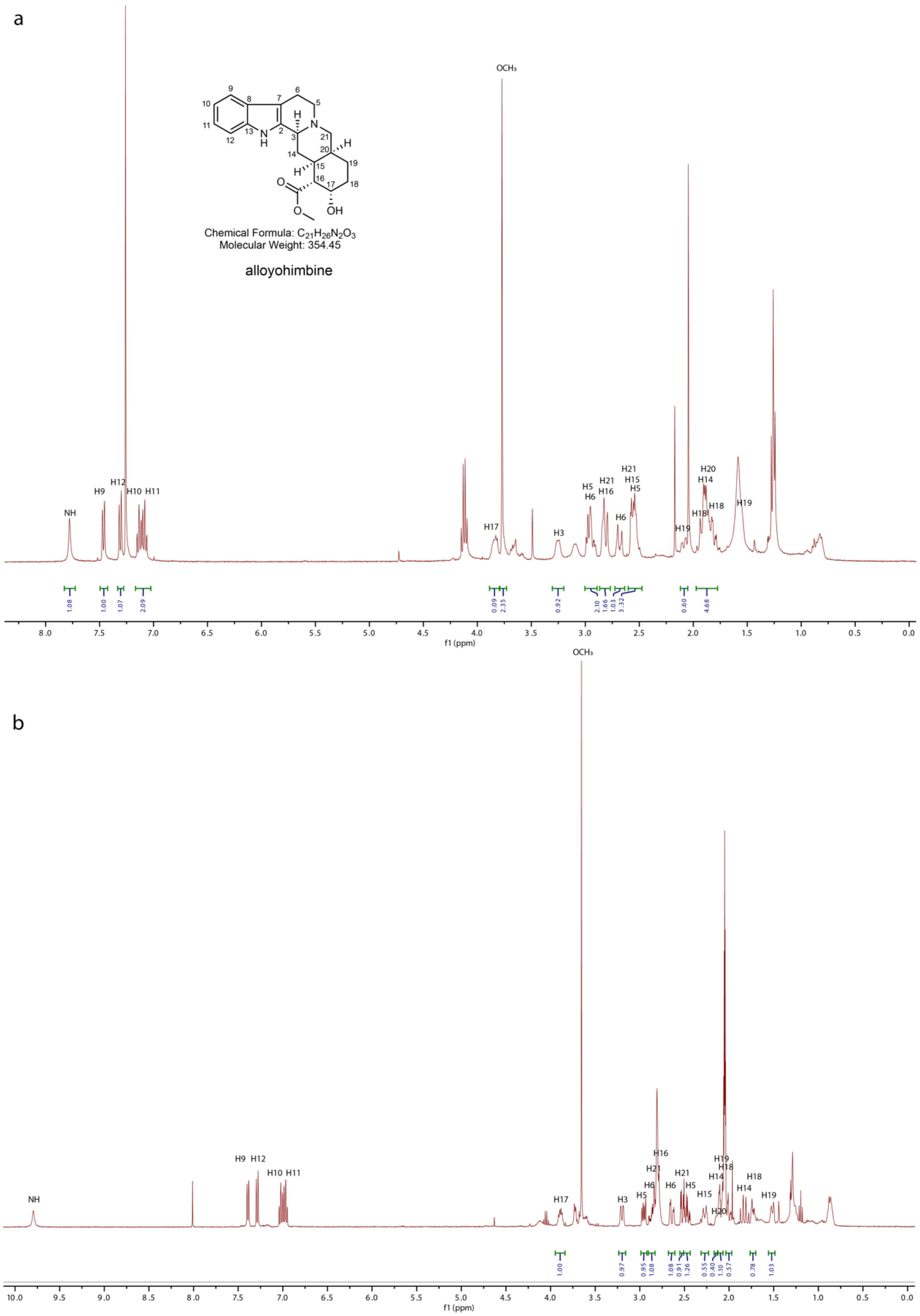

**Supplementary Figure 21. $^1$H NMR spectra of alloyohimbine in $CDCl_3$ (a) & acetone-*d6* (b).**

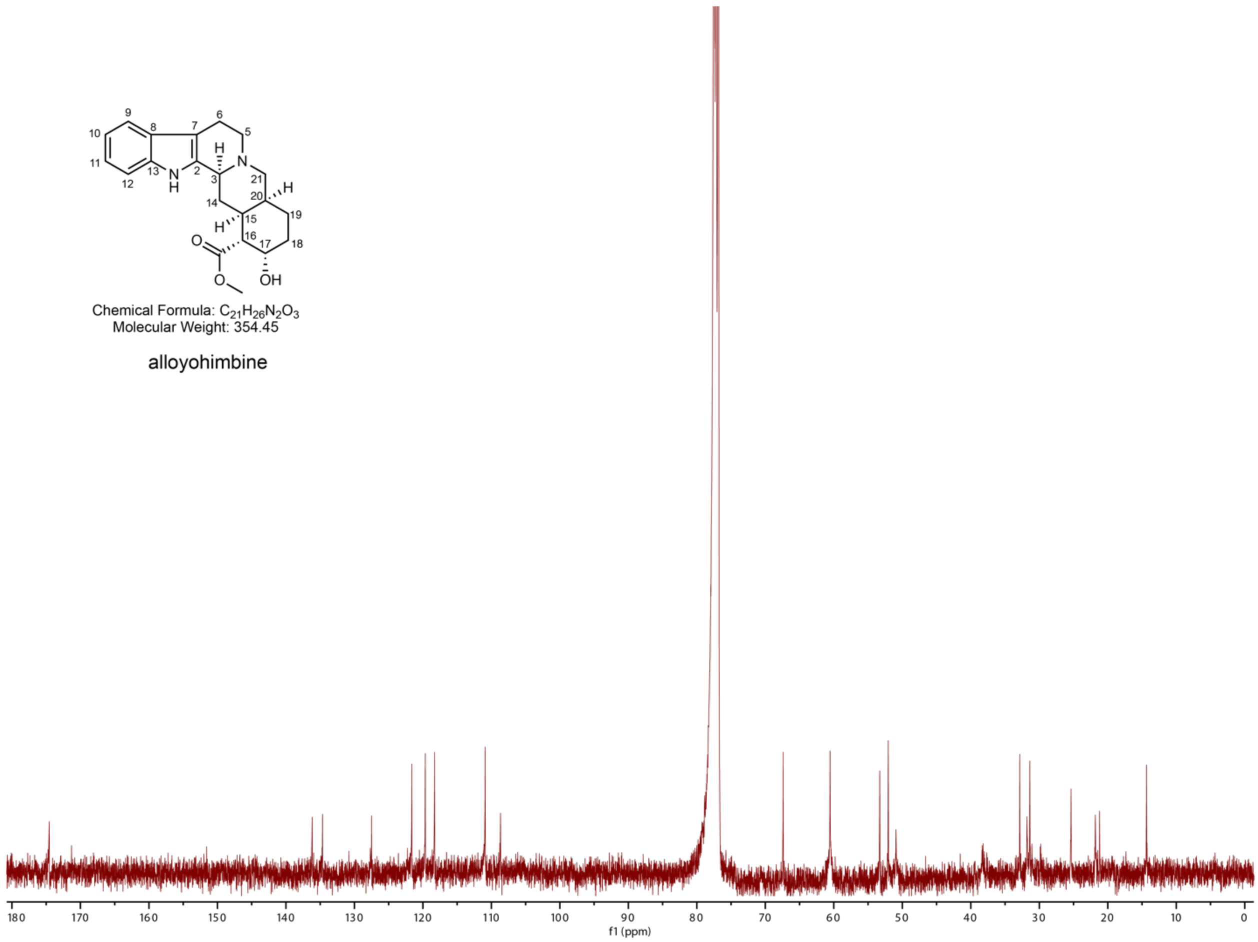

**Supplementary Figure 22. $^{13}C$ NMR spectra of alloyohimbine in $CDCl_3$.**

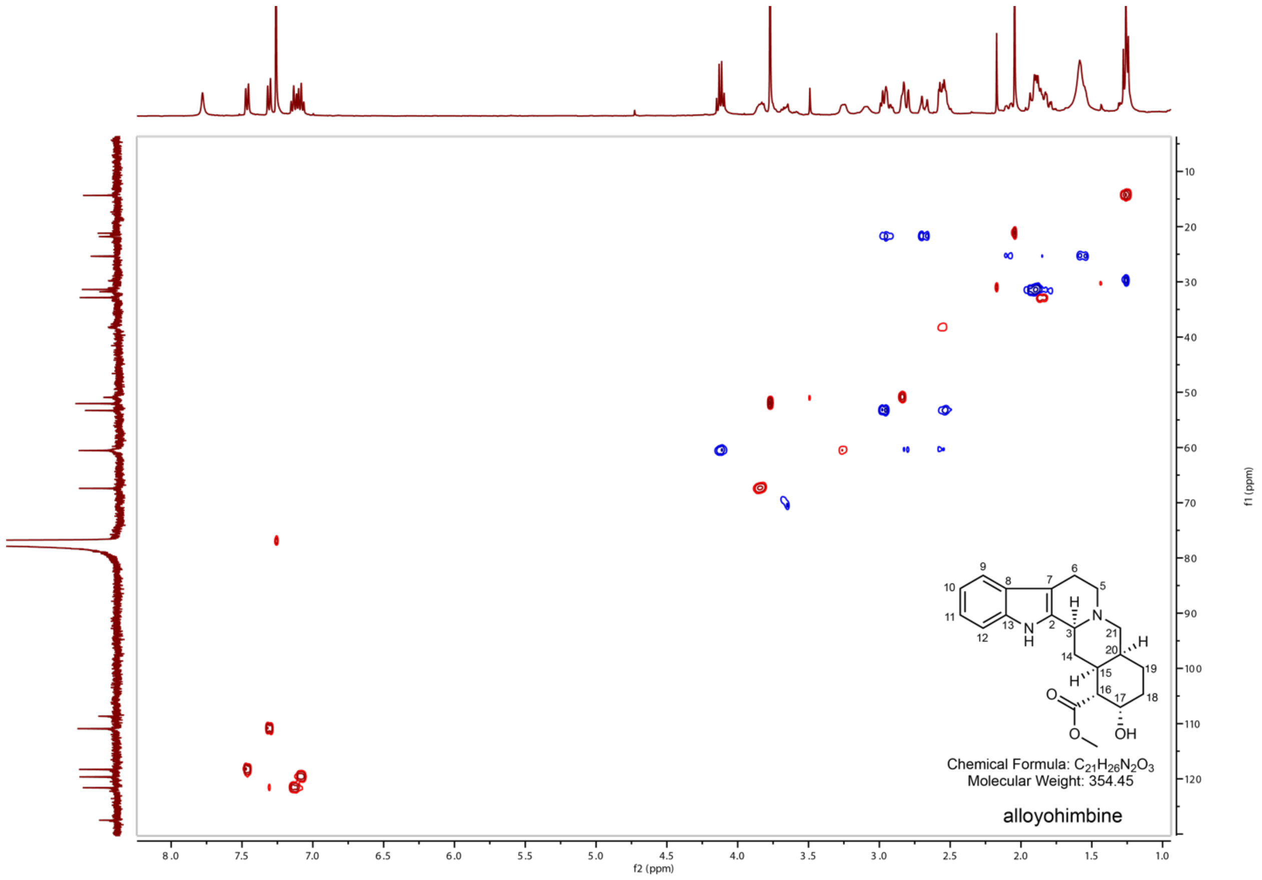

**Supplementary Figure 23. HSQC NMR spectra of alloyohimbine in $CDCl_3$.**

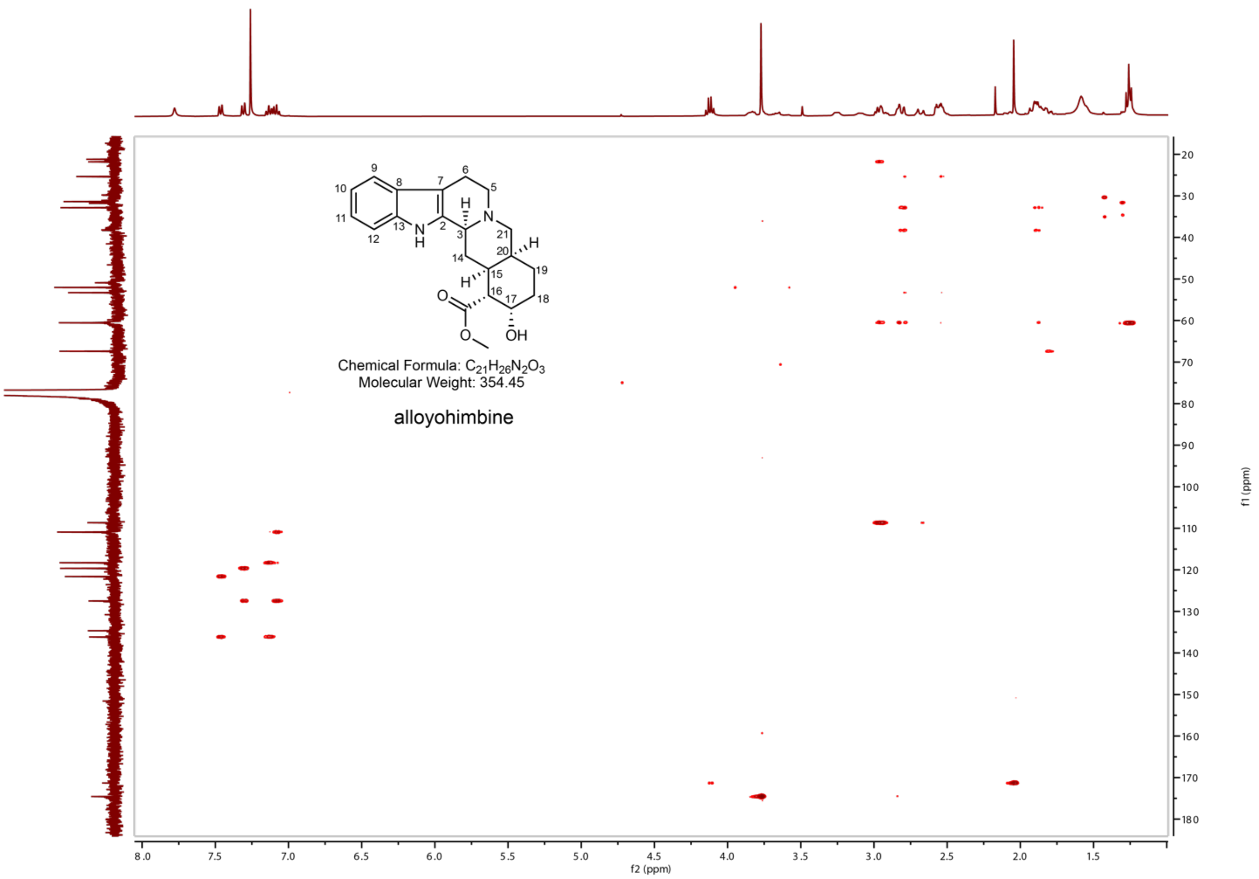

**Supplementary Figure 24. HMBC NMR spectra of alloyohimbine in $CDCl_3$.**

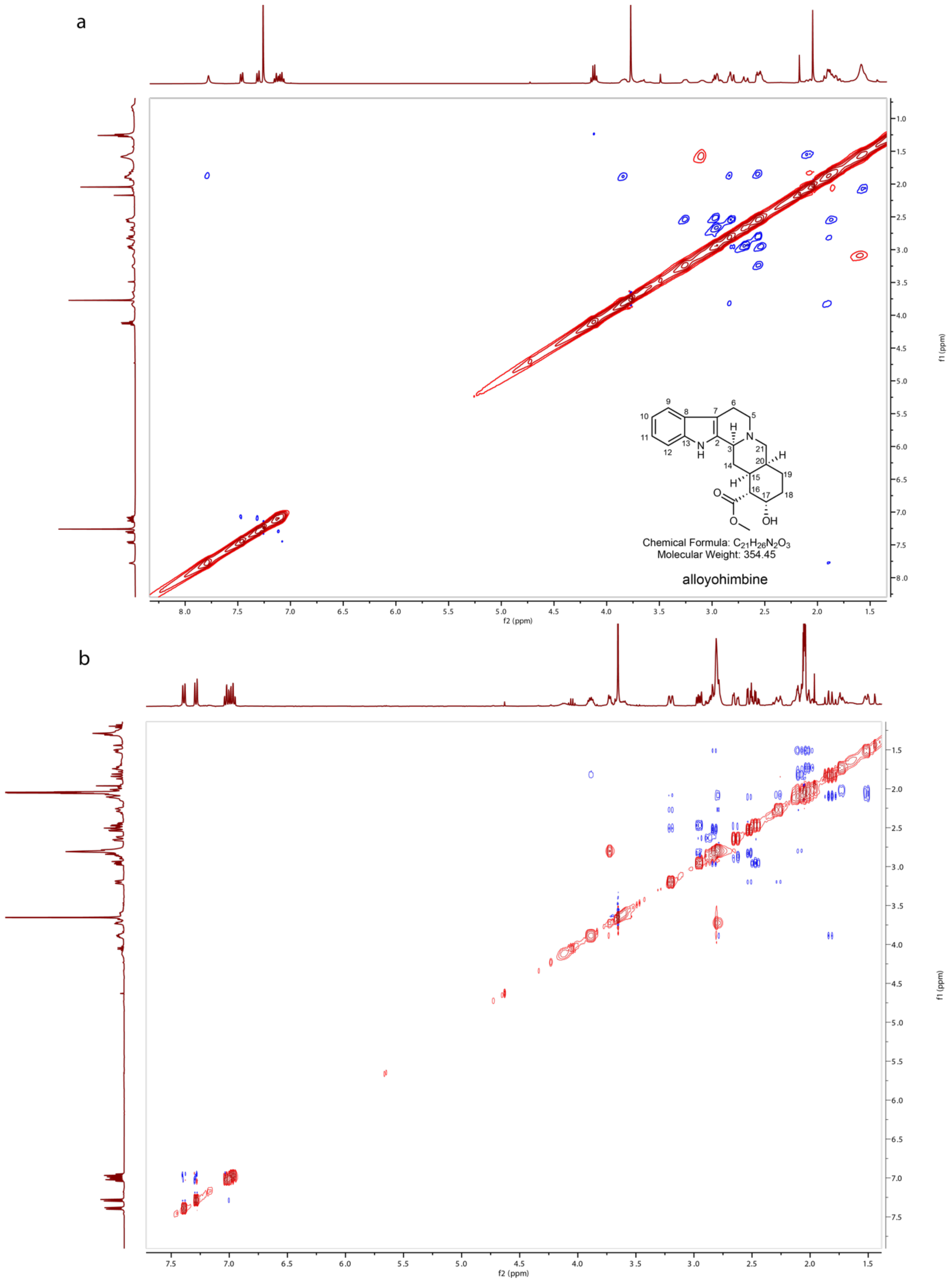

**Supplementary Figure 25. NOESY NMR spectra of alloyohimbine in $CDCl_3$ (a) and acetone-*d6* (b).**

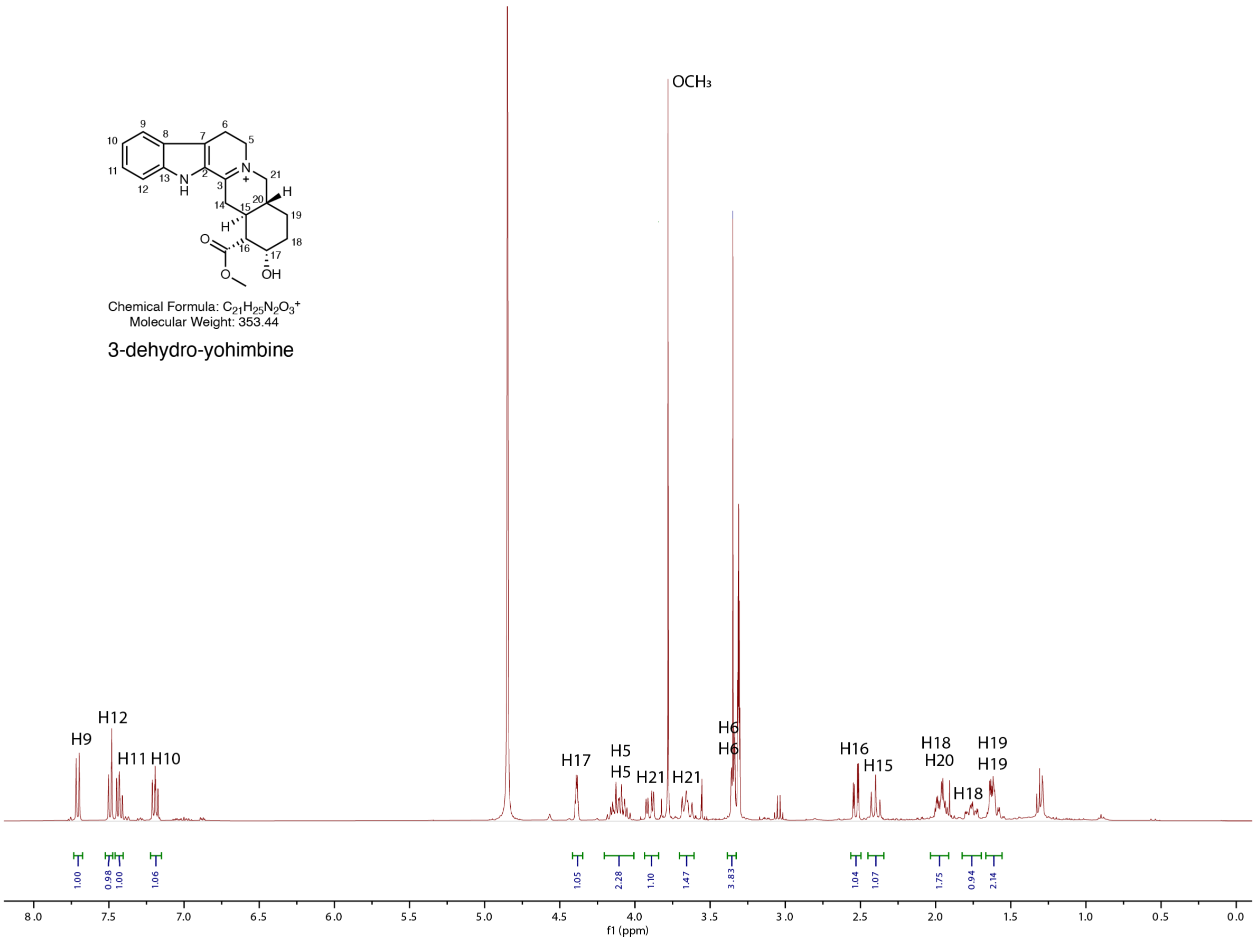

**Supplementary Figure 26. $^1$H NMR spectra of 3-dehydro-yohimbine in MeOD.**
The H14 methylene signals are absent, likely due to tautomeric exchange between the 3,4(*N*)- and 3,14-dehydro forms in MeOD.

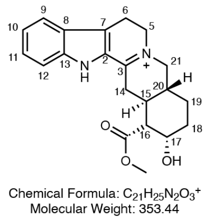

3-dehydro-yohimbine

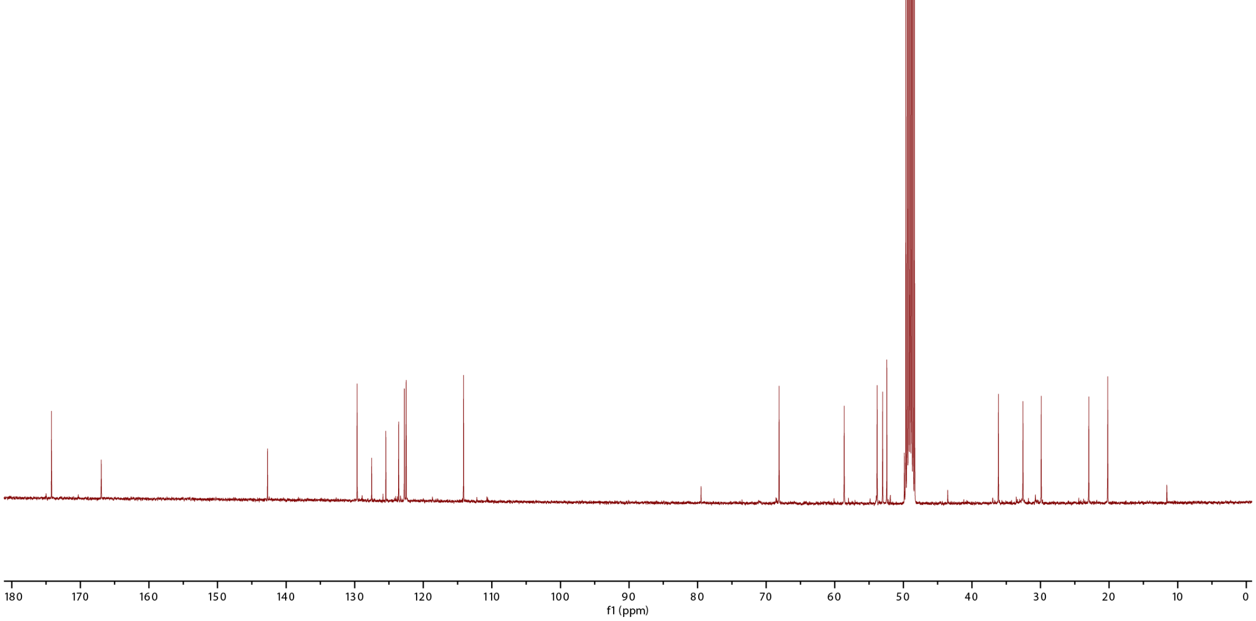

**Supplementary Figure 27. $^{13}C$ NMR spectra of 3-dehydro- yohimbine in MeOD.**
The C14 methylene signal is absent, likely due to tautomeric exchange between the 3,4(*N*)- and 3,14-dehydro forms in MeOD.

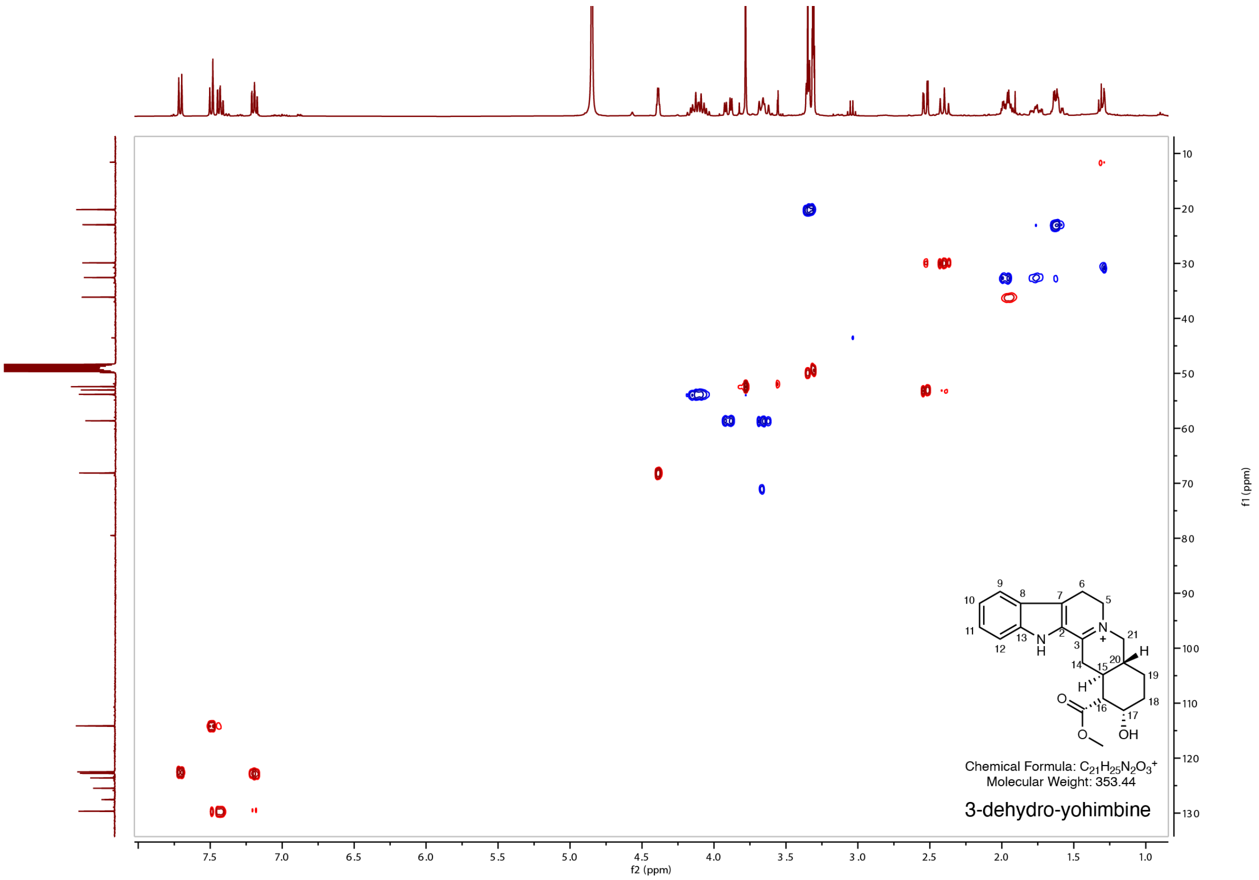

**Supplementary Figure 28. HSQC NMR spectra of 3-dehydro-yohimbine in MeOD.**

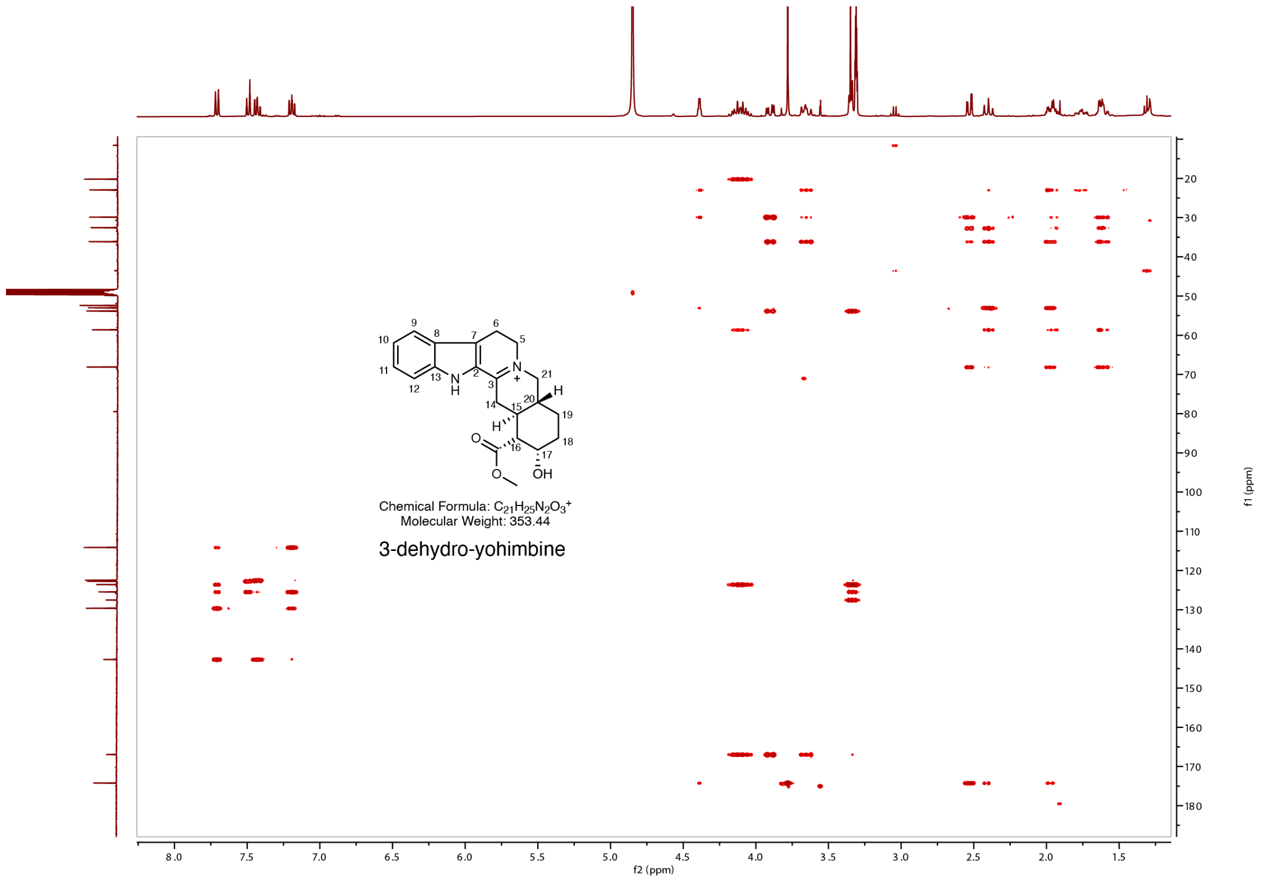

**Supplementary Figure 29. HMBC NMR spectra of 3-dehydro-yohimbine in MeOD.**

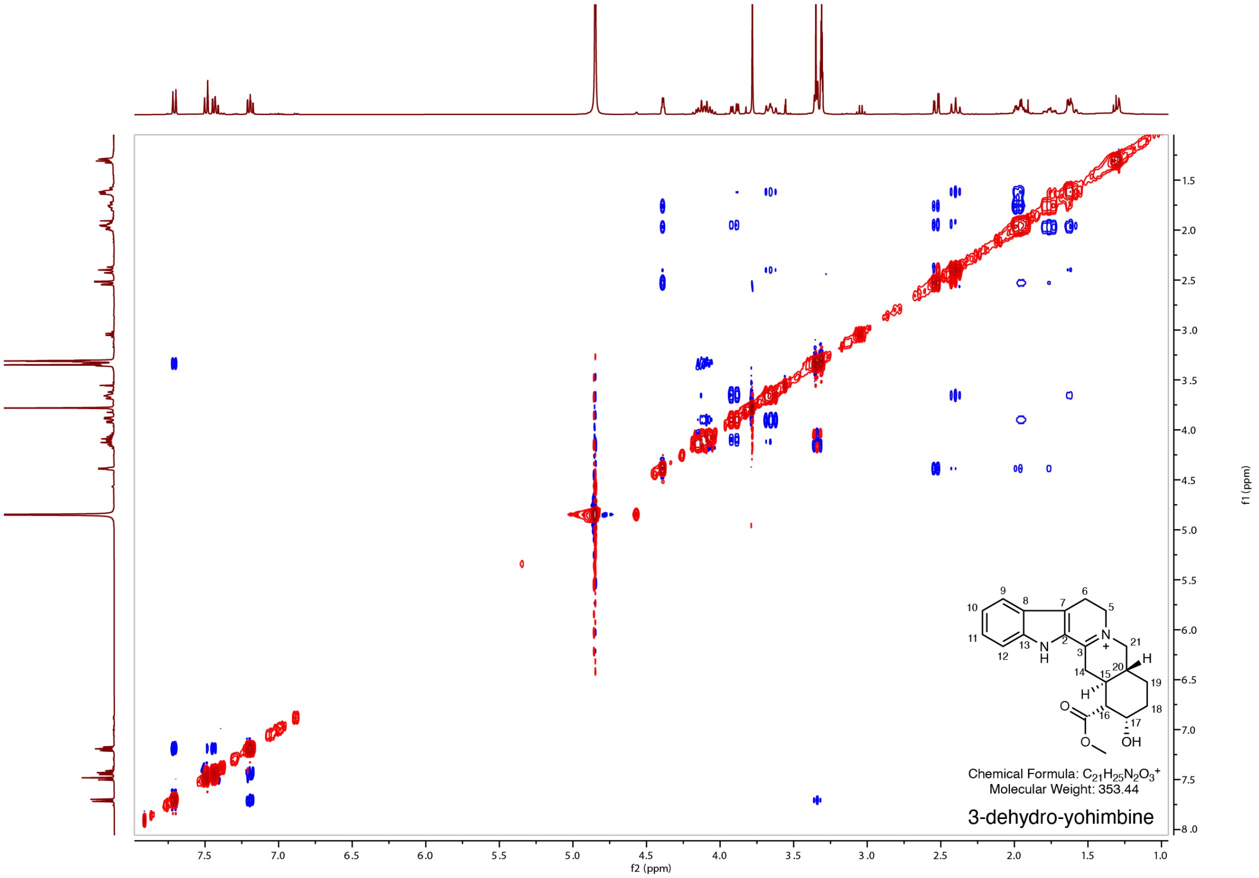

**Supplementary Figure 30. NOESY NMR spectra of 3-dehydro-yohimbine in MeOD.**

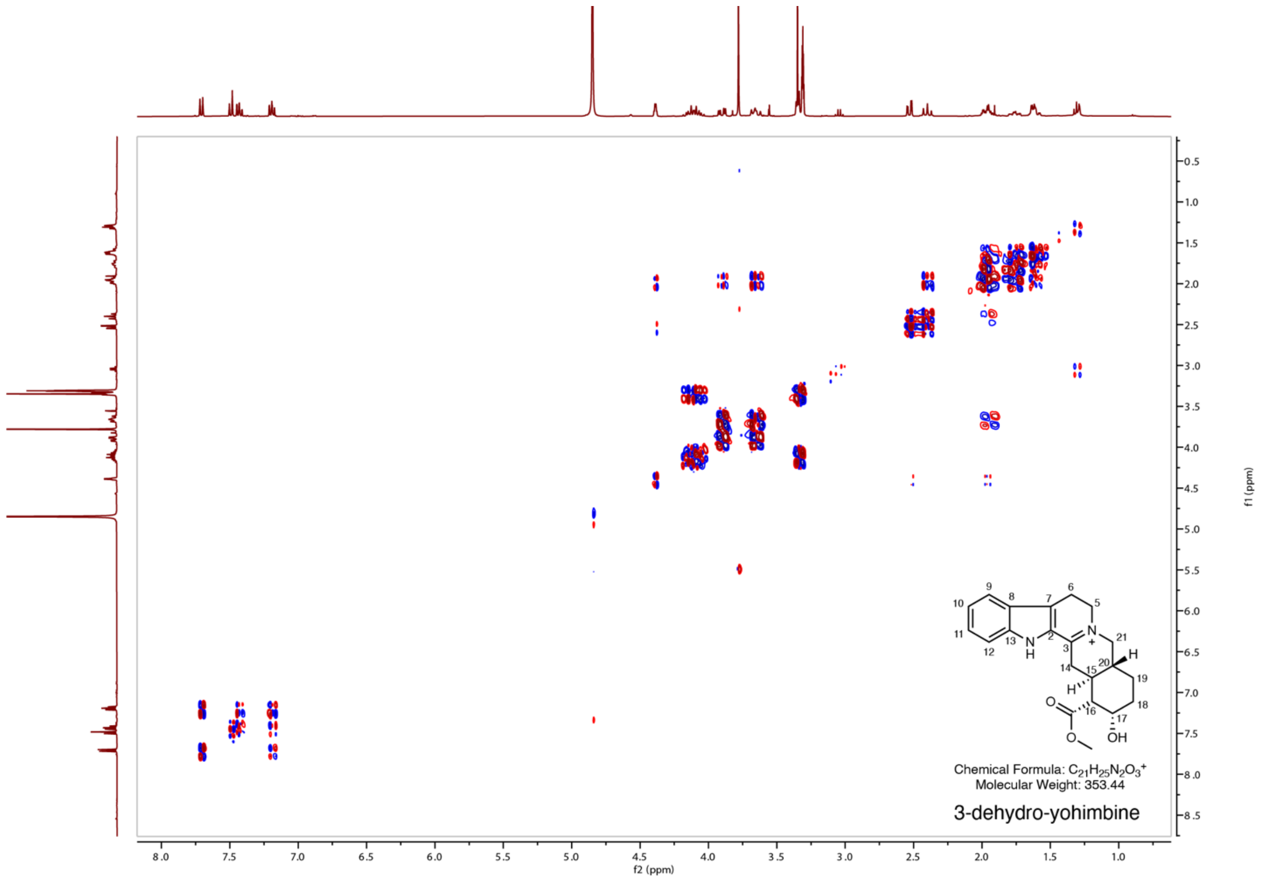

**Supplementary Figure 31. COSY NMR spectra of 3-dehydro-yohimbine in MeOD.**

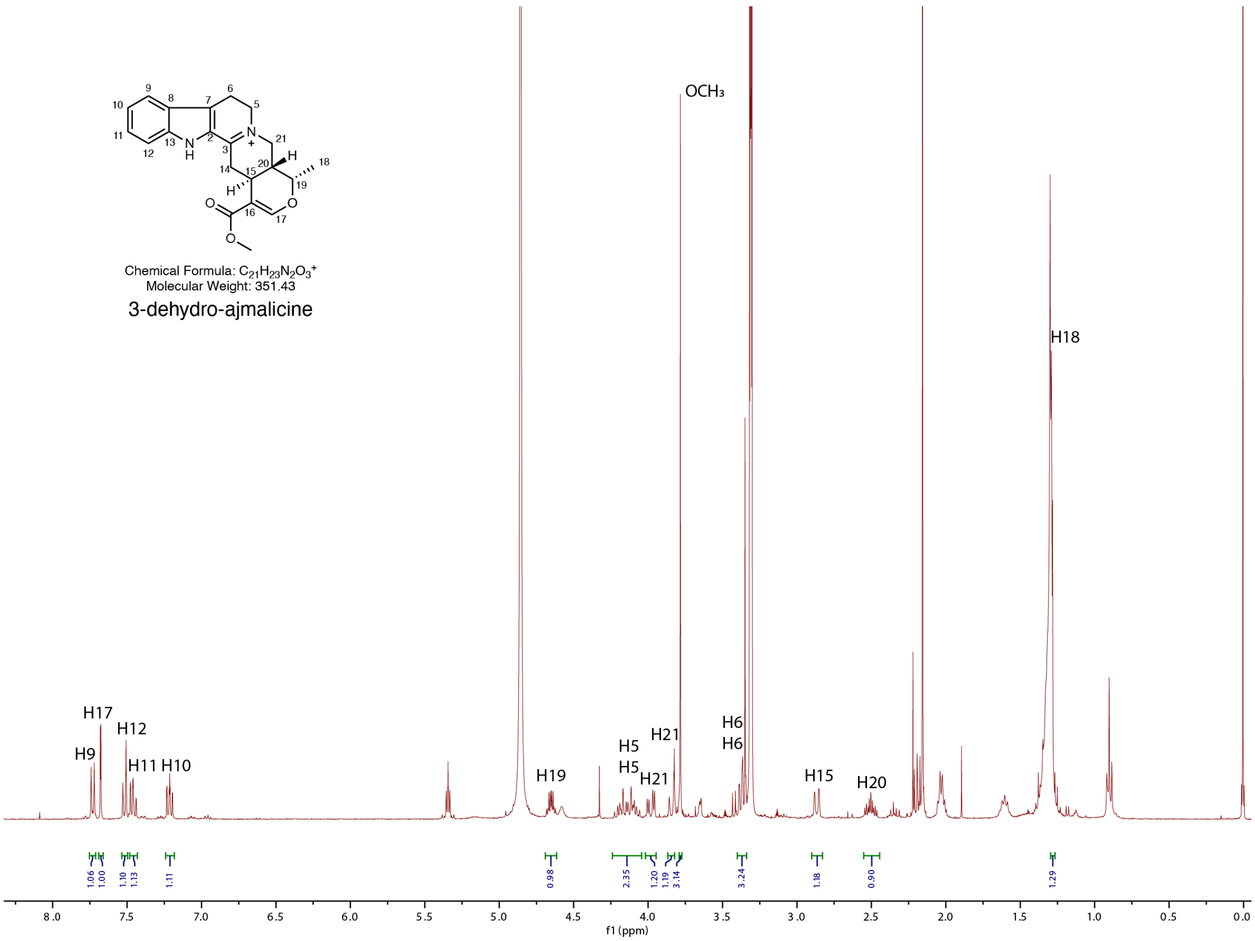

**Supplementary Figure 32. [1]H NMR spectra of 3-dehydro-ajmalicine in MeOD.**
The H14 methylene signals are absent, likely due to tautomeric exchange between the 3,4(*N*)- and 3,14-dehydro forms in MeOD.

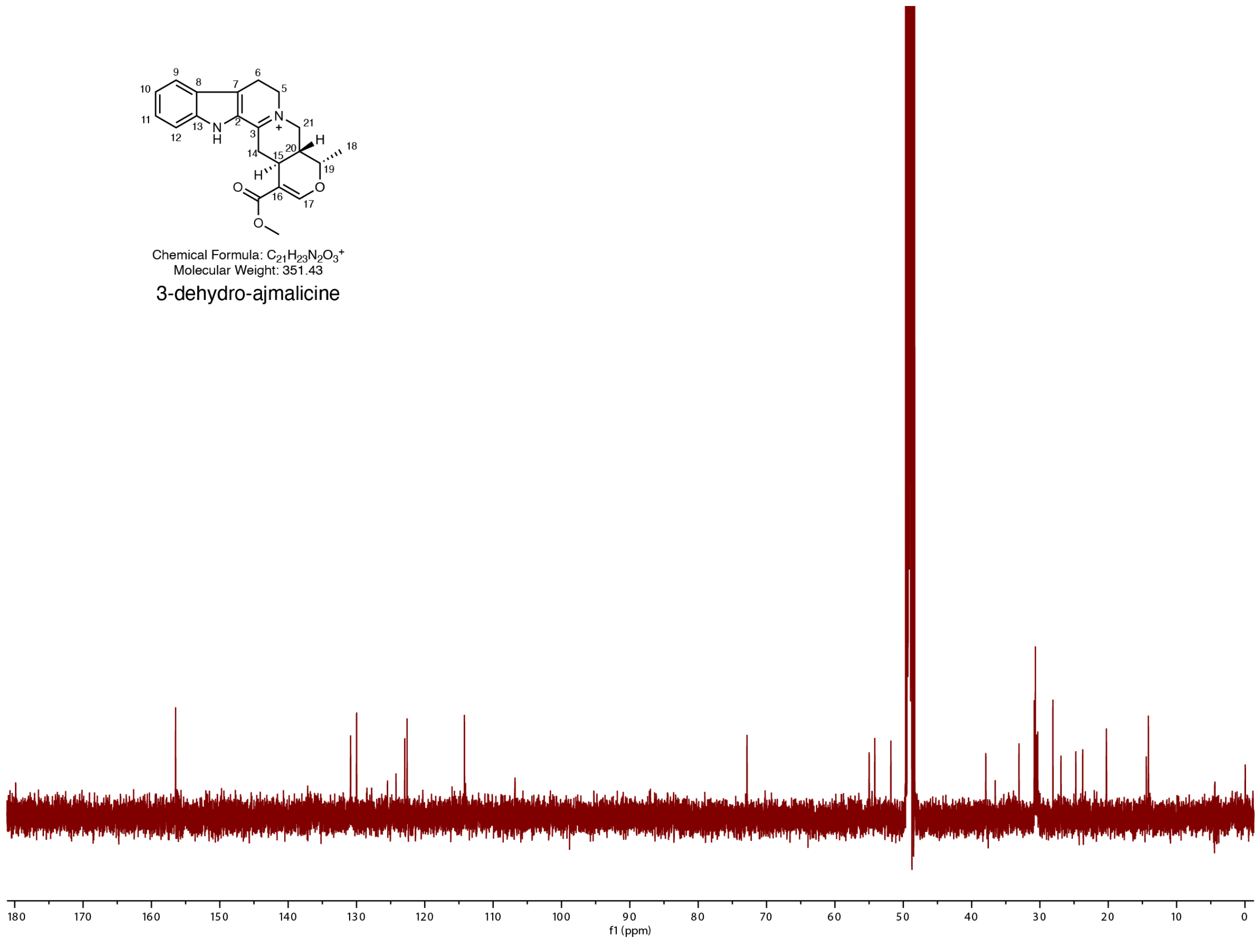

**Supplementary Figure 33. $^{13}$C NMR spectra of 3-dehydro-ajmalicine in MeOD.**
The C3, and C14 signals are absent, likely due to tautomeric exchange between the 3,4(*N*)- and 3,14-dehydro forms in MeOD.

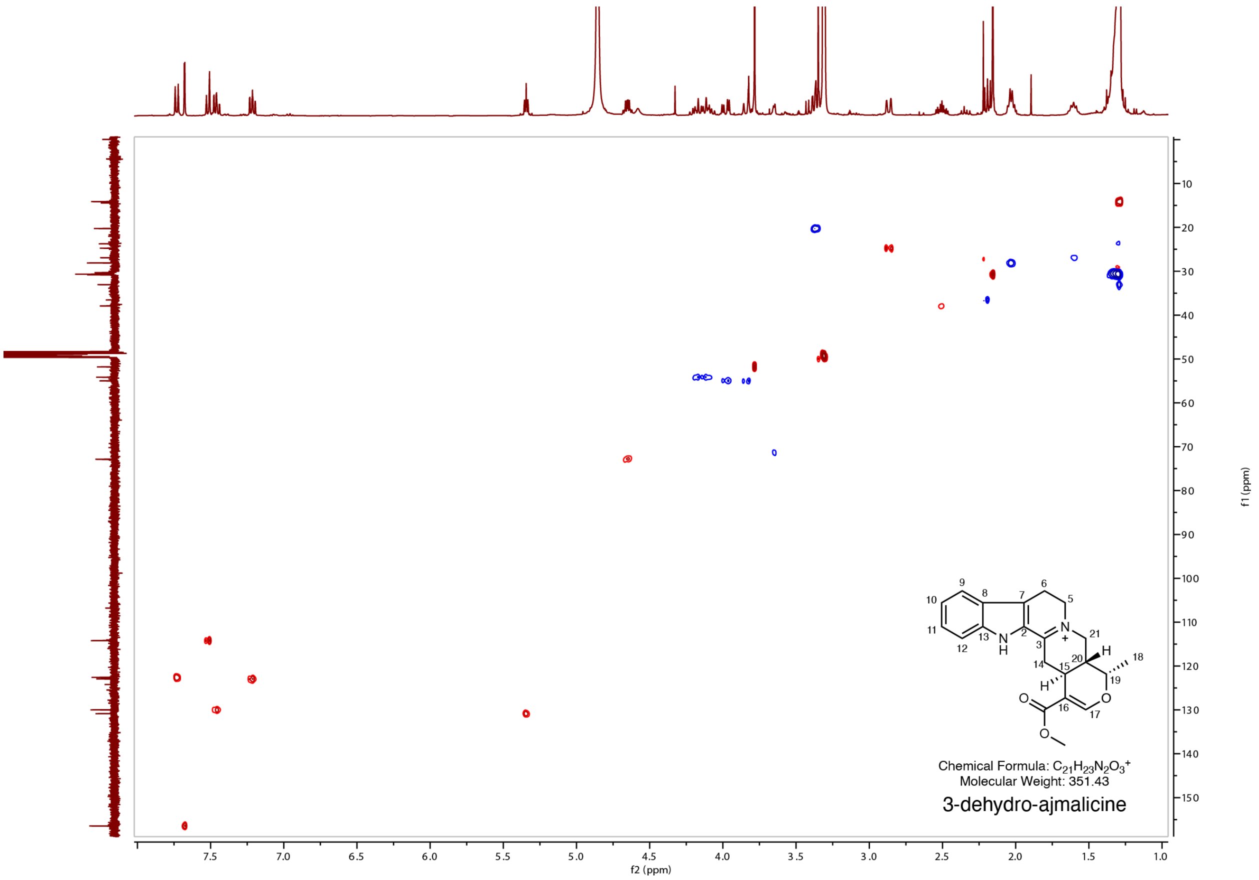

**Supplementary Figure 34. HSQC NMR spectra of 3-dehydro-ajmalicine in MeOD.**

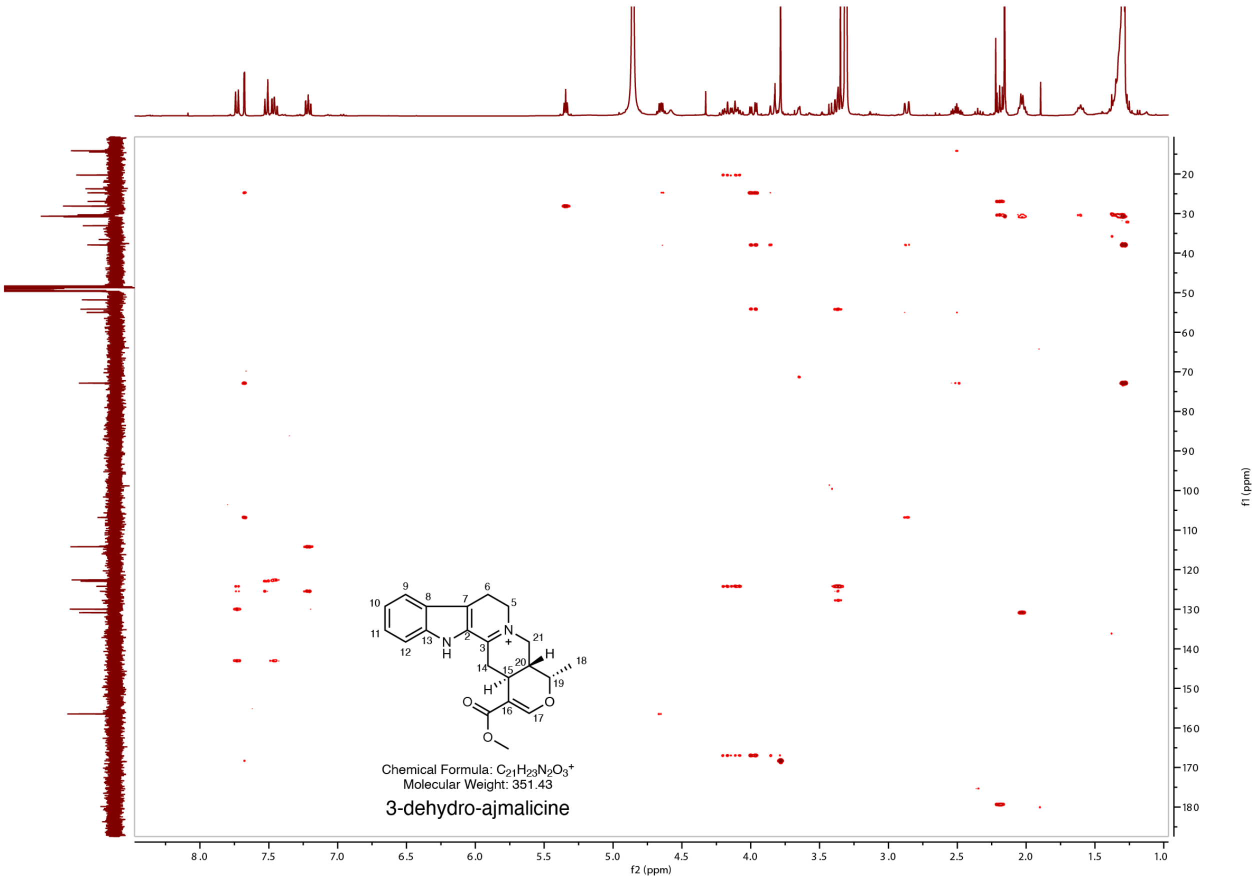

**Supplementary Figure 35. HMBC NMR spectra of 3-dehydro-ajmalicine in MeOD.**

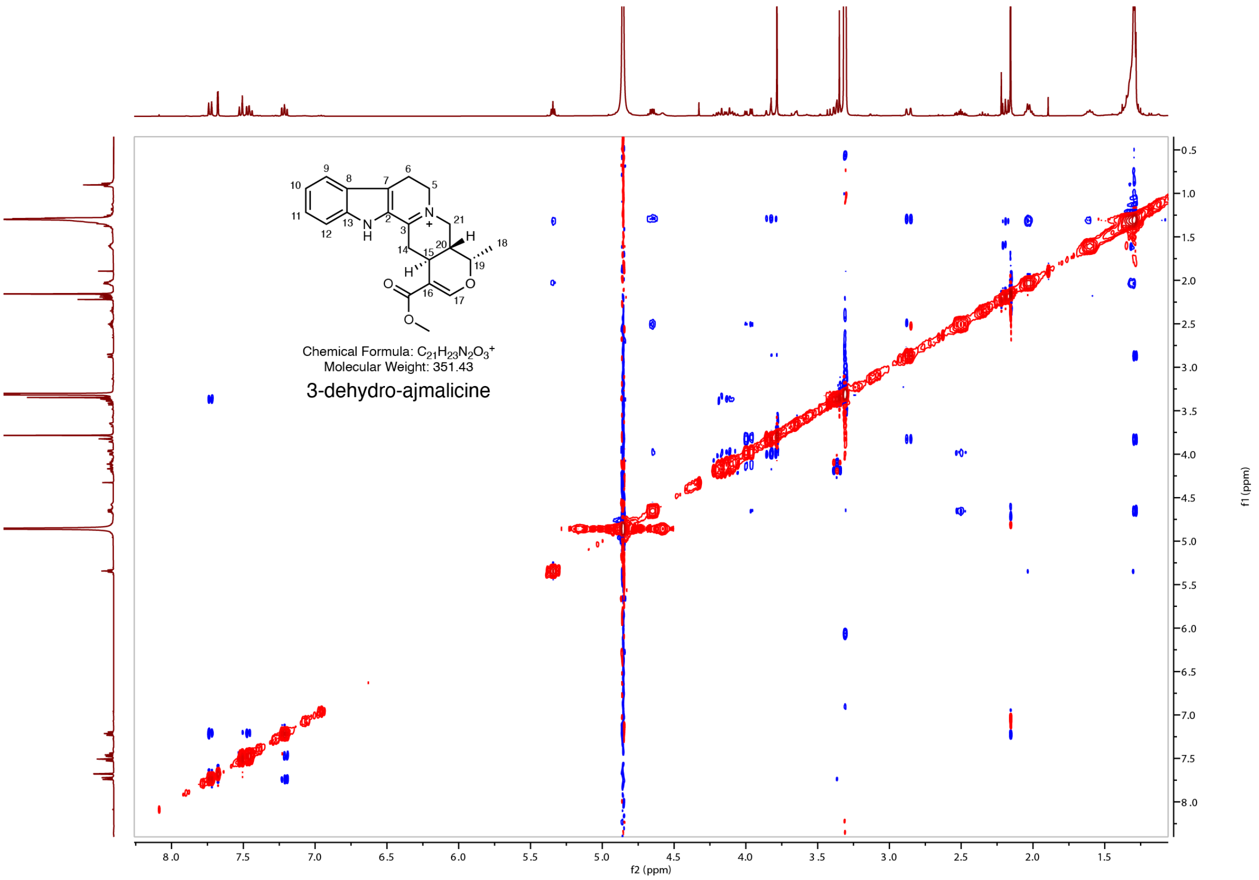

**Supplementary Figure 36. NOESY NMR spectra of 3-dehydro-ajmalicine in MeOD.**

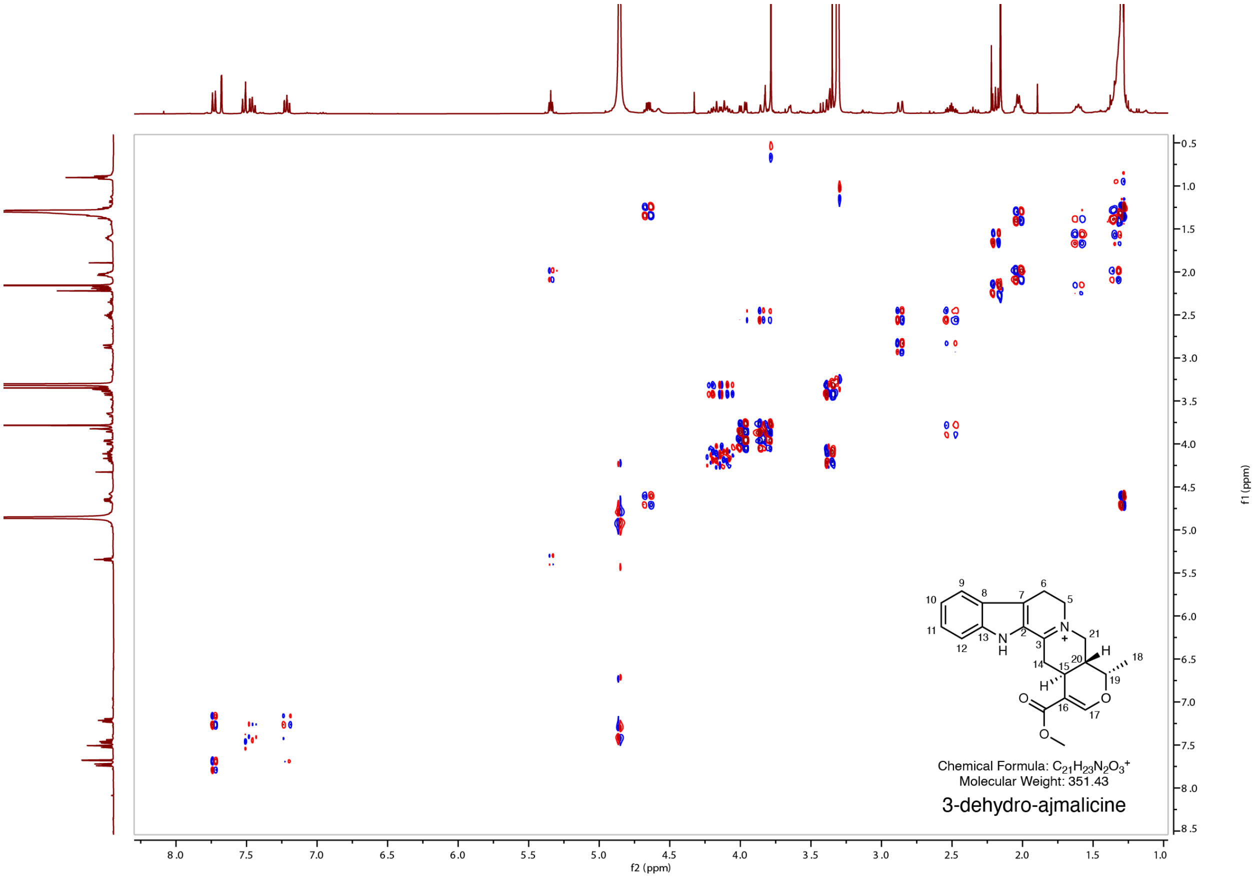

**Supplementary Figure 37. COSY NMR spectra of 3-dehydro-ajmalicine in MeOD.**

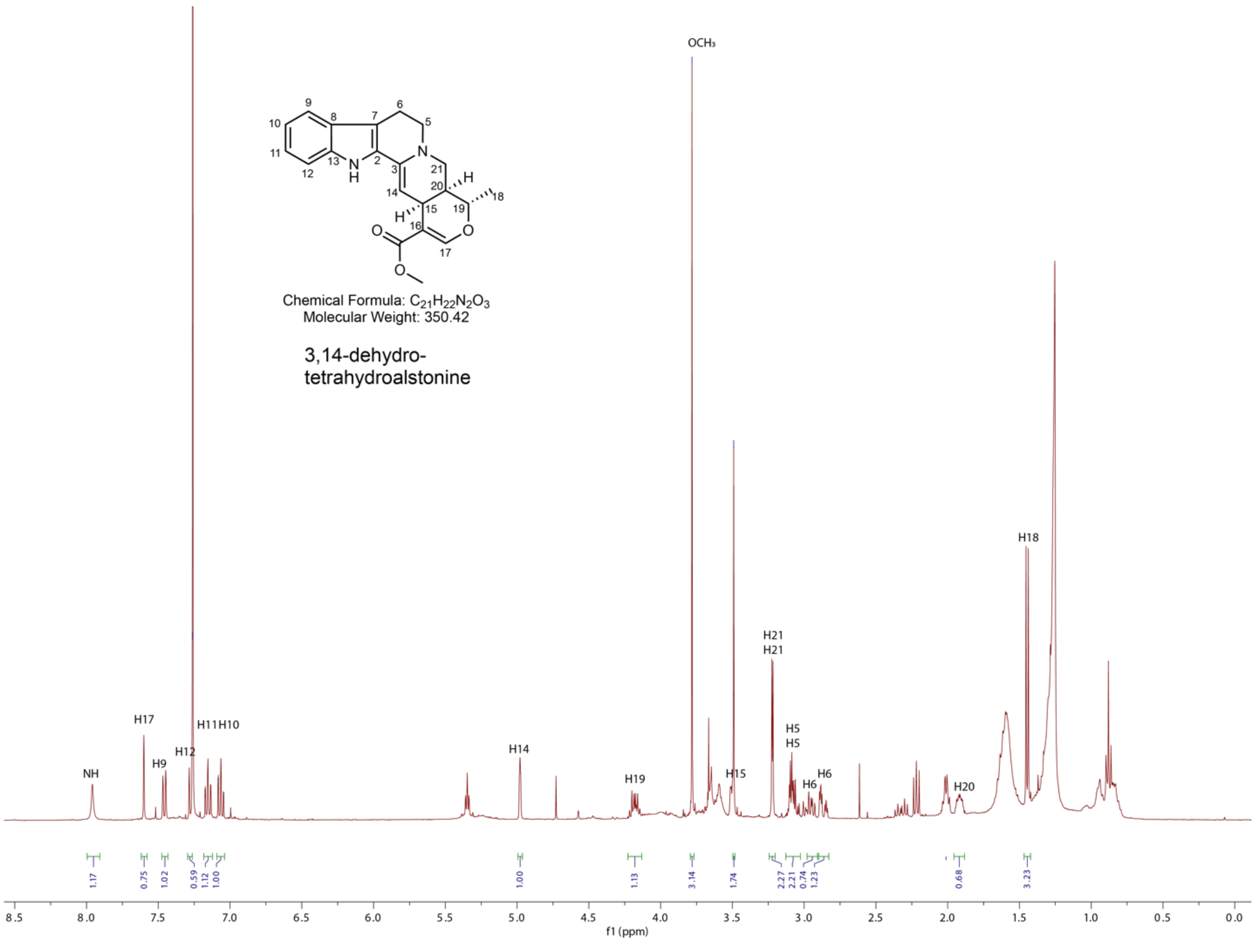

**Supplementary Figure 38.** **[1]H NMR spectra of 3,14-dehydro-tetrahydroalstonine in $CDCl_3$.**

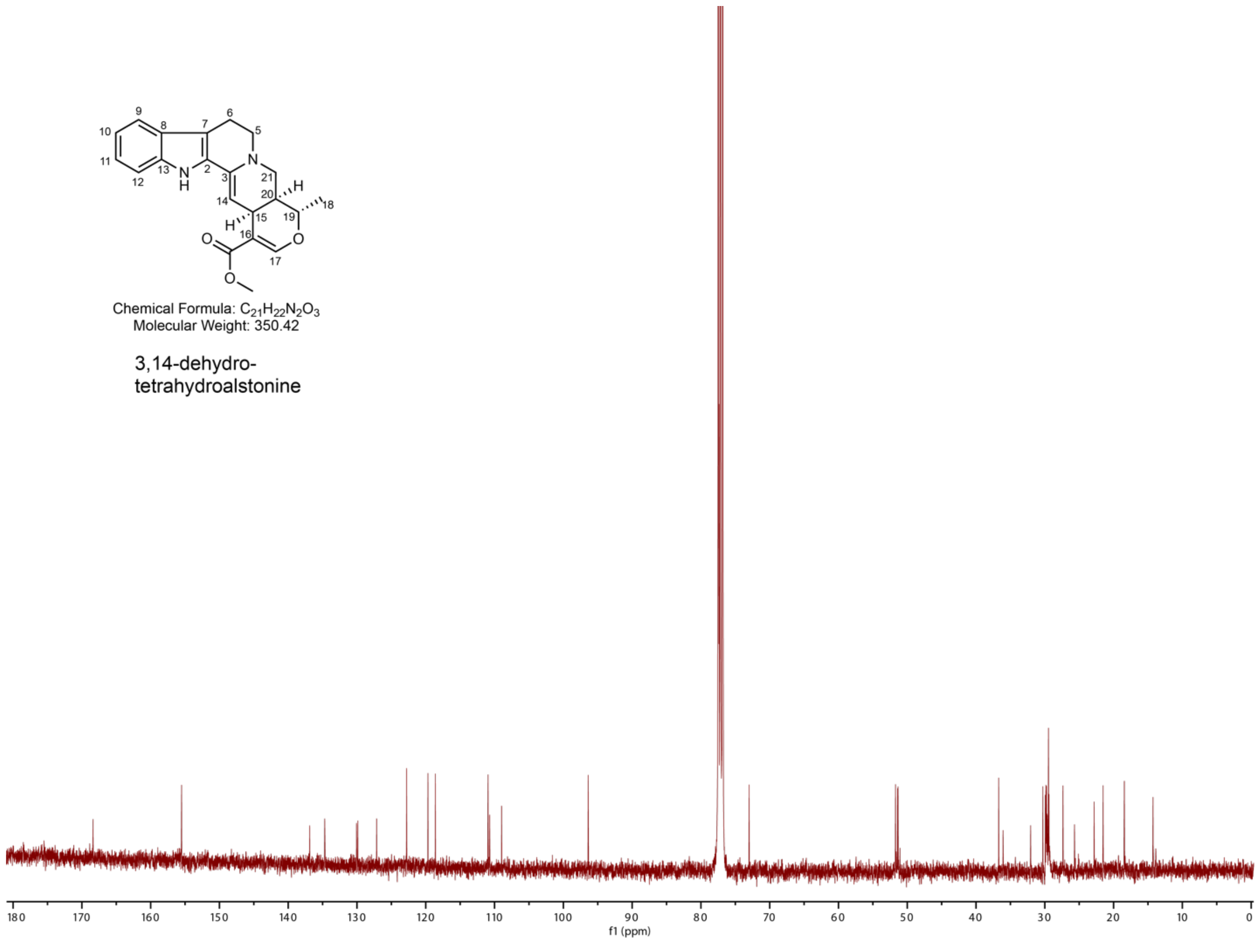

**Supplementary Figure 39. $^{13}C$ NMR spectra of 3,14-dehydro-tetrahydroalstonine in $CDCl_3$.**

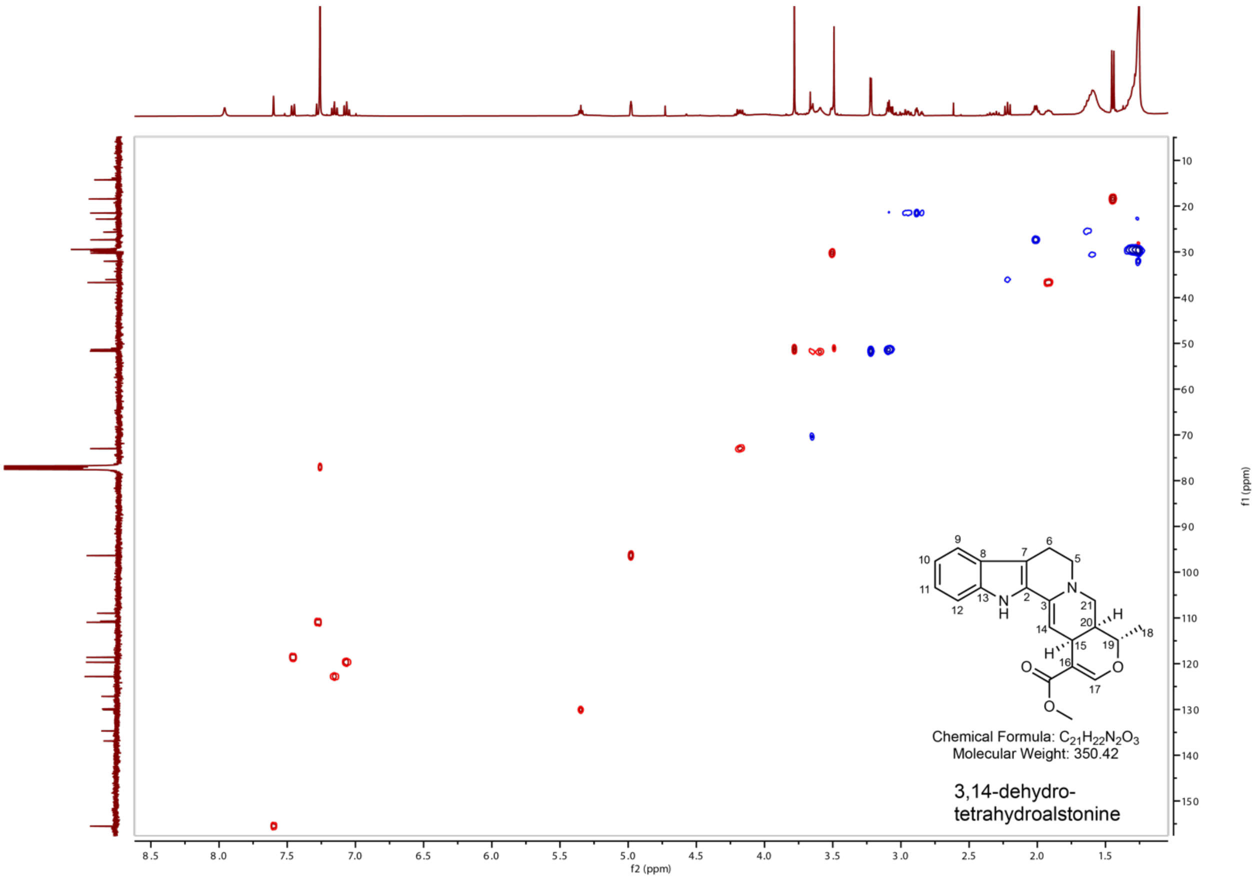

**Supplementary Figure 40. HSQC NMR spectra of 3,14-dehydro-tetrahydroalstonine in $CDCl_3$.**

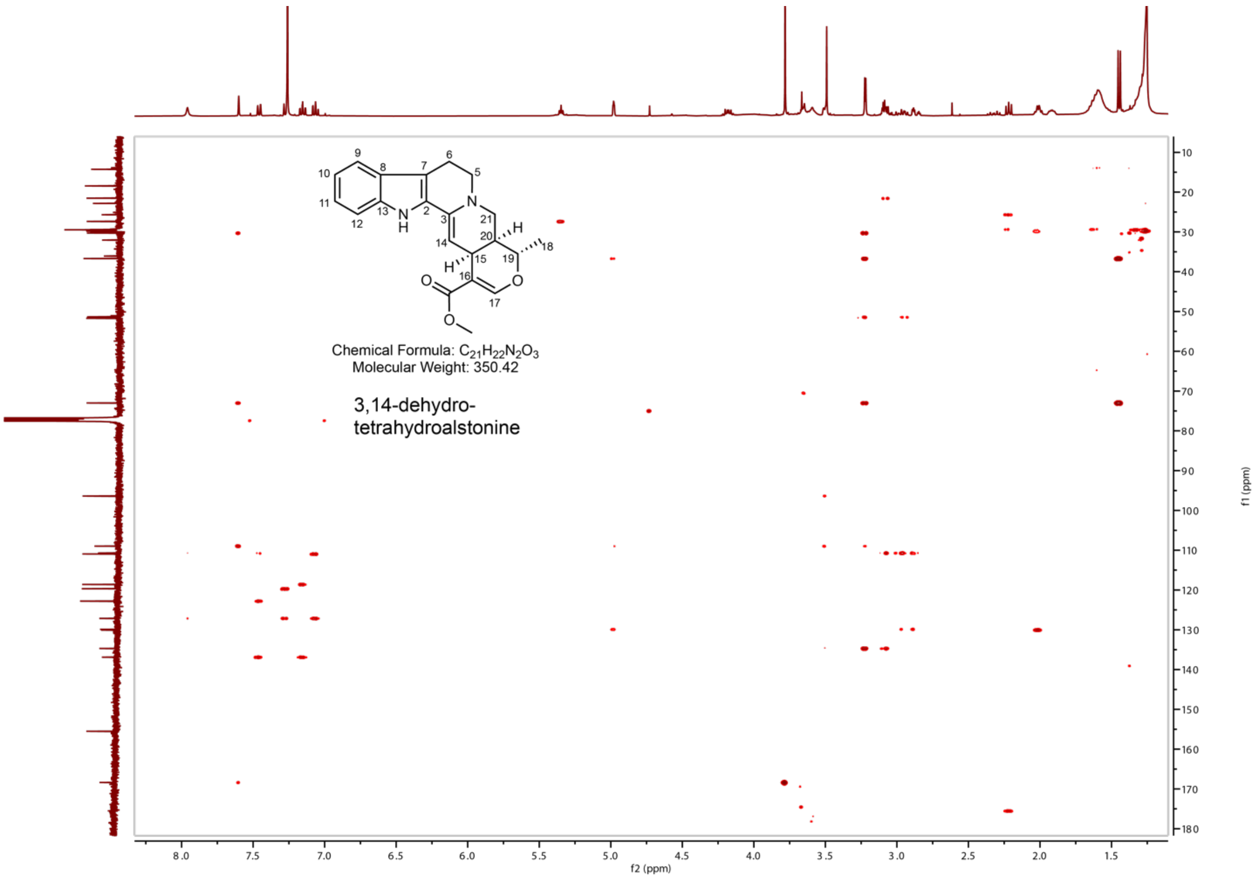

**Supplementary Figure 41. HMBC NMR spectra of 3,14-dehydro-tetrahydroalstonine in $CDCl_3$.**

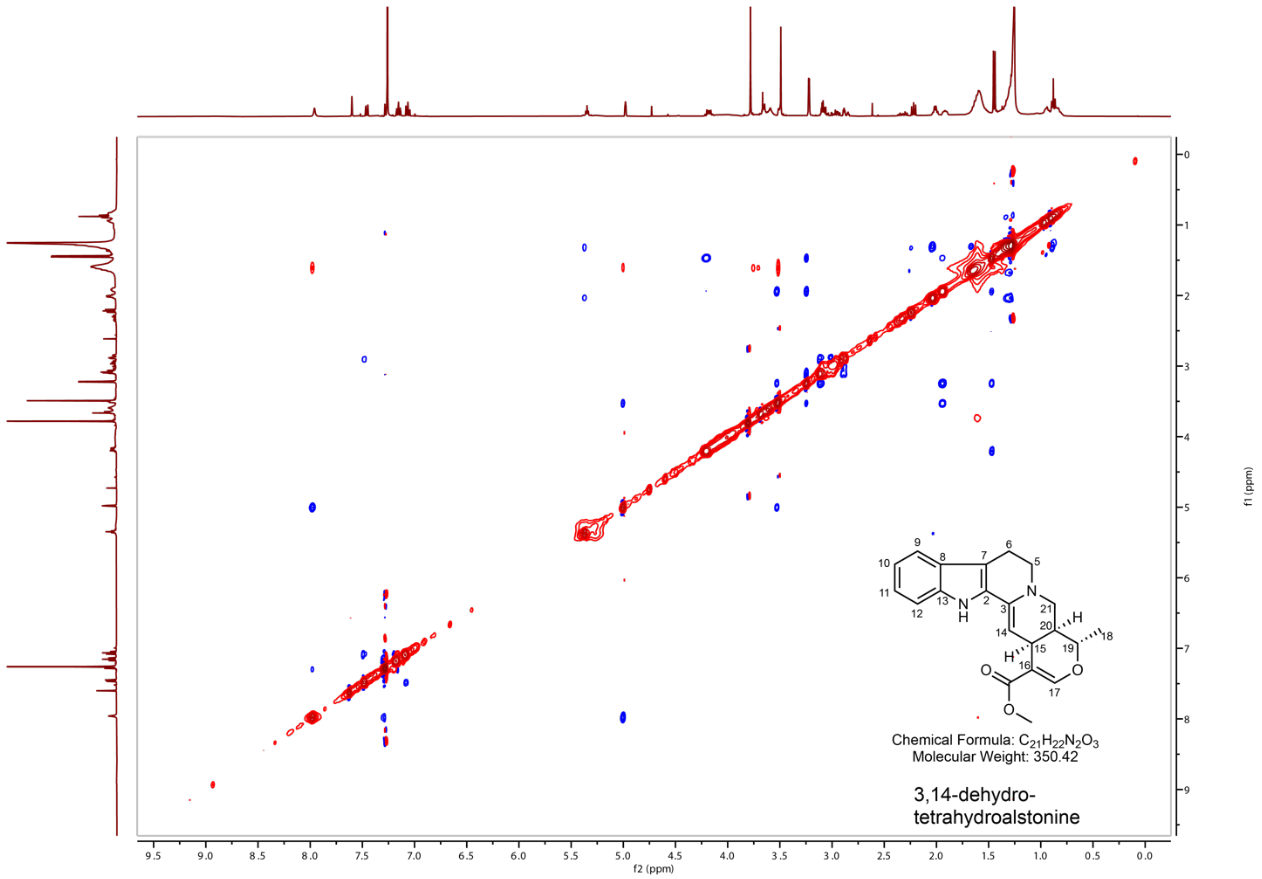

**Supplementary Figure 42. NOESY NMR spectra of 3,14-dehydro-tetrahydroalstonine in $CDCl_3$.**

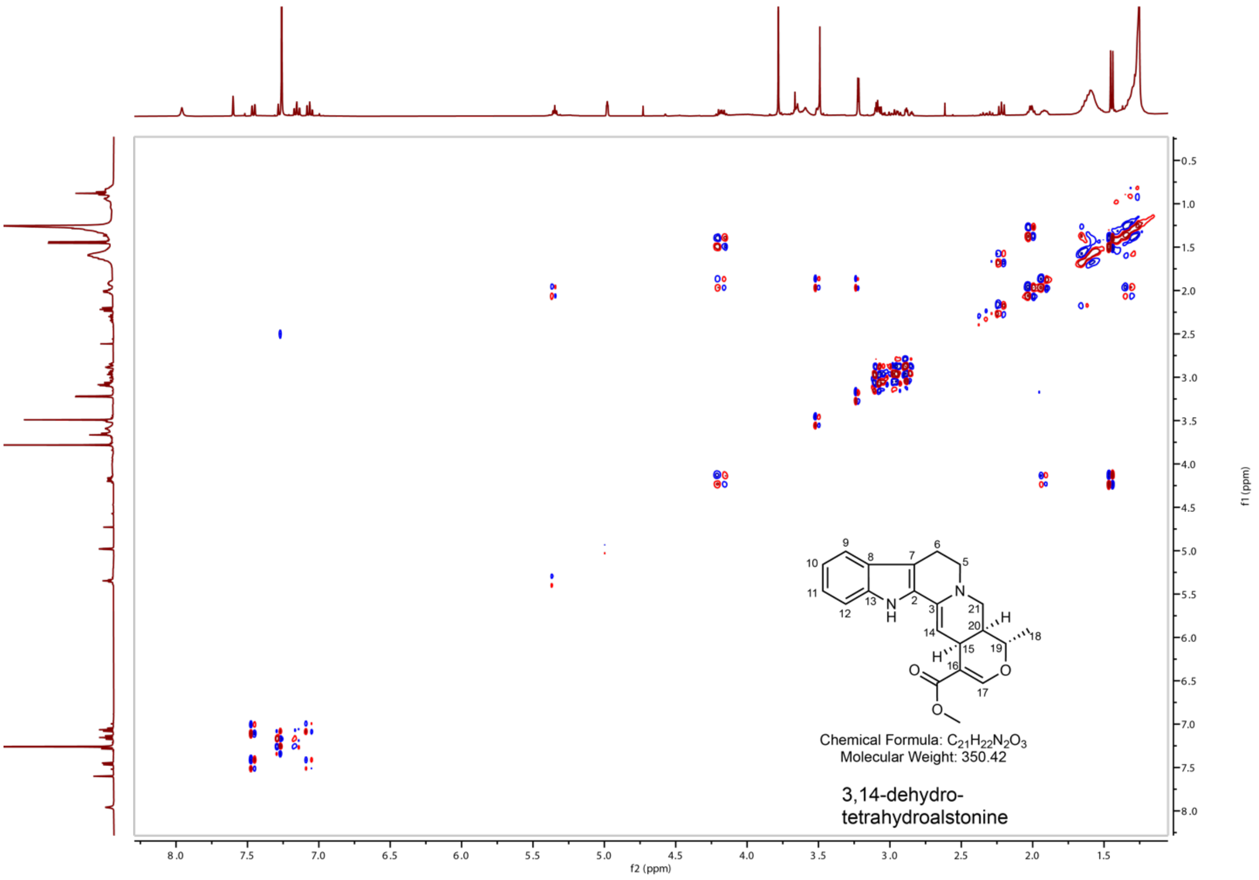

**Supplementary Figure 43. COSY NMR spectra of 3,14-dehydro-tetrahydroalstonine in $CDCl_3$.**

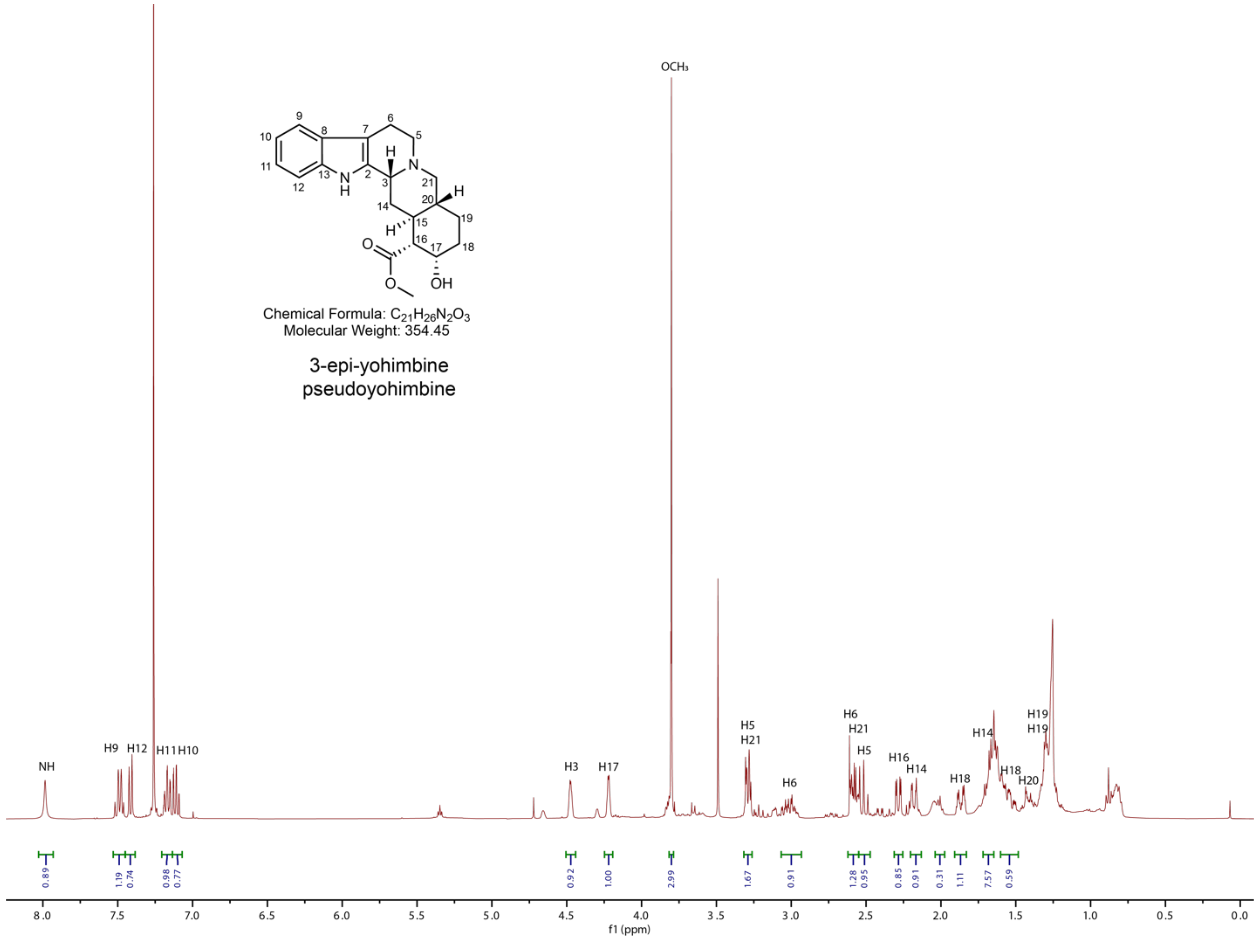

**Supplementary Figure 44. ¹H NMR spectra of 3-epi-yohimbine (pseudoyohimbine) in CDCl3.**

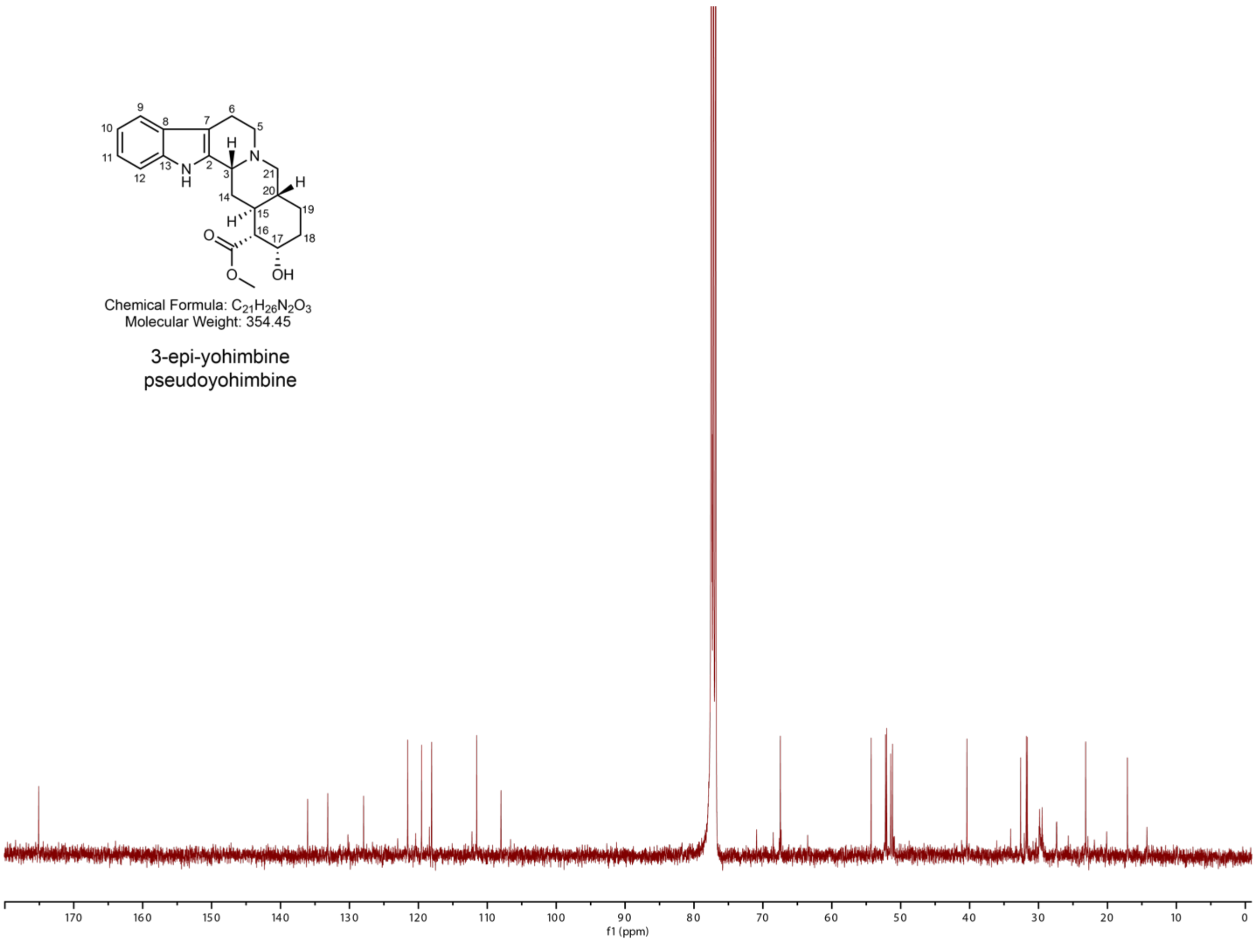

**Supplementary Figure 45. $^{13}$C NMR spectra of 3-epi-yohimbine (pseudoyohimbine) in $CDCl_3$.**

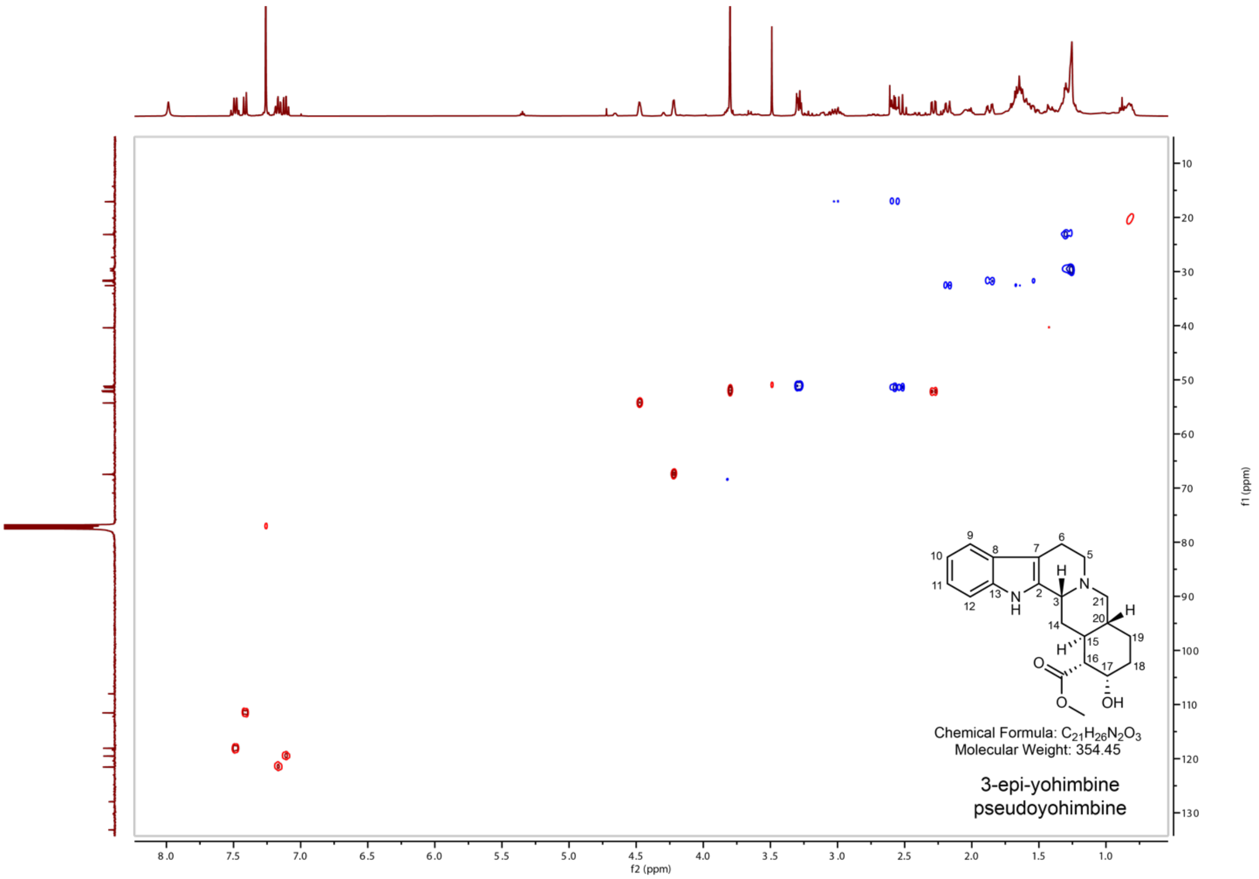

**Supplementary Figure 46. HSQC NMR spectra of 3-epi-yohimbine (pseudoyohimbine) in $CDCl_3$.**

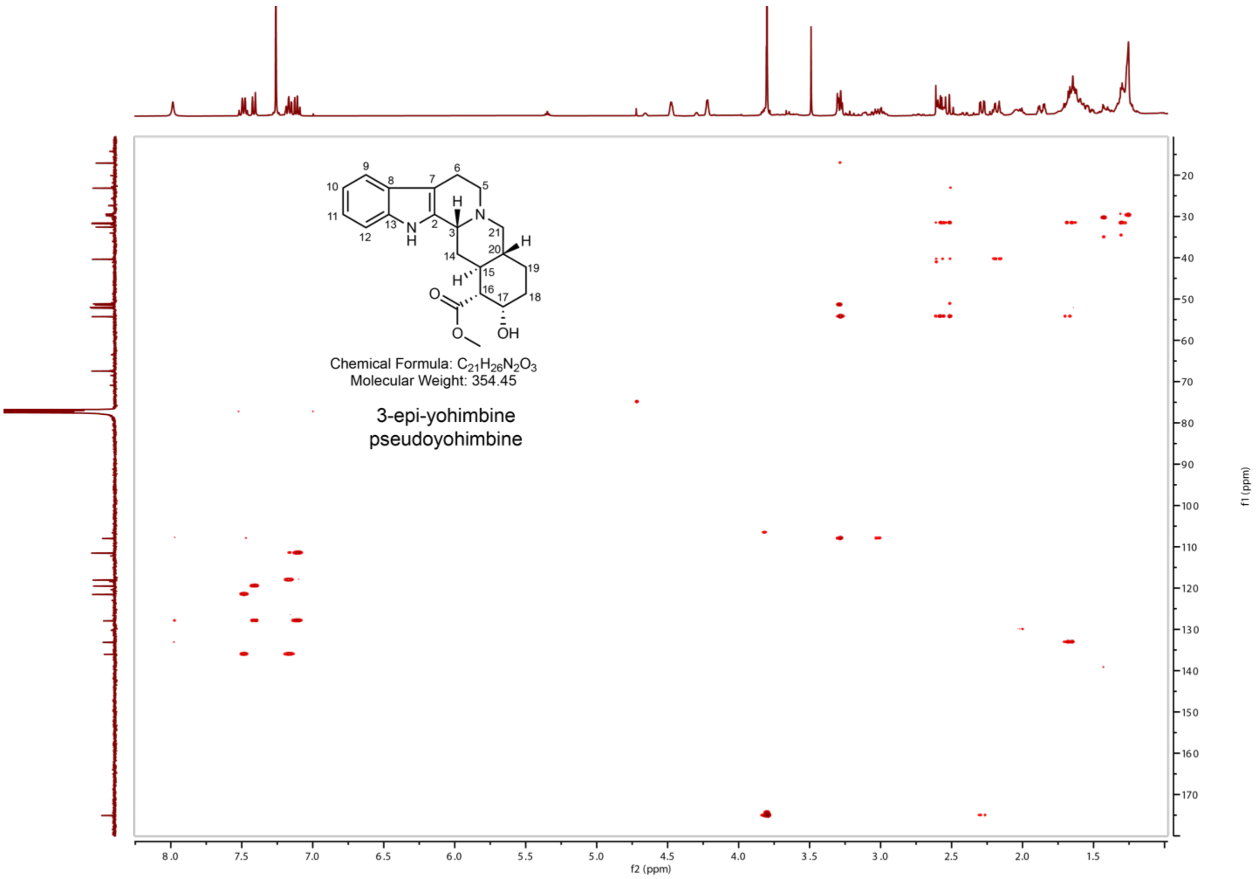

**Supplementary Figure 47. HMBC NMR spectra of 3-epi-yohimbine (pseudoyohimbine) in CDCl3.**

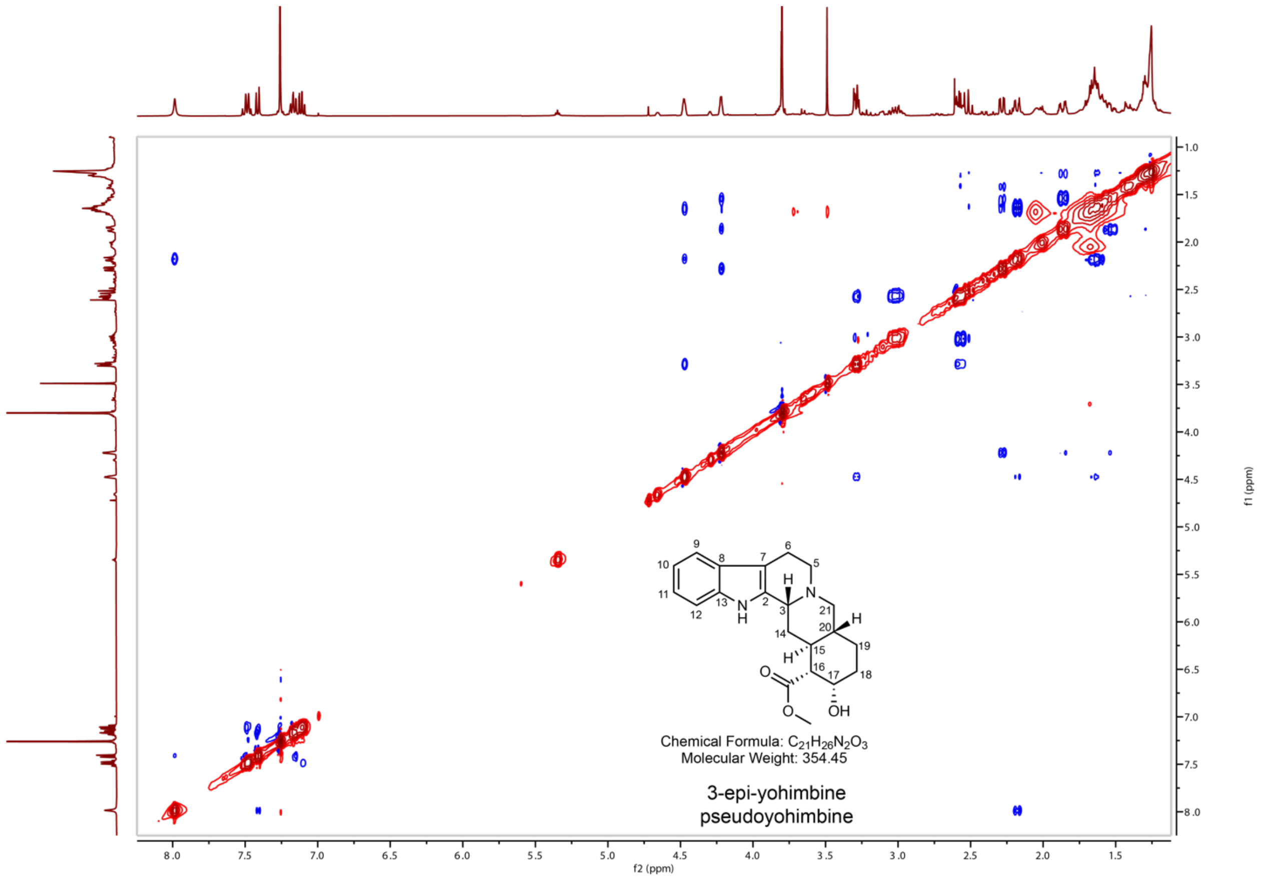

**Supplementary Figure 48. NOESY NMR spectra of 3-epi-yohimbine (pseudoyohimbine) in $CDCl_3$.**

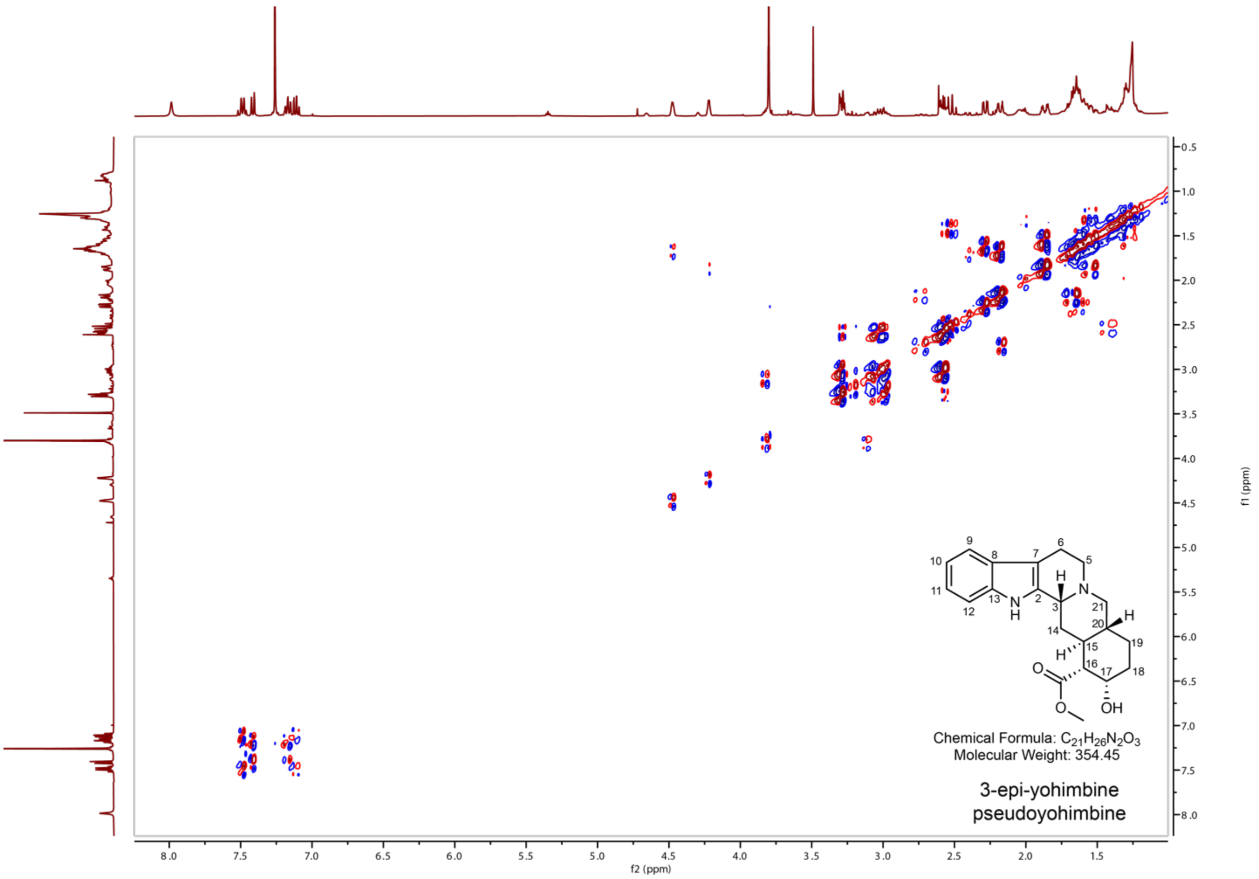

**Supplementary Figure 49. COSY NMR spectra of 3-epi-yohimbine (pseudoyohimbine) in $CDCl_3$.**

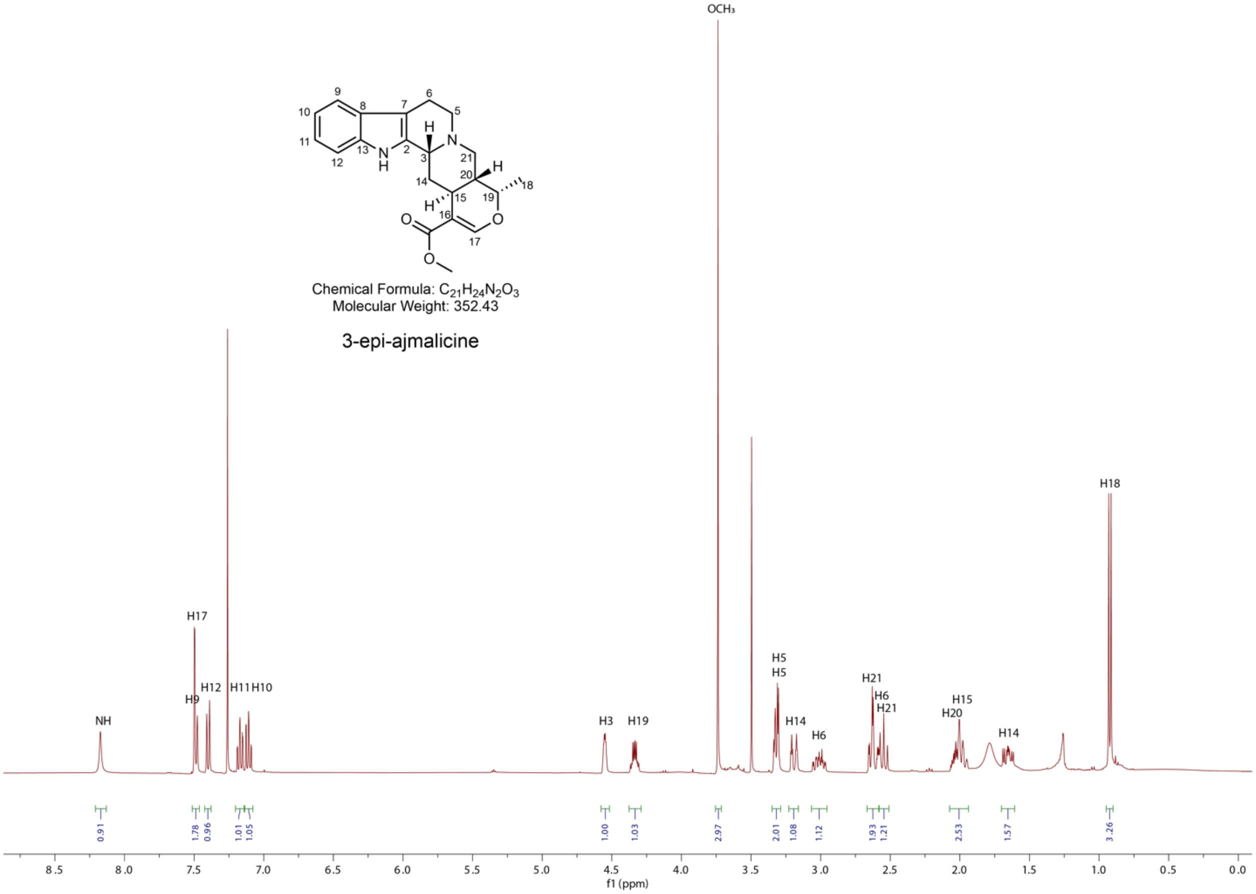

**Supplementary Figure 50. $^{1}H$ NMR spectra of 3-epi-ajmalicine in $CDCl_3$.**

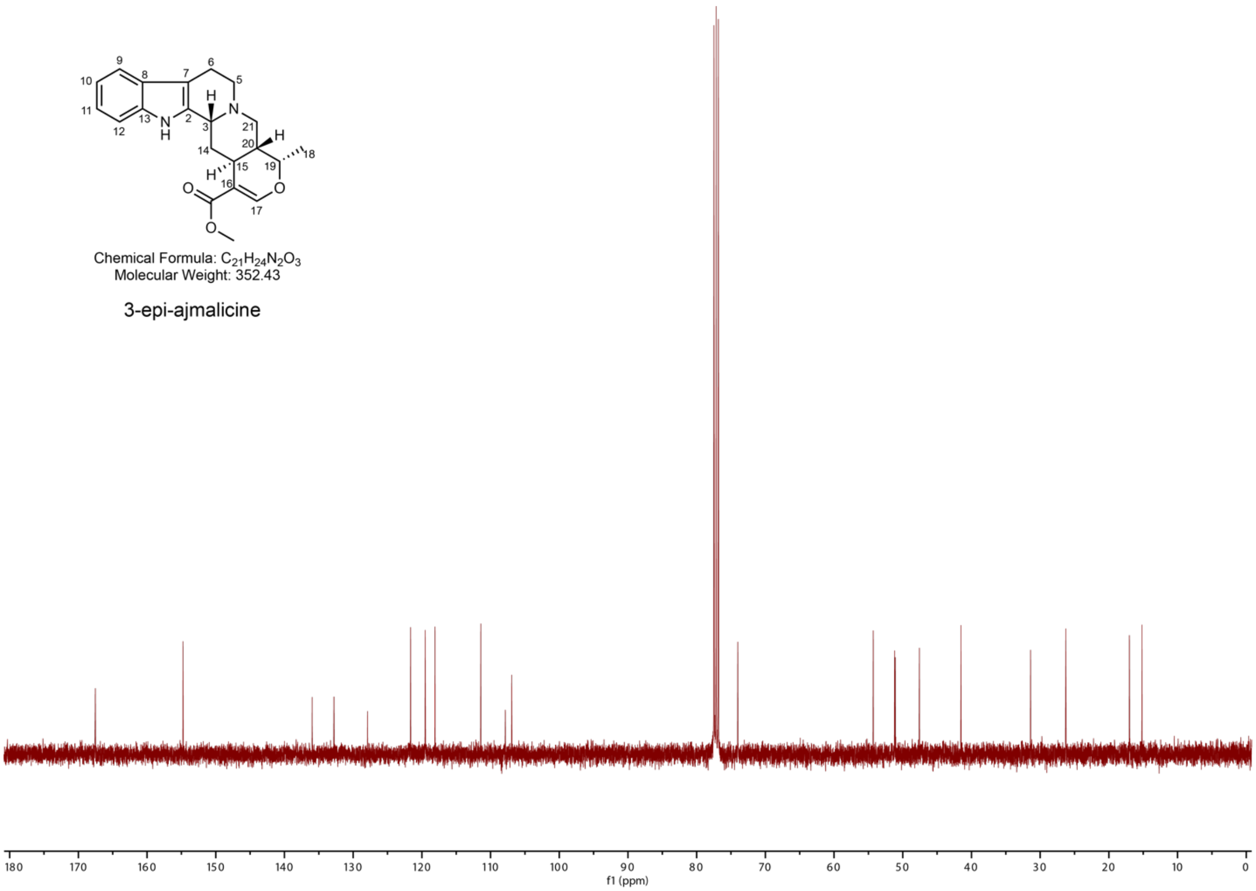

**Supplementary Figure 51. $^{13}C$ NMR spectra of 3-epi-ajmalicine in $CDCl_3$.**

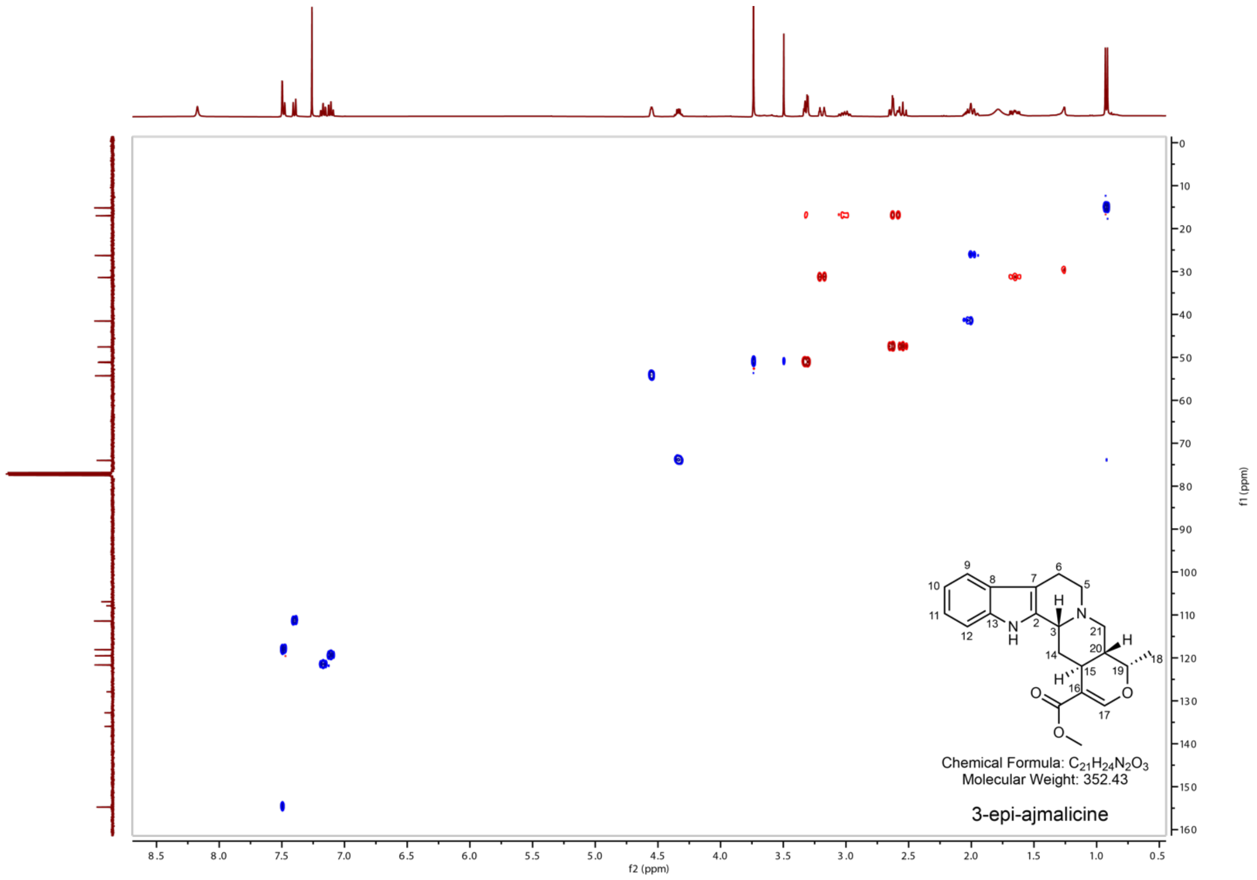

**Supplementary Figure 52. HSQC NMR spectra of 3-epi-ajmalicine in $CDCl_3$.**

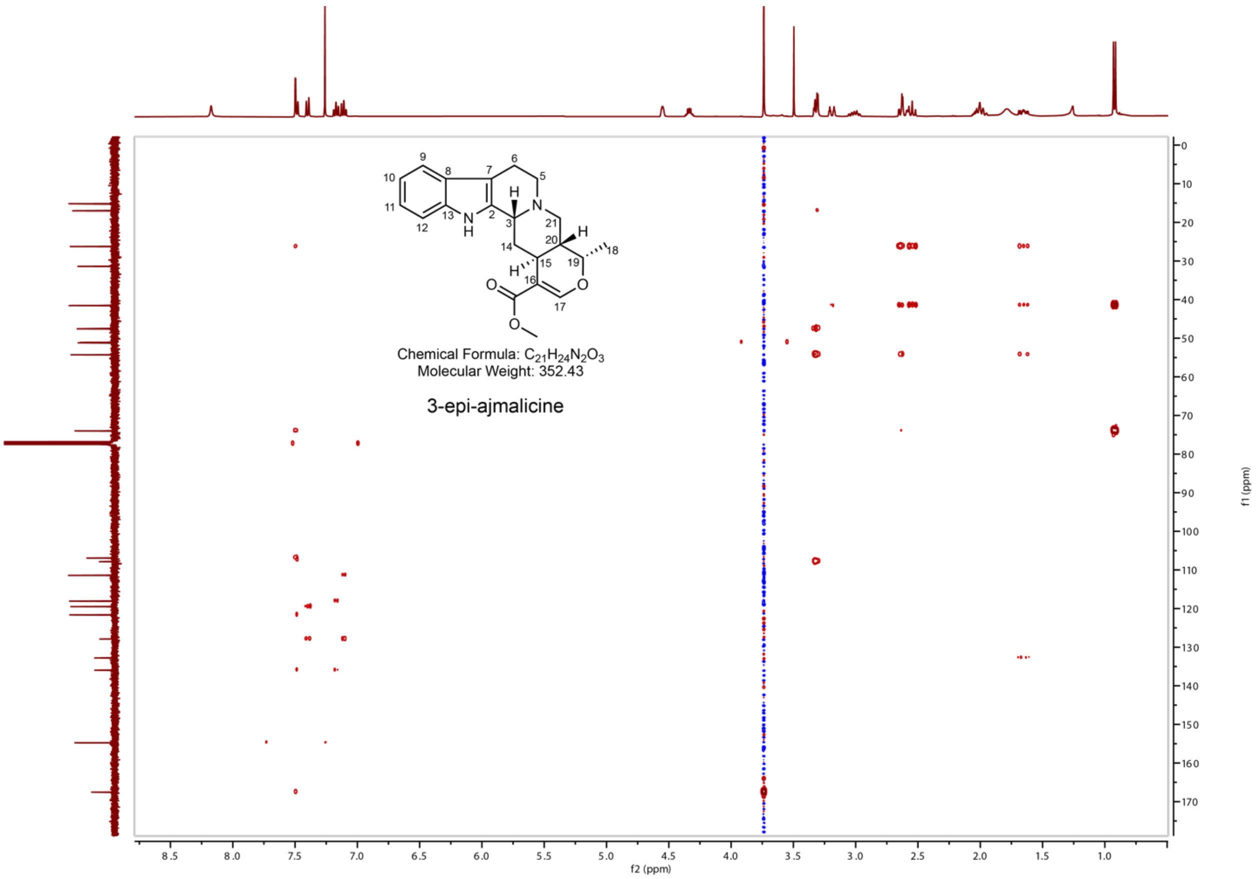

**Supplementary Figure 53. HMBC NMR spectra of 3-epi-ajmalicine in $CDCl_3$.**

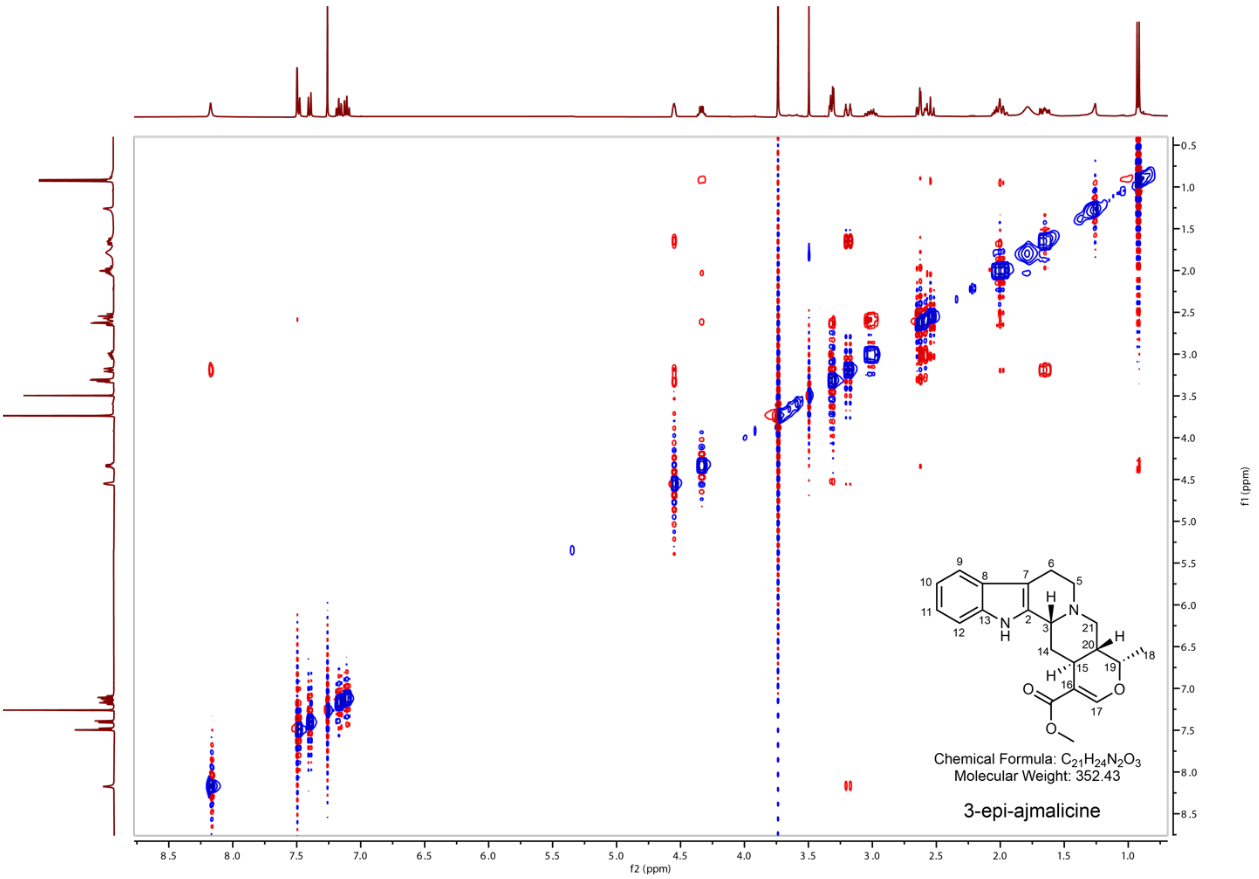

**Supplementary Figure 54. NOESY NMR spectra of 3-epi-ajmalicine in $CDCl_3$.**

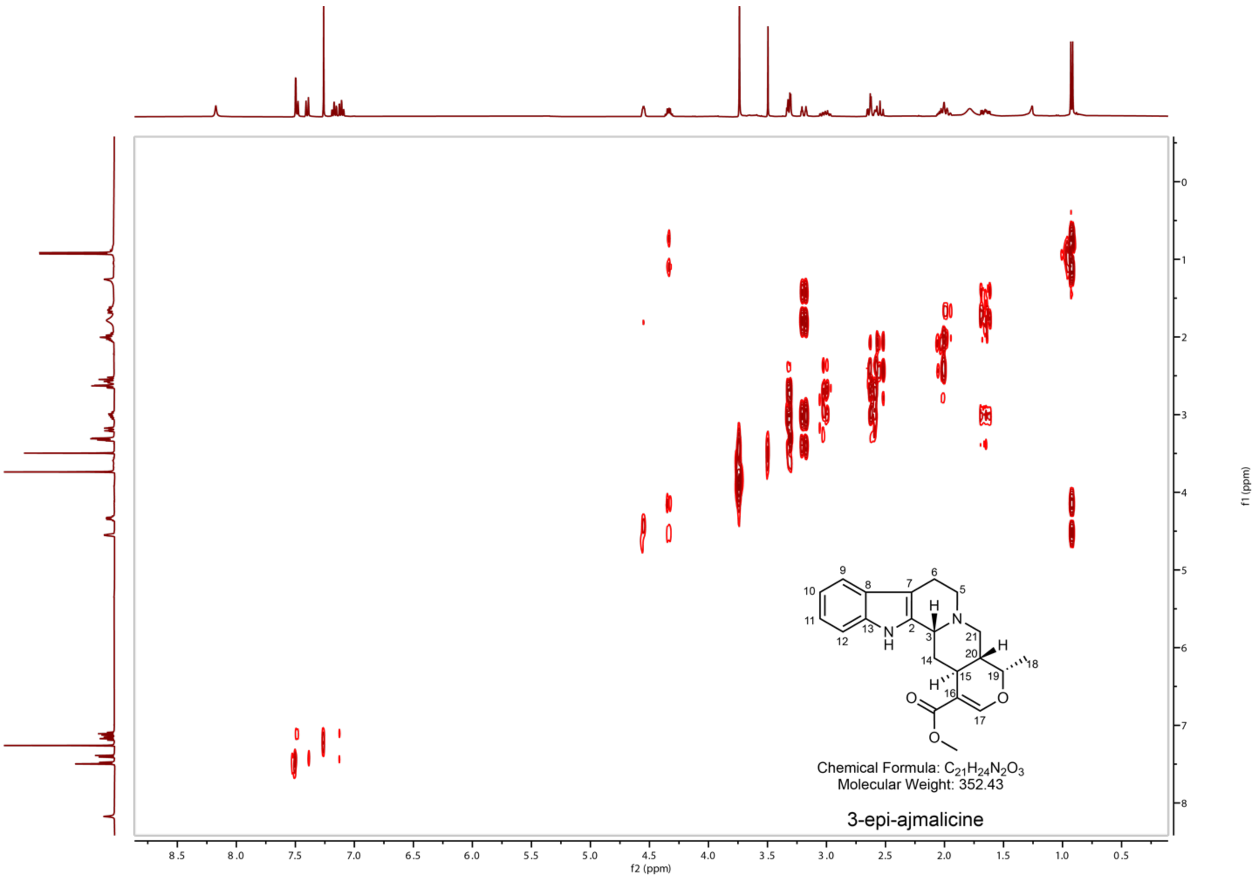

**Supplementary Figure 55. COSY NMR spectra of 3-epi-ajmalicine in $CDCl_3$.**

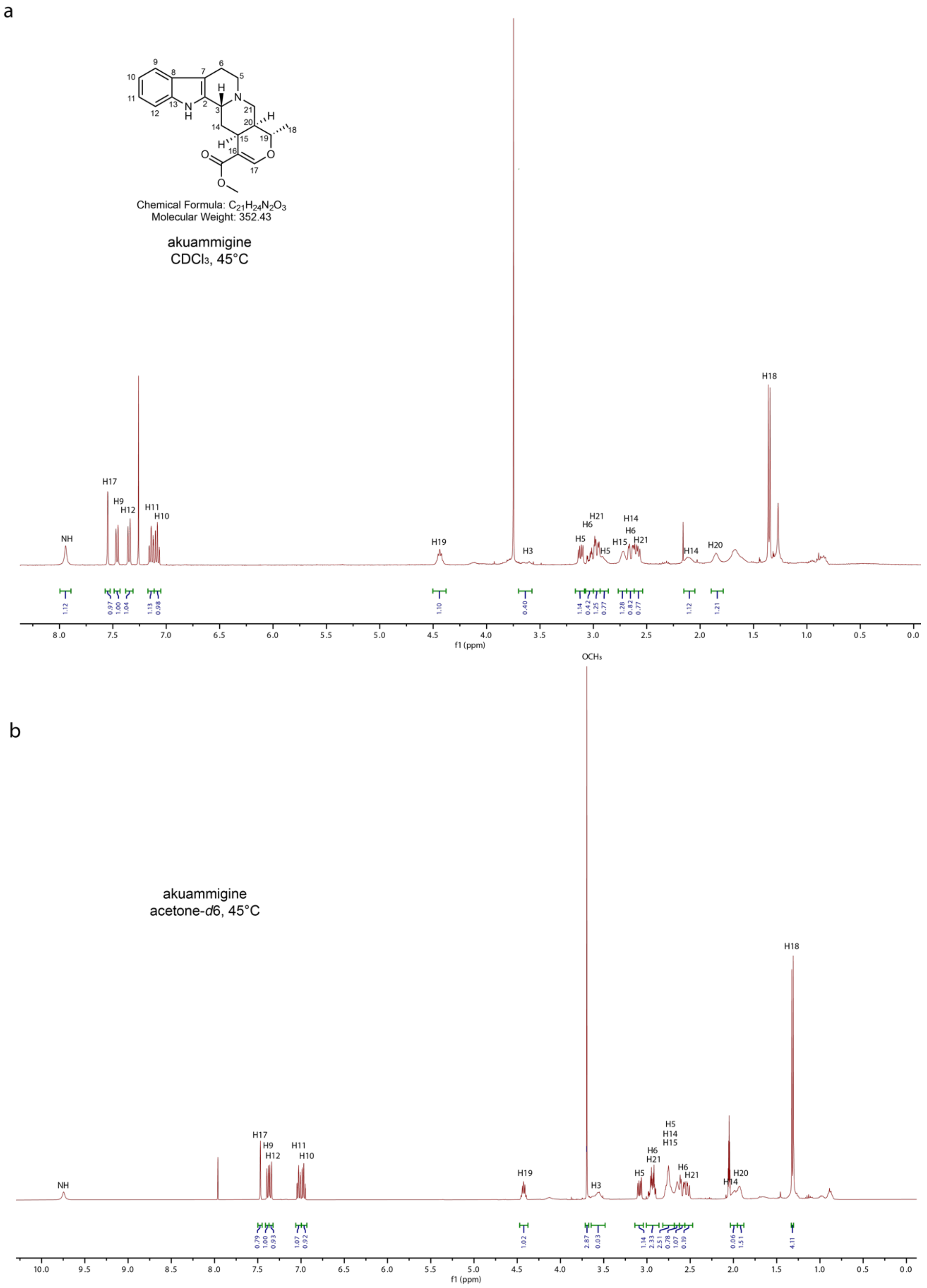

**Supplementary Figure 56.** **$^{1}$H NMR spectra of 3-epi-tetrahydroalstonine (akuammigine) in $CDCl_3$ and acetone-*d6* (b) at 45 °C.**

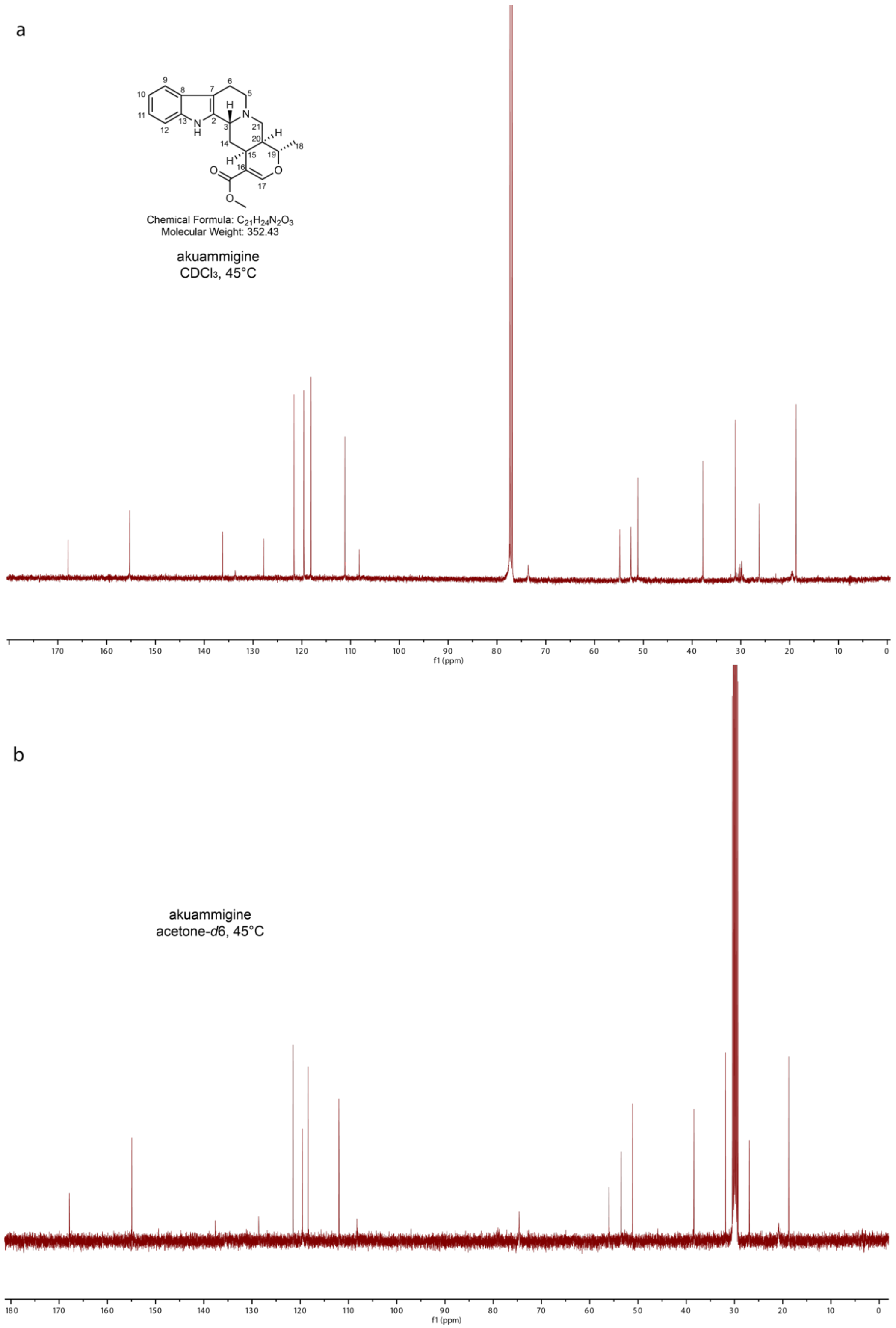

**Supplementary Figure 57. $^{13}$C NMR spectra of 3-epi-tetrahydroalstonine (akuammigine) in $CDCl_3$ and acetone-*d6* (b) at 45 °C.**

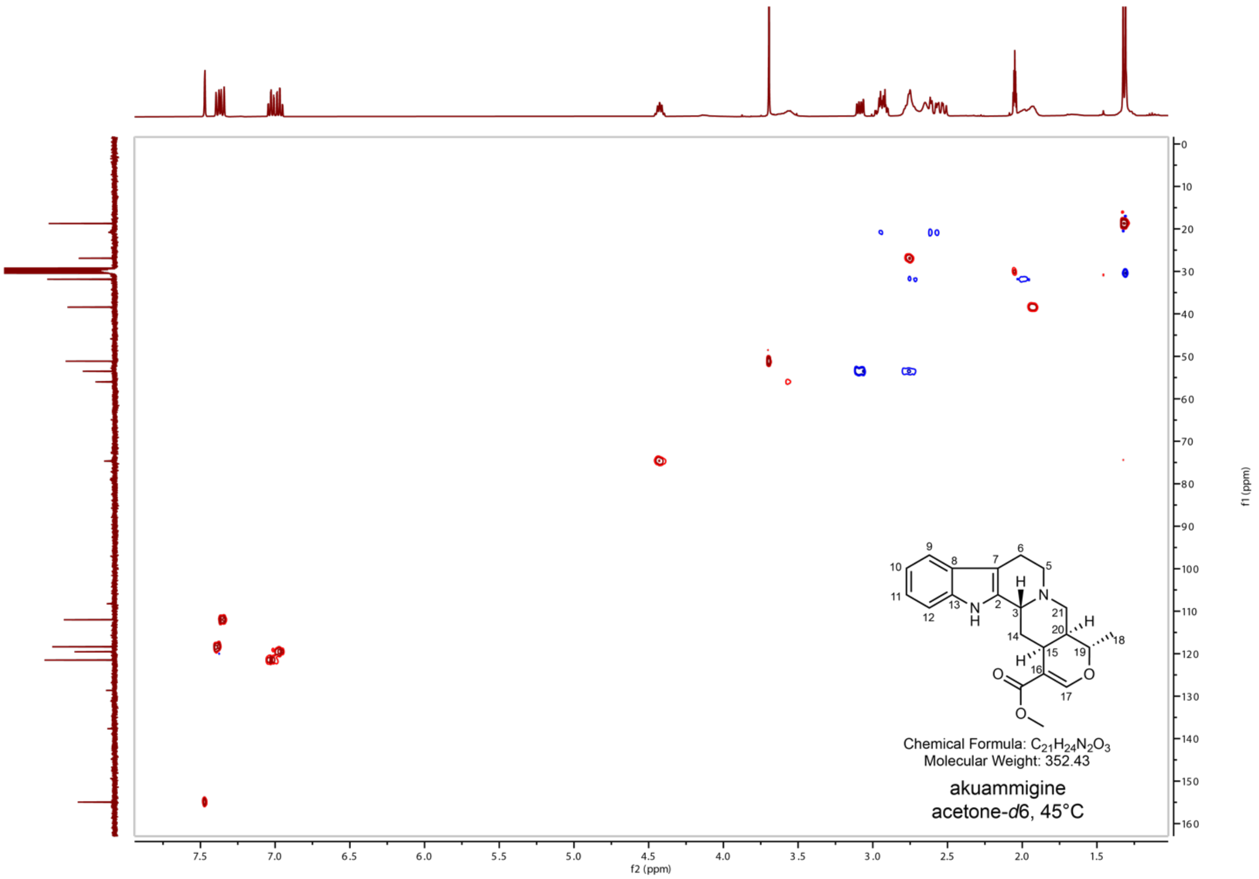

**Supplementary Figure 58. HSQC NMR spectra of 3-epi-tetrahydroalstonine (akuammigine) in acetone-*d6* 45 °C.**

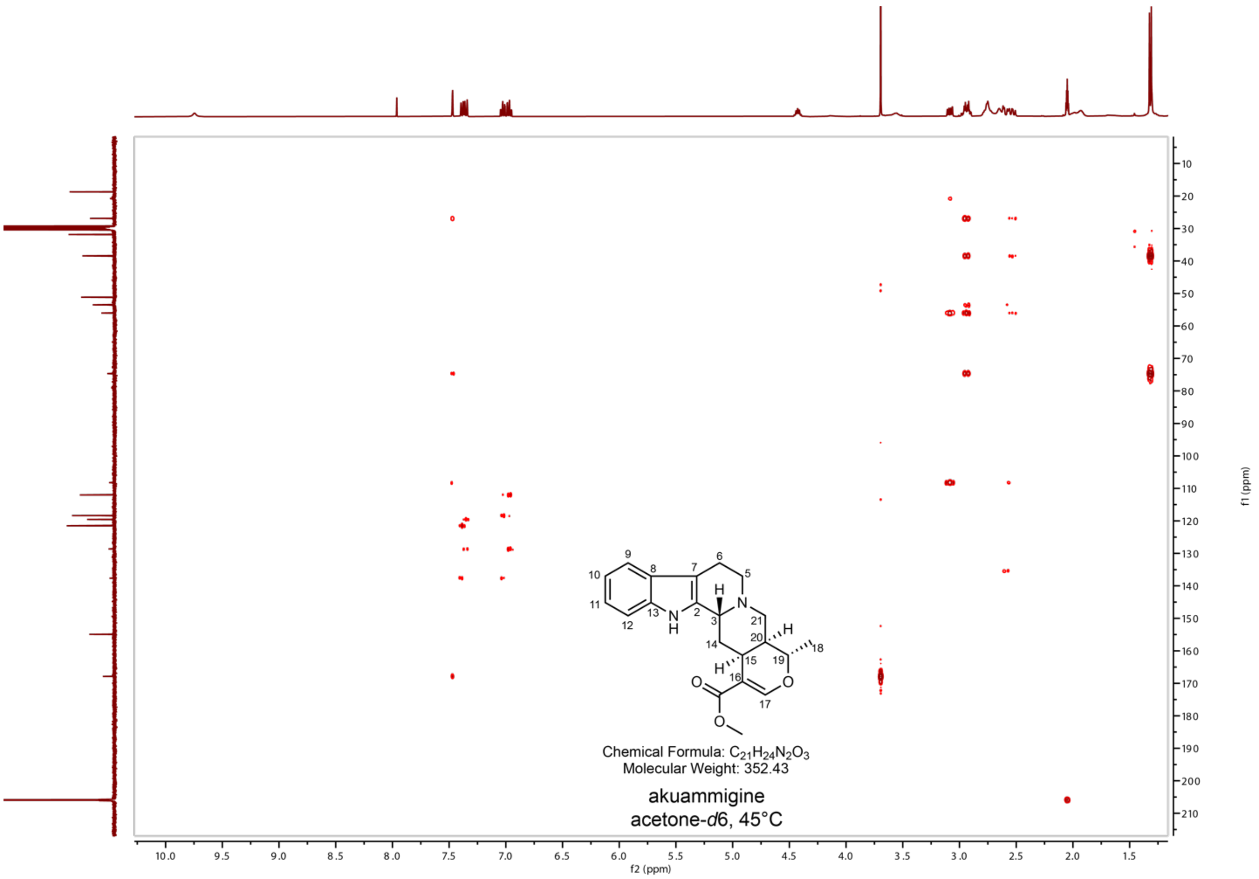

**Supplementary Figure 59. HMBC NMR spectra of 3-epi-tetrahydroalstonine (akuammigine) in acetone-*d6* 45 °C.**

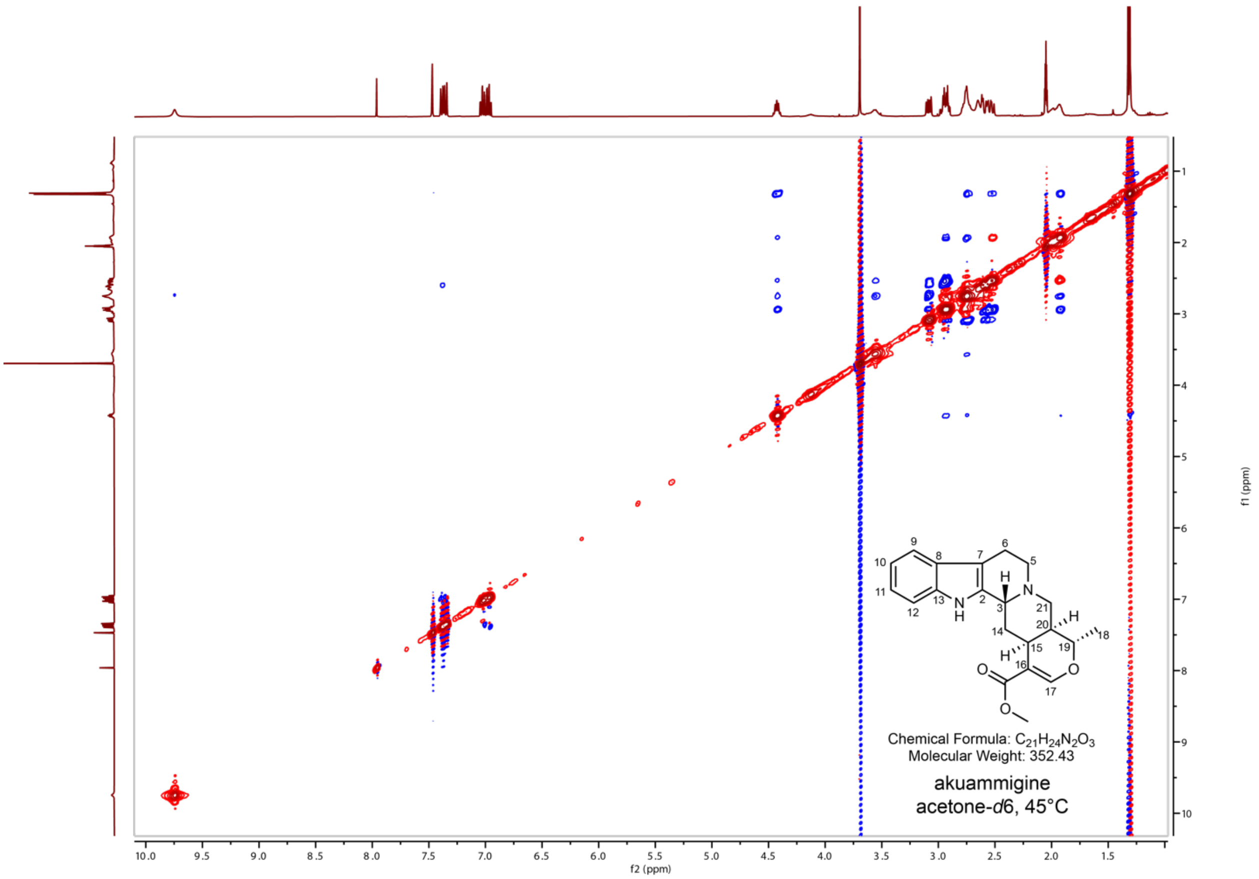

**Supplementary Figure 60. NOESY NMR spectra of 3-epi-tetrahydroalstonine (akuammigine) in acetone-*d6* 45 °C.**

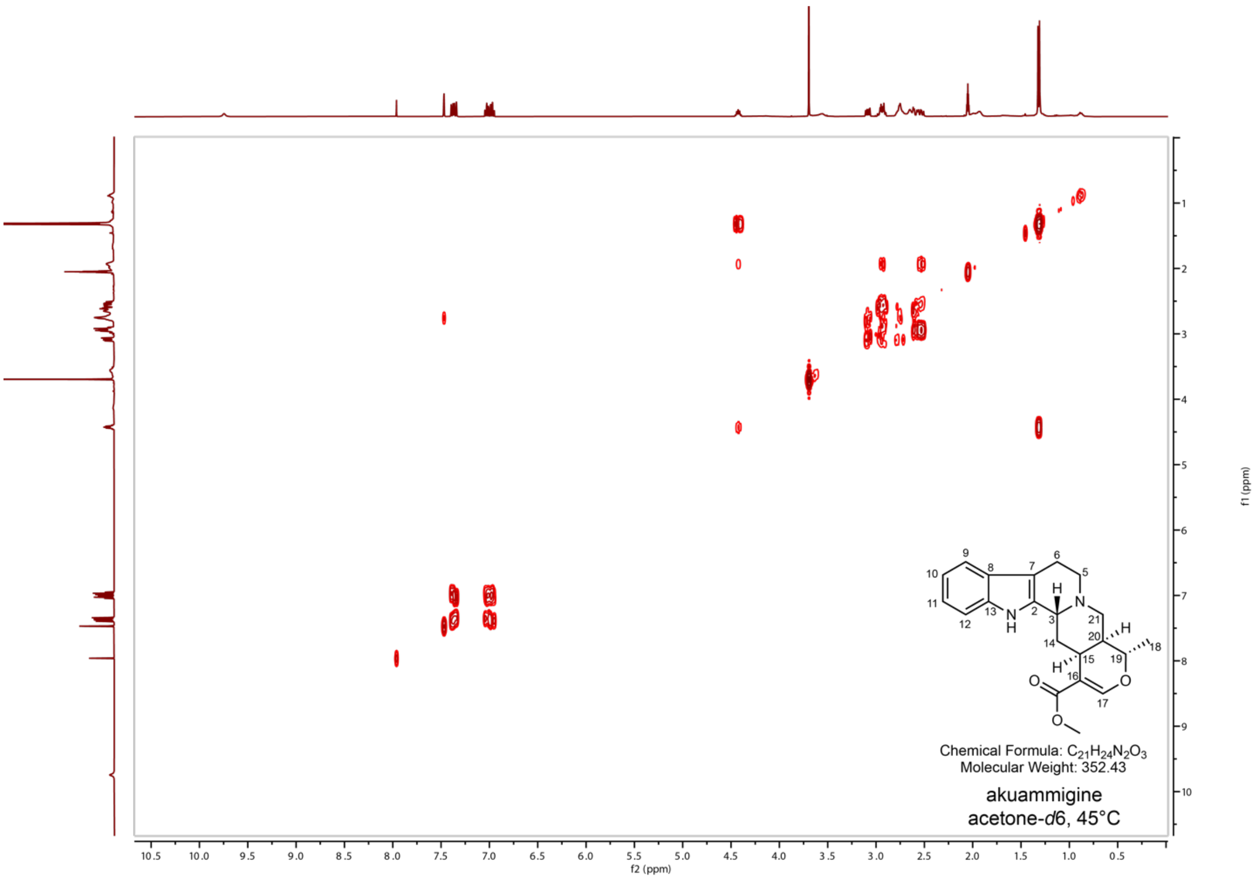

**Supplementary Figure 61. COSY NMR spectra of 3-epi-tetrahydroalstonine (akuammigine) in acetone-*d6* 45 °C.**

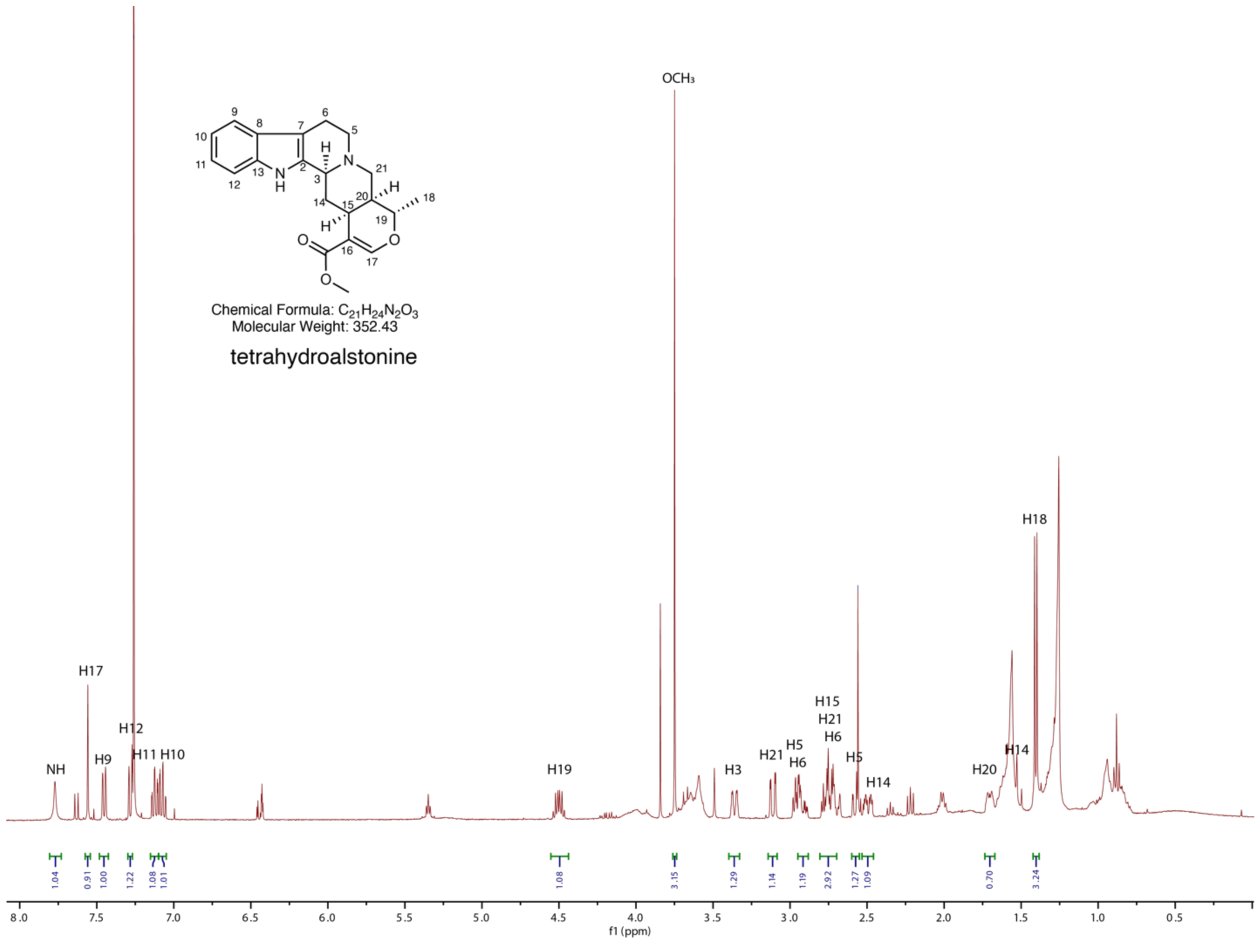

**Supplementary Figure 62. $^{1}H$ NMR spectra of tetrahydroalstonine in $CDCl_3$.**

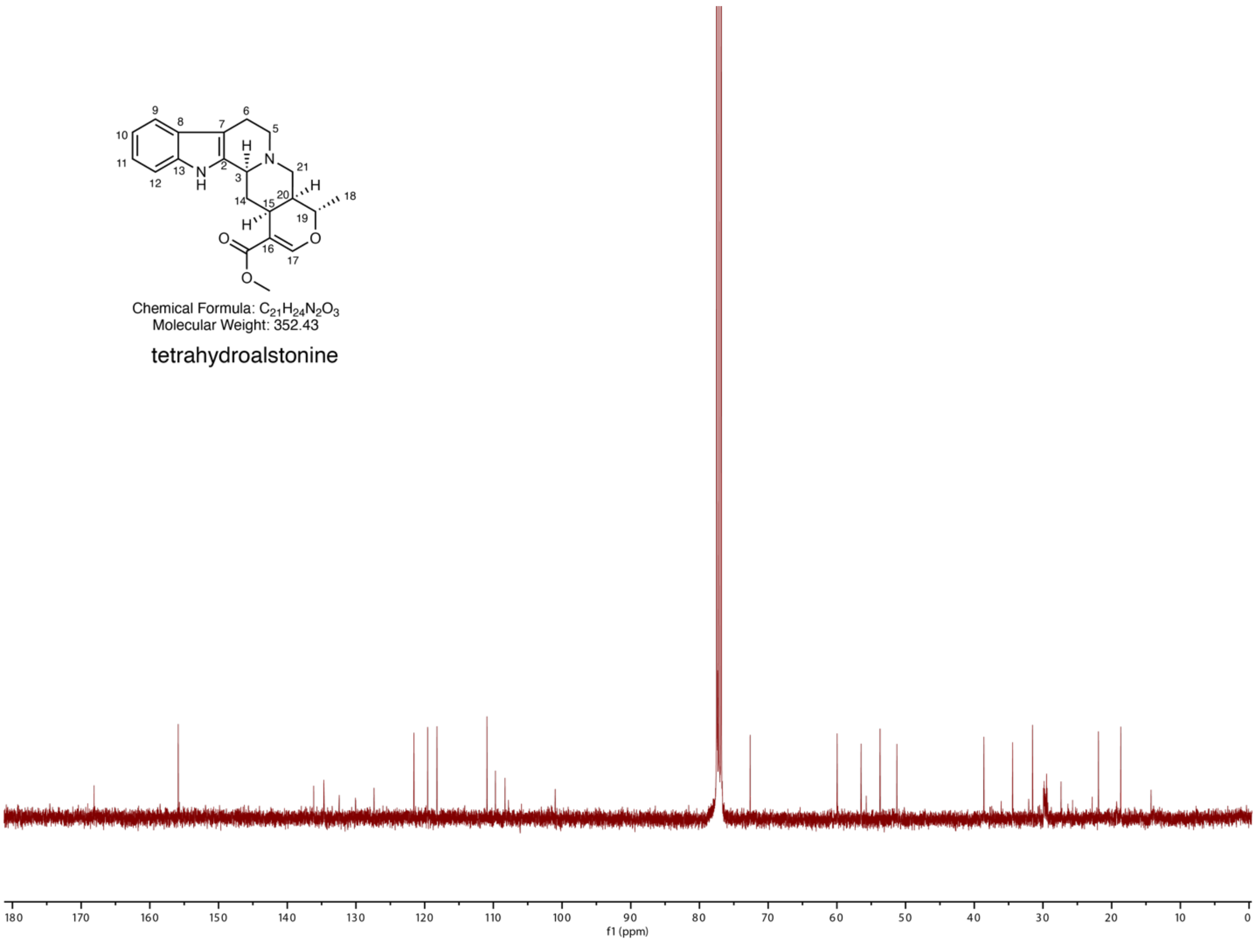

**Supplementary Figure 63. $^{13}C$ NMR spectra of tetrahydroalstonine in $CDCl_3$.**

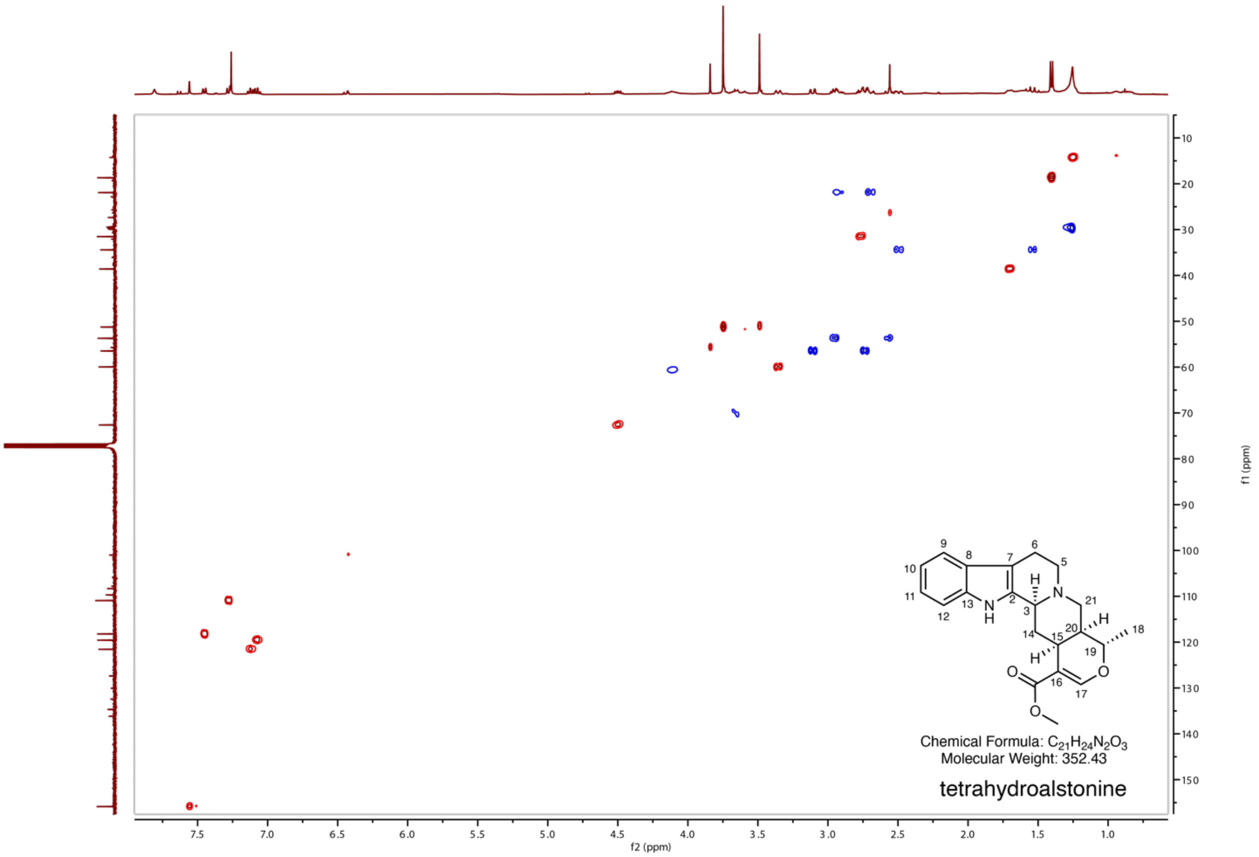

**Supplementary Figure 64. HSQC NMR spectra of tetrahydroalstonine in $CDCl_3$.**

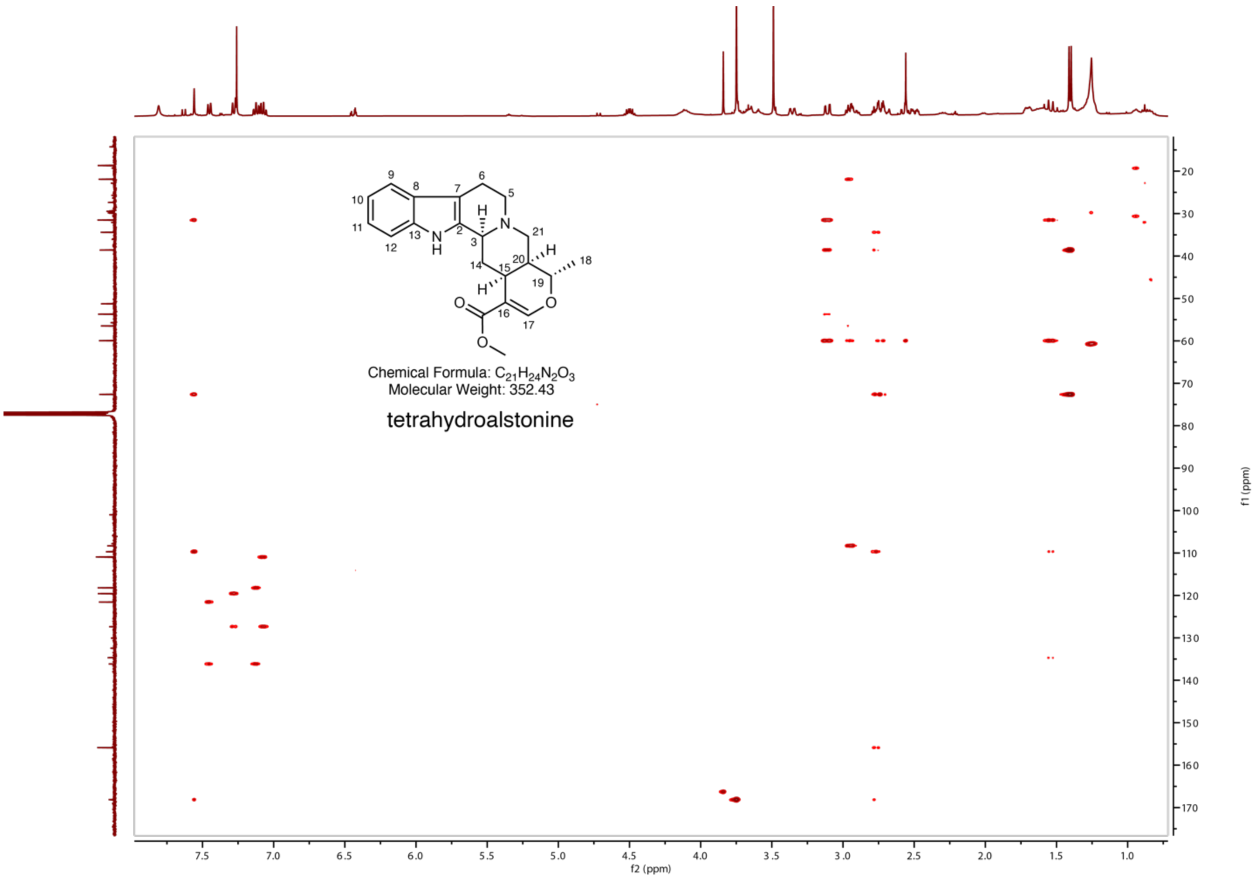

**Supplementary Figure 65. HMBC NMR spectra of tetrahydroalstonine in $CDCl_3$.**

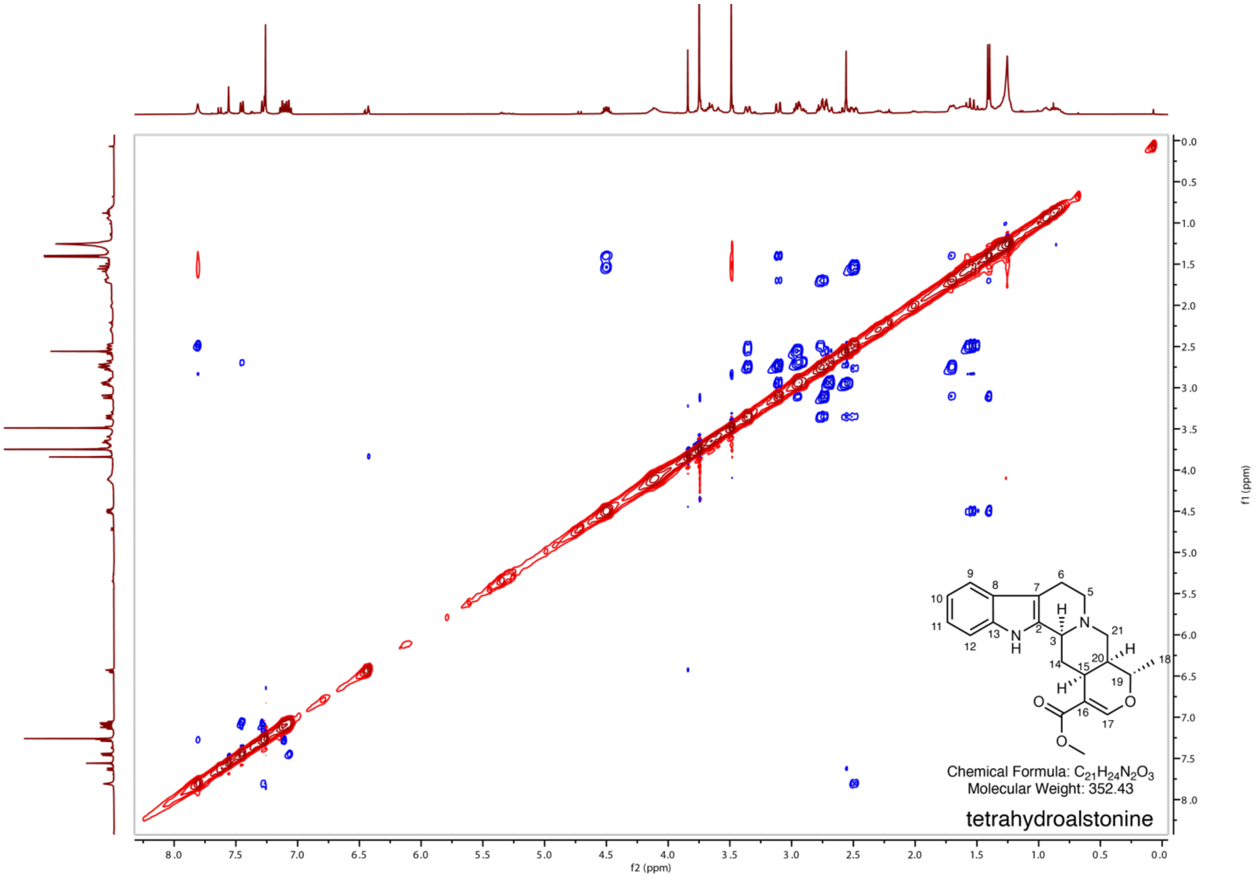

**Supplementary Figure 66. NOESY NMR spectra of tetrahydroalstonine in $CDCl_3$.**

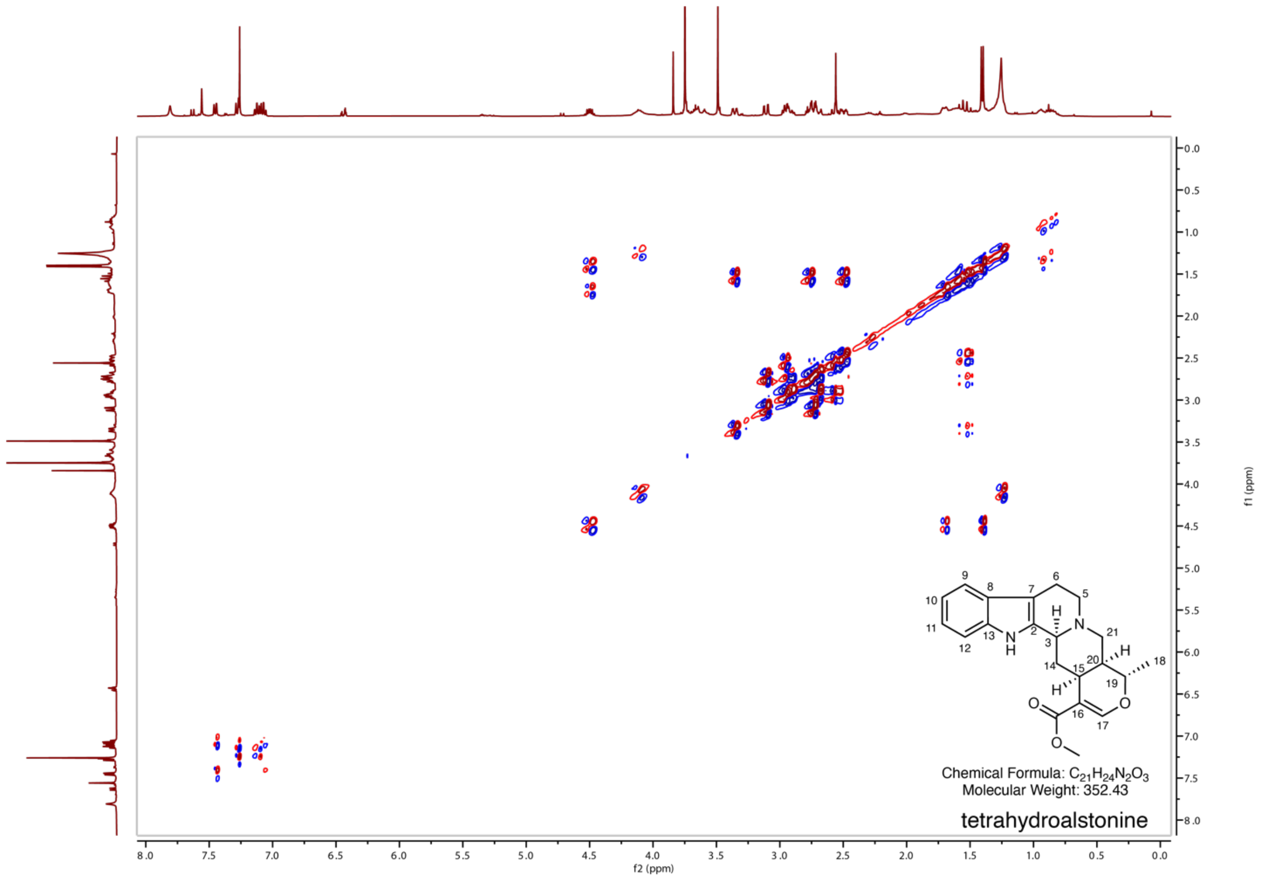

**Supplementary Figure 67. COSY NMR spectra of tetrahydroalstonine in $CDCl_3$.**

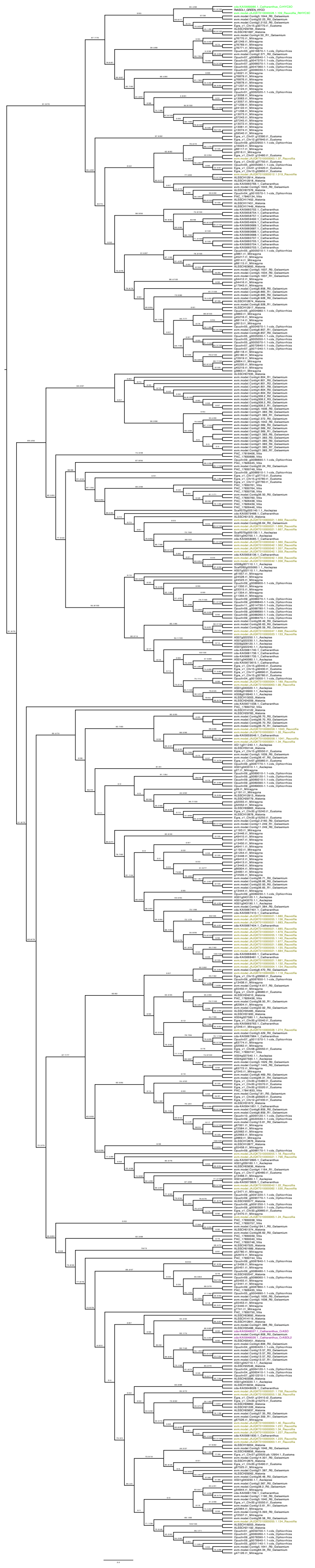

**Supplementary Figure 68. Midpoint-rooted phylogenetic tree of HYC3O, ASO, and other BBE-like homologs in members of Gentianales and *Vitis vinifera*.**
See attached PDF file in Supplementary data for a larger representation of the phylogenetic tree. Gene family representatives from *the Rauvolfia tetraphylla* reserpine BGC were used as CoGeBLAST query sequences against the proteomes of *Mitragyna speciosa, Gelsemium elegans, Catharanthus roseus, Alstonia scholaris, Rauvolfia tetraphylla, Asclepias syriaca, Ophiorrhiza pumila, Eustoma grandiflorum,* and *Vitis vinifera*. Tip colors (green) correspond to syntenic gene colorations on the MCScan graphics. Ochre tips are used for all other *Rauvolfia* genes, as well as genes that are shared between species but not part of a defined BGC or functionally characterized cluster. Sequences were aligned as proteins using Muscle and trimmed using GBlocks through the sequence processing tool Seaview, and the tree was reconstructed with IQTree 1.6.12 using the LG substitution model and 1000 bootstrap replicates for support values.

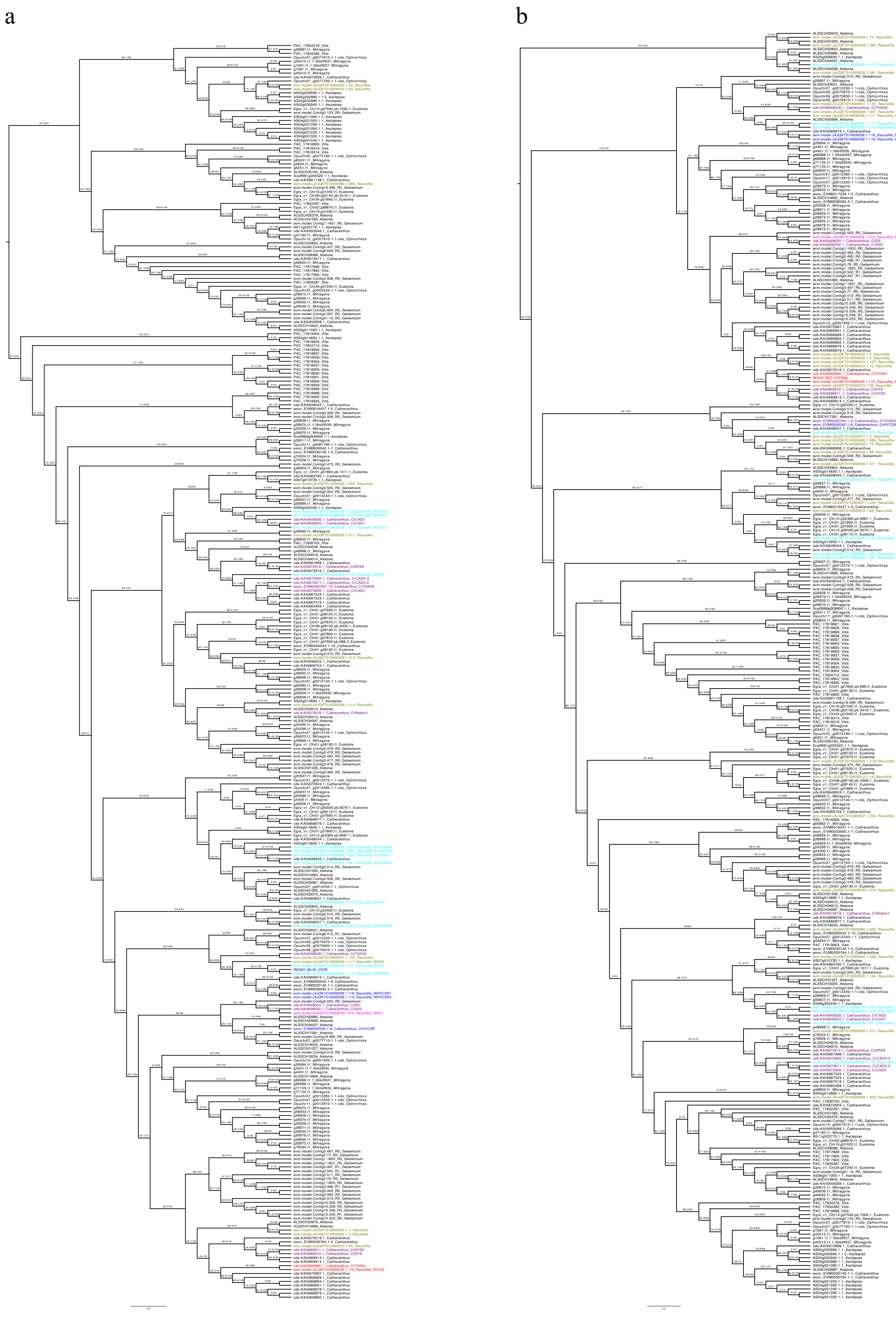
a
b

**Supplementary Figure 69. Midpoint-rooted phylogenetic trees of (a) HYC3R with other CAD-like reductases, and (b) YOS/GS with other CAD-like reductases.**
See attached PDF file in Supplementary data for a larger representation of the phylogenetic tree. Gene family representatives from *the Rauvolfia tetraphylla* reserpine BGC were used as CoGeBLAST query sequences against the proteomes of *Mitragyna speciosa, Gelsemium elegans, Catharanthus roseus, Alstonia scholaris, Rauvolfia tetraphylla, Asclepias syriaca, Ophiorrhiza pumila, Eustoma grandiflorum,* and *Vitis vinifera*. Tip colors correspond to syntenic gene colorations on the MCScan graphics, except for proteins from the Redox1/DPAS and HYS/THAS4 clusters, which are colored in purple, and the GS gene is highlighted in magenta. Ochre tips are used for all other *Rauvolfia* genes, as well as genes that are shared between species but not part of a defined BGC or functionally characterized cluster; these may include homologs of *Rauvolfia*-specific *CAD/CAD-like* genes (cyan). Sequences were aligned as proteins using Muscle and trimmed using GBlocks through the sequence processing tool Seaview, and the trees were reconstructed with IQTree 1.6.12 using the LG substitution model and 1000 bootstrap replicates for support values.

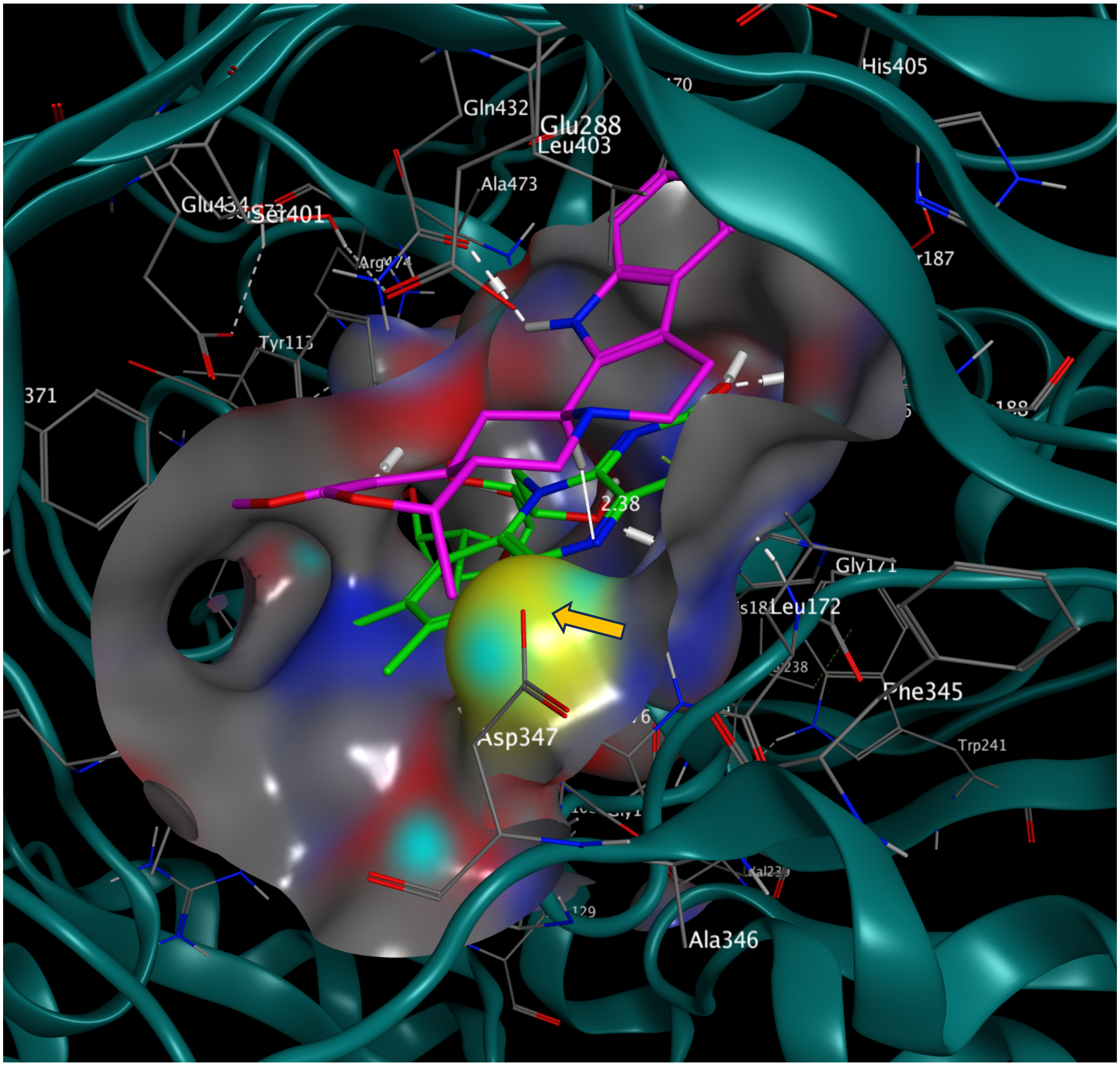

**Supplementary Figure 70. Ajmalicine at the active site of RtHYC3O.**
The hydrogen bonds (white dashed lines) between Q432 and indole NH, along with Tyr113 and the carbonyl, ensure the correct substrate orientation, facilitating close contact between substrate H3 and FAD N5 (2.38 Å). Docking score -7.3854 kcal/mol. The active site surface is illustrated with the following color scheme: oxygen (red), nitrogen (blue), sulfur (yellow), and carbon (grey). FAD is shown in green, and the alkaloid substrate is shown in magenta. Hydrogen bonds are indicated by white dashed lines. The orange arrows indicate the surface of C173 that covalently bonds FAD.

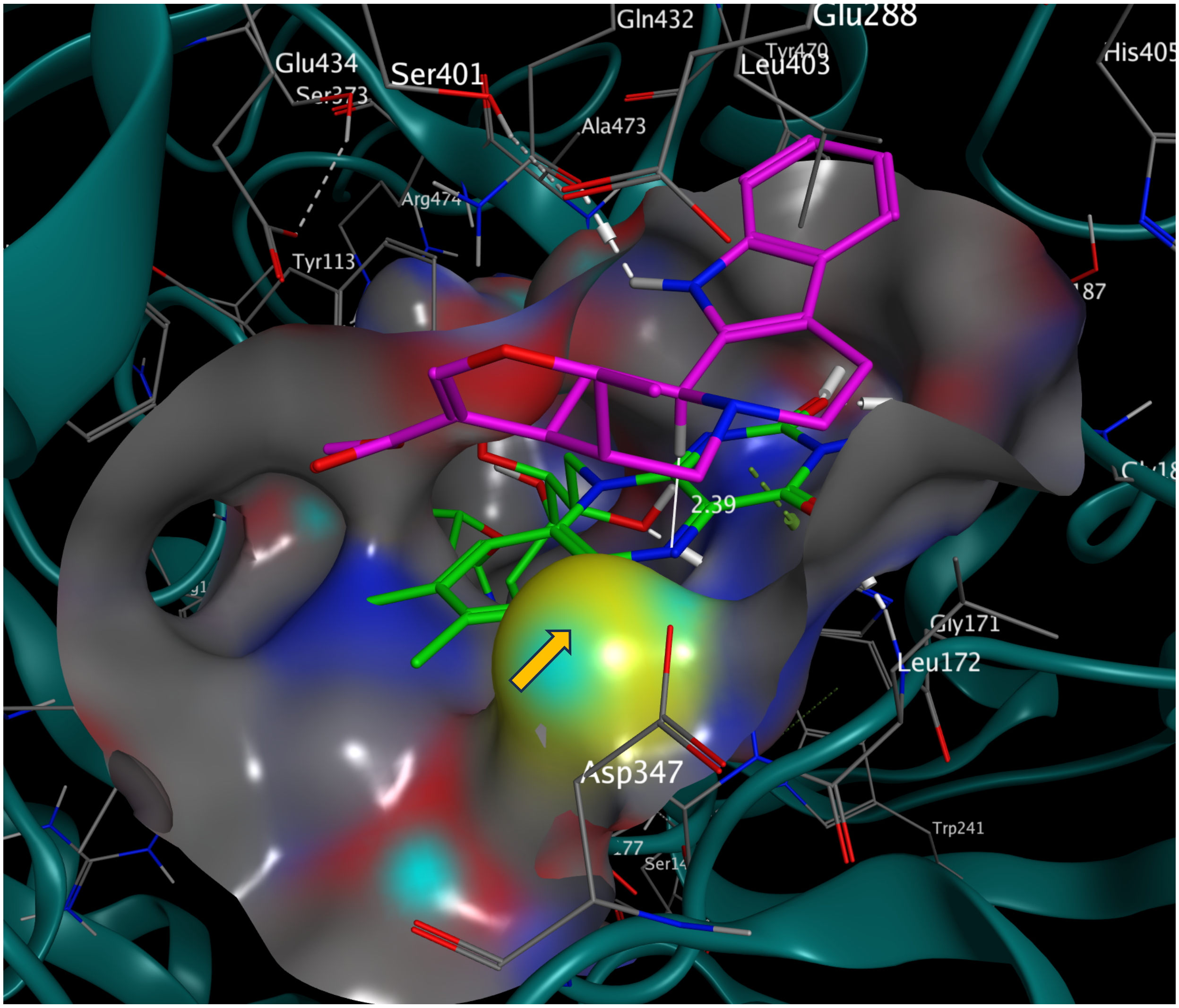

**Supplementary Figure 71. Tetrahydroalstonine at the active site of RtHYC3O.**
The hydrogen bonds (white dashed lines) between Q432 and indole NH ensures the correct substrate orientation, facilitating close contact between substrate H3 and FAD N5 (2.39 Å). Docking score -7.1937 kcal/mol. The active site surface is illustrated with the following color scheme: oxygen (red), nitrogen (blue), sulfur (yellow), and carbon (grey). FAD is shown in green, and the alkaloid substrate is shown in magenta. Hydrogen bonds are indicated by white dashed lines. The orange arrows indicate the surface of C173 that covalently bonds FAD.

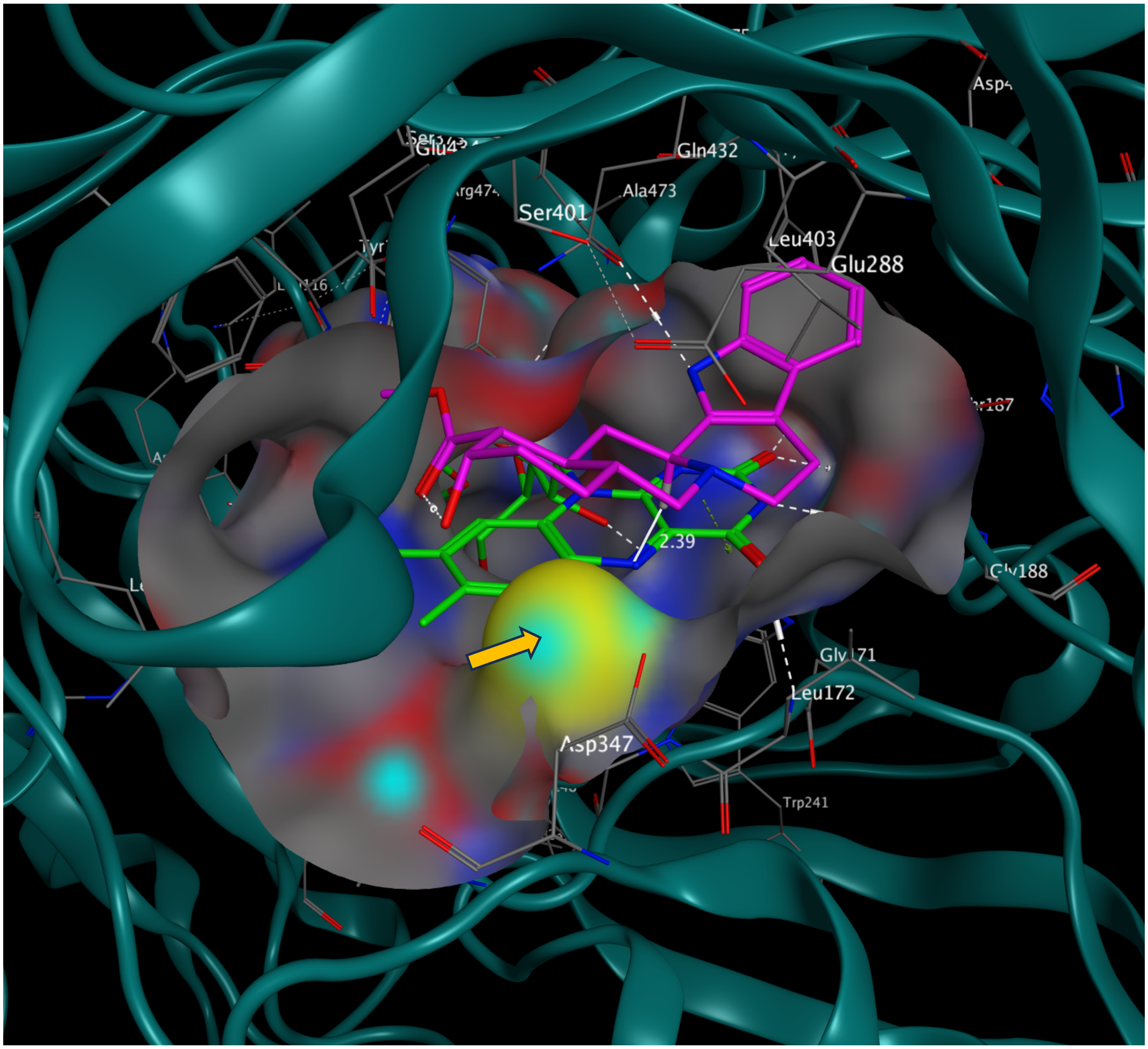

**Supplementary Figure 72. Yohimbine at the active site of RtHYC3O.**
The hydrogen bonds (white dashed lines) between Q432 and indole NH ensure the correct substrate orientation, facilitating close contact between substrate H3 and FAD N5 (2.39 Å). Docking score -7.0916 kcal/mol. The active site surface is illustrated with the following color scheme: oxygen (red), nitrogen (blue), sulfur (yellow), and carbon (grey). FAD is shown in green, and the alkaloid substrate is shown in magenta. Hydrogen bonds are indicated by white dashed lines. The orange arrows indicate the surface of C173 that covalently bonds FAD.

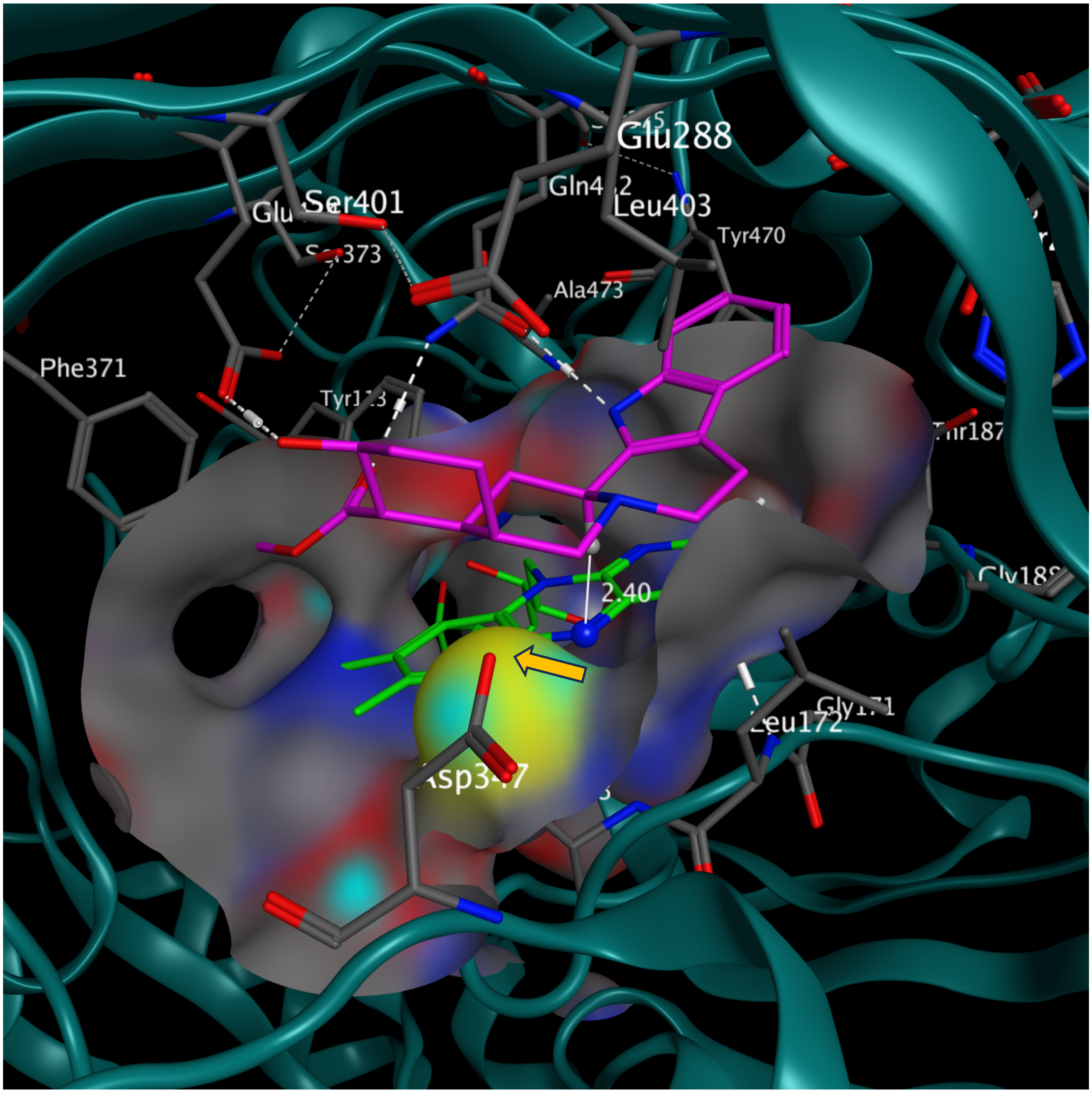

**Supplementary Figure 73. Rauwolscine at the active site of RtHYC3O.**
Three hydrogen bonds (white dashed lines) between Q432/E434 and rauwolscine's indole, carbomethoxy, and 17-OH ensured the correct substrate orientation, facilitating close contact between substrate H3 and FAD N5 (2.40 Å). Docking score -7.4154 kcal/mol. The active site surface is illustrated with the following color scheme: oxygen (red), nitrogen (blue), sulfur (yellow), and carbon (grey). FAD is shown in green, and the alkaloid substrate is shown in magenta. Hydrogen bonds are indicated by white dashed lines. The orange arrows indicate the surface of C173 that covalently bonds FAD.

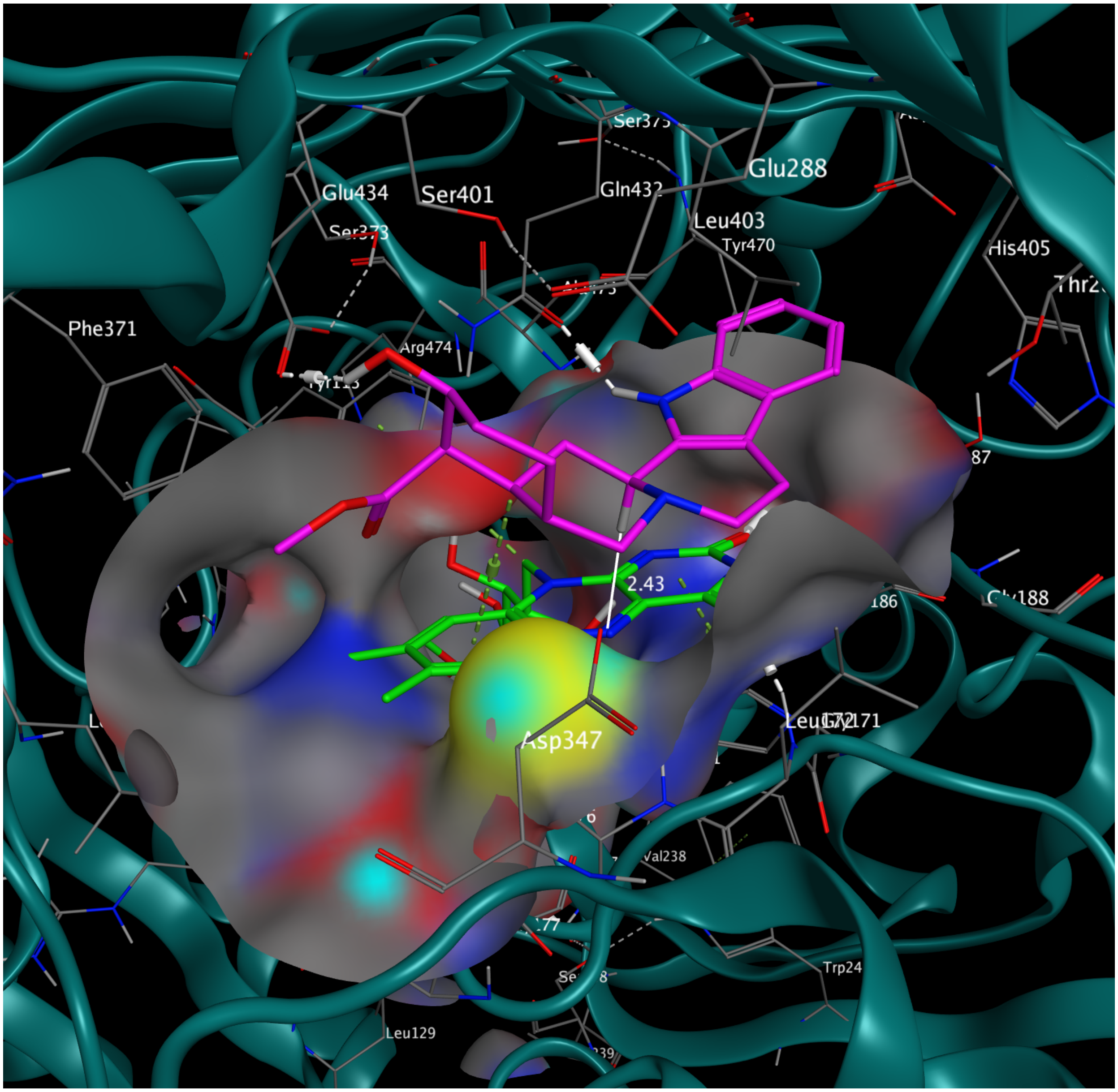

**Supplementary Figure 74. Alloyohimbine at the active site of RtHYC3O.**
The hydrogen bonds (white dashed lines) between Q432 and indole NH, as well as E434 and 17-OH ensure the correct substrate orientation, facilitating close contact between substrate H3 and FAD N5 (2.43 Å). Docking score -7.2922 kcal/mol. The active site surface is illustrated with the following color scheme: oxygen (red), nitrogen (blue), sulfur (yellow), and carbon (grey). FAD is shown in green, and the alkaloid substrate is shown in magenta. Hydrogen bonds are indicated by white dashed lines. The orange arrows indicate the surface of C173 that covalently bonds FAD.

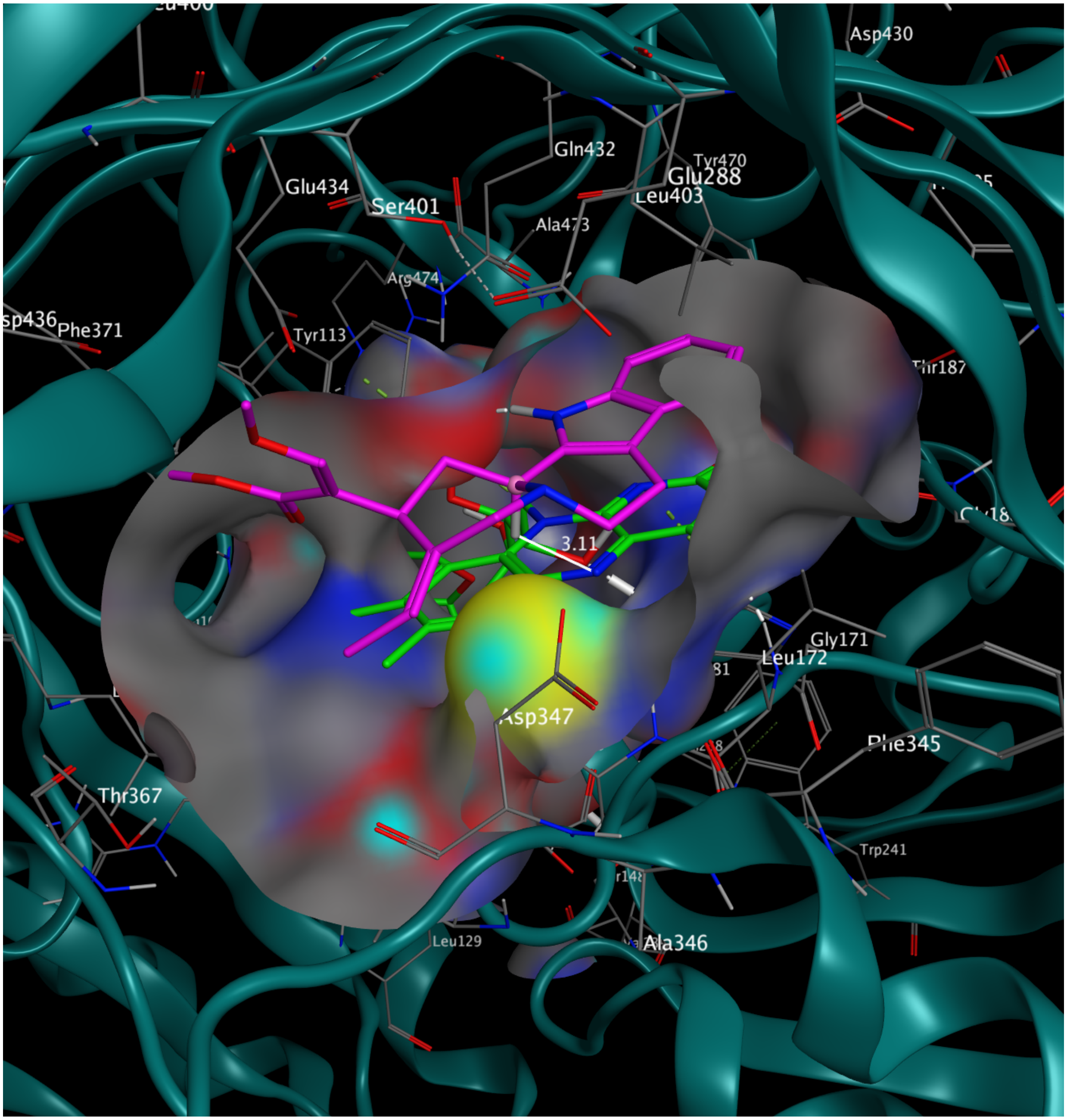

**Supplementary Figure 75. Geissoschizine methyl ether at the active site of RtHYC3O.** The hydrogen bonds (white dashed lines) between Q432 and indole NH contributes to the correct substrate orientation, facilitating reactionn between substrate H3 and FAD N5. The relatively larger distance (3.11 Å) is consistent with lower substrate preference. Docking score -7.3815 kcal/mol. The active site surface is illustrated with the following color scheme: oxygen (red), nitrogen (blue), sulfur (yellow), and carbon (grey). FAD is shown in green, and the alkaloid substrate is shown in magenta. Hydrogen bonds are indicated by white dashed lines. The orange arrows indicate the surface of C173 that covalently bonds FAD.

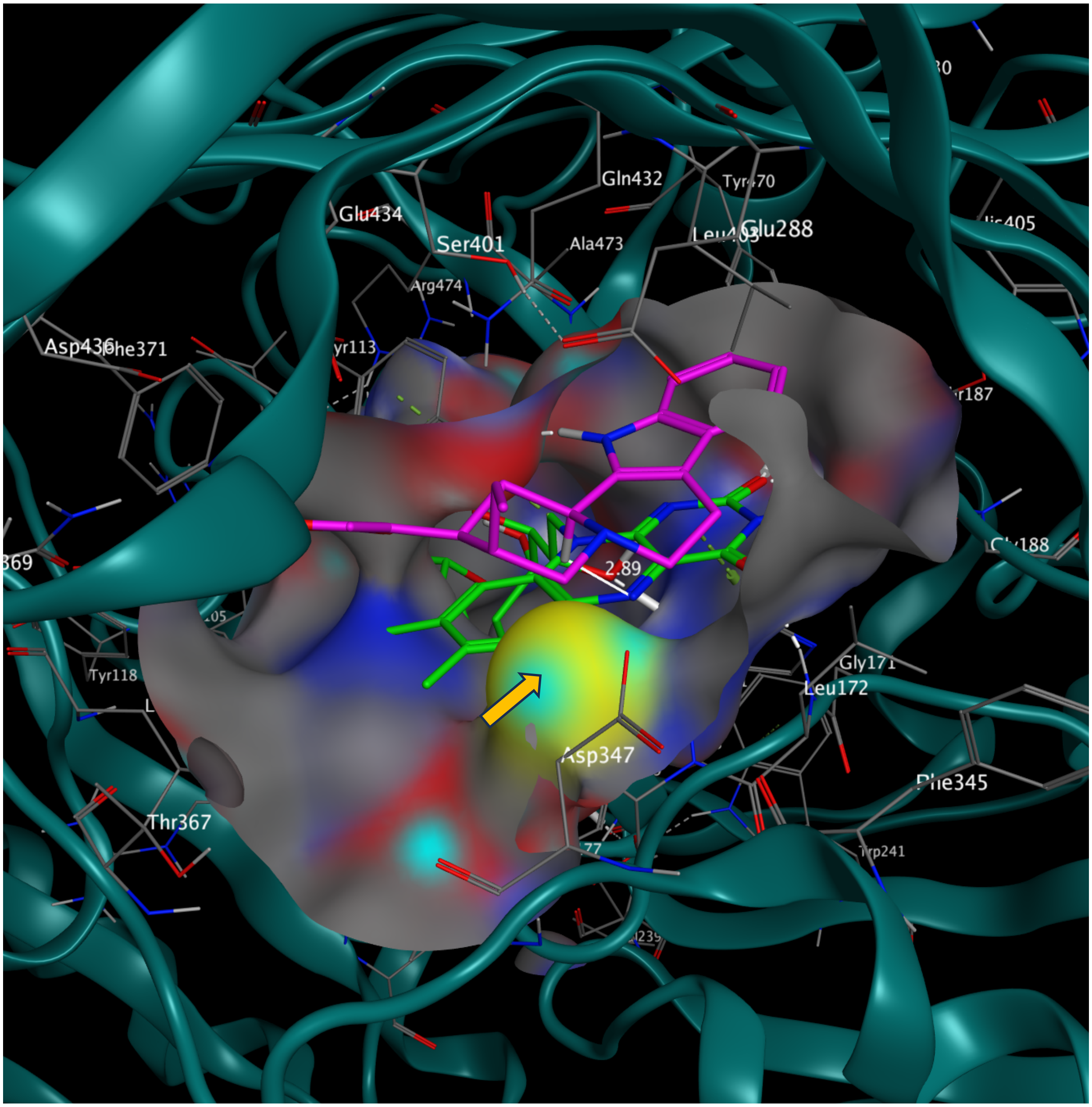

**Supplementary Figure 76. Corynanthedine at the active site of RtHYC3O.**
The hydrogen bonds (white dashed lines) between Q432 and indole NH contributes to the correct substrate orientation, facilitating reactionn between substrate H3 and FAD N5. The relatively larger distance (2.97 Å) is consistent with lower substrate preference. Docking score -7.5552 kcal/mol. The active site surface is illustrated with the following color scheme: oxygen (red), nitrogen (blue), sulfur (yellow), and carbon (grey). FAD is shown in green, and the alkaloid substrate is shown in magenta. Hydrogen bonds are indicated by white dashed lines. The orange arrows indicate the surface of C173 that covalently bonds FAD.

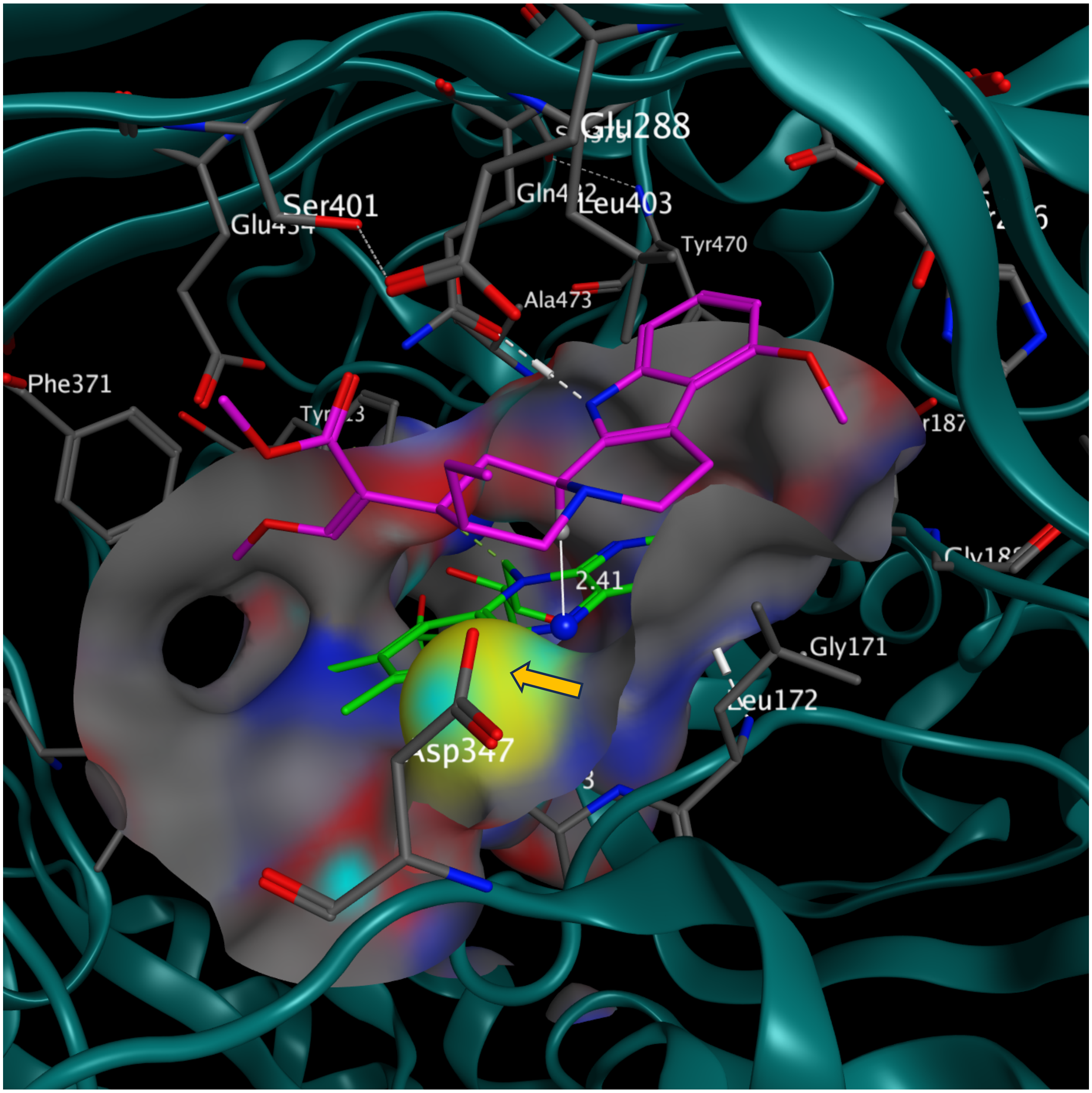

**Supplementary Figure 77. Mitragynine at the active site of RtHYC3O.**
Three hydrogen bonds (white dashed lines) between Q432 and indole NH ensures the correct substrate orientation, facilitating close contact between substrate H3 and FAD N5 (2.41 Å). The 9-methoxyl group fits into a spacious binding pocket. Docking score -8.0934 kcal/mol. The active site surface is illustrated with the following color scheme: oxygen (red), nitrogen (blue), sulfur (yellow), and carbon (grey). FAD is shown in green, and the alkaloid substrate is shown in magenta. Hydrogen bonds are indicated by white dashed lines. The orange arrows indicate the surface of C173 that covalently bonds FAD.

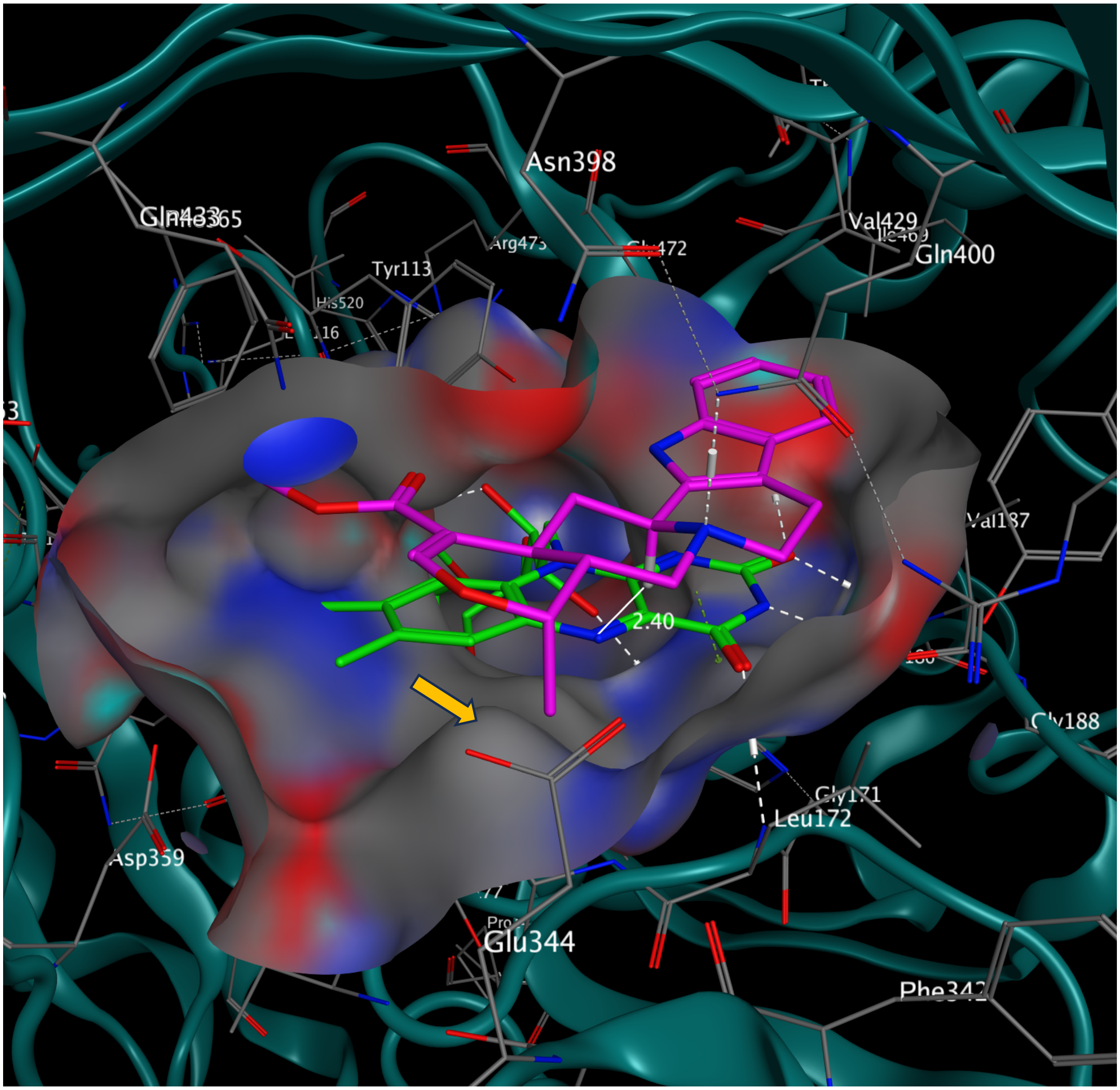

**Supplementary Figure 78. Ajmalicine at the active site of HpHYC3O.**
The C173A substitution (orange arrow) contributed to a spacious active site accommodating all seven tested substrates. Despite a single FAD-His covalent bond, FAD conformation is still tightly oriented by multiple hydrogen bonds (dotted lines) with HpHYC3O. Hydrogen bonds (white dashed lines) among Q400, R286, N398, and that between Q400 and substrate's N4 facilitated proper docking, facilitating close contact between substrate H3 and FAD N5 (2.40 Å). Docking score -8.2801 kcal/mol. The active site surface is illustrated with the following color scheme: oxygen (red), nitrogen (blue), sulfur (yellow), and carbon (grey). FAD is shown in green, and the alkaloid substrate is shown in magenta. Hydrogen bonds are indicated by white dashed lines.

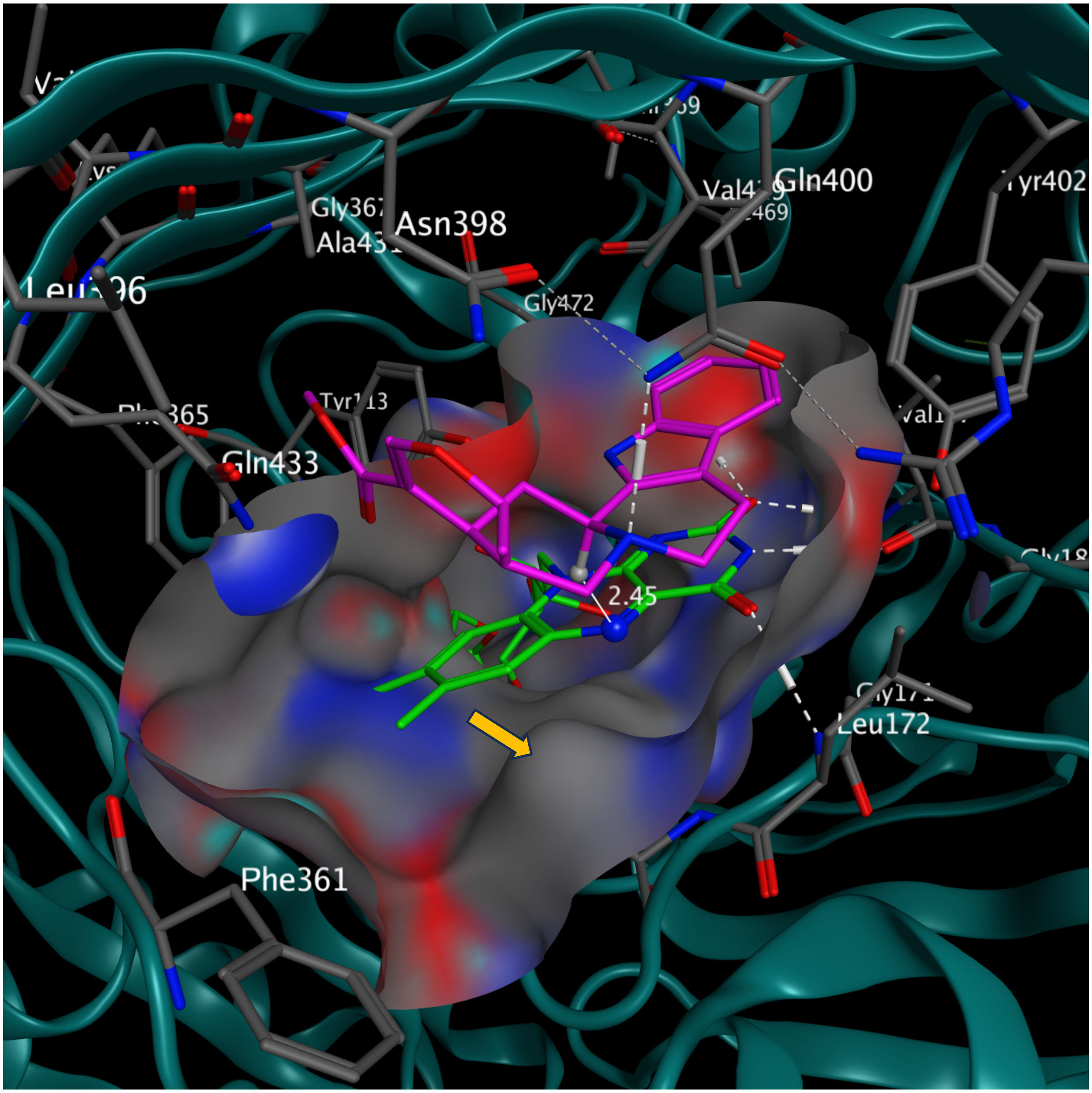

**Supplementary Figure 79. Tetrahydroalstonine at the active site of HpHYC3O.**
The C173A substitution (orange arrow) contributed to a spacious active site accommodating all seven tested substrates. Despite a single FAD-His covalent bond, FAD conformation is still tightly oriented by multiple hydrogen bonds (dotted lines) with HpHYC3O. Hydrogen bonds (white dashed lines) among Q400, R286, N398, and that between Q400 and substrate's N4 facilitated proper docking, facilitating close contact between substrate H3 and FAD N5 (2.45 Å). Docking score -8.2109 kcal/mol. The active site surface is illustrated with the following color scheme: oxygen (red), nitrogen (blue), sulfur (yellow), and carbon (grey). FAD is shown in green, and the alkaloid substrate is shown in magenta. Hydrogen bonds are indicated by white dashed lines.

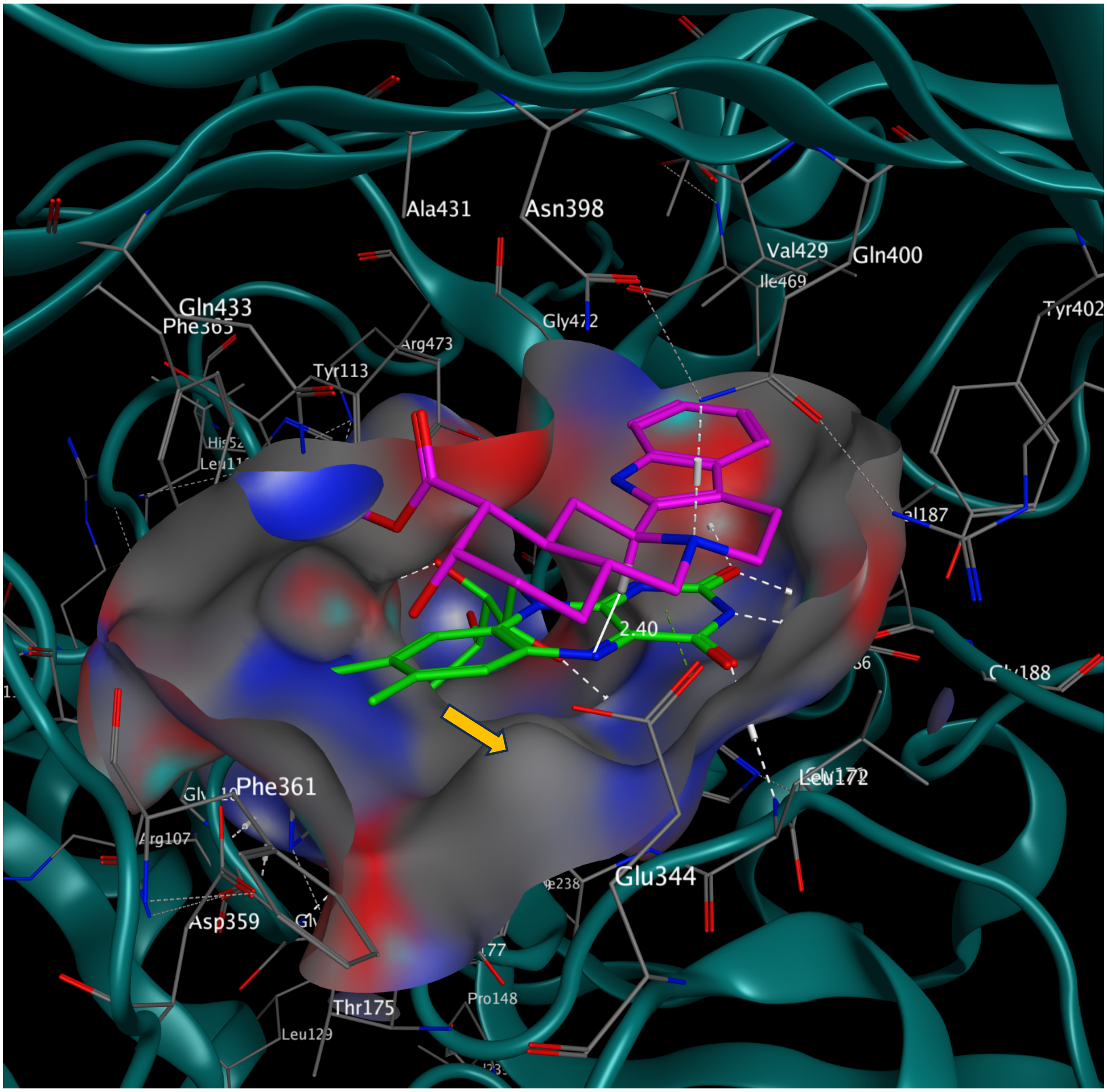

**Supplementary Figure 80. Yohimbine at the active site of HpHYC3O.**
The C173A substitution (orange arrow) contributed to a spacious active site accommodating all seven tested substrates. Despite a single FAD-His covalent bond, FAD conformation is still tightly oriented by multiple hydrogen bonds (dotted lines) with HpHYC3O. Hydrogen bonds (white dashed lines) among Q400, R286, N398, and that between Q400 and substrate's N4 facilitated proper docking, facilitating close contact between substrate H3 and FAD N5 (2.40 Å). Docking score -8.1000 kcal/mol. The active site surface is illustrated with the following color scheme: oxygen (red), nitrogen (blue), sulfur (yellow), and carbon (grey). FAD is shown in green, and the alkaloid substrate is shown in magenta. Hydrogen bonds are indicated by white dashed lines.

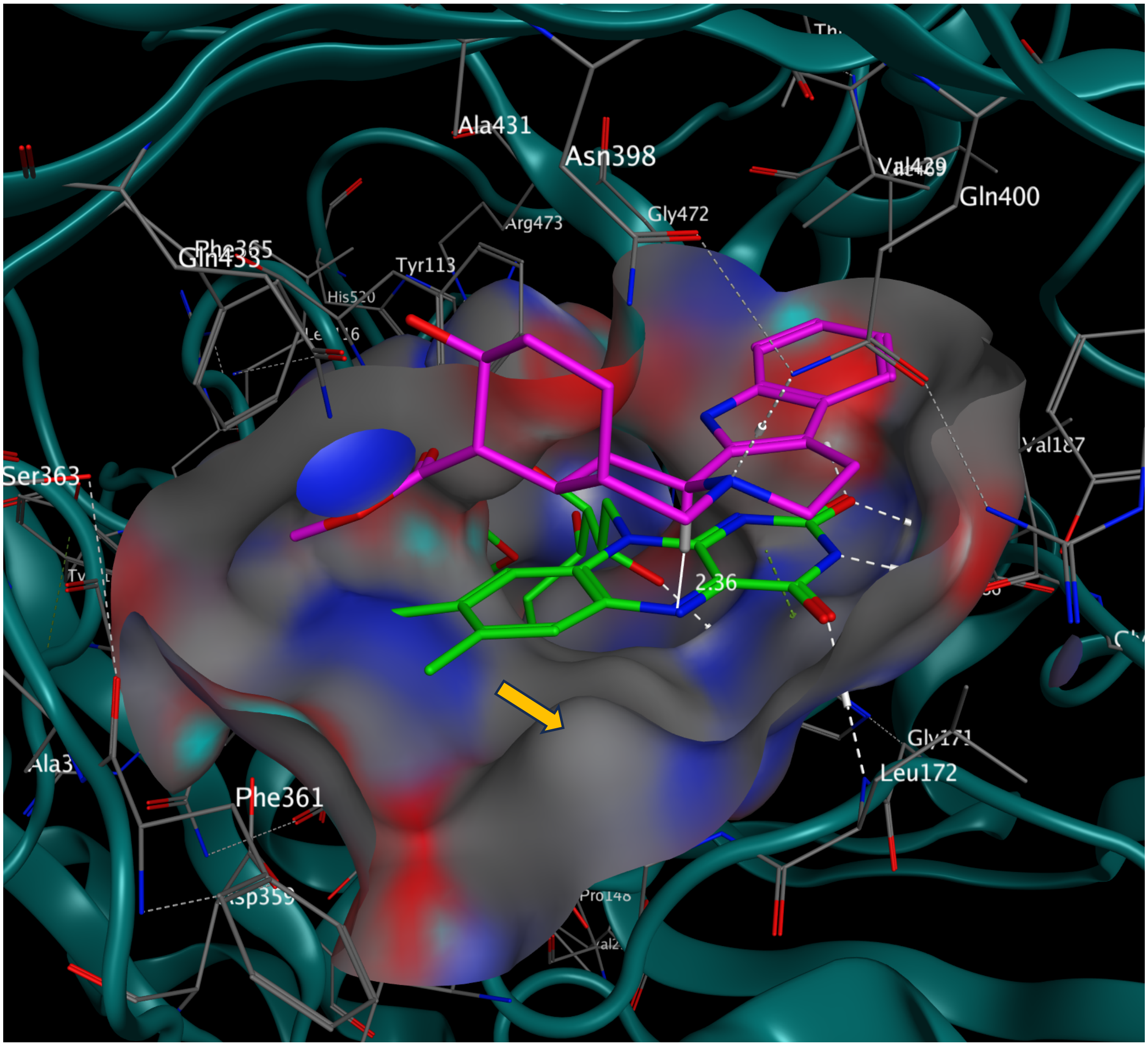

**Supplementary Figure 81. Rauwolscine at the active site of HpHYC3O.**
The C173A substitution (orange arrow) contributed to a spacious active site accommodating all seven tested substrates. Despite a single FAD-His covalent bond, FAD conformation is still tightly oriented by multiple hydrogen bonds (dotted lines) with HpHYC3O. Hydrogen bonds (white dashed lines) among Q400, R286, N398, and that between Q400 and substrate's N4 facilitated proper docking, facilitating close contact between substrate H3 and FAD N5 (2.36 Å). Docking score -8.4088 kcal/mol. The active site surface is illustrated with the following color scheme: oxygen (red), nitrogen (blue), sulfur (yellow), and carbon (grey). FAD is shown in green, and the alkaloid substrate is shown in magenta. Hydrogen bonds are indicated by white dashed lines.

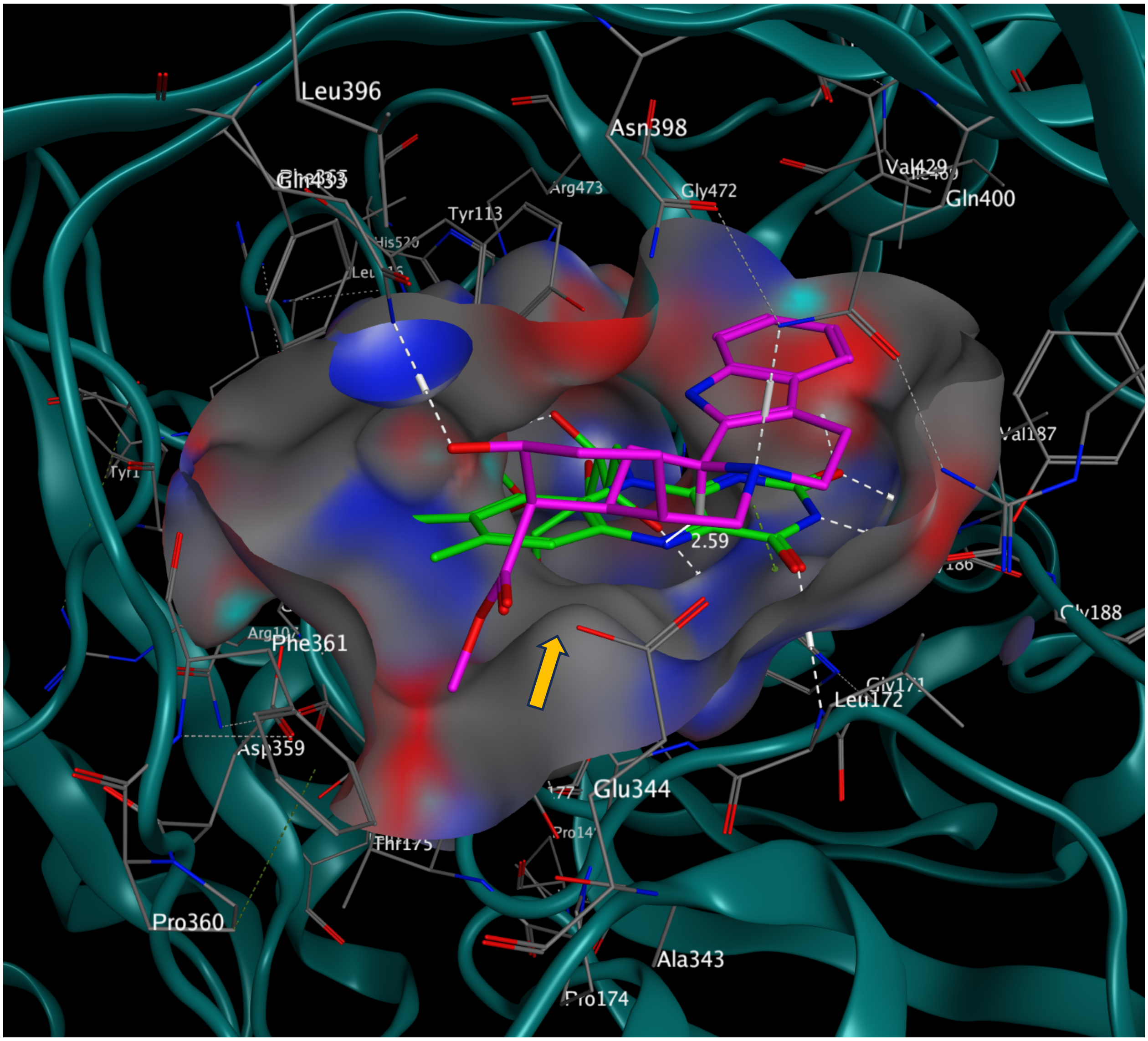

**Supplementary Figure 82. Alloyohimbine at the active site of HpHYC3O.**
The C173A substitution (orange arrow) contributed to a spacious active site accommodating all seven tested substrates. Despite a single FAD-His covalent bond, FAD conformation is still tightly oriented by multiple hydrogen bonds (dotted lines) with HpHYC3O. Hydrogen bonds (white dashed lines) among Q400, R286, N398, and that between Q400 and substrate's N4, along with Q433 and 17-OH, facilitated proper docking, facilitating close contact between substrate H3 and FAD N5 (2.59 Å). Docking score -8.0375 kcal/mol. The active site surface is illustrated with the following color scheme: oxygen (red), nitrogen (blue), sulfur (yellow), and carbon (grey). FAD is shown in green, and the alkaloid substrate is shown in magenta. Hydrogen bonds are indicated by white dashed lines.

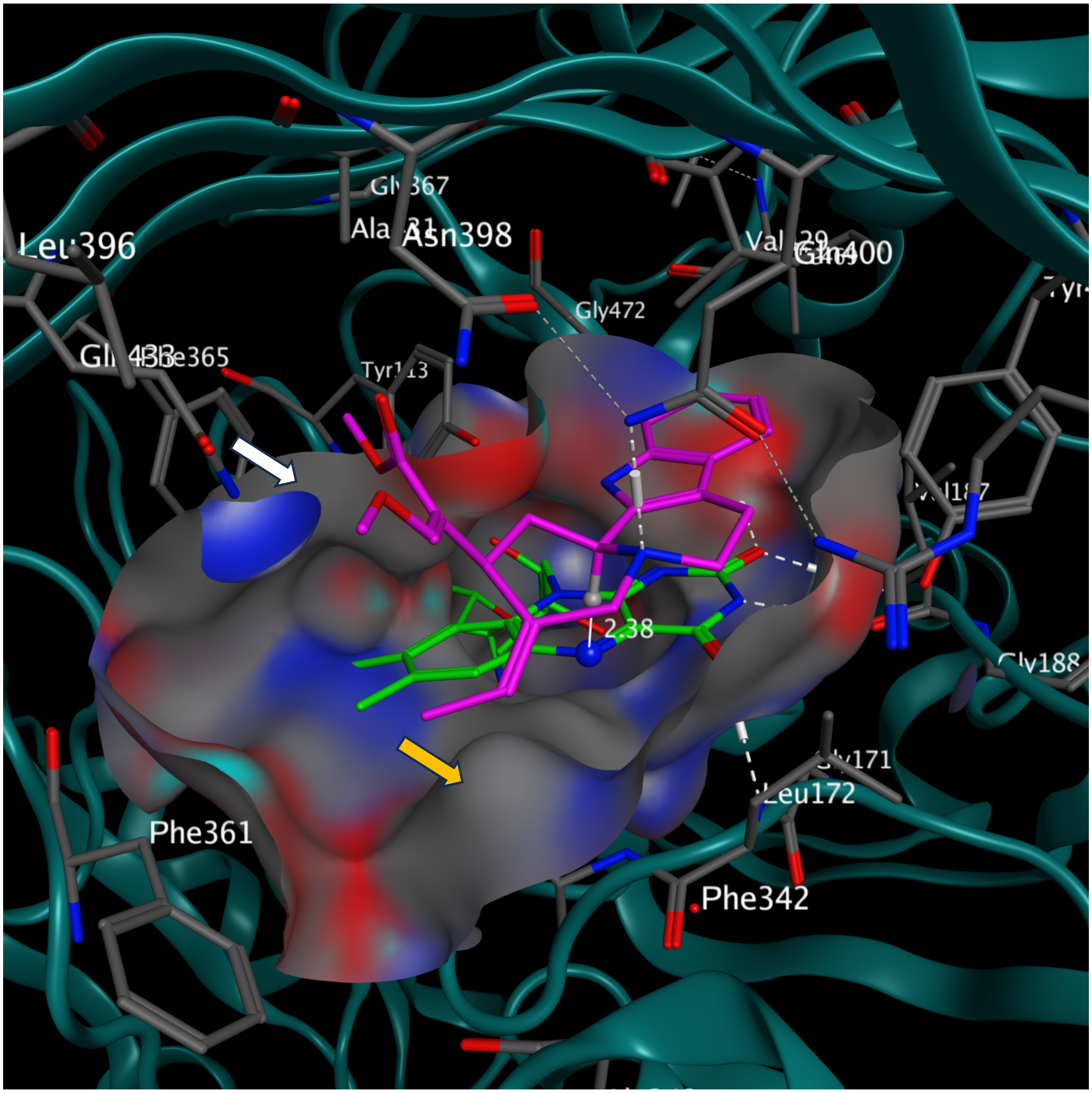

**Supplementary Figure 83. Geissoschizine methyl ether at the active site of HpHYC3O.** The C173A substitution (orange arrow) contributed to a spacious active site accommodating all seven tested substrates. The steric hindrance between C173 and the ethylidene side chain would not allow proper geissoschizine docking. The E431A substitution (white arrow) in HpHYC3O also created a binding pocket that could accommodate the carbomethoxy group in geissoschizine methyl ether. Despite a single FAD-His covalent bond, FAD conformation is still tightly oriented by multiple hydrogen bonds (dotted lines) with HpHYC3O. Hydrogen bonds (white dashed lines) among Q400, R286, N398, and that between Q400 and substrate's N4 facilitated proper docking, facilitating close contact between substrate H3 and FAD N5 (2.38 Å). Docking score -8.4512 kcal/mol. The active site surface is illustrated with the following color scheme: oxygen (red), nitrogen (blue), sulfur (yellow), and carbon (grey). FAD is shown in green, and the alkaloid substrate is shown in magenta. Hydrogen bonds are indicated by white dashed lines.

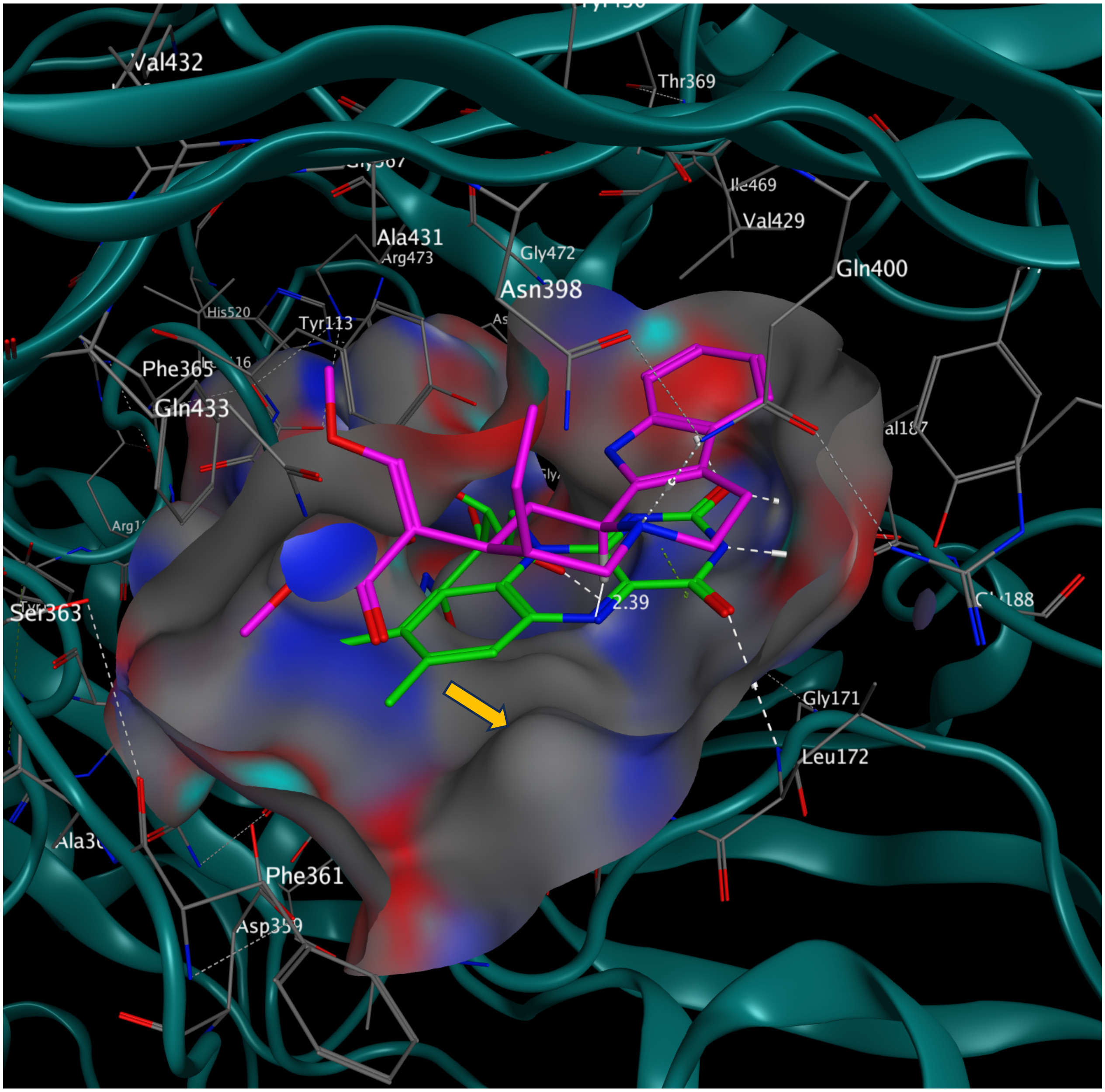

**Supplementary Figure 84. Corynantheidine at the active site of HpHYC3O.**
The C173A substitution (orange arrow) contributed to a spacious active site accommodating all seven tested substrates. Despite a single FAD-His covalent bond, FAD conformation is still tightly oriented by multiple hydrogen bonds (dotted lines) with HpHYC3O. Hydrogen bonds (white dashed lines) among Q400, R286, N398, and that between Q400 and substrate's N4 facilitated proper docking, facilitating close contact between substrate H3 and FAD N5 (2.39 Å). Docking score -8.7615 kcal/mol. The active site surface is illustrated with the following color scheme: oxygen (red), nitrogen (blue), sulfur (yellow), and carbon (grey). FAD is shown in green, and the alkaloid substrate is shown in magenta. Hydrogen bonds are indicated by white dashed lines.

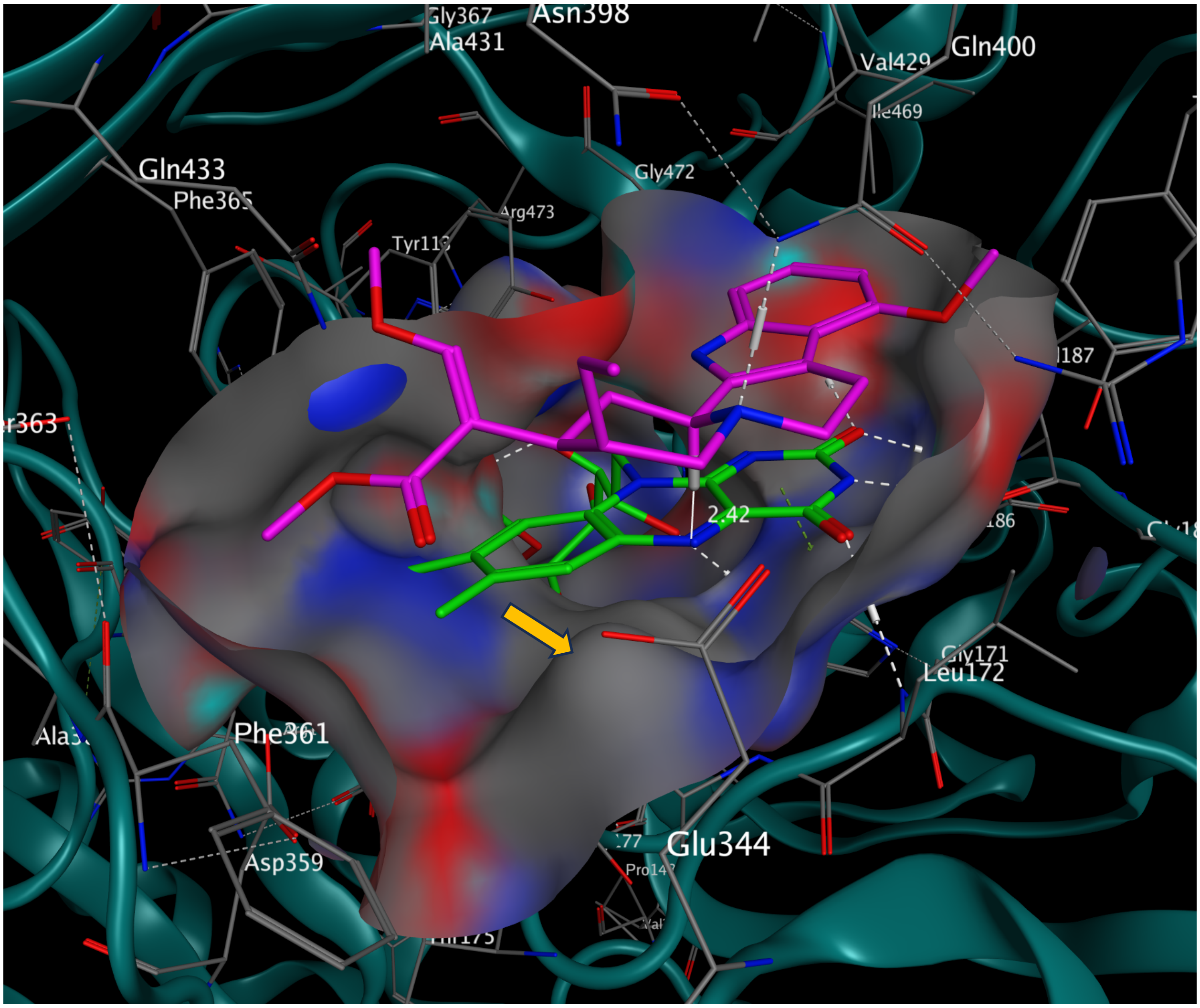

**Supplementary Figure 85. Mitragynine at the active site of HpHYC3O.**
The C173A substitution (orange arrow) contributed to a spacious active site accommodating all seven tested substrates. Despite a single FAD-His covalent bond, FAD conformation is still tightly oriented by multiple hydrogen bonds (dotted lines) with HpHYC3O. Hydrogen bonds (white dashed lines) among Q400, R286, N398, and that between Q400 and substrate's N4 facilitated proper docking, facilitating close contact between substrate H3 and FAD N5 (2.42 Å). Docking score -8.9653 kcal/mol. The active site surface is illustrated with the following color scheme: oxygen (red), nitrogen (blue), sulfur (yellow), and carbon (grey). FAD is shown in green, and the alkaloid substrate is shown in magenta. Hydrogen bonds are indicated by white dashed lines.

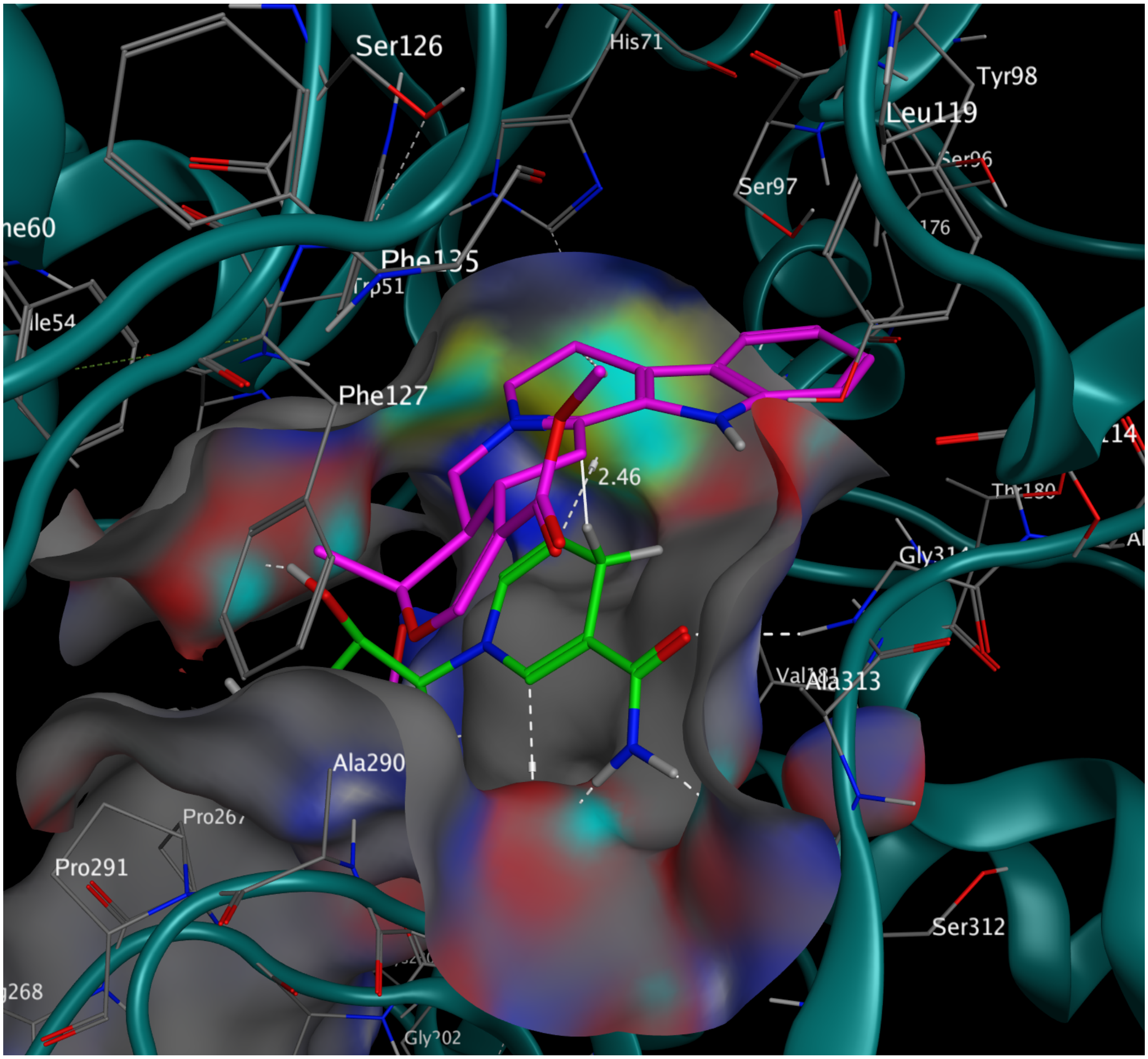

**Supplementary Figure 86. 3-dehydro-ajmalicine at the active site of RsHYC3R.**
The substrate binding is mainly facilitated by van der Waals interactions at the active site, conducive for C3 hydride transfer from NADPH (2.46 Å, white solid line). Docking score -6.5944 kcal/mol. The active site surface is illustrated with the following color scheme: oxygen (red), nitrogen (blue), sulfur (yellow), and carbon (grey). NADPH is shown in green, and the alkaloid substrate is shown in magenta. Hydrogen bonds are indicated by white dashed lines.

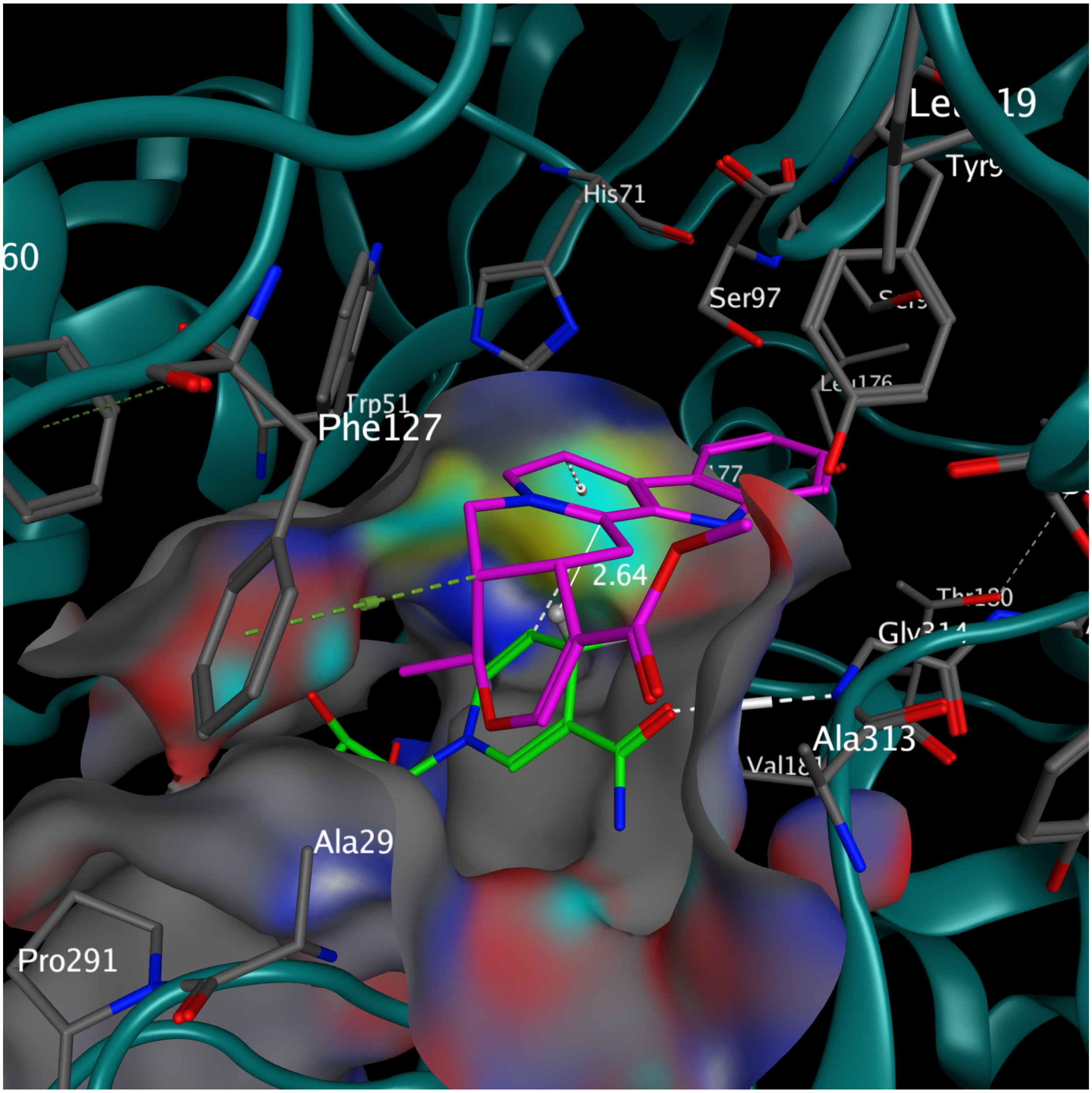

**Supplementary Figure 87. 3-dehydro-tetrahydroalstonine at the active site of RsHYC3R.** The substrate binding is mainly facilitated by van der Waals interactions at the active site, conducive for C3 hydride transfer from NADPH (2.64 Å, white solid line). Docking score -6.8943 kcal/mol. The active site surface is illustrated with the following color scheme: oxygen (red), nitrogen (blue), sulfur (yellow), and carbon (grey). NADPH is shown in green, and the alkaloid substrate is shown in magenta. Hydrogen bonds are indicated by white dashed lines.

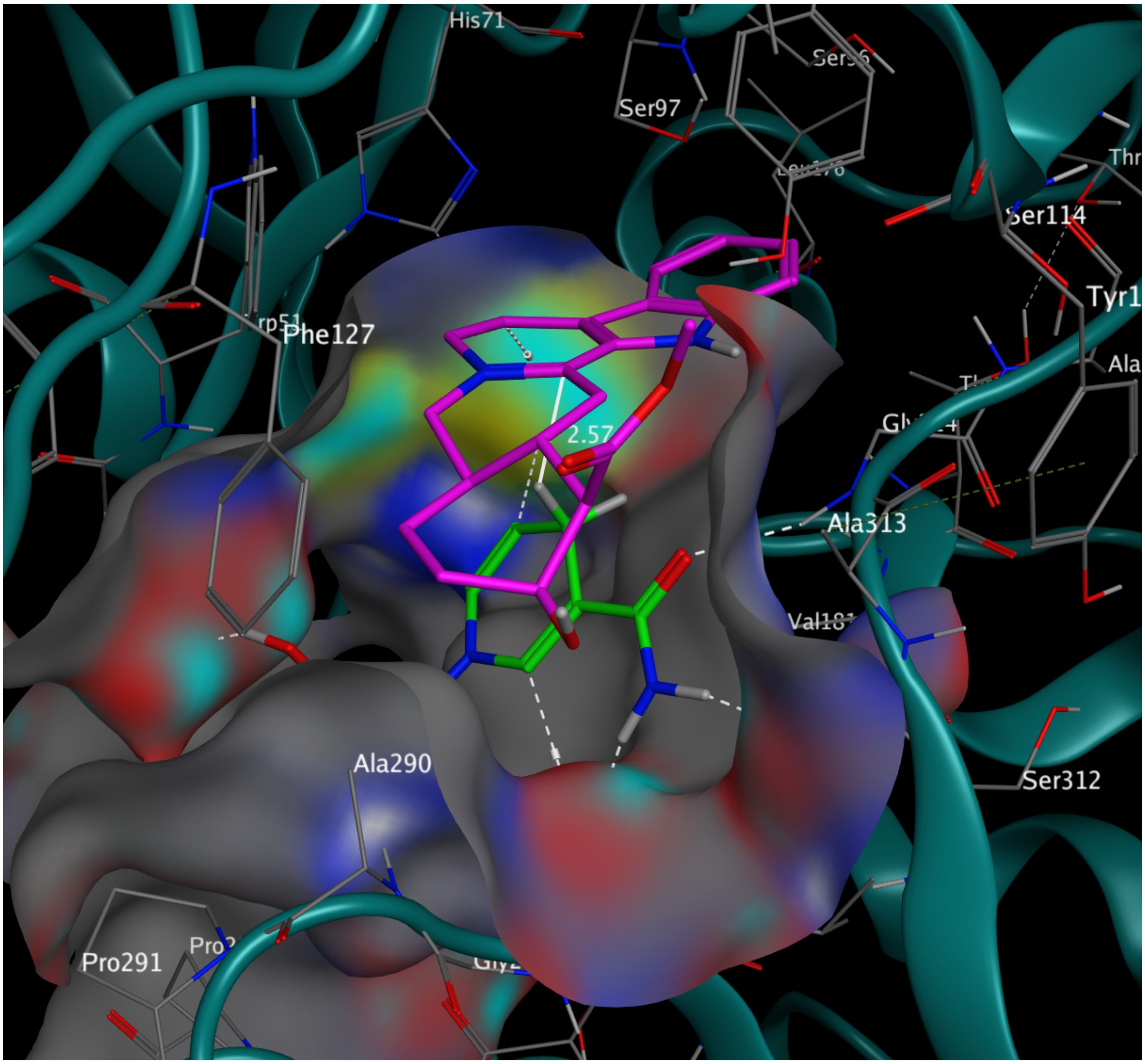

**Supplementary Figure 88. 3-dehydro-yohimbine at the active site of RsHYC3R.**
The substrate binding is mainly facilitated by van der Waals interactions at the active site, conducive for C3 hydride transfer from NADPH (2.57 Å, white solid line). Docking score -6.8923 kcal/mol. The active site surface is illustrated with the following color scheme: oxygen (red), nitrogen (blue), sulfur (yellow), and carbon (grey). NADPH is shown in green, and the alkaloid substrate is shown in magenta. Hydrogen bonds are indicated by white dashed lines.

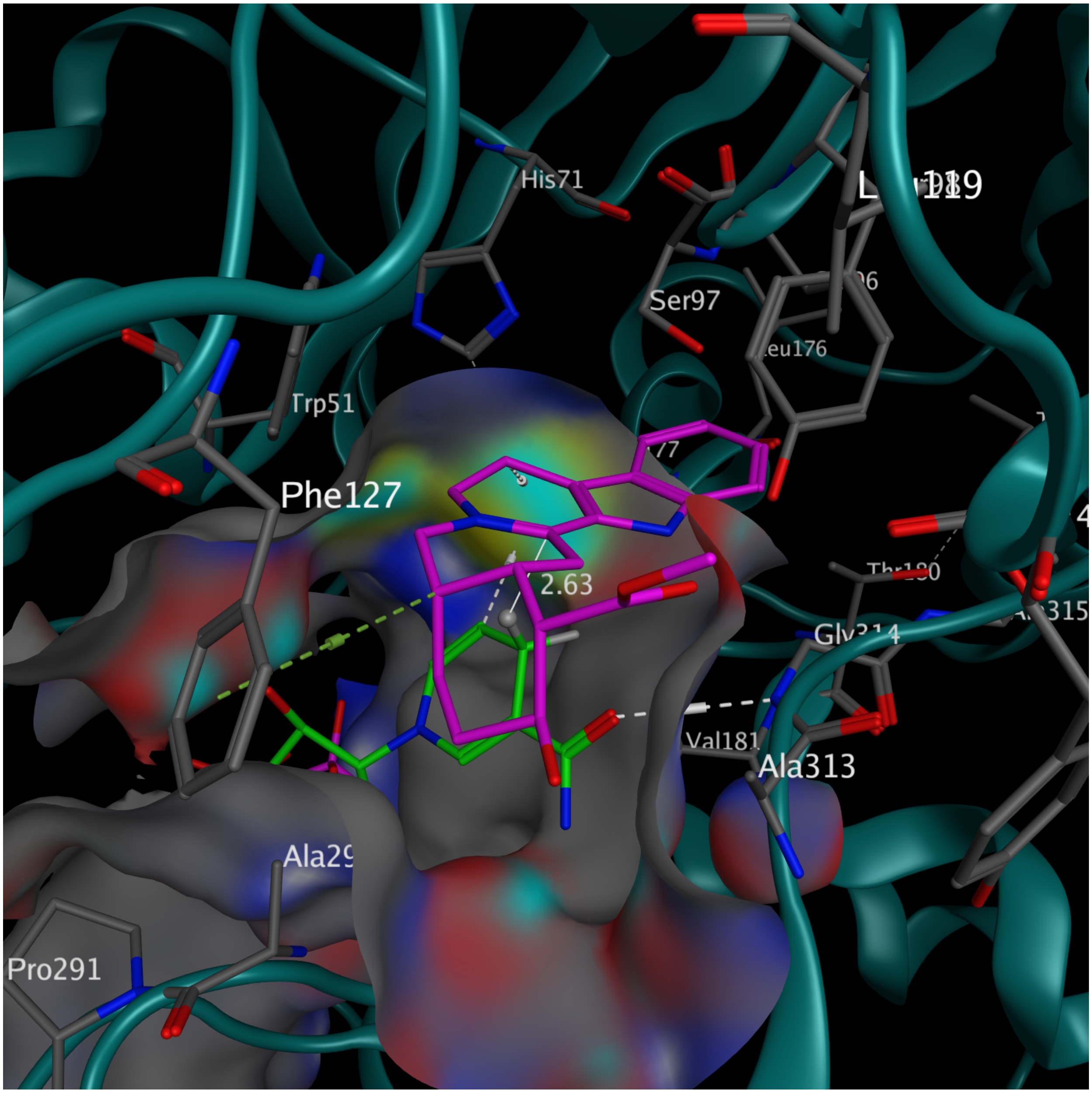

**Supplementary Figure 89. 3-dehydro-rauwolscine at the active site of RsHYC3R.**
The substrate binding is mainly facilitated by van der Waals interactions at the active site, conducive for C3 hydride transfer from NADPH (2.63 Å, white solid line). Docking score -6.7754 kcal/mol. The active site surface is illustrated with the following color scheme: oxygen (red), nitrogen (blue), sulfur (yellow), and carbon (grey). NADPH is shown in green, and the alkaloid substrate is shown in magenta. Hydrogen bonds are indicated by white dashed lines.

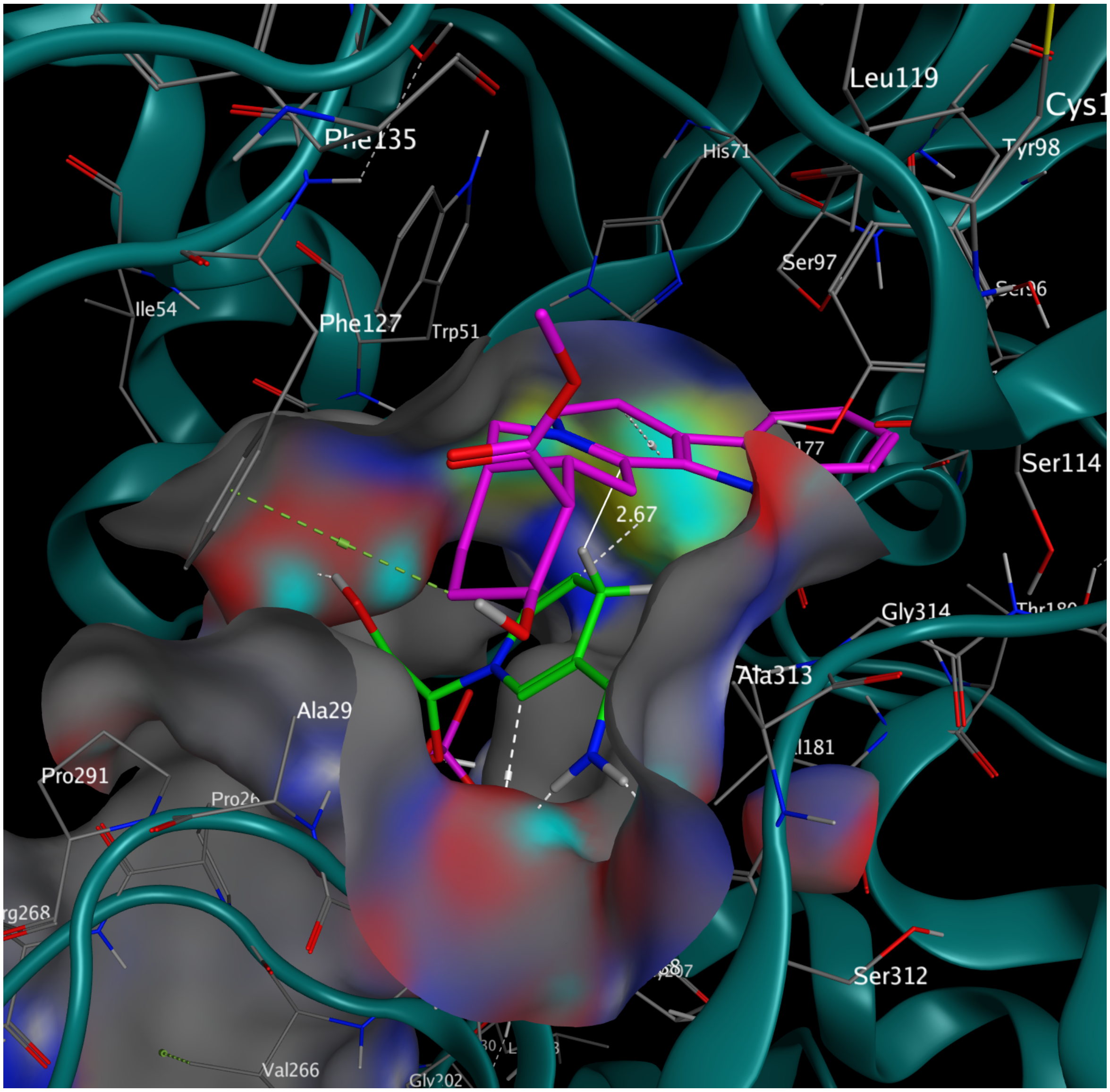

**Supplementary Figure 90. 3-dehydro-alloyohimbine at the active site of RsHYC3R.** The substrate binding is mainly facilitated by van der Waals interactions at the active site, conducive for C3 hydride transfer from NADPH (2.67 Å, white solid line). Docking score -6.7306 kcal/mol. The active site surface is illustrated with the following color scheme: oxygen (red), nitrogen (blue), sulfur (yellow), and carbon (grey). NADPH is shown in green, and the alkaloid substrate is shown in magenta. Hydrogen bonds are indicated by white dashed lines.

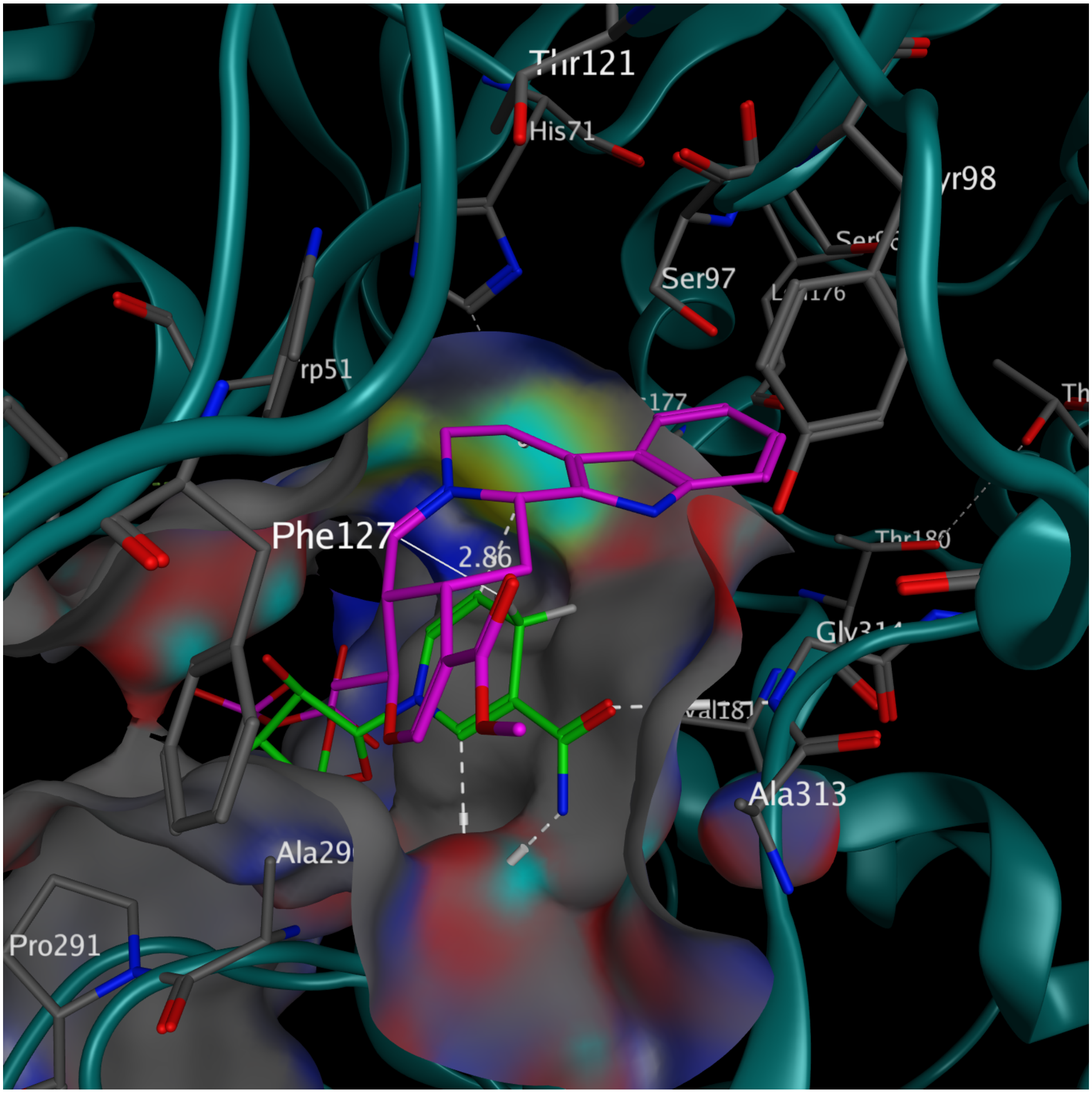

**Supplementary Figure 91. 4,21-dehydro-tetrahydroalstonine at the active site of RsHYC3R.** The substrate binding is mainly facilitated by van der Waals interactions at the active site, conducive for C3 hydride transfer from NADPH (2.86 Å, white solid line). The spacious binding site allow proper docking of both 4,21-dehydro-tetrahydroalstonine and 3, *N*4-dehydro-tetrahydroalstoninne. Docking score -6.8911 kcal/mol. The active site surface is illustrated with the following color scheme: oxygen (red), nitrogen (blue), sulfur (yellow), and carbon (grey). NADPH is shown in green, and the alkaloid substrate is shown in magenta. Hydrogen bonds are indicated by white dashed lines.

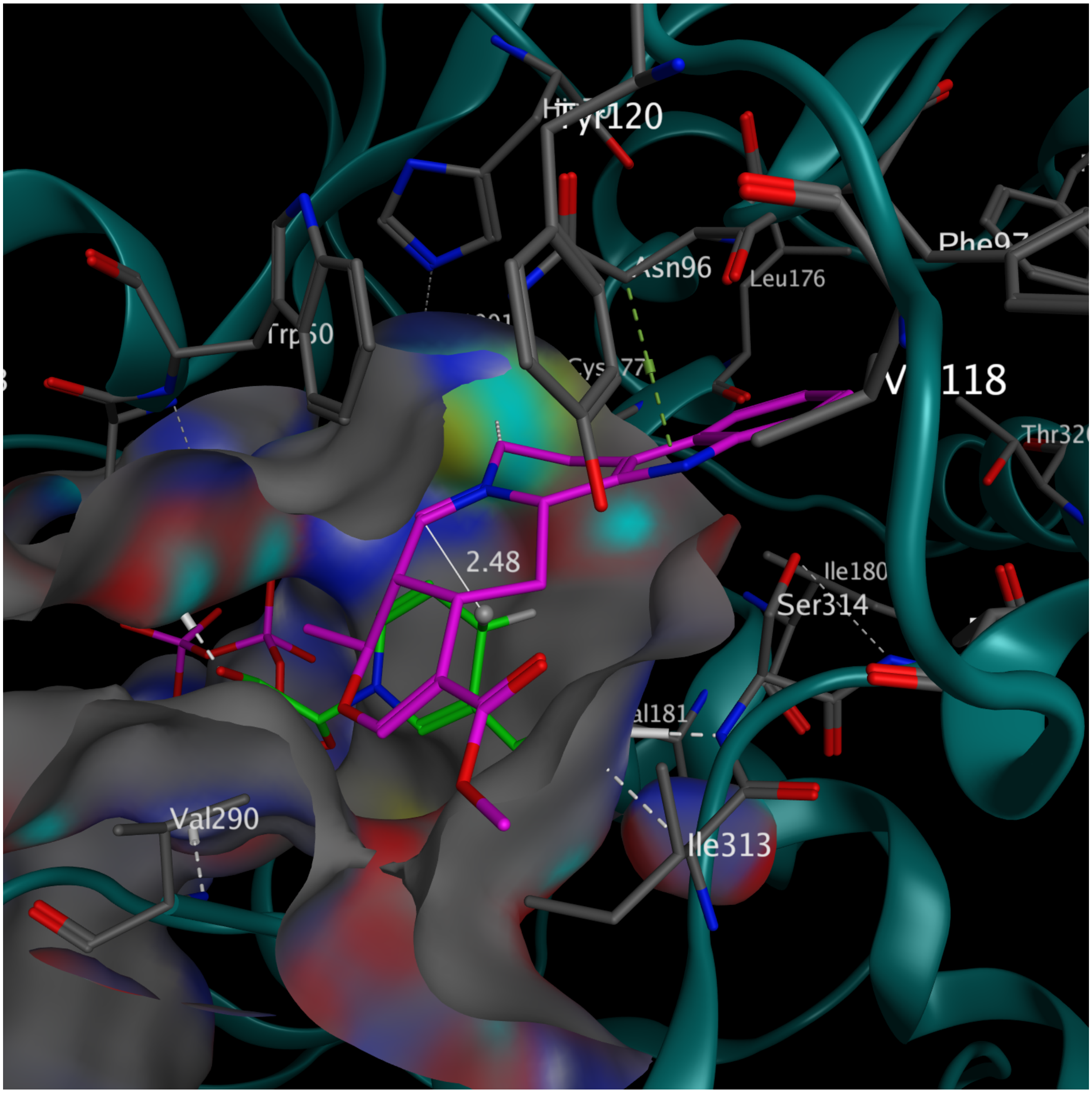

**Supplementary Figure 92. 4,21-dehydro-tetrahydroalstonine at the active site of CrTHAS2.** The substrate binding is mainly facilitated by van der Waals interactions at the active site, conducive for C3 hydride transfer from NADPH (2.48 Å, white solid line). While the active site accommodates 4,21-dehydro-tetrahydroalstonine, it is too constricted for docking 3, *N*4-dehydro-tetrahydroalstoninne, at least partially caused by the bulky V290, I313, and W60. Docking score -6.6511 kcal/mol. The active site surface is illustrated with the following color scheme: oxygen (red), nitrogen (blue), sulfur (yellow), and carbon (grey). NADPH is shown in green, and the alkaloid substrate is shown in magenta. Hydrogen bonds are indicated by white dashed lines.

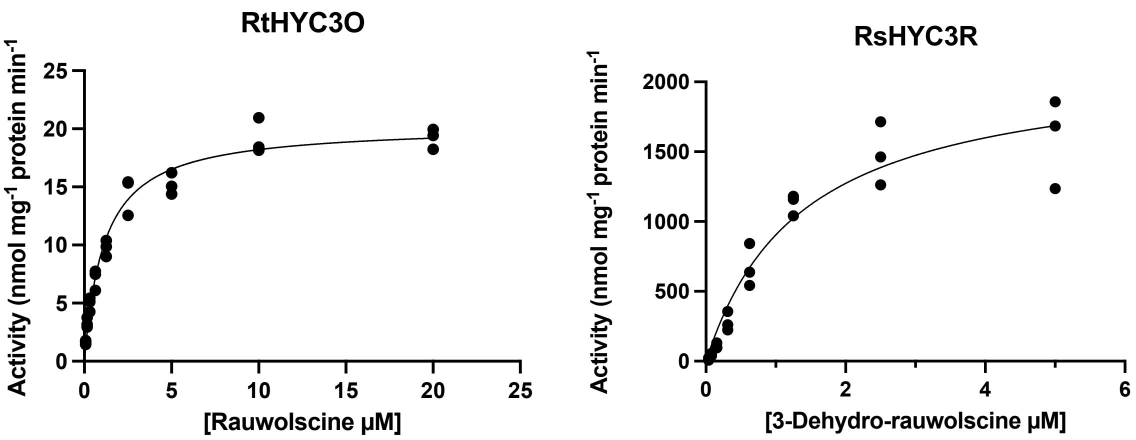

**Supplementary Figure 93. Saturation kinetics for RtHYC3O (left panel) and RsHYC3R (right panel).**

Both enzymes exhibited high binding affinity to their respective substrates. The $K_M$ value for RtHYC3O with rauwolscine was 1.15 µM (95% confidence interval [CI]: 0.927-1.42 µM), while the $K_M$ for RsHYC3R with 3-dehydro-rauwolscine was 1.38 µM (95% CI: 0.932-2.08 µM). Data was fitted using non-linear regression in Prism GraphPad 10.4.0. See Source data for the dataset used for kinetics calculation.

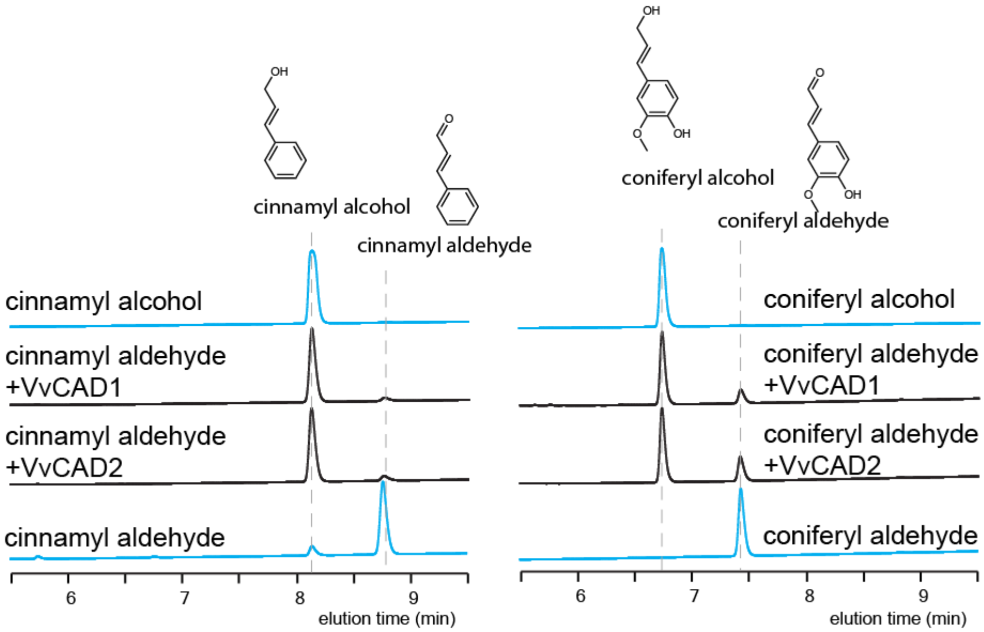

**Supplementary Figure 94.** ***Vitis vinifera*** **cinnamyl alcohol dehydrogenase (VvCAD1 and 2) catalyzes formation of cinnamyl and coniferyl alcohol from their aldehydes.**
Affinity purified his-tagged reductases from *E. coli* were used in in vitro assays with NADPH and monolignol aldehydes. The reduction data were obtained and visualized using liquid chromatography at 260 nm.

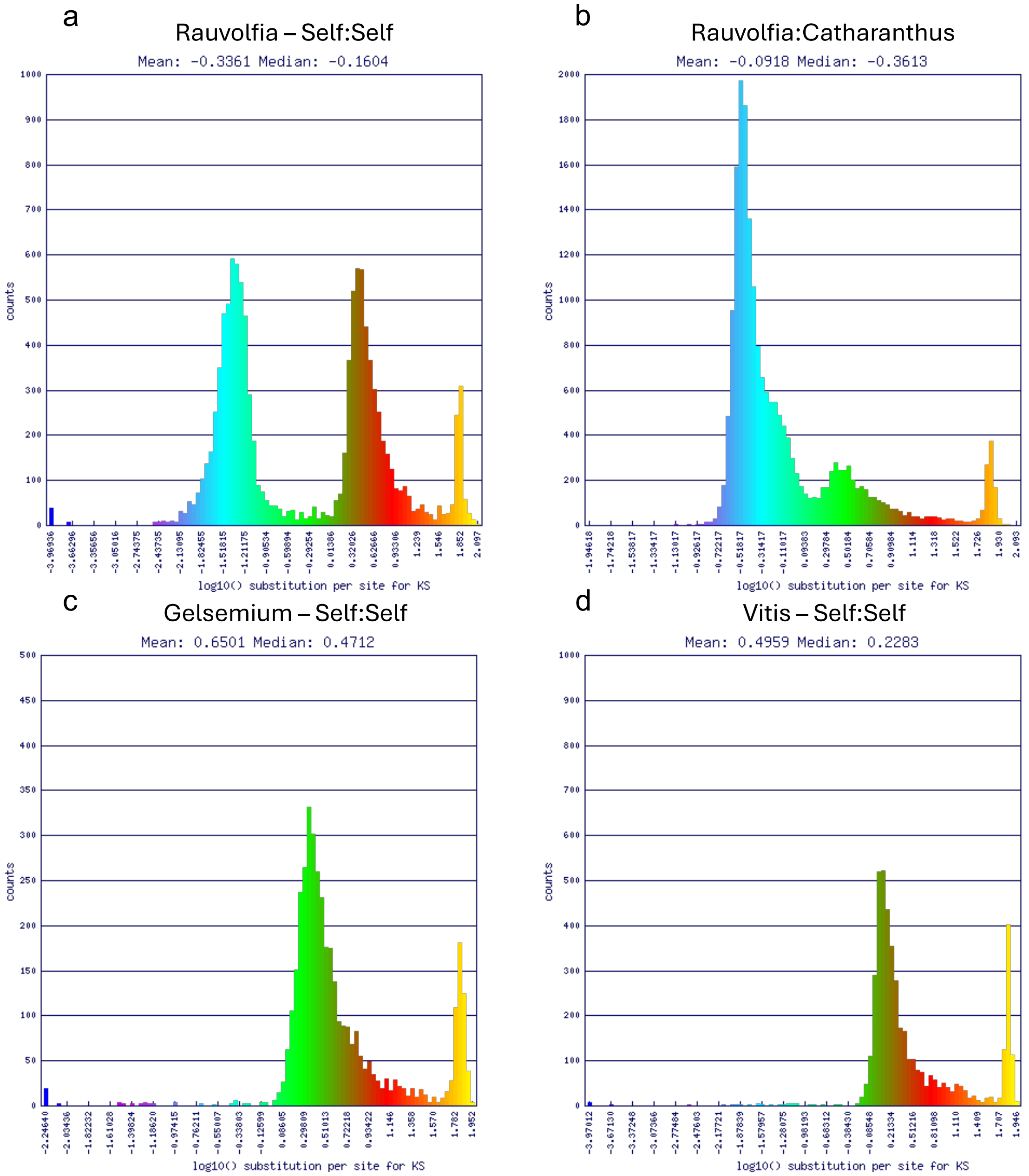

**Supplementary Figure 95. Ks histograms from CoGe SynMap analyses suggest a partially diploid nature of the published *Rauvolfia tetraphylla* genome assembly [1] .**
(**a**) self:self comparison of *Rauvolfia tetraphylla* reveal two Ks peaks: one at extremely low Ks (blue) and another at high Ks (red), with a similar number of gene pairs in both peaks. The high Ks peak corresponds to the ancient gamma hexaploidy event that occurred in the stem lineage of core eudicots. The peak is also observed at a comparable Ks value in three other analyses: (**b**) *Rauvolfia tatraphylla*:*Catharanthus roseus* (green), (**c**) *Gelsemium elegans*:*Gelsemium elegans* (green), and (**d**) *Vitis vinifera*:*Vitis vinifera* (green). If the low Ks peak in the *R. tetraphylla* self:self comparison had arisen from a recent polyploidy event, it would e expected to contain substantially more gene pairs than the ancient *gamma* peak, since fractionation would have heavily affected that ancient event. The extremely high Ks peaks (orange) observed in all panels represent typical background noise associated with Ks estimation in SynMap analyses.

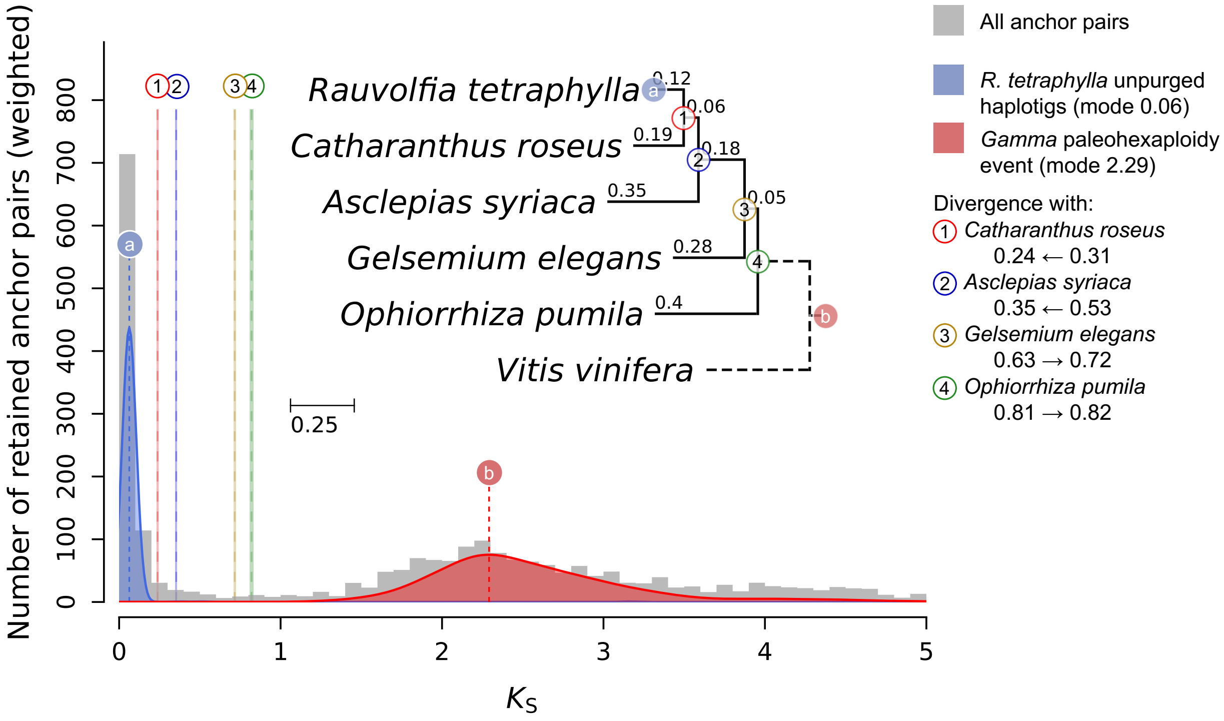

**Supplementary Fig. 96. A Ksrates analysis supports the conclusion that the *Rauvolfia tetraphylla* assembly [1] is partially diploid.**
Ksrates applies rate smoothing across a phylogenetic tree to place species divergence events relative to whole-genome duplications (WGDs), based on distributions of synonymous substitution rates (Ks). The analysis reveals two major Ks peaks: a very low Ks peak in *R. tetraphylla* (blue), potentially reflecting a recent duplication or assembly artifact, and a higher Ks peak (red) corresponding to the ancient gamma hexaploidy event at the base of all core eudicots.

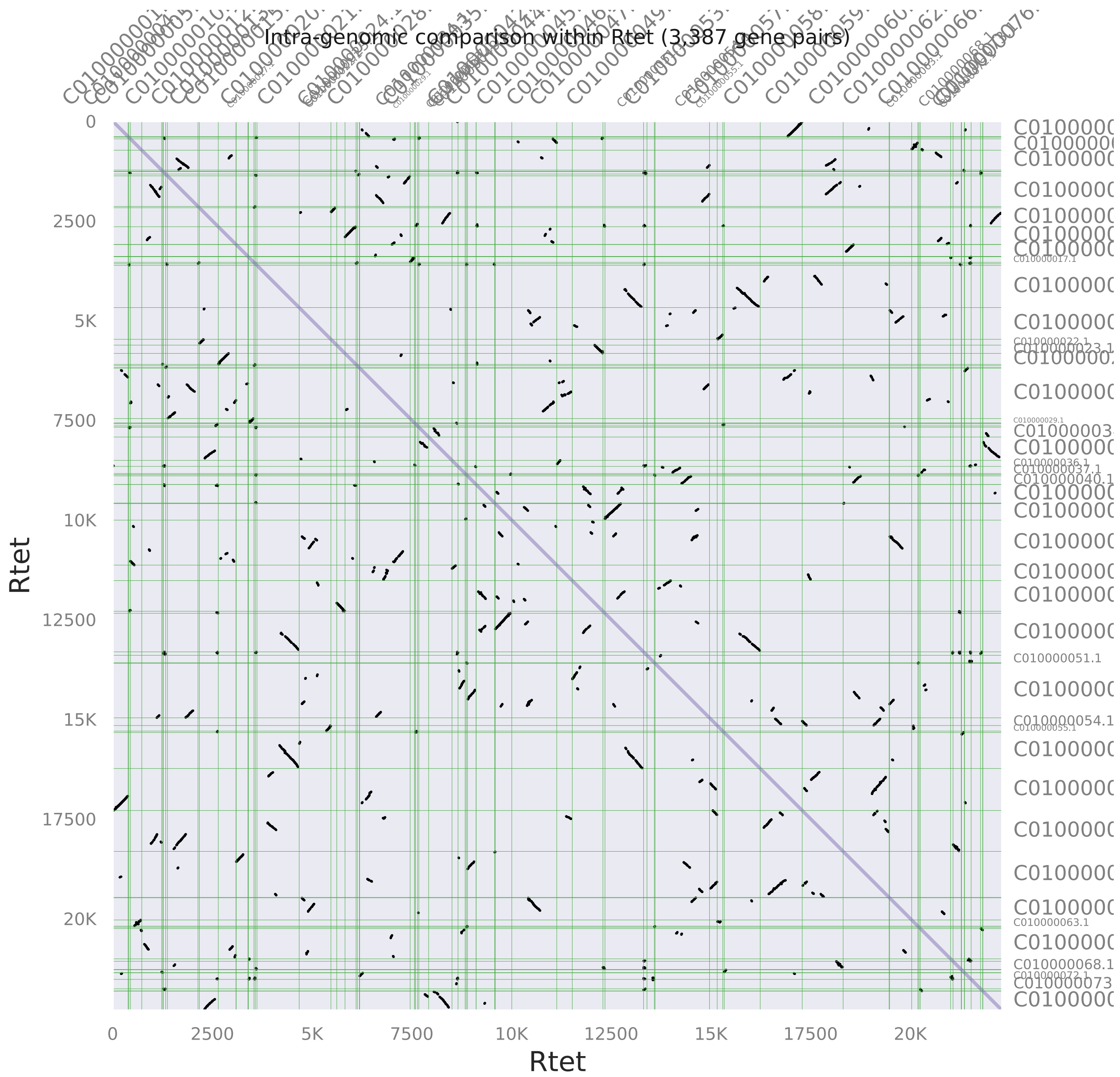

**Supplementary Figure 97. MCscan self:self syntenic dotplot *Rauvolfia* shows off-diagonal homologous blocks, indicating extended haplotypic regions in the published assembly [1] .**

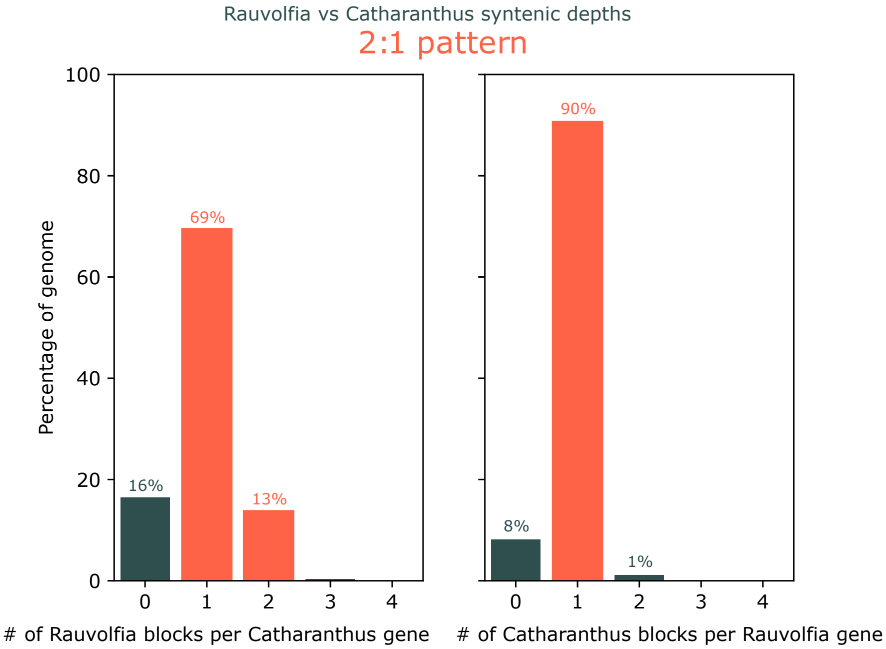

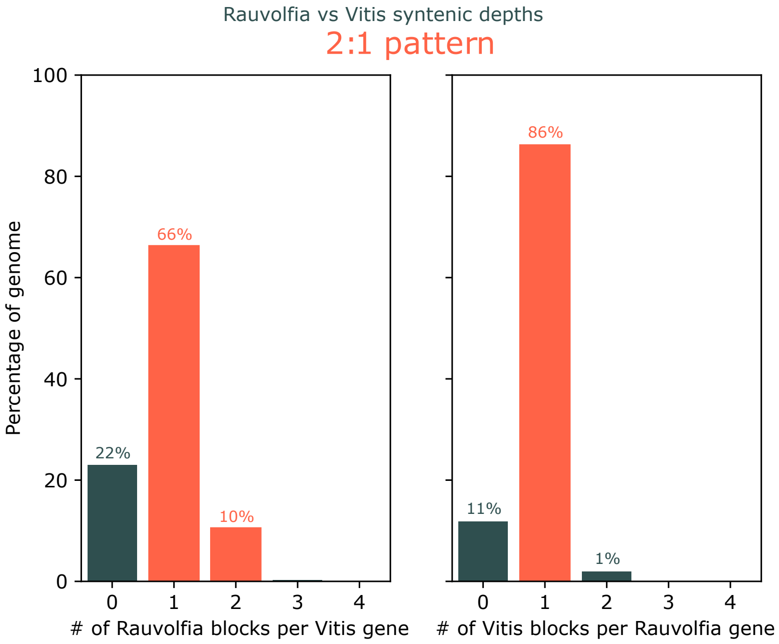

**Supplementary Figure 98. MCScan syntenic depth histograms of *Rauvolfia tetraphylla* against *Catharanthus roseus* and the reverse (top), and *Rauvolfia tetraphylla* against *Vitis vinifera* and the reverse (bottom), reveal some duplicated blocks in the published *Rauvolfia tetraphylla* genome [1].** Only 13% of the *Rauvolfia tetraphylla* genome is indicated as duplicated in comparison with *Catharanthus roseus*, and 10% against *Vitis vinifera*. These low proportions are insufficient to support the presence of a recent whole-genome duplication in *R. tetraphylla*, reinforcing conclusions from Ks-based and dotplot analyses that the genome is partially diploid.

**Supplementary table 1. $^{1}$H NMR chemical shifts of alkaloids in this study.**

| | Tetrahydro-alstonine | 3,14-dehydro-tetrahydro-alstonine | Akuammigine (3-epi-tetrahydroalstonine) | | | 3-epi-ajmalicine | | 3-dehydro-ajmalicine | pseudoyohimbine (3-epiyohimbine) | | 3-dehydro-yohimbine | 3-epi-rauwolscine | Rauwolscine | 3-dehydro-rauwolscine | alloyohimbine | |
|---|---|---|---|---|---|---|---|---|---|---|---|---|---|---|---|---|
| | This study<br>$CDCl_3$ | This study<br>$CDCl_3$ | This study 45°C<br>acetone-*d6* | This study 45°C<br>$CDCl_3$ | Ref[3] above RT<br>$CDCl_3$ | This study<br>$CDCl_3$ | Ref[4]<br>$CDCl_3$ | This study<br>MeOD | This study<br>$CDCl_3$ | Ref[5]<br>$CDCl_3$ | This study<br>MeOD | This study<br>$CDCl_3$ | This study<br>$CDCl_3$ | This study<br>MeOD | This study<br>acetone-*d6* | This study<br>$CDCl_3$ |
| NH | 7.77 s | 7.96 s | 9.74 s | 7.94 s | 7.84 s | 8.17 s | - | | 7.99 br s | 8.11 | - | 7.69 br s | 7.69 br s | - | 9.79 s | 7.78 s |
| 3 | 3.36 ddd | - | 3.56 br m | 3.60 br m | 3.8 | 4.55 qt | 4.55 m | - | 4.47 qt | 4.46 | - | 3.15 d | 3.15 d | - | 3.20 d | 3.25 br s |
| 5 | 2.56 m<br>2.96 ddd | 3.09 dd<br>3.07 ddd | 2.75 m<br>3.08 dd | 2.91 br s<br>3.12 dd | 2.8-3.2 m<br>(2H) | 3.32 dd | 3.30 m | 4.14 m<br>(2H) | 2.53 dd<br>3.29 m | 2.54<br>3.28 | 4.11 m<br>(2H) | 2.52 dd<br>2.96 dd | 2.52 dd<br>2.96 dd | 4.05 m<br>4.11 m | 2.47 dt<br>2.95 dd | 2.54 m<br>2.97 dd |
| 6 | 2.70 m<br>2.94 ddd | 2.87 ddd<br>2.95 dd | 2.59 dd<br>2.93 m | 2.65 dd<br>3.02 m | 2.8-3.2 m<br>(2H) | 2.55 dd<br>3.01 dddd | 2.55 m<br>3.00 m | 3.37 ddd<br>(2H) | 2.57 m<br>3.02 m | 2.60<br>3.02 | 3.34 m<br>(2H) | 2.68 m<br>2.95 m | 2.68 m<br>2.95 m | 3.32 | 2.64 dt<br>2.85 dddd | 2.68 br d<br>2.95 m |
| 9 | 7.45 d | 7.46 d | 7.39 d | 7.46 d | 7.46 dbr | 7.49 dd | 7.49 dd | 7.73 dt | 7.49 br d | 7.47 | 7.71 dt | 7.46 d | 7.46 d | 7.72 dt | 7.39 d | 7.46 d |
| 10 | 7.07 dt | 7.06 dt | 6.97 dt | 7.08 dt | 7.08 dd | 7.11 dt | 7.10 dt | 7.21 dt | 7.11 dt | 7.09 | 7.19 dt | 7.08 dt | 7.08 dt | 7.20 dt | 6.97 dt | 7.08 t |
| 11 | 7.13 dt | 7.15 dt | 7.03 dt | 7.14 dt | 7.14 dd | 7.17 dt | 7.17 dt | 7.46 dt | 7.17 dt | 7.15 | 7.43 dt | 7.13 dt | 7.13 dt | 7.45 dt | 7.05 dt | 7.13 t |
| 12 | 7.28 d | 7.27 d | 7.35 d | 7.35 d | 7.35 dbr | 7.40 dd | 7.40 dd | 7.52 dt | 7.41 dt | 7.4 | 7.49 dt | 7.31 d | 7.31 d | 7.52 dt | 7.29 d | 7.31 d |
| 14 | 2.49 dt<br>1.54 dd | 4.98 dd | 1.98 br s<br>2.74 br m | 2.11 br s<br>2.64 m | 2.14 m<br>2.8-3.2 m | 1.65 ddd<br>3.19 dt | 1.65 ddd<br>3.19 d | N.D.* | 1.65 m<br>2.18 m | 1.66<br>2.23 | N.D. * | 1.60 m<br>1.71 q | 1.60 m<br>1.71 q | N.D. * | 1.83 q<br>2.09 dt | 1.89 m<br>1.89 m |
| 15 | 2.77 m | 3.50 m | 2.75 br m | 2.72 br s | 2.75 m | 1.99 dddd | 2.00 m | 2.87 d | 1.62 m | 1.61 | 2.39 t | 2.44 ddd | 2.44 ddd | 2.28 m | 2.27 ddd | 2.55 m |
| 16 | - | - | - | - | - | - | - | - | 2.29 dd | 2.28 | 2.53 dd | 2.55 m | 2.55 m | 2.67 dd | 2.79 m | 2.84 br s |
| 17 | 7.56 s | 7.60 s | 7.47 s | 7.55 d | 7.54 s | 7.50 d | 7.50 d | 7.68 d | 4.22 qt | 4.19 | 4.39 qt | 4.00 dt | 4.00 dt | 4.10 m | 3.88 quin | 3.84 br s |
| 18 | 1.41 d | 1.45 d | 1.32 d | 1.35 e | 1.37 d | 0.92 d | 0.92 d | 1.28 d | 1.53 ddd<br>1.87 ddd | 1.54<br>1.84 | 1.76 ddd<br>1.97 ddd | 1.37 m<br>2.05 m | 1.37 m<br>2.05 m | 1.48 m<br>2.11 ddd | 1.73 m<br>2.00 m | 1.81 m<br>1.92 m |
| 19 | 4.50 dq | 4.18 dq | 4.43 quin | 4.44 quin | 4.44 qd | 4.34 dq | 4.33 dq | 4.65 ddd | 1.28 m<br>1.29 m | 1.25<br>1.30 | 1.62 m 2H | 1.56 m<br>2.11 ddd | 1.56 m<br>2.11 ddd | 1.72 m<br>1.72 m | 1.52 m<br>2.02 m | 1.56 m<br>2.09 br d |
| 20 | 1.70 m | 1.92 m | 1.93 br s | 1.85 br s | 1.86 m | 2.02 ddd | 2.00 m | 2.51 ddd | 1.41 m | 1.41 | 1.95 m | 1.82 m | 1.82 m | 2.78 ddd | 2.13 m | 1.85 m |
| 21 | 2.74 m<br>3.11 dd | 3.22 d | 2.53 ddd<br>2.93 m | 2.59 dd<br>2.97 dd | 2.5-2.7 m<br>2.8-3.2 m | 2.56 t<br>2.63 dt | 2.50t<br>2.63 t | 3.84 br d<br>3.98 dd | 2.56 m<br>3.29 m | 2.55<br>3.26 | 3.66 dd<br>3.90 dd | 2.60 dd<br>2.85 dd | 2.60 dd<br>2.85 dd | 3.76 d<br>4.16 br d | 2.53 dd<br>2.82 m | 2.56 m<br>2.81 br d |
| $OCH_3$ | 3.75 s | 3.78 s | 3.69 s | 3.75 s | 3.75 s | 3.74 s | 3.73 s | 3.78 s | 3.58 s | 3.79 | 3.78 s | 3.84 s | 3.84 s | 3.80 s | 3.65 s | 3.77 s |

* Not detected. The H14 signals were not detected in 3-dehydro-ajmalicine, -yohimbine, or -rauwolscine in MeOD, likely due to equilibrium between the 3-dehydro (H14 as methylene) and 3,14-dehydro (H14 as alkene) forms, resulting exchange-mediated loss in the NMR spectrum.

**Supplementary table 2. $^{13}$C NMR chemical shifts of alkaloids in this study.**

| | Tetrahydro-alstonine | 3,14-dehydro-tetrahydro-alstonine | akuammigine (3-epi-tetrahydroalstonine) | | | 3-epi-ajmalicine | | 3-dehydro-ajmalicine | 3-epi-yohimbine (pseudoyohimbine) | | 3-dehydro-yohimbine | 3-epi-rauwolscine | | Rauwol-scine | 3-dehydro-rauwol-scine | alloyohimbine | | |
|---|---|---|---|---|---|---|---|---|---|---|---|---|---|---|---|---|---|---|
| | This study CDCl₃ | This study CDCl₃ | This study 45°C acetone-*d6* | This study 45°C CDCl₃ | Ref [6] above RT CDCl₃ | This study CDCl₃ | Ref [4] CDCl₃ | This study MeOD | This study CDCl₃ | Ref [5] CDCl₃ | This study MeOD | This study CDCl₃ | Ref [7] CDCl₃ | This study CDCl₃ | This study MeOD | This study acetone-*d6* | This study CDCl₃ | Ref [6] CDCl₃ |
| 2 | 134.68 | 129.9 | 135.4 | 133.6 | 132.8 | 132.8 | 132.50 | 127.77 | 133.2 | 123.3 | 127.52 | 132.2 | 132 | 134.6 | 127.73 | - | 134.7 | 134.4 |
| 3 | 59.98 | 134.7 | 55.98 | 54.81 | 54.5 | 54.26 | 54.10 | N.D.* | 54.29 | 54.3 | 166.95 | 54.04 | 53.8 | 60.32 | 166.14 | 60.84 | 60.54 | 60.1 |
| 5 | 53.72 | 51.41 | 53.55 | 52.54 | 52.2 | 51.07 | 50.90 | 54.16 | 51.19 | 51 | 53.82 | 51.22 | 51 | 53.34 | 54.23 | 53.19 | 53.29 | 52.8 |
| 6 | 21.92 | 21.53 | 20.84 | 19.49 | 19.2 | 16.98 | 16.80 | 20.27 | 17.1 | 16.9 | 20.19 | 16.88 | 16.6 | 21.75 | 20.18 | 21.69 | 21.8 | 21.3 |
| 7 | 108.27 | 110.7 | 108.3 | 108.2 | 106.8 | 106.9 | 106.70 | 124.20 | 108 | 107.2 | 123.58 | 108.5 | 108 | 108.6 | 123.94 | - | 108.7 | 107.1 |
| 8 | 127.32 | 127.1 | 128.6 | 127.9 | 127.2 | 127.9 | 127.70 | 125.48 | 128 | 127.5 | 125.47 | 127.9 | 127.6 | 127.5 | 125.46 | - | 127.5 | 126.8 |
| 9 | 118.21 | 118.6 | 118.3 | 118.1 | 117.7 | 118.1 | 117.90 | 122.93 | 118 | 117.9 | 122.49 | 118.2 | 118 | 118.3 | 122.52 | 117.6 | 118.3 | 117.5 |
| 10 | 119.56 | 119.7 | 119.6 | 119.6 | 119.1 | 119.5 | 119.30 | 122.59 | 119.5 | 119.2 | 122.76 | 119.7 | 119.6 | 119.6 | 122.85 | 118.6 | 119.7 | 118.6 |
| 11 | 121.56 | 122.8 | 121.5 | 121.6 | 121.2 | 121.6 | 121.40 | 129.96 | 121.5 | 121.4 | 129.66 | 121.7 | 121.7 | 121.6 | 129.82 | 120.7 | 121.6 | 120.5 |
| 12 | 110.93 | 110.9 | 112 | 111.2 | 110.8 | 111.4 | 111.20 | 114.21 | 111.5 | 111.4 | 114.13 | 111 | 110.9 | 110.8 | 114.2 | 110.9 | 110.9 | 110.6 |
| 13 | 136.14 | 136.9 | 137.6 | 136.2 | 135.7 | 136 | 135.80 | 142.98 | 136.1 | 136.1 | 142.73 | 135.8 | 135.7 | 136.1 | 142.89 | - | 136.2 | 135.8 |
| 14 | 34.41 | 96.39 | 31.85 | 31.1 | 30.6 | 31.37 | 31.20 | N.D. * | 32.6 | 32.1 | N.D. * | 23.88 | 23.6 | 27.84 | N.D. * | 31.41 | 31.38 | 31 |
| 15 | 31.52 | 30.29 | 25.95 | 26.2 | 25.7 | 26.27 | 26.00 | 24.75 | 31.62 | 31.3 | 29.88 | 32.67 | 32.2 | 38.04 | 34.34 | 38.25 | 38.17 | 37.4 |
| 16 | 109.62 | 109 | 108.3 | 107.2 | 107.6 | 107.8 | 107.60 | 106.63 | 52.2 | 52.1 | 53.02 | 54.14 | 54 | 54.83 | 54.13 | 51.54 | 50.92 | 50.6 |
| 17 | 155.87 | 155.5 | 155 | 155.3 | 154.8 | 154.8 | 154.60 | 156.47 | 67.47 | 67.2 | 68.12 | 66.09 | 65.8 | 66.13 | 66.37 | 66.38 | 67.4 | 66.7 |
| 18 | 18.67 | 18.44 | 18.75 | 18.74 | 18.4 | 15.2 | 15.00 | 14.12 | 31.77 | 31.6 | 32.56 | 33.45 | 33.2 | 33.2 | 34.67 | 30.78 | 31.79 | 30.2 |
| 19 | 72.62 | 72.99 | 74.7 | 73.59 | 73.2 | 74.02 | 73.80 | 72.86 | 23.16 | 23.1 | 22.94 | 24.41 | 24.1 | 24.63 | 24.87 | 25.11 | 25.35 | 24.8 |
| 20 | 38.60 | 36.7 | 38.41 | 37.8 | 37.2 | 41.54 | 41.30 | 37.91 | 40.39 | 39.9 | 36.13 | 36.06 | 35.6 | 36.69 | 31.62 | 32.32 | 32.85 | 32 |
| 21 | 56.48 | 51.7 | N.D. | N.D. | 50.3 | 47.56 | 47.30 | 54.99 | 51.45 | 51 | 58.62 | 49.86 | 49.5 | 60.64 | 58.42 | 60.5 | 60.35 | 59.6 |
| $CH_3$-O-**<u>CO</u>** | 168.12 | 168.4 | 167.9 | 168 | 167.5 | 167.5 | 167.30 | 179.41 | 175.1 | 175.4 | 174.23 | 174.9 | 173.8 | 174.8 | 174.55 | - | 174.6 | 174 |
| **<u>$CH_3$</u>**-O-CO | 51.26 | 51.33 | 51.13 | 51.18 | 50.9 | 51.19 | 50.90 | 51.81 | 52.04 | 51.9 | 52.41 | 52.08 | 52 | 52.04 | 52.47 | 50.57 | 52.05 | 51.5 |

* Not detected. The C14 signals were not detected in 3-dehydro-ajmalicine, -yohimbine, or -rauwolscine in MeOD, likely due to equilibrium between the 3-dehydro (C14 as methylene) and 3,14-dehydro (C14 as alkene) forms, resulting exchange-mediated loss in the NMR spectrum. While the C3 signals were identified for 3-dehydro-yohimbine and -rauwolscine, the C3 signal for 3-dehydro-ajmalicine was not detected, likely due to exchange effects.

**Supplementary table 3. Primers used in this study**

| # | primers | sequence 5'-3' |
|---|---|---|
| 1 | RsHYC3R-BamHI-F | ATAGGATCCGATGGCAGCAGCAGAAAC |
| 2 | RsHYC3R-SalI-R | GAAGTCGACTTGAAATGCAGCATCTATGC |
| 3 | RsYOS-BamHI-F | ATAGGATCCAATGGCTTCAGAGTCGCCGGA |
| 4 | RsYOS-SalI-R | GAAGTCGACTTATGCCGATTTGAGAGTGTTTC |
| 5 | CrHYS-BamHI-F | TTCGGATCCAATGGCTGAGGGATTAATGGCT |
| 6 | CrHYS-SalI-R | TTCGTCGACTTAAAGCGATTTGAGAGTGTTTCC |
| 7 | CrDCS-BamHI-F | AACGGATCCTATGGCAATGGCTTCAAAGTCAC |
| 8 | CrDCS-SalI-R | CAAGTCGACTTAATTTGATTTCAGAGTGTTCCCT |
| 9 | CrTHAS2-BamHI-F | CGAGGATCCAATGTCTTCAAAATCAGCAAAACC |
| 10 | CrTHAS2-SalI-R | GGTGTCGACCTAAGCAGATTTCAATGTGTTTTC |
| 11 | CrHYC3R-BamHI-F | ATAGGATCCAATGGCAGTTCCATCGGCAGAA |
| 12 | CrHYC3R-SalI-R | GAAGTCGACTAACTCTAAACAGATCCCAAAGA |
| 13 | RtHYC3O-BamHI-F | gtcaaggagaaaaaaccccggatccATGGAAACAGAAGTCCGTATGGT |
| 14 | RtHYC3O-His-R | aaatcaacttctgttccatgtCTAATGATGATGATGATGGTGCACAGATGCAAGAATGTATAGAG |
